# Enhancing biocompatibility of rhodamine fluorescent probes by a neighbouring group effect

**DOI:** 10.1101/2020.03.11.986919

**Authors:** Jonas Bucevičius, Georgij Kostiuk, Rūta Gerasimaitė, Tanja Gilat, Gražvydas Lukinavičius

**Affiliations:** Chromatin Labeling and Imaging Group, Department of NanoBiophotonics, Max Planck Institute for Biophysical Chemistry, Am Fassberg 11, 37077 Göttingen, Germany

## Abstract

Fluorescence microscopy is an essential tool for understanding dynamic processes in living cells and organisms. However, many fluorescent probes for labelling cellular structures suffer from unspecific interactions and low cell permeability. Herein, we demonstrate that the neighbouring group effect which results from positioning an amide group next to a carboxyl group in the benzene ring of rhodamines dramatically increases cell permeability of the rhodamine-based probes through stabilizing a fluorophore in a hydrophobic spirolactone state. Based on this principle, we create probes targeting tubulin, actin and DNA. Their superb staining intensity, tuned toxicity and specificity allows long-term 3D confocal and STED nanoscopy with sub-30 nm resolution. As a result, the real microtubule diameter of 23 nm was resolved inside a living cell for the first time. Due to their unrestricted cell permeability and efficient accumulation on tubulin, the new probes produce high contrast images at sub-nanomolar concentrations.

## Introduction

Modern super-resolution fluorescence microscopy and nanoscopy techniques permit observation of biological processes in living organisms down to the molecular level ^1, 2, 3^. However, their success depends on the availability of biocompatible fluorescent probes for specific labelling of cellular structures. Usually, these probes are constructed by coupling a fluorescent dye to a small molecule ligand via a linker to target a protein of interest. The linker and the fluorophore make significant contributions to the final properties of such probes, often resulting in compromised cell permeability and off-targeting ^4, 5^. Recent studies dedicate a lot of attention to the rhodamine class fluorophores that can cycle between a non-fluorescent spirolactone and a fluorescent zwitterion form. This property is exploited to generate fluorogenic probes, which are sensitive to a number of environmental factors: pH, ion concentration, enzymatic activity and its local microenvironment polarity or light ^6, 7^. Since the spirolactone is a more hydrophobic molecule compared to the zwitterion, its presence makes rhodamine-based fluorescent probes more cell-permeable ^8, 9^. Regular rhodamines and carbon-substituted analogues (carbopyronines) have the equilibrium shifted towards zwitterion form, which results in relatively poor membrane permeability ^8, 9^. To overcome this issue, several studies attempted to induce spirolactone form preference by introducing electron-withdrawing groups into the xanthene core ^10, 11, 12^ or in the benzoic acid substituent ^13^. However, these approaches result in a bulkier core structure and alter the physicochemical properties of the dyes. We propose a way for modulating the spirolactone-zwitterion equilibrium without any change in the fluorescent dye core structure by exploiting the neighbouring group effect (NGE), i.e. the phenomenon that two neighbouring carboxylic groups can influence each other via steric, electrostatic or H-bond interactions ^14, 15^..

Here, we introduce a new positional isomer with NGE to increase the cell permeability and fluorogenicity of regular rhodamines without changing their spectroscopic properties. We demonstrate that excellent quality of wash-free multi-colour live-cell images can be obtained at low concentrations of probes with wide-field, confocal and super-resolution microscopes.

## Results

### Chemical synthesis of 4′-carboxyl rhodamines

For a long time, the existence of a novel isomeric class of 4’-carboxyrhodamines was debated due to the synthesis challenge arising from steric hindrance and the *ortho*-effect altered reactivity of adjacent carbonyl groups ^16^. Isomeric mixtures of 5/6-carboxyrhodamines and fluoresceins are usually obtained by the classical condensation process of phthalic anhydrides and 3-dimethylaminophenol or resorcinol respectively ^17, 18^. But condensation between resorcinol and 1,2,3-benzenetricarboxylic acid or its homologs yields solely 7’-carboxyfluorescein and not 4’-carboxyfluorescein ^19^. Another widely applied and regioselective synthesis strategy, which can be applied in the synthesis of 4’-carboxyrhodamines, involves addition of aryl-lithium species to the 3,6-substituted-xanthone ^20^.We used formation of di-*tert*-butyl ester **1** for the protection of 3-bromophtalic acid carboxyl groups for the halogen-lithium exchange reaction (Fig. 1a). However, the *tert*-butyl protecting group has low tolerance to strong nucleophiles and the lithium-halogen exchange must be carried out at lower than usual temperatures, between −90 and - 120 °C. Such a temperatures are not compatible with the reactivity of xanthones and their C/Si analogues possessing strong electron donating dialkyl amino groups at positions 3 and 6. When the halogen exchange was attempted at −78 °C, mainly homo-addition products were formed. To solve this issue, we performed the reaction at ∼-116 °C with the more reactive silyl protected 3,6-dihydroxyxanthone **2** and its C (**3**) and Si (**4**) analogues, and in the later stage converted the resulting fluoresceins to rhodamines using the strategy developed by J. B. Grimm *et al* ^21^. To achieve this, TBDMS protecting groups on compounds **5-7** were removed with TBAF in THF and the obtained fluorescein class compounds **8-10** were reacted with triflic anhydride in the presence of pyridine in DCM to obtain the triflates **11-13**. Finally, the triflate groups were converted to the desired (di)methylamino groups (compounds **14-16**) by a Pd-catalyzed C-N cross coupling reaction and the desired 4’-carboxyrhodmines were obtained by quantative deprotection with a 1:1 mixture of DCM/TFA (Fig. 1a).

**Fig. 1.**
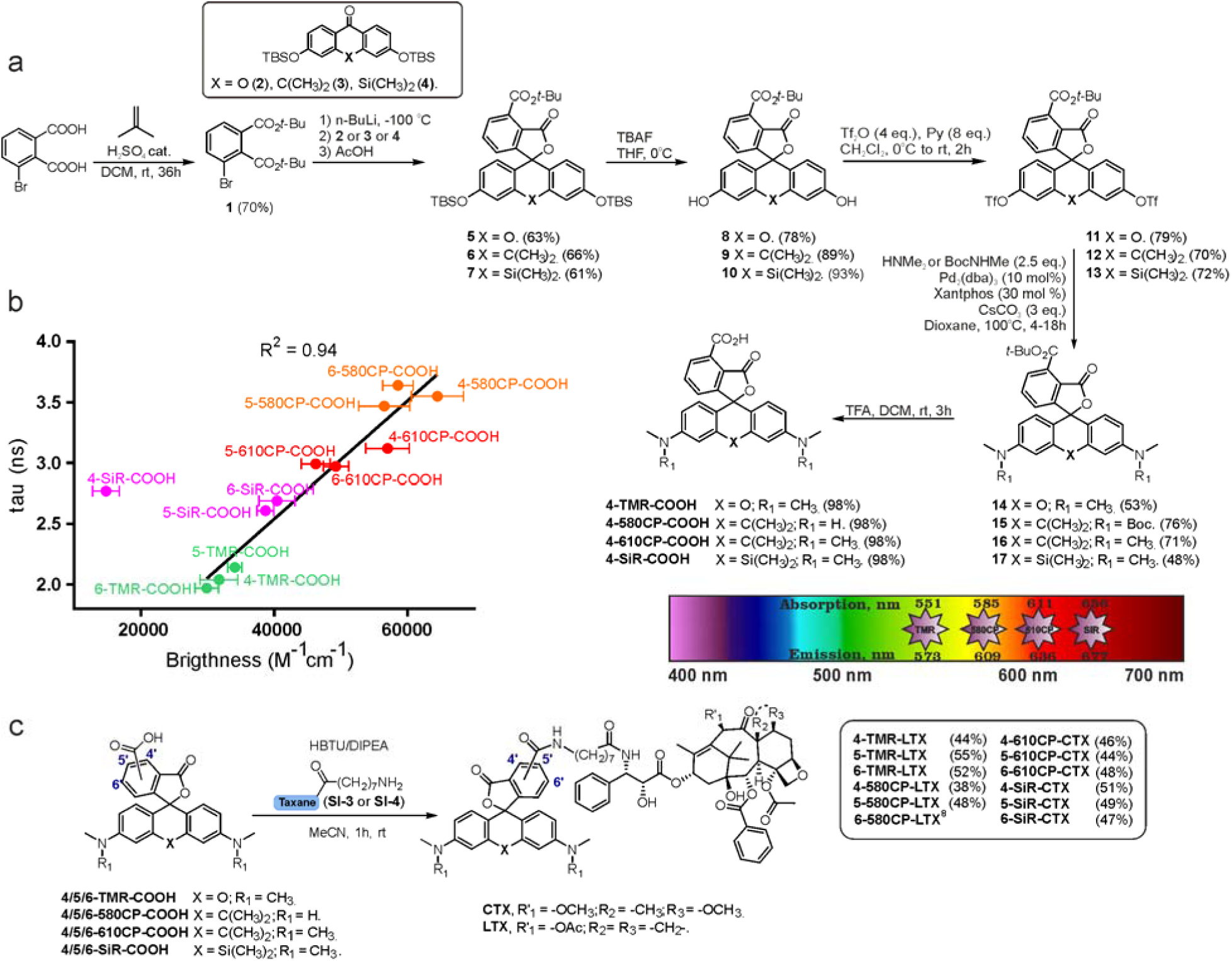
Synthesis of rhodamine 4’-isomers, probes and their photophysical properties. **a**, Synthesis scheme of **4-TMR-COOH, 4-580CP-COOH, 4-610CP-COOH** and **4-SiR-COOH** dyes. **b**, Correlation between brightness and fluorescence lifetime of 4’/5’/6’-carboxyrhodamine dyes. **c**, Synthesis scheme for fluorescent probes based on taxane ligands for tubulin imaging.

### In vitro characterization of neighbouring group effect in 4′-carboxyrhodamines

4’-, 5’- and 6’-carboxyrhodamine regioisomers have almost identical absorption, fluorescence emission spectra and quantum yield (QY) in phosphate buffered saline (PBS) (Supplementary Table 1). In agreement with previously published observations, we obtained an excellent correlation between QY and fluorescence lifetime (Supplementary Fig. 1) ^22^. Even better correlation was obtained between fluorophore brightness (product of QY and extinction coefficient) and fluorescence lifetime with the exception of the **4-SiR-COOH** dye, which showed a significantly lower extinction coefficient (Fig. 1b).

To characterize the ability of the fluorophores to switch between spirolactone and zwitterion states, we measured the absorbance in water-1,4-dioxane mixtures with known dielectric constant ^23^.We fitted the absorbance change to the dose response equation and obtained the ^dye^D_50_ value, which indicates the dielectric constant at which absorbance is halved (Supplementary Table 2 and Supplementary Fig. 2).^Dye^ D_50_ of **4-SiR-COOH** was significantly higher than the ^dye^D_50_ of 5’- and 6’-isomers.For all other fluorophores, the ^dye^D_50_ of all three isomers were very close.

**Fig. 2.**
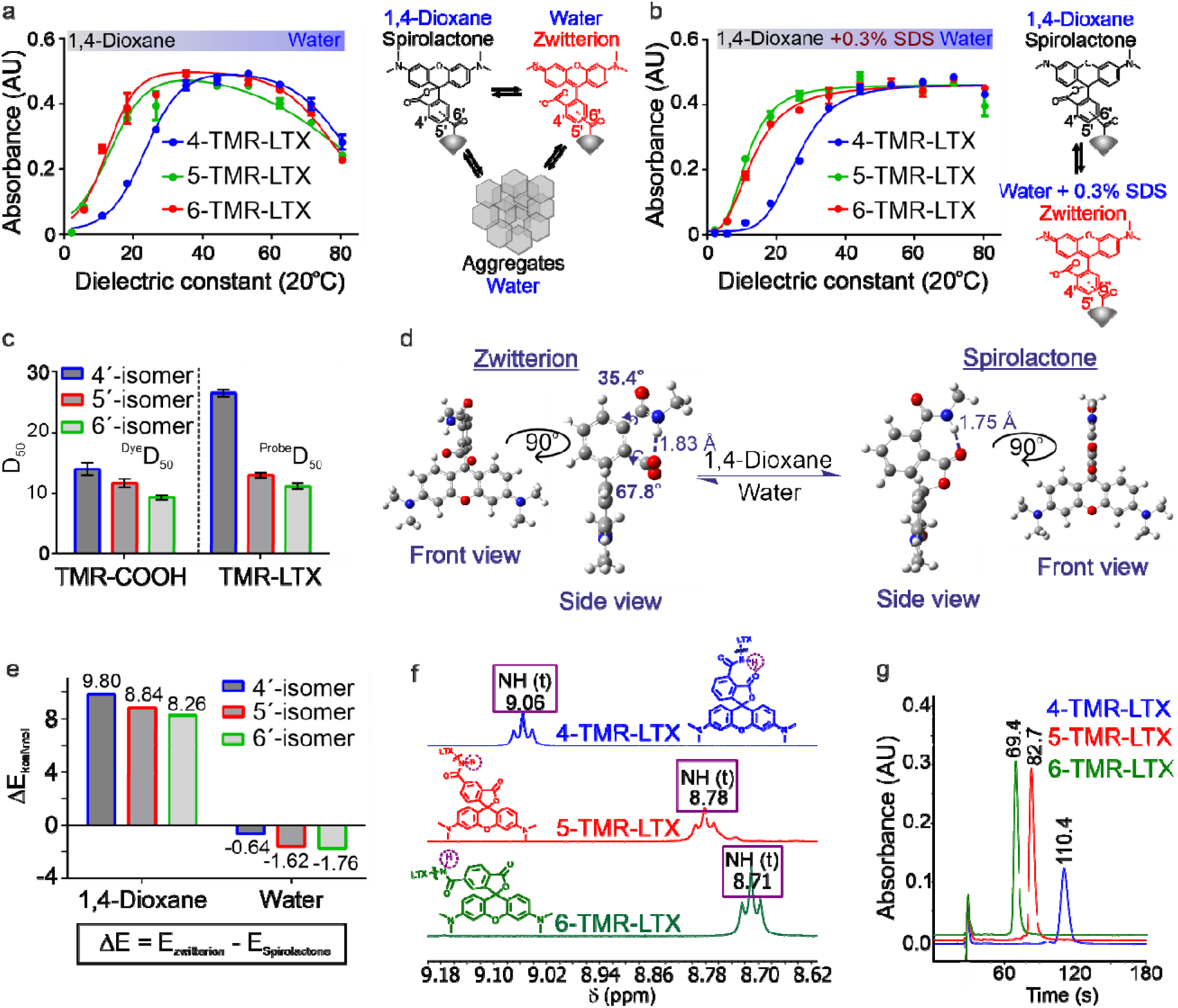
Neighbouring group effect in the fluorescent probes. **a**, Plot showing absorbance of **TMR-LTX** positional isomers at λ_max_ versus dielectric constant of **1**,4-dioxane-water mixtures. Data points were fitted to bell-shaped dose response curve, which describes two consecutive processes spirolactonization and aggregation. Data presented as mean ± s.d., N = 3 independent experiments each n = 3. **b**, Plot showing absorbance of **TMR-LTX** positional isomers at λ_max_ versus dielectric constan of 1,4-dioxane-water mixtures with constant 0.3% SDS additive. D_50_ value was obtained by fitting to EC_50_ dose-response. Data presented as mean ± s.d., N = 3 independent experiments, each n = 3. **c**, ^Dye^D_50_ values of positional isomers of **TMR-COOH** and ^probe^D_50_ of **TMR-LTX. d**, DFT optimized geometries of model 4’-regioisomer fluorescent probe in spirolactone and zwitterion forms with truncated linker and targeting moiety. **e**, DFT calculated potential energy differences between spirolactone and zwitterion of model 4’/5’/6’-regiosomer probes in 1,4-dioxane and water. **f**, Chemical shifts of the amide proton of **TMR-LTX** regioisomeric probes**. g**, Comparison of retention times of **TMR-LTX** regioisomeric probes in HPLC analysis with SB-C18 column under isocratic elution conditions (75:25 MeOH : H_2_O 25m HCOONH_4_ pH = 3.6).

We have previously demonstrated that the performance of the tubulin probes depends on the chosen taxane ligand, and that the optimal ligand is different for each fluorophore ^8^. Based on this knowledge, we synthesized a new series of tubulin probes by coupling the 4’-regioisomers of **TMR-COOH** and **580CP-COOH** fluorophores to a Larotaxel derivative (**LTX-C8-NH_2_**, (**SI-4**)) and **610CP-COOH** and **SiR-COOH** to a Cabazitaxel derivative (**CTX-C8-NH**_2_, (**SI-3**)) (Fig. 1c). 1,4-Dioxane-water titration of **TMR-LTX** probes yielded bell-shaped dose response curves, indicating two transitions corresponding to zwitterion formation followed by aggregation (Fig. 2a). These processes are described by ^probe^ D_50_ and ^probe^A_50_ parameters indicating dielectric constant at which absorbance is halved, respectively (Supplementary Table 3). If the same titration is performed keeping the SDS concentration constant at 0.3%, the aggregates cannot form and the data can be fitted to a simple EC_50_ equation (Fig. 2b, Supplementary Fig. 3 and Supplementary Table 3).

**Fig. 3.**
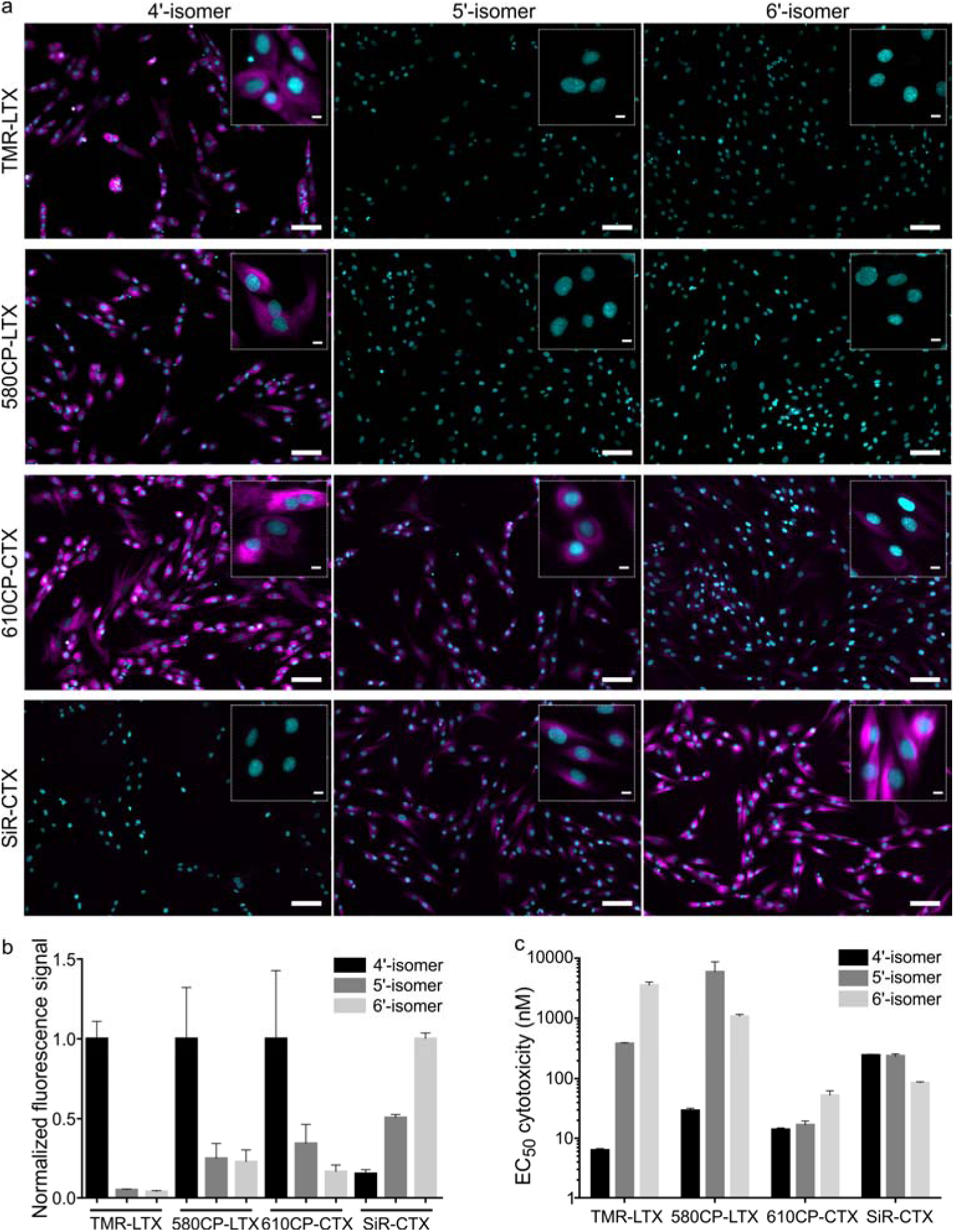
Performance of tubulin fluorescent probes based on 4’-isomers of rhodamines. **a**, Wide-field fluorescence microscopy of living primary fibroblasts stained with 100 nM **TMR-LTX** isomers for 1h at 37°C. Cells were washed once with HBSS and imaged in DMEM growth media. Insets shows zoomed-in images. Scale bars: 100 µm (large field of view), 10 µm (inset). Hoechst staining is shown in cyan and all tubulin probes are in magenta**. b**, Quantification of fluorescence signal in the cytoplasm of living cells stained with tubulin probes. Data is presented as mean ± s.e.m., N = 3 independent experiments, each time n > 100 cells were quantified. **c**, Cytotoxicity of tubulin fluorescent probes presented as half maximal effective concentration (EC_50_) after 24h incubation at 37°C in growth media. Cytotoxicity was determined as fraction of cells containing less than a single set of genetic material (sub G1 DNA content). Data is presented as mean ± s.e.m., N = 3 independent experiments, each time n > 100 cells were quantified.

For 5’- and 6’-regioisomers, ^probe^ D_50_ were only marginally higher than the ^dye^ D_50_ of corresponding fluorophores. ^probe^ D_50_ of 4’-regioisomers was significantly increased compared to the corresponding ^dye^ D_50_ (Supplementary Tables 2 and 3). This indicates that the conversion of carboxyl group to amide induces higher spirolactone content at equilibrium and thus implies higher hydrophobicity of 4’-probes (Fig. 2c).

In order to better understand how NGE alters the spirolactone-zwitterion equilibrium, we modelled *in silico* 4’/5’/6’-carboxamidetetramethylrhodamine methylamides (**4-TMR-NHMe, 5-TMR-NHMe** and **6-TMR-NHMe**) with truncated ligand and linker. Geometrical structure optimisations pointed out unique properties of the 4’-isomer: the spirolactone form can form an intramolecular H-bond since the distance between the carboxamide proton and carbonyl is only 1.75Å, the carboxylate is 68°twisted with respect to the benzene ring because of steric hindrance in the zwitterion form (Fig. 2d and Supplementary Fig. 4). Comparison of calculated total potential energies of the molecules in the modelled solvent environment representing two uttermost points in 1,4-dioxane (ε_r_ = 2.21) and water (ε_r_ = 78.35) indicate that the spirolactone form of **4-TMR-NHMe** in 1,4-dioxane is stabilised and the zwitterion form in water is destabilised in comparison with other regioisomers (Fig. 2e). As the stabilisation of spirolactone form could be attributed to the formation of intramolecular hydrogen bond, the stabilising effect could be considered diminishing upon increase of water content. Therefore the observed shift of D_50_ value in 4’-carboxamide isomers is mostly arising from the destabilisation of the zwitterion form in water environment (Supplementary Table 4).

**Fig. 4.**
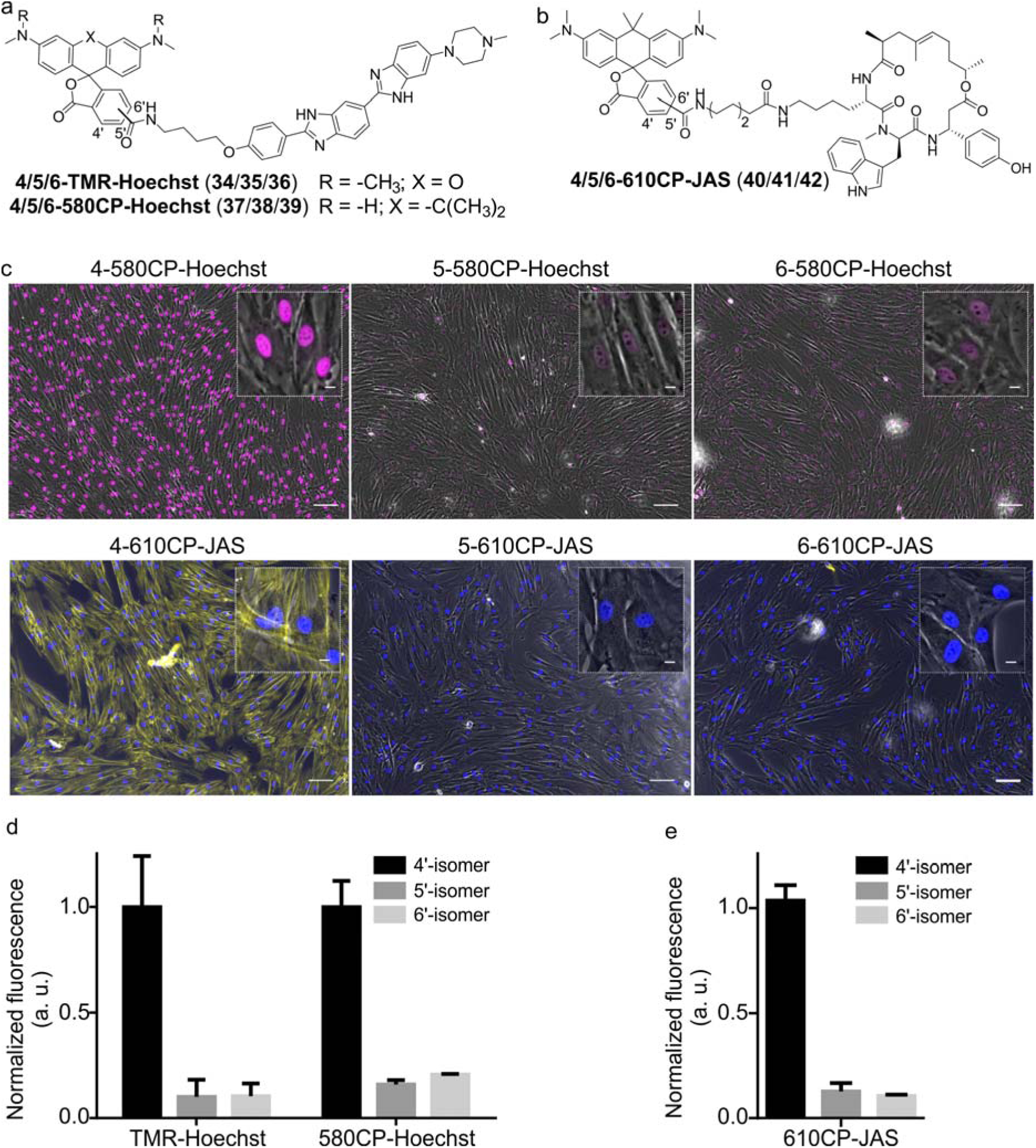
Performance of DNA and actin fluorescent probes based on 4’-isomer of rhodamines. **a**, Structure of DNA probes showing positional isomerism of the attachment point. **b**, Structure of actin probes and positional isomerism of the attachment point. **c**, Wide-field microscopy images of living primary fibroblasts. The cells were stained with 100 nM **4/5/6-580CP-Hoechst** (magenta) or 100 nM **4/5/6-610CP-JAS** (yellow) for 1h at 37°C, washed once with HBSS and imaged in DMEM media. Inserts show zoomed-in images. Scale bars: 100 µm (large field of view), 10 µm (insets). Overlay with light transmission (grey) is shown. **d**, Quantification of DNA probes’ fluorescence signal in the nuclei. Data presented as mean ± s.d., N = 3 independent experiments, each n > 100 cells. **f**, Quantification of **4/5/6-610CP-JAS** fluorescence signal in the cytoplasm of living cells. Data presented as mean ± s.d., N = 3 independent experiments, each n > 100 cells.

NMR spectroscopy confirmed the presence of the NGE: the amide proton in the 4’ –isomer probes is deshielded and downfield shifted by 0.3 - 0.4 ppm compared to 5’- and 6’-regioisomers. This shift does not depend on the xanthene ring system or on the attached targeting ligand (Fig. 2f, Supplementary Table 5).

The higher ^probe^D_50_ values of 4’-regioisomers suggest a higher percentage of the hydrophobic spirolactone form under equilibrium conditions which is retained longer in the reverse phase C_18_ HPLC column because of the stronger interaction with resin ^25^. We compared retention times of the probes under isocratic elution conditions and observed a high correlation between ^probe^D_50_ and the retention times (Supplementary Fig. 5). The 4’-regioisomer derivatives displayed significantly longer elution times compared to 5’- and 6’-regioisomer based probes (Fig. 2g and Supplementary Table 6).

**Fig. 5.**
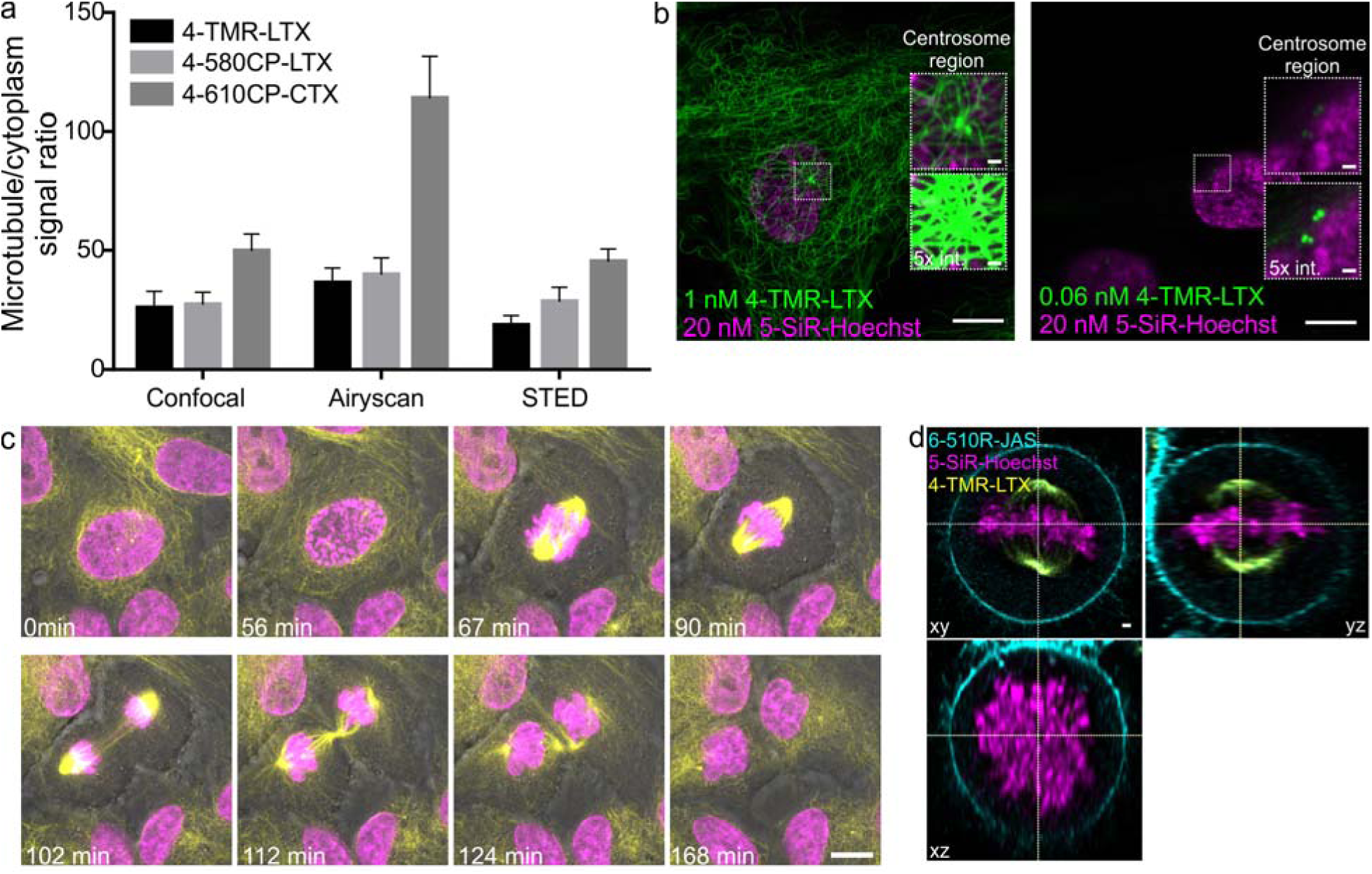
Confocal and Airyscan imaging of living cells stained with rhodamine 4’-isomer probes. **a**, Single microtubule and cytoplasm fluorescence signal ratio in living human fibroblasts stained with 100 nM of indicated probe for 1h at 37°C and imaged with the indicated microscopy method without probe removal. Data is presented as mean ± s.d., N ≥ 3 independent fields of view, each time n ≥ 20 microtubules. **b**, Zeiss Airyscan images of human primary fibroblasts stained with **5-SiR-Hoechst** and **4-TMR-LTX** for 24 h at 37°C in growth media at indicated concentrations. Images acquired without probe removal. **c**, Cell cycle of human primary fibroblasts stained with 1 nM **4-TMR-LTX** (yellow) and 10 nM **5-610CP-Hoechst** (magenta). Numbers in the lower left corner indicate time. Scale bar: 10 µm. **d**, Three-color ZEISS Airyscan image of living HeLa cell at metaphase stained with 3 nM **4-TMR-LTX** (yellow), 20 nM **5-SiR-Hoechst** (magenta) and 1000 nM **6-510R-JAS** (cyan). Scale bar: 1 µm.

Altogether, our experiments demonstrated that the NGE influences the hydrophobicity of the probes and encouraged us to test if this leads to a better cell membrane permeability.

### Characterization of probe-target interaction in vitro

Rhodamine probes are prone to aggregation, which significantly contributes to their fluorogenicity and might impair interaction with the target ^8^. We observed a high correlation between fluorescence increase after binding to tubulin and after SDS addition, which indicates that affinity towards tubulin is sufficient to dissociate the probe from the aggregates (Supplementary Fig. 6). In both cases, fluorogenicity changes in the ordering TMR < 580CP ≈ 610CP < SiR. Next, we carried out a tubulin polymerization assay which showed that all probes, except **5-SiR-CTX**, were able to stabilize microtubules and are functional (Supplementary Fig. 7).

**Fig. 6.**
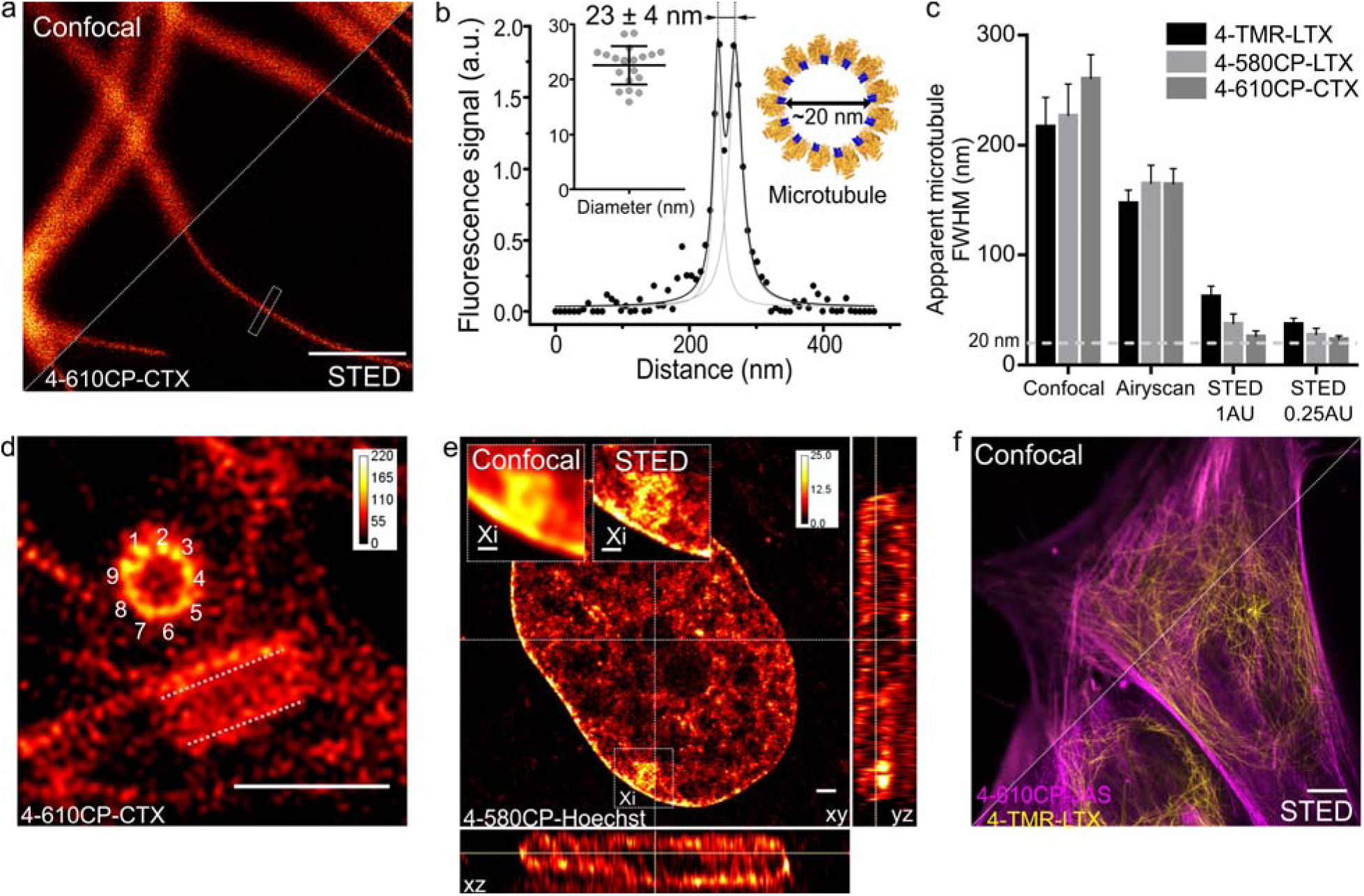
STED nanoscopy imaging of living cells stained with rhodamine 4’-isomer probes. **a**, Confocal and STED images taken with 0.25 AU pinhole of microtubules in human fibroblasts stained with 100 nM **4-610CP-CTX** for 1h at 37°C. Scale bar: 1 µm. **b**, Fluorescence intensity profile at the rectangle in panel **a**. Insets show measured (mean ± s.d., N = 20) and predicted diameter from cryo-electronmicroscopy model of tubulin (orange) bound to paclitaxel (blue). **c**, Apparent microtubule FWHM measured by the indicated microscopy methods. Human fibroblasts stained with 100 nM probes for 1h at 37°C. Data presented as mean ± s.d., N ≥ 3 independent fields of view, each time n ≥ 10 microtubules. **d**, Nine-fold symmetry of centriole resolved in the deconvolved STED DyMIN^43^ image of U-2 OS cell stained with 1 µM **4-610CP-CTX** for 1h at 37°C. White dashed lines mark a second centriole. Scale bar: 1 µm. **e**, Deconvolved STED image of human female fibroblast nucleus stained with 100 nM **4-580CP-Hoechst** showing the inactivated X chromosome (Xi). Insets - zoomed-in confocal and STED images of Xi region. Scale bars: 500 nm (insets), 1 µm (main image). **f**, Two-colour STED no-wash image of human fibroblasts stained with 100 nM **4-610CP-JAS** and 10 nM **4-TMR-LTX** for 1h at 37°C. Scale bar: 10 µm.

### Performance of 4′-carboxyrhodamines tubulin probes in living cells

Cabazitaxel or Larotaxel are anticancer drugs displaying superior resistance to efflux pumps and brain blood permeability ^26, 27^. The cytotoxicity of tubulin probes is implied by the design, which exploits the attachment of fluorophores to the 3’-position of taxanes ^8^. We hypothesized that NGE enhanced spirolactonization should increase the cell membrane permeability, resulting in stronger staining of tubulin and higher cytotoxicity. Thus, we stained HeLa, U-2 OS and primary human dermal fibroblasts with the series of tubulin probes at 100 nM concentration (Fig. 3a and Supplementary Fig. 8). All probes based on 4⍰-carboxamide dyes, with the exception of **4-SiR-CTX**, stained the cells much more strongly compared to the 5’- or 6’-regioisomer. The improvement of staining ranged from 3-up to 20-fold for primary human fibroblasts (Fig. 3b). In good agreement with these results, all 4’-regiosiomer probes, except **4-SiR-CTX**, demonstrated higher cytotoxicity in HeLa cells compared to the 5’- and 6’-regioisomer probes (Fig. 3c, Supplementary Fig. 9 and Supplementary Table 6).

Not only poor membrane permeability, but also active efflux might impede cell staining. To test the effect of multidrug-resistance (MDR) efflux pumps, we took advantage of U-2 OS cells, which are known to express MDR proteins susceptible to a broad specificity inhibitor Verapamil ^28^. Staining with **TMR, 580CP** and **610CP** 4’-regioisomer probes were independent of Verapamil, suggesting that these probes are not good substrates of MDR efflux pumps. In contrast, the best performing silicon-rhodamine probe **6-SiR-CTX** was prone to the action of efflux pumps and required Verapamil for efficient staining (Supplementary Fig. 10).

### Probes targeting DNA and actin

Superior cell membrane permeability of NGE carrying tubulin probes encouraged us to test the versatility of this approach with the probes targeting DNA and actin. To this end, we synthesized **4-TMR-Hoechst, 4-580CP-Hoechst** and **4-610CP-JAS**. All these probes stained their targets more intensively compared to 5’- and 6’-regioisomers in living cells (Fig. 4, Supplementary Fig. 11 and 12). DNA probes showed similar affinity to previously reported 5’- and 6’-regioisomers and a single DNA binding mode similarly as the 5‘-regioisomer (Supplementary Fig. 11b) ^29^.

Hoechst and its derivatives interfere with DNA synthesis leading to changes in S and G2/M phases ^29^. Jasplakinolide is known to cause multinucleation and apoptosis causing DNA fragmentation^30.^. Knowing these phenotypes, we monitored alterations in the cell cycle progression and identified the cytotoxicity threshold (Supplementary Fig. 11d and 12c).

### Confocal and Airyscan microscopy of living cells

Confocal images of tubulin stained with 4’-regioisomers in living cells showed excellent contrast exceeding 50-fold single microtubule to cytoplasm signal ratio even if excess of the probe was not removed (Fig. 5a and Supplementary Fig. 13). Airyscan images benefited from increased resolution (∼150 nm) and reached up to 100-fold single microtubule to cytoplasm signal ratio (Fig. 5a and Supplementary Fig. 14). This clearly demonstrated excellent permeability, high affinity and selectivity of the taxane probes. A similar improvement was observed for DNA and actin probes (Fig. 4, Supplementary Fig. 15 and Supplementary movie 1).

The **4-TMR-LTX** probe shows a very low cytotoxicity threshold (EC_50_ = 6.2 ± 0.5 nM, Fig. 3c), approaching the cytotoxicity of taxanes ranging from 1 - 10 nM^31^ (Supplementary Table 7). Taxanes are known to accumulate inside the cells up to 1000-fold ^32^. Indeed, we were able to image fibroblasts stained with sub-nanomolar concentrations of **4-TMR-LTX** (Fig. 5b). Despite its relatively low fluorogenicity, a washing step was not required to achieve high contrast. Noteworthy, at picomolar concentrations, **4-TMR-LTX** selectively stains centrosome, likely due to a higher local tubulin concentration in this organelle (Fig. 5b, and Supplementary movie 2).

The spectral separation between **TMR** and **610CP** allows two colour sequential imaging (Supplementary Fig. 16). We could resolve all cell cycle stages in 3D while imaging HeLa cells stained with **4-TMR-LTX** and **5-610CP**-**Hoechst** (Fig. 5c and Supplementary movie 3). We exploited **6-510R-JAS** ^33^, **4-TMR-LTX** and **5-SiR-Hoechst** ^29^ for three colour sequential Airyscan Z-stack imaging of mitotic HeLa cells meeting Nyquist criteria (50-50-150 nm in x-y-z axes) without significant bleaching (Fig. 5d and Supplementary movie 4).

### Super-resolution STED microscopy of living cells

Finally, we applied our new probes in STED nanoscopy. The fluorescence of **TMR** based probes could be inhibited by 660 and 775 nm lasers resulting in significant improvement of the resolution and yielding apparent microtubule diameter below 80 nm in all the tested cases (Supplementary Fig. 17). Despite significant fluorescence quenching by the STED laser (10 MW/cm^2^), we could reach up to 45-fold single microtubule to cytoplasm signal ratio while staining with **4-610CP-CTX** (Supplementary Fig. 18 and Fig. 5a). We took advantage of the no-wash conditions and recorded STED images with up to a 100 frames time-course without significant bleaching (Supplementary movie 5). Good labeling efficiency and brightness of the probes allowed acquisition of STED images with 0.25 Airy units pinhole at increased resolution (Fig. 6a-c). For the first time we could resolve the microtubule diameter of 23 ± 4 nm in living cells. This value closely matches the dimensions measured with cryo-electron microscopy (Fig. 6b) ^34^. In addition, we obtained excellent quality images of tubulin in the centrosome demonstrating the 9-fold symmetry of this organelle (Fig. 6d). 3D STED allowed to resolve, independently of orientation, the tube-like structure of centrioles stained with **4-610CP-CTX** in living cells without probe removal (Supplementary Fig. 19 and Supplementary movie 6). Using the **4-580CP-Hoechst** probe we visualized chromatin domain clusters of the inactivated X chromosome in primary human fibroblasts (Fig. 6e). Finally, we demonstrate two-color STED imaging using **4-TMR-LTX** and **4-610CP-JAS** probes (Fig. 6f and Supplementary Fig. 16d).

## Discussion

Due to their excellent photophysical properties and biocompatibility, rhodamines are among the most popular dyes used for synthesis of fluorescent probes for live-cell imaging. Controlling spirolactonization of rhodamine dyes facilitates cell permeability and increases fluorogenicity of the final probes. Herein we exploited the neighbouring group effect (NGE) occurring in the carboxyrhodamines positional 4’-conjugates for enhancing spirolactonization without any modifications of the dye’s core structure. This led to increased hydrophobicity and significantly enhanced cell permeability, demonstrating a significant contribution of the fluorophore configuration to the properties of the final probe.

Several studies have used ^dye^D_50_ to guide rational fluorophore design. ^11, 12, 13, 35^. In our case this would not have worked; with the exception of **4-SiR-COOH**, ^dye^ D_50_ values of 4’-, 5’- and 6’-isomers are very close to each other, thus implying no major improvement at the free fluorophore level. For the probes, the situation is entirely different: after conjugation to the targeting ligand, one of the neighbouring carboxylic groups is replaced with the carboxamide group, which enhances NGE and considerably increases ^probe^D_50_ of 4’-isomers. Furthermore, even the ^probe^D_50_ value is a poor predictor for probe performance, as the optimal spirolactonization propensity is different for each probe. For example, **4-TMR-LTX** efficiently stains cells at concentration as low as 0.06 nM, which clearly points out its high cell membrane permeability, despite the fact that its ^probe^D_50_ = 26.5 ± 0.5 is the lowest among 4’-isomers.

NGE dramatically enhances probes based on more hydrophilic **4-TMR-COOH, 4-580CP-COOH** and **4-610CP-COOH** dyes, but not on a more hydrophobic **4-SiR-COOH**. In **4-SiR-CTX**, NGE pushes ^probe^D_50_ outside the water polarity range, causing dominance of a non-absorbing spirolactone, which results in decreased solubility and increased aggregation in water. This manifests in low fluorescence, weak polymerized tubulin stabilization *in vitro*, low cytotoxicity and poor staining. Thus, **6-SiR-CTX**, which has ^probe^D_50_ close to the dielectric constant value of water, is the best performing SiR-based probe.

The unprecedented permeability of new tubulin probes allowed imaging at nanomolar - picomolar concentrations, which is in sharp contrast to micromolar-high concentrations reported in other studies ^11, 12, 13, 35, 36, 37, 38^. At such low concentrations, no washing step was required even with probes of low fluorogenicity, such as **4-TMR-LTX** which can be used as a selective stain for the centrosome at picomolar range.

On the other side, high cell permeability of **4-610CP-CTX** allowed exceptionally dense labelling of microtubules and provided sufficiently strong signal for STED imaging with the pinhole set to an extreme value of 0.25 AU. Thereby, to the best our knowledge for the first time, we resolved microtubule diameters of 22 ± 3 nm in living cells, which matches the value obtained by cryo-EM^34^.

We demonstrated the generality of the NGE by synthesizing probes targeting DNA and actin, which can also operate at low nanomolar concentrations indicating unrestricted permeability, very high affinity and specificity of the new probes. This decreases the chances for unspecific probe accumulation in the membranes. Although invisible, such off-targeting might disturb physiological processes.

In conclusion, we employed the neighbouring group effect in the 4’-regioisomer of rhodamine fluorophores to create, to our best knowledge, the most efficient probes for tubulin, actin and DNA labelling in living cells (Supplementary Table 7). Their excellent spectroscopic properties, outstanding cell permeability and specificity makes them powerful tools for many types of microscopy techniques, including super-resolution methods. The simple approach of positional isomerism is applicable to rhodamine- and fluorescein-type fluorophores, does not require modification of the xanthene ring system and can be used for generating new probes and for improving the performance of existing ones.

## Acknowledgement

The authors are grateful to Dr. Vladimir Belov, Jan Seikowski, Jens Schimpfhauser and Jürgen Bienert for the NMR/ESI-MS measurements of numerous probes and providing Boc-JAS actin targeting moiety. They also acknowledge Dr. Peter Lenart and Dr. Antonio Politi (Live-cell imaging facility) for the possibility to perform live-cell confocal and Airyscan imaging. JB and GL are grateful to the Max Planck Society for a Nobel Laureate Fellowship. GK acknowledges the Max Planck Institute for Biophysical Chemistry for a Manfred Eigen Fellowship, the iNEXT consortium (PID: 7187, Horizon 2020 program) for the possibility to perform imaging experiments in the Advanced Light Microscopy Facility at EMBL in Heidelberg and is very grateful to Dr. Sebastian Schnorrenberg for guidance. The authors acknowledge Dr. Stefan W. Hell, Dr. Steffen Sahl and Jaydev Jethwa for critical reading of the manuscript.

## Author contributions

JB and GL conceived and planned the study. JB, GK, RG, TG and GL performed the experiments. JB, GK, RG and GL performed the data analysis. All authors contributed to manuscript writing. The manuscript was written with contributions from all authors.

## Data availability

The data that support the findings of this study are available from the corresponding author upon request.

## Methods

### Quantum chemical calculations

The initial geometries of the studied molecules were generated by using a molecular mechanics method (force field MMF94, steepest descent algorithm) and a systematic conformational analysis was implemented in Avogadro 1.1.1 software. The minimum energy conformer geometries found by molecular mechanics were further optimized with the Gaussian 09 program package^24^ by means of density functional theory (DFT) using the Becke, 3-parameter, Lee–Yang–Parr (B3LYP)^39, 40^ exchange-correlation hybrid functional with 6-311G++(d,p) basis set^41^, including the polarizable continuum model^42^ in 1,4-dioxane or water. Further, harmonic vibrational frequencies were calculated to verify the stability of the optimized geometries. All the calculated vibrational frequencies were real (positive) indicating the true minimum of the calculated total potential energy of the optimized system. The computation was performed at the High Performance Computing center in Göttingen provided by GWDG.

### Maintenance and preparation of cells

Human primary dermal fibroblasts and HeLa cells were cultured in high-glucose DMEM (Thermo Fisher, #31966047) and 10% FBS (Life Technologies, #10270-106) with a presence of 1% Penicillin-Streptomycin (Sigma, #P0781) in a humidified 5% CO_2_ incubator at 37 °C. The cells were split every 3-4 days or at confluence. Cells were seeded in glass bottom 12-well plates (MatTek, #P12G-1.0-14-F) before imaging experiments. The U2OS cells were cultured in McCoys 5A medium (Thermo fisher #16600082) and 10% FBS (Merck Biochrom, #S0615) with a presence of 1% Sodium pyruvate (Sigma, #S8636) and 1% of Penicillin-Streptomycin (Sigma #P0781) in a humidified 5% CO_2_ incubator at 37 °C.

Cells were stained with the fluorescent probes in DMEM (Thermo Fisher, #31053-028) supplemented with 10% FBS (Thermo Fisher, #10082139) at 37 °C and 5% CO_2_. If needed, the cells were washed 2 times with HBSS (Lonza, #BE10-527F) and imaged in DMEM with 10% FBS. No-wash experiments were performed in DMEM with 10% FBS after probe addition and incubation for the indicated period of time. For Airyscan experiments, cells were seeded in a 10-well plate (Greiner bio-one, #543079) 1 day before staining. The probes were applied to the cells in DMEM medium, and after 1h or overnight incubation cells were imaged without washing.

### STED microscope with 775 nm laser

Comparative confocal and STED images were acquired on an Abberior STED 775 QUAD scanning microscope (Abberior Instruments GmbH) equipped with 561 nm and 640 nm 40 MHz pulsed excitation lasers, a pulsed 775 nm 40 MHz STED laser, and an UPlanSApo 100x/1.40 Oil objective. The following detection windows were used: for the TMR/580CP channel 615 / 20 nm, and for the 610CP/SiR channel 685 / 70 nm. In this setup, voxel size was 15 – 30 nm in the xy plane and 150 nm in z-axis for 2D STED images. For the STED imaging with 0.25 AU pinhole, pixel size set to 7 × 7 nm and images acquired with 2 line accumulation. Laser powers were optimized for each sample. 3D STED images were acquired using pinhole set to 0.8 AU, voxel size set to 40 × 40 × 40 nm, 3D STED doughnut set to 90%, with single line accumulation and xzy scanning mode.

Estimation of the STED effect was performed by varying the STED laser power from 0 to 100% while measuring cells stained with tubulin probes. Obtained data were fitted using GraphPad Prism 6 to the following equation:

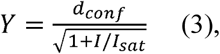

where *d*_*conf*_ – confocal resolution, *I* – STED laser intensity power, *I*_*sat*_ – saturating STED laser intensity power.

### STED microscope with 660 nm laser

STED images with 660 nm STED laser were acquired on a Leica TCS SP8 STED 3X scanning microscope (Leica microsystems) equipped with HC PL APO CS2 100x/1.40 Oil objective and the incubator set to 37°C and 5% CO_2_. The 4-TMR-LTX probe was excited with the 561 nm, 80 MHz pulsed line of a white light laser, de-excited with a continuous wave 660 nm STED laser and detected in the 600 / 50 nm window. In this setup, the pixel size was set to 23 nm in the xy plane. The STED effect was estimated as described above.

### Airyscan microscope

Live-cell imaging was performed on a Zeiss LSM880 system (Carl Zeiss) equipped with oil immersion Plan-Apochromat 63X/1.40 Oil Corr M27 objective and the incubator set to 37°C and 5% CO_2_. The following excitation lasers in combination with confocal or Airyscan acquisition regimes were used: argon laser 488 nm for 510R probes, 561 nm diode laser for TMR and 580CP probes, 633 nm diode laser for 610CP and SiR probes. For TMR and 610CP, a combination of BP 570 - 620 + LP 645 filters with dichroic beam splitter SBS SP 615 in the TMR channel was used. The TMR, 510R and SiR combination was excited and scanned in sequence using BP 495 - 550 + LP 570 filters. Laser powers were optimized for each sample. Detailed excitation and detection schemes are shown in Supplementary Figure 16.

## Refenrences

1. Sahl SJ, Hell SW, Jakobs S. Fluorescence nanoscopy in cell biology. Nat Rev Mol Cell Biol 2017, 18(11): 685–701.

2. Keereweer S, Van Driel PB, Snoeks TJ, Kerrebijn JD, Baatenburg de Jong RJ, Vahrmeijer AL, et al. Optical image-guided cancer surgery: challenges and limitations. Clin Cancer Res 2013, 19(14): 3745–3754.

3. Hell SW. Nanoscopy with Focused Light (Nobel Lecture). Angew Chem Int Ed Engl 2015, 54(28): 8054–8066.

4. Wang L, Frei MS, Salim A, Johnsson K. Small-Molecule Fluorescent Probes for Live-Cell Super-Resolution Microscopy. J Am Chem Soc 2019, 141(7): 2770–2781.

5. van de Linde S, Heilemann M, Sauer M. Live-cell super-resolution imaging with synthetic fluorophores. Annu Rev Phys Chem 2012, 63: 519–540.

6. Uno SN, Tiwari DK, Kamiya M, Arai Y, Nagai T, Urano Y. A guide to use photocontrollable fluorescent proteins and synthetic smart fluorophores for nanoscopy. Microscopy (Oxf) 2015, 64(4): 263–277.

7. Zhang G, Zheng S, Liu H, Chen PR. Illuminating biological processes through site-specific protein labeling. Chem Soc Rev 2015, 44(11): 3405–3417.

8. Lukinavičius G, Mitronova GY, Schnorrenberg S, Butkevich AN, Barthel H, Belov VN, et al. Fluorescent dyes and probes for super-resolution microscopy of microtubules and tracheoles in living cells and tissues. Chem Sci 2018, 9(13): 3324–3334.

9. Lukinavičius G, Reymond L, Umezawa K, Sallin O, D’Este E, Gottfert F, et al. Fluorogenic Probes for Multicolor Imaging in Living Cells. J Am Chem Soc 2016, 138(30): 9365–9368.

10. Butkevich AN, Belov VN, Kolmakov K, Sokolov VV, Shojaei H, Sidenstein SC, et al. Hydroxylated Fluorescent Dyes for Live-Cell Labeling: Synthesis, Spectra and Super-Resolution STED. Chemistry – A European Journal 2017, 23(50): 12114–12119.

11. Butkevich AN, Mitronova GY, Sidenstein SC, Klocke JL, Kamin D, Meineke DN, et al. Fluorescent Rhodamines and Fluorogenic Carbopyronines for Super-Resolution STED Microscopy in Living Cells. Angew Chem Int Ed Engl 2016, 55(10): 3290–3294.

12. Zheng Q, Ayala AX, Chung I, Weigel AV, Ranjan A, Falco N, et al. Rational Design of Fluorogenic and Spontaneously Blinking Labels for Super-Resolution Imaging. ACS Cent Sci 2019, 5(9): 1602–1613.

13. Wang L, Tran M, D’Este E, Roberti J, Koch B, Xue L, et al. A general strategy to develop cell permeable and fluorogenic probes for multi-colour nanoscopy. bioRxiv 2019: 690867.

14. Teh EJ, Leong YK, Liu Y. Isomerism and solubility of benzene mono- and dicarboxylic acid: its effect on alumina dispersions. Langmuir 2011, 27(1): 49–58.

15. Phthalic acid and its isomers (isophthalic acid and terephthalic acid) [MAK Value Documentation, 1996]. The MAK-Collection for Occupational Health and Safety, pp 242–246.

16. Beija M, Afonso CA, Martinho JM. Synthesis and applications of Rhodamine derivatives as fluorescent probes. Chem Soc Rev 2009, 38(8): 2410–2433.

17. Scala-Valéro C, Doizi D, Guillaumet G. Synthesis of isomers of rhodamine 575 and rhodamine 6G as new laser dyes. Tetrahedron Letters 1999, 40(26): 4803–4806.

18. Baeyer A. Ueber eine neue Klasse von Farbstoffen. Berichte der deutschen chemischen Gesellschaft 1871, 4(2): 555–558.

19. Deo C, Sheu SH, Seo J, Clapham DE, Lavis LD. Isomeric Tuning Yields Bright and Targetable Red Ca(2+) Indicators. J Am Chem Soc 2019, 141(35): 13734–13738.

20. Grimm JB, Sung AJ, Legant WR, Hulamm P, Matlosz SM, Betzig E, et al. Carbofluoresceins and carborhodamines as scaffolds for high-contrast fluorogenic probes. ACS Chem Biol 2013, 8(6): 1303–1310.

21. Grimm JB, Lavis LD. Synthesis of rhodamines from fluoresceins using Pd-catalyzed C-N cross-coupling. Org Lett 2011, 13(24): 6354–6357.

22. Savarese M, Aliberti A, De Santo I, Battista E, Causa F, Netti PA, et al. Fluorescence lifetimes and quantum yields of rhodamine derivatives: new insights from theory and experiment. J Phys Chem A 2012, 116(28): 7491–7497.

23. Åkerlöf G, Short AO. The Dielectric Constant of Dioxane—Water Mixtures between 0 and 80°. J Am Chem Soc 1936, 58(7): 1241–1243.

24. Frisch MJ, Trucks GW, Schlegel HB, Scuseria GE, Robb MA, Cheeseman JR, et al. Gaussian 09 Rev. D.01. Wallingford, CT; 2009.

25. Valko K, Bevan C, Reynolds D. Chromatographic Hydrophobicity Index by Fast-Gradient RP-HPLC: A High-Throughput Alternative to log P/log D. Anal Chem 1997, 69(11): 2022–2029.

26. Abidi A. Cabazitaxel: A novel taxane for metastatic castration-resistant prostate cancer-current implications and future prospects. J Pharmacol Pharmacother 2013, 4(4): 230–237.

27. Metzger-Filho O, Moulin C, de Azambuja E, Ahmad A. Larotaxel: broadening the road with new taxanes. Expert Opin Investig Drugs 2009, 18(8): 1183–1189.

28. Choi CH. ABC transporters as multidrug resistance mechanisms and the development of chemosensitizers for their reversal. Cancer Cell Int 2005, 5: 30.

29. Bucevičius J, Keller-Findeisen J, Gilat T, Hell SW, Lukinavičius G. Rhodamine-Hoechst positional isomers for highly efficient staining of heterochromatin. Chem Sci 2019, 10(7): 1962–1970.

30. Moulding DA, Blundell MP, Spiller DG, White MR, Cory GO, Calle Y, et al. Unregulated actin polymerization by WASp causes defects of mitosis and cytokinesis in X-linked neutropenia. J Exp Med 2007, 204(9): 2213–2224.

31. Ren S, Wang Y, Wang J, Gao D, Zhang M, Ding N, et al. Synthesis and biological evaluation of novel larotaxel analogues. Eur J Med Chem 2018, 156: 692–710.

32. Jordan MA, Toso RJ, Thrower D, Wilson L. Mechanism of mitotic block and inhibition of cell proliferation by taxol at low concentrations. Proc Natl Acad Sci U S A 1993, 90(20): 9552–9556.

33. Grimm F, Nizamov S, Belov VN. Green-Emitting Rhodamine Dyes for Vital Labeling of Cell Organelles Using STED Super-Resolution Microscopy. Chembiochem 2019, 20(17): 2248–2254.

34. Kellogg EH, Hejab NMA, Howes S, Northcote P, Miller JH, Diaz JF, et al. Insights into the Distinct Mechanisms of Action of Taxane and Non-Taxane Microtubule Stabilizers from Cryo-EM Structures. J Mol Biol 2017, 429(5): 633–646.

35. Grimm JB, Muthusamy AK, Liang Y, Brown TA, Lemon WC, Patel R, et al. A general method to fine-tune fluorophores for live-cell and in vivo imaging. Nat Methods 2017, 14(10): 987–994.

36. Lukinavičius G, Blaukopf C, Pershagen E, Schena A, Reymond L, Derivery E, et al. SiR-Hoechst is a far-red DNA stain for live-cell nanoscopy. Nat Commun 2015, 6: 8497.

37. Lukinavičius G, Reymond L, D’Este E, Masharina A, Gottfert F, Ta H, et al. Fluorogenic probes for live-cell imaging of the cytoskeleton. Nat Methods 2014, 11(7): 731–733.

38. Lukinavičius G, Umezawa K, Olivier N, Honigmann A, Yang G, Plass T, et al. A near-infrared fluorophore for live-cell super-resolution microscopy of cellular proteins. Nat Chem 2013, 5(2): 132–139.

39. Becke AD. Density-functional thermochemistry. III. The role of exact exchange. The Journal of Chemical Physics 1993, 98(7): 5648–5652.

40. Lee C, Yang W, Parr RG. Development of the Colle-Salvetti correlation-energy formula into a functional of the electron density. Phys Rev B Condens Matter 1988, 37(2): 785–789.

41. Davidson ER, Feller D. Basis set selection for molecular calculations. Chemical Reviews 1986, 86(4): 681–696.

42. Tomasi J, Mennucci B, Cammi R. Quantum Mechanical Continuum Solvation Models. Chemical Reviews 2005, 105(8): 2999–3094.

43. Heine J, Reuss M, Harke B, D’Este E, Sahl SJ, Hell SW. Adaptive-illumination STED nanoscopy. Proc Natl Acad Sci U S A 2017, 114(37): 9797–9802.

