## Supplementary material for "Enhancing biocompatibility of rhodamine fluorescent probes by a neighbouring group effect"

### Supplementary inventory

#### Supplementary figures

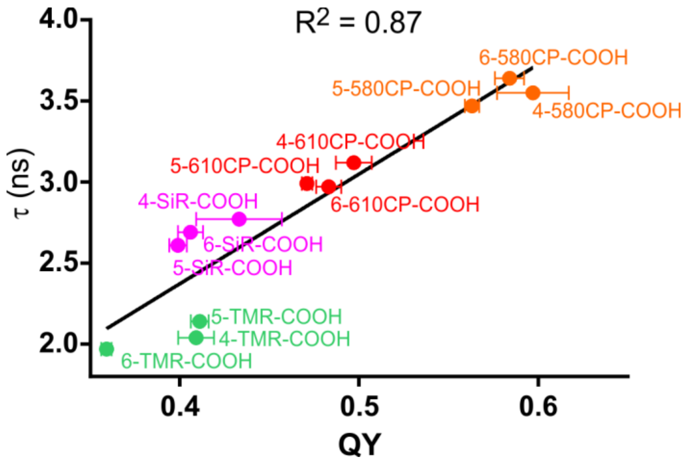

**Supplementary Figure 1.** Correlation between fluorescent dye quantum yield and florescence lifetime. Data points are presented as mean ± s.d. of three independently repeated experiments (N=3).

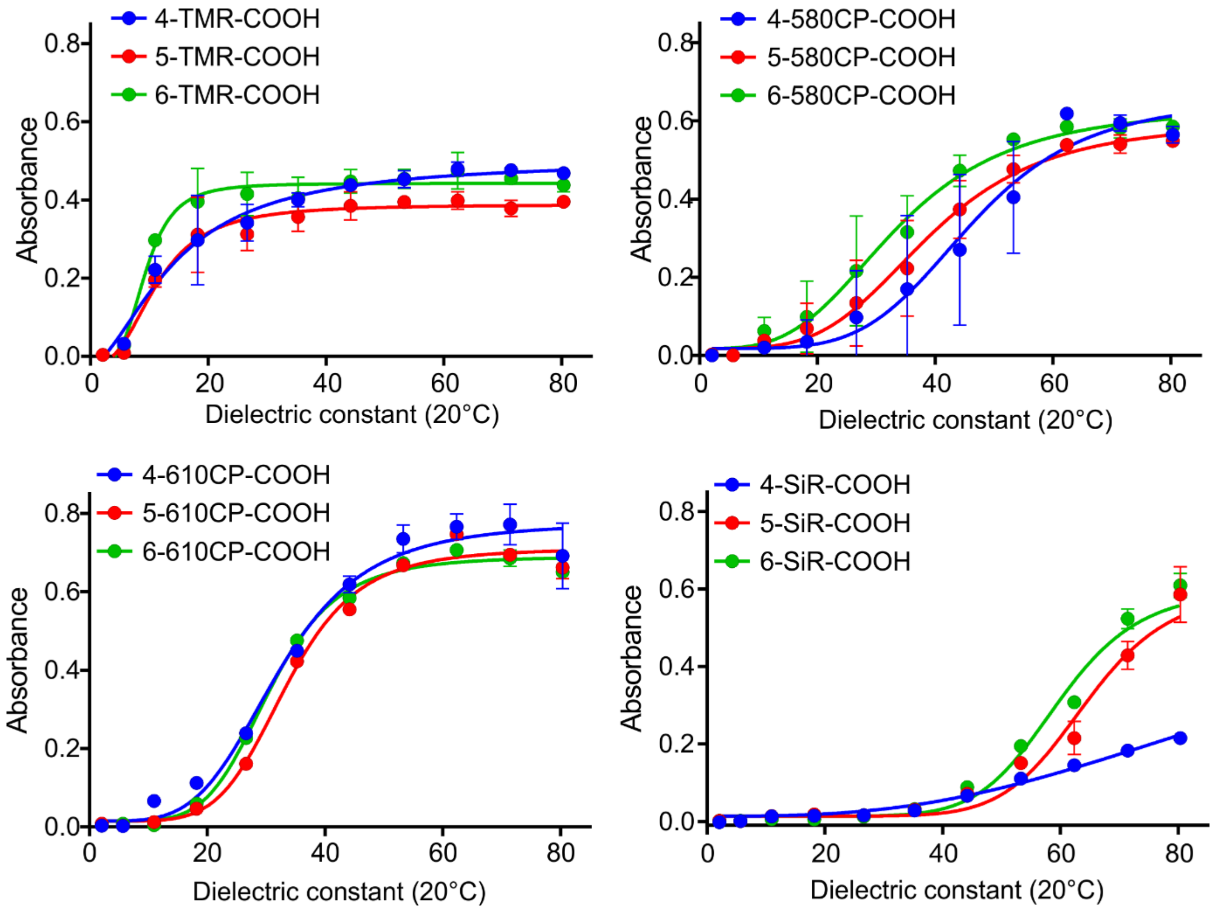

**Supplementary Figure 2.** Plots showing absorbance of dyes' positional isomers at λmax versus dielectric constant (D) of 1,4-dioxane-water mixtures. $D_{50}$ value was obtained by fitting to dose-response equation and corresponds to the D value that provokes half of the maximal absorbance. Data points are presented as mean ± s.d. of three independently repeated experiments (N=3).

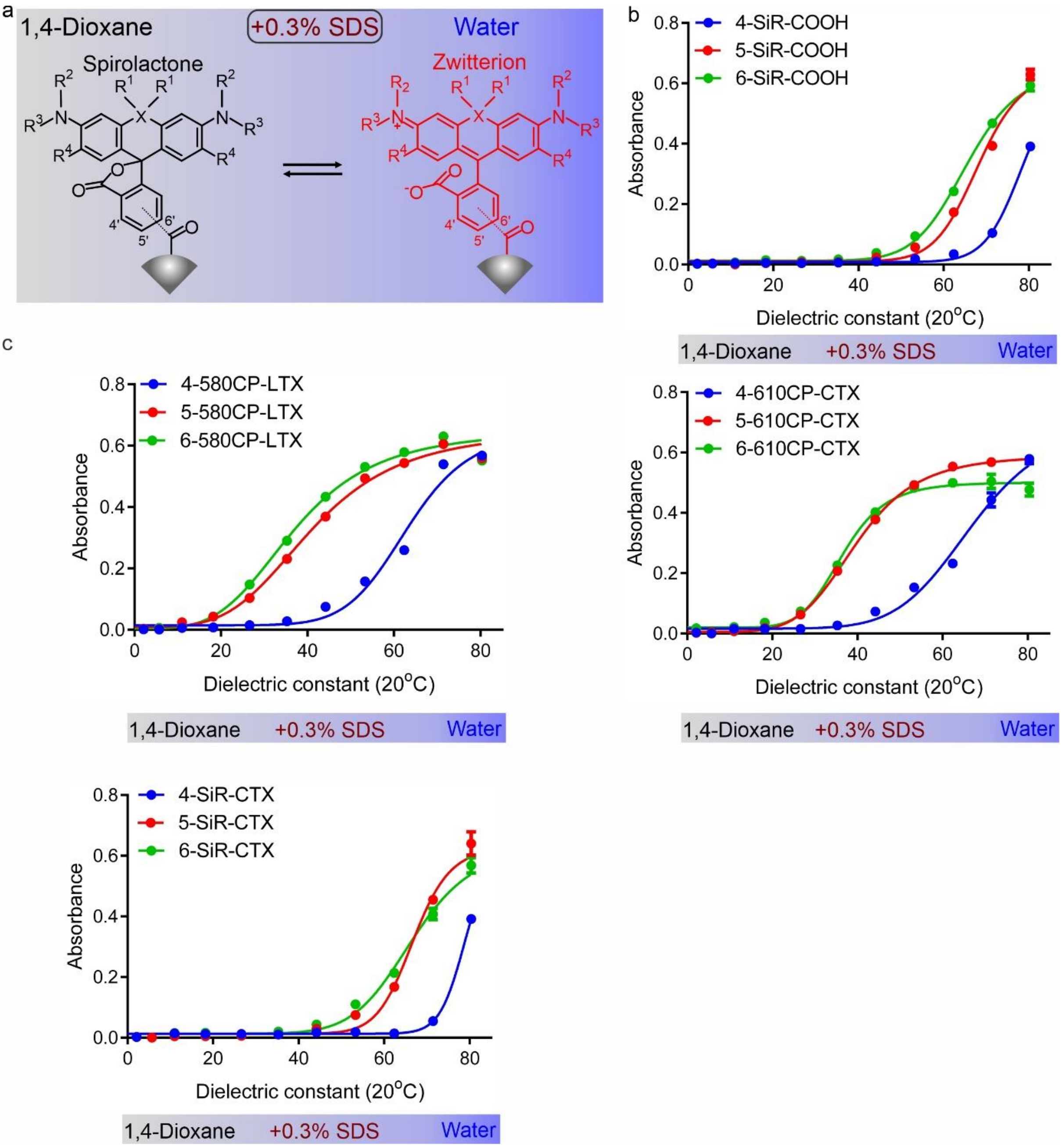

**Supplementary Figure 3.** Behaviour of the tubulin probes in 1,4-dioxane-water mixtures containing 0.3% SDS. **a.** Principal scheme of the mechanism of chromogenic properties. **b.** Plot showing absorbance of SiR-COOH positional isomers at λmax versus dielectric constant (D) of 1,4-dioxane-water mixtures with constant 0.3% SDS additive. $D_{50}$ value was obtained by fitting to dose-response equation and corresponds to the D value that provokes half of the maximal absorbance. Data points are presented as mean ± s.d. of three independently repeated experiments (N=3). **c.** Plots showing absorbance of probes' positional isomers at λmax versus dielectric constant (D) of 1,4-dioxane-water mixtures with constant 0.3% SDS additive. $D_{50}$ values were obtained by fitting to dose-response equation and corresponds to the D value that provokes half of the maximal absorbance. Data points are presented as mean ± s.d. of three independently repeated experiments (N=3).

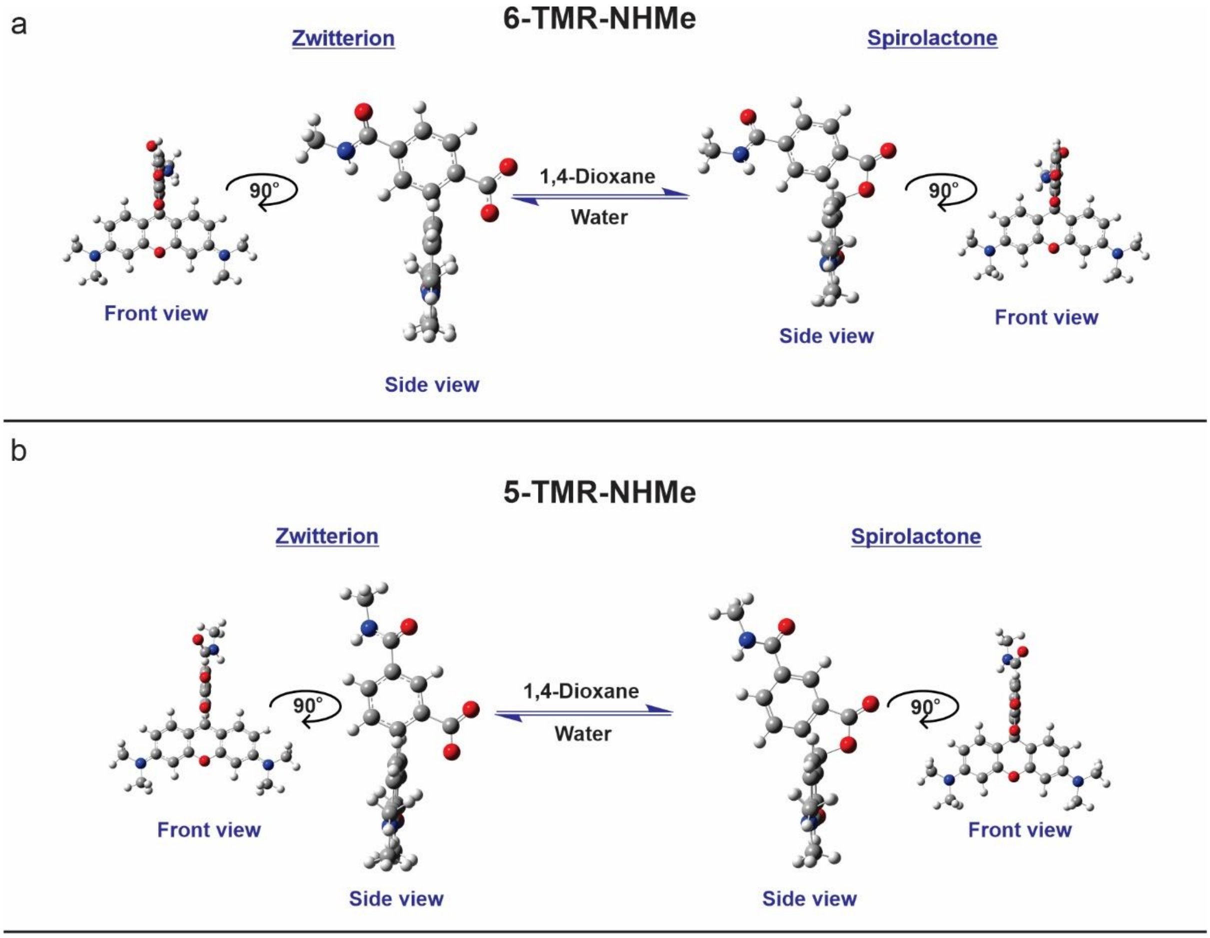

**Supplementary Figure 4**. DFT optimized geometries of model compounds with truncated linker and targeting moiety. **a.** DFT optimized geometries of model compound **6-TMR-NHMe** in both spirolactone and zwitterion forms in front and side views. **b.** DFT optimized geometries of model compound **5-TMR-NHMe** in both spirolactone and zwitterion forms in front and side views.

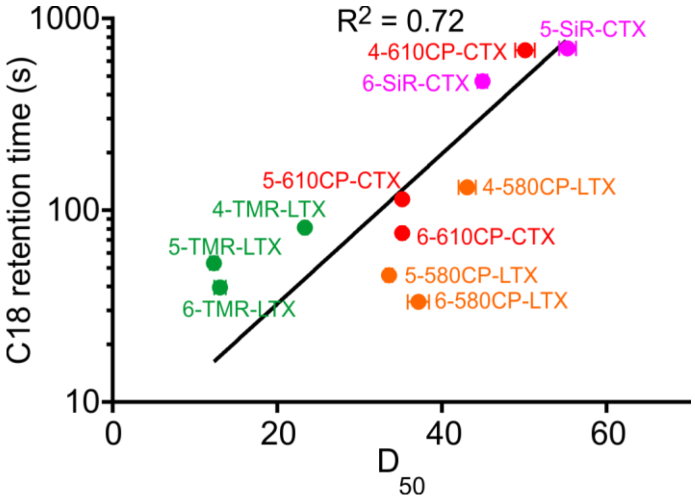

**Supplementary Figure 5**. Correlation between $D_{50}$ values and retention times on C18 HPLC column of the tubulin probes. Data points are presented as mean ± s.d. of three independently repeated experiments (N=3).

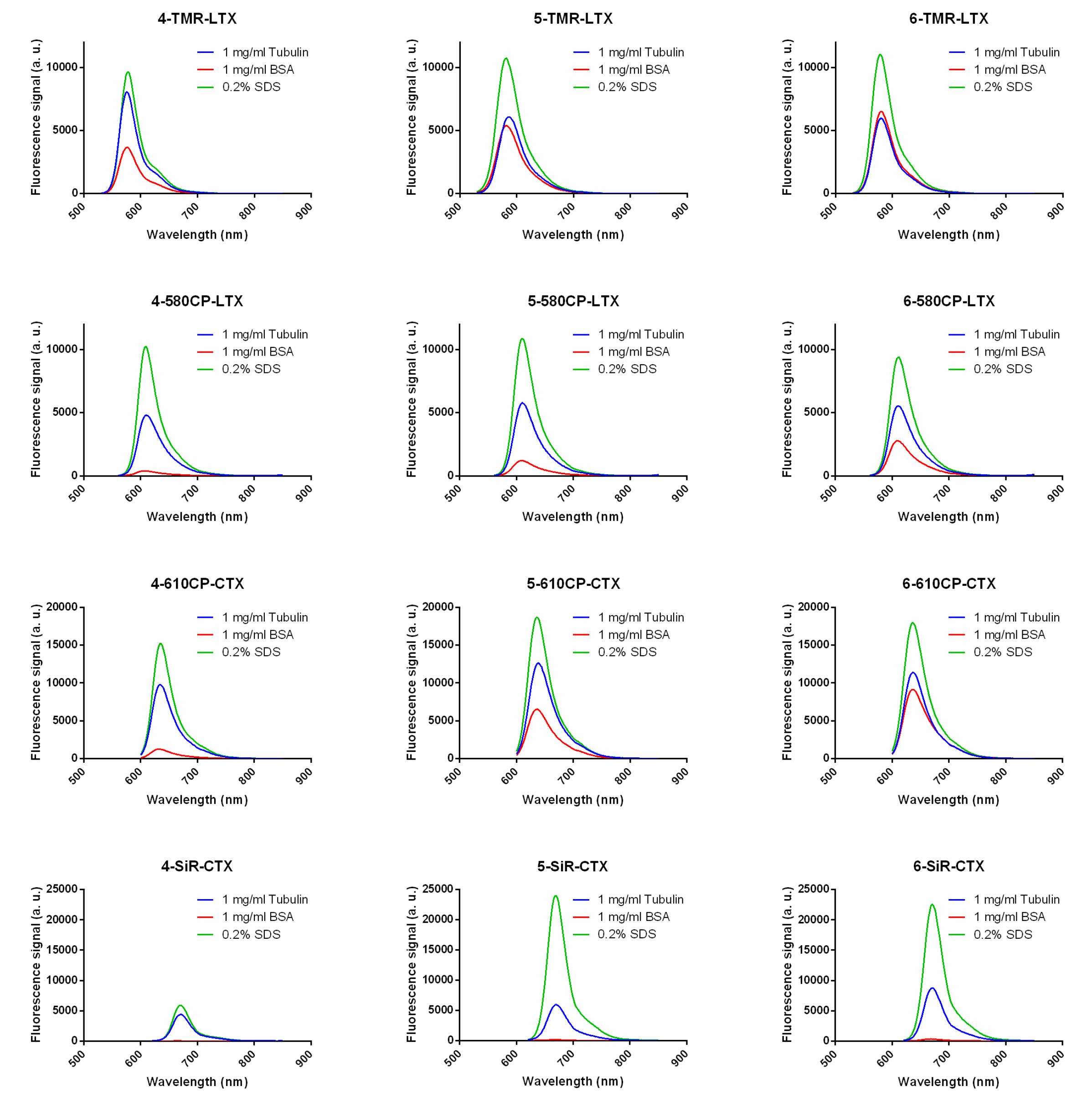

**Supplementary Figure 6.** Fluorescence spectra of the tubulin probes. Spectra were recorded after incubating 2 µM probes with 1 mg/ml tubulin + 1 mM GTP (blue), 1 mg/ml BSA (red) or 0.2% SDS (green) at 37 °C for 3 h to ensure complete tubulin polymerization. Spectra are presented as averages of three independently repeated experiments (N=3).

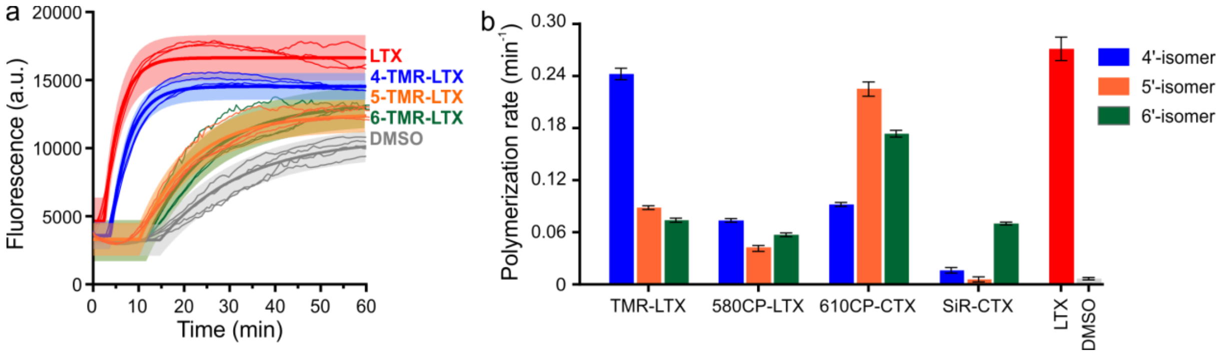

**Supplementary Figure 7.** Stimulation of tubulin polymerization in vitro properties by fluorescent probes. **a.** time-courses of tubulin polymerization in the presence of 3 µM TMR-LTX probes at 37°C. Individual traces from 3 or 4 independent experiments are shown (thin lines), together with the fitting curve (thick line); 95% prediction intervals are shaded. **b.** tubulin polymerization rate constants, obtained from the traces analogous to those shown in a. The time-courses were fitted into "plateau followed by one-phase association" function, and the derived rate constants are expressed as the best-fit values ± standard error.

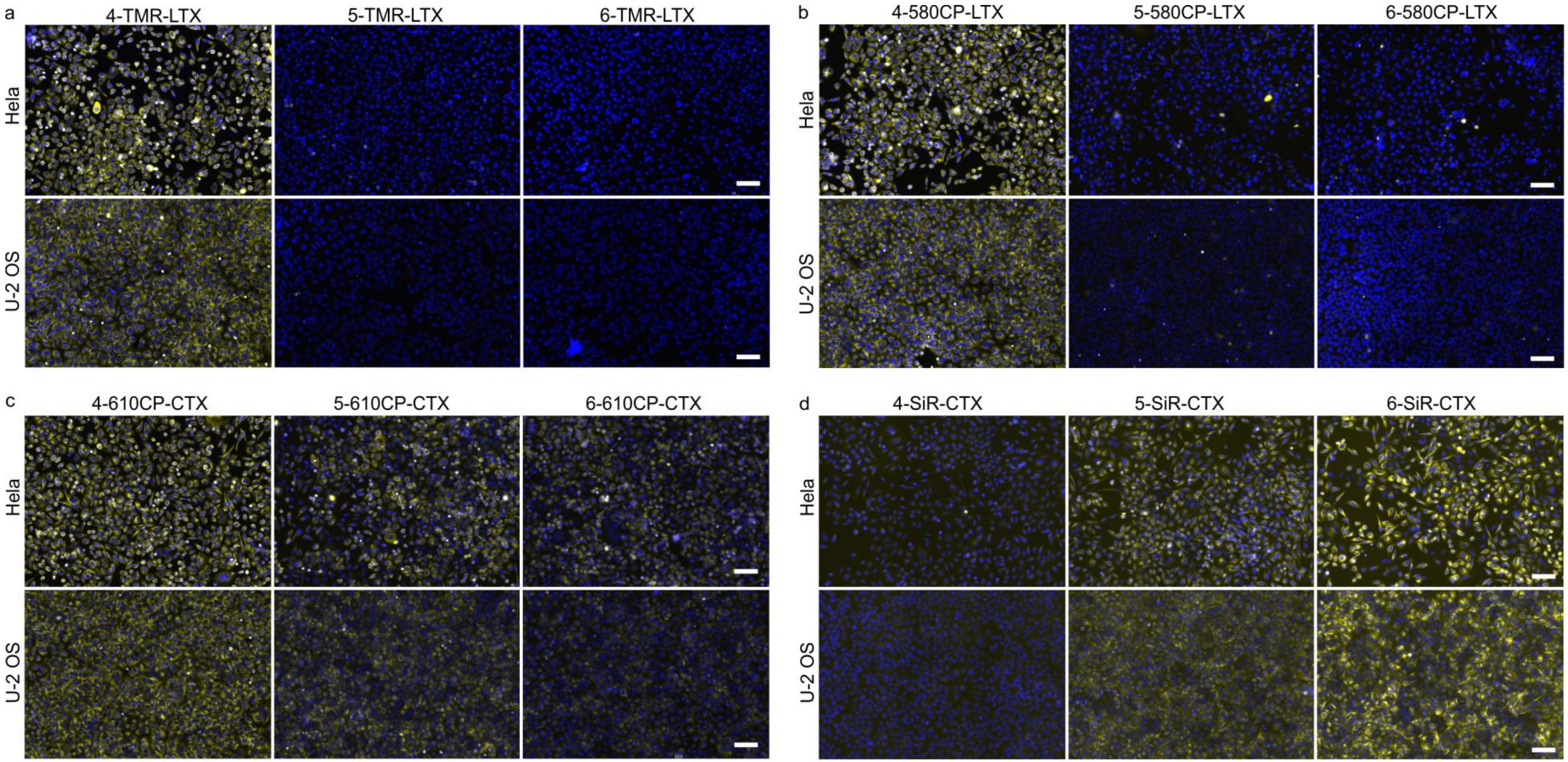

**Supplementary Figure 8.** Wide field fluorescence microscopy images of living Hela and U-2 OS cell lines stained with the tubulin probes. **a. TMR-LTX**; **b. 580CP-LTX**; **c. 610CP-CTX** and **d. SiR-CTX**. Living cells were stained with a mixture of 100 nM probe (yellow) and 0.1 µg/ml Hoechst 33342 (blue) in DMEM growth medium containing 10% FBS at 37 °C for 1 h, washed once with HBSS and imaged on Biotek Lionheart FX automated microscope. Scale bars 100 µm.

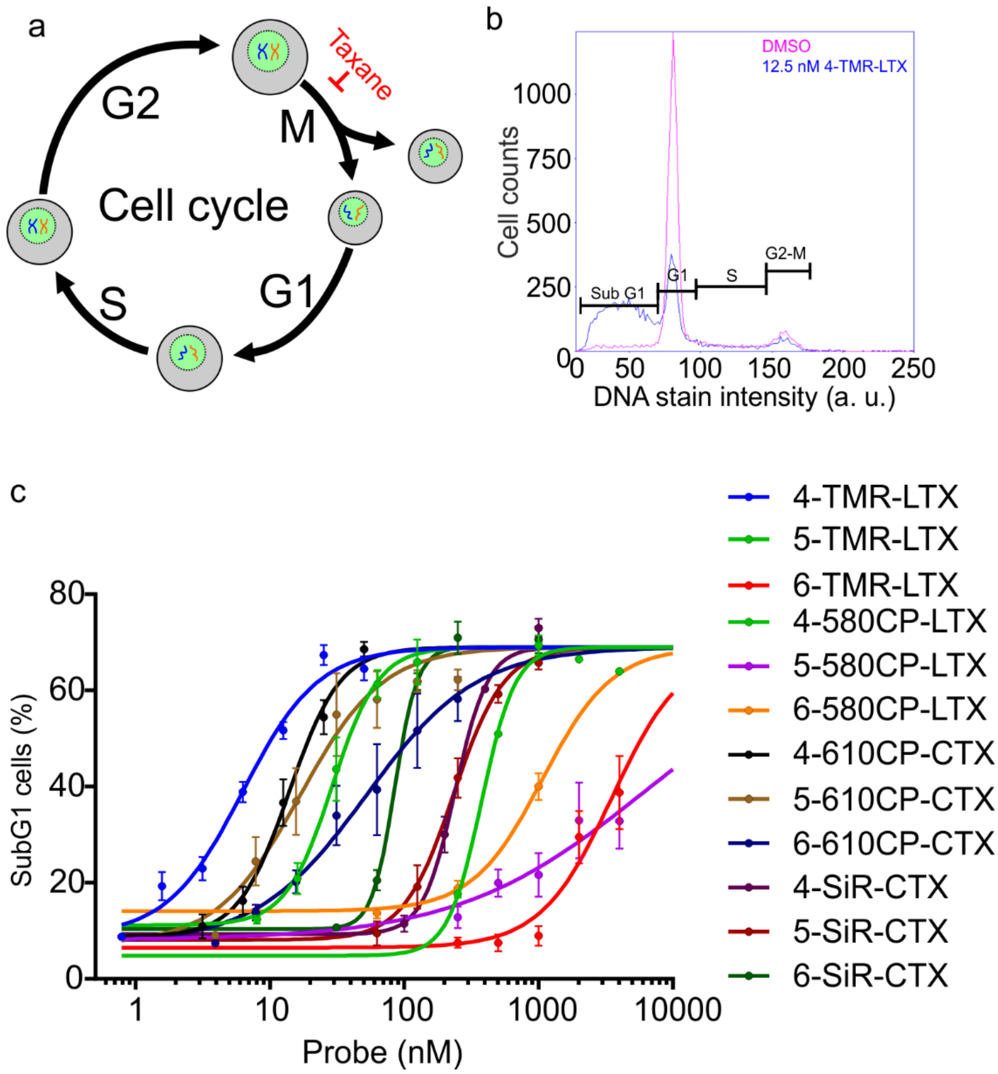

**Supplementary Figure 9.** Cell cycle perturbations in HeLa cells induced by fluorescent tubulin probes. **a.** Cytotoxicity of taxanes results from the inhibition of the cell cycle at the stage of mitosis (M). **b.** A representative histogram of DNA content distribution in HeLa cell population treated with DMSO or 12.5 nM **4-TMR-LTX** for 24 h. Indicated cell cycle phases (Sub G1, G1, S and G2-M) are identified by the amount of DNA in the measured cells. **c.** Accumulation of subG1 phase HeLa cells upon treatment with tubulin probes. Data were fitted to the dose response curve to obtain $IC_{50}$. Note, the maximum percentage of subG1 cells was fixed to 69% and shared between all datasets, while minimal value was not fixed. These fitting conditions allowed estimation of the probe toxicity even if no saturation was reached for probes: **6-TMR-LTX**, **5-580CP-LTX** and **6-580CP-LTX**.

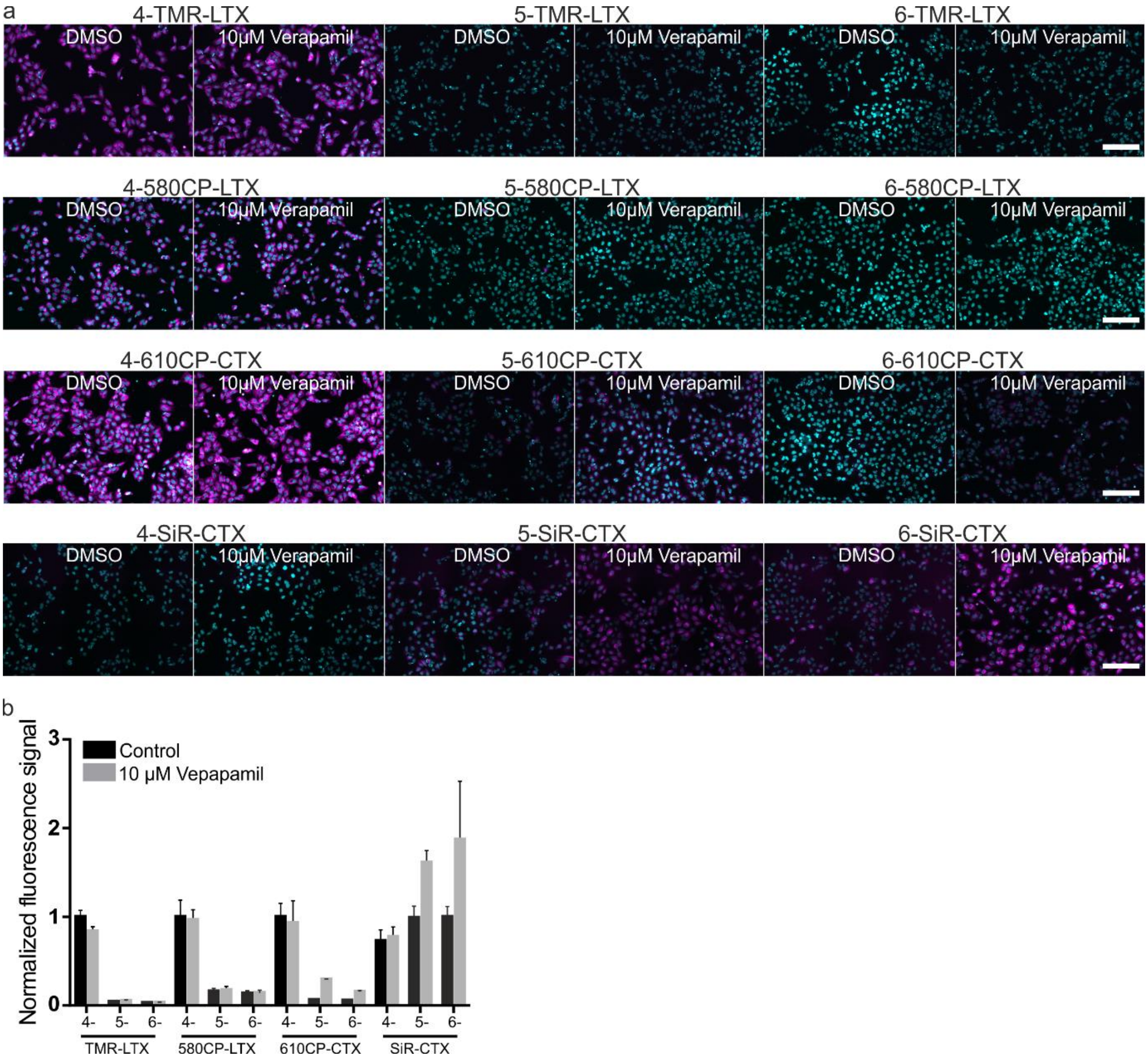

**Supplementary Figure 10.** Effect of efflux pump inhibitor (verapamil) on the staining efficiency of living U-2 OS cells with tubulin probes. **a.** Wide field fluorescence microscopy images of living U-2 OS cells stained with the fluorescent tubulin probes in the absence and presence of verapamil. Living cells were stained with 100 nM probes alone or a mixtures of probe and 10 µM Verapamil in DMEM growth medium containing 10% FBS. The cells were incubated at 37 °C for 1 h and washed once with HBSS and imaged using the same excitation powers for the six represented images: three isomers with and without Verapamil. Scale bars: 100 µm. **b.** Quantification of fluorescence signal in the cytoplasm of living cells stained with tubulin probes. Data are presented as mean ± s.d., N = 3 independent experiments, each time n > 100 cells were quantified.

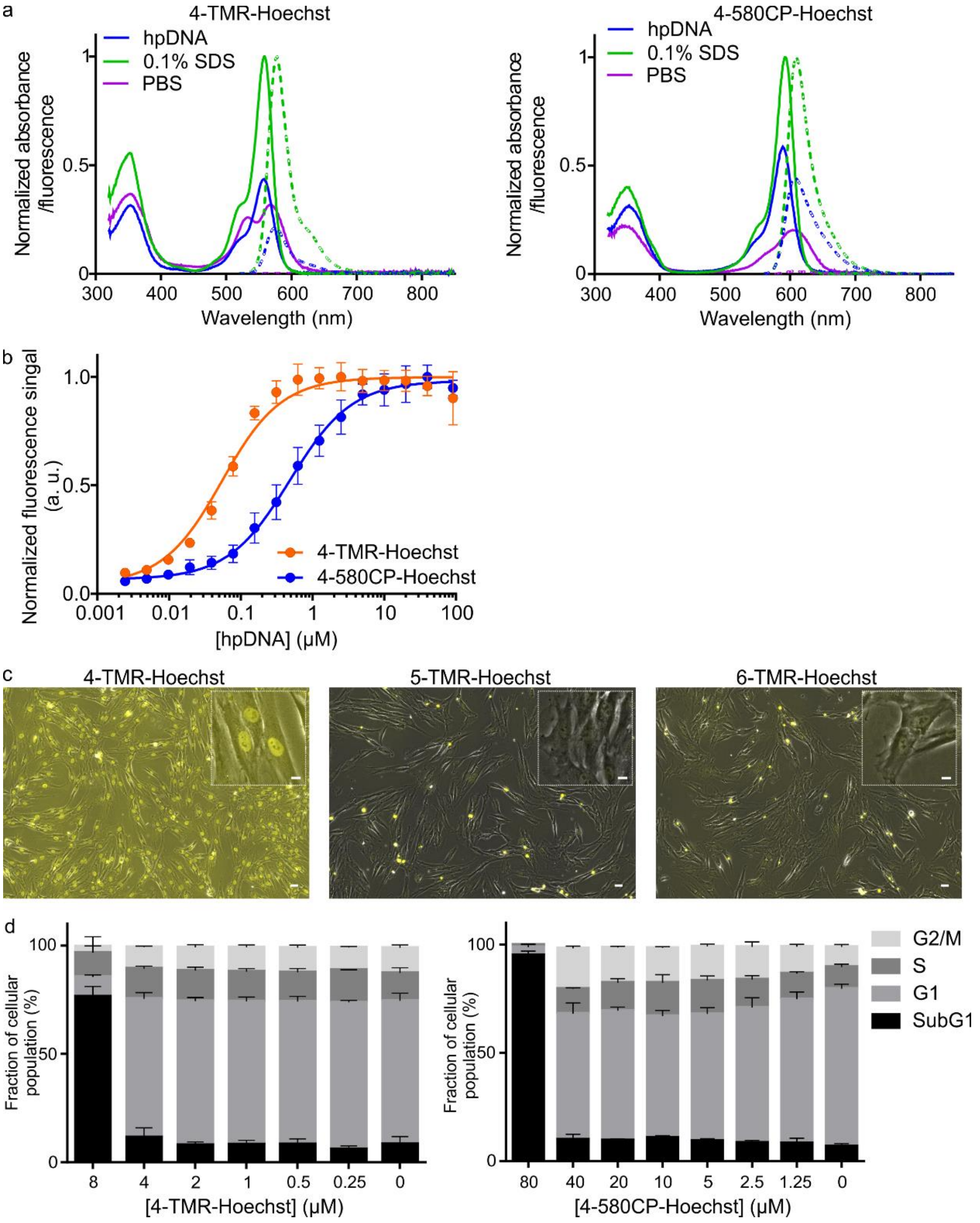

**Supplementary Figure 11.** Properties of fluorescent DNA probes. **a.** Absorption (solid line) and emission (dashed line) spectra of the DNA probes. Spectra recorded in PBS, PBS containing 0.1 % SDS or target 30 μM hpDNA. **b.** Titration of 10 nM **4-TMR-Hoechst** and **4-580CP-Hoechst** with hpDNA. The data points are fitted to a single site binding equation. Data are presented as mean values with standard deviations, N = 3 independent experiments. **c.** Wide-field microscopy images showing overlay of the light transmission (grey) and fluorescence (yellow) channels. Living primary fibroblasts were stained with 100 nM **4**/**5**/**6-TMR-Hoechst** probes for 1h at 37°C. Cells were washed once with HBSS and imaged in growth DMEM media. Inserts show zoomed-in images. Scale bars: large field of view - 30 μm, insert -10 μm. **d.** Cytotoxicity measurements of the DNA probes. HeLa cells were incubated with the indicated concentrations of the DNA probes at 37 °C for 24 h in a humidified 5% $CO_2$ incubator. Experimental data are averages of three independent experiments (N=3, n ≥ 9000 cells) and presented as means with standard deviations.

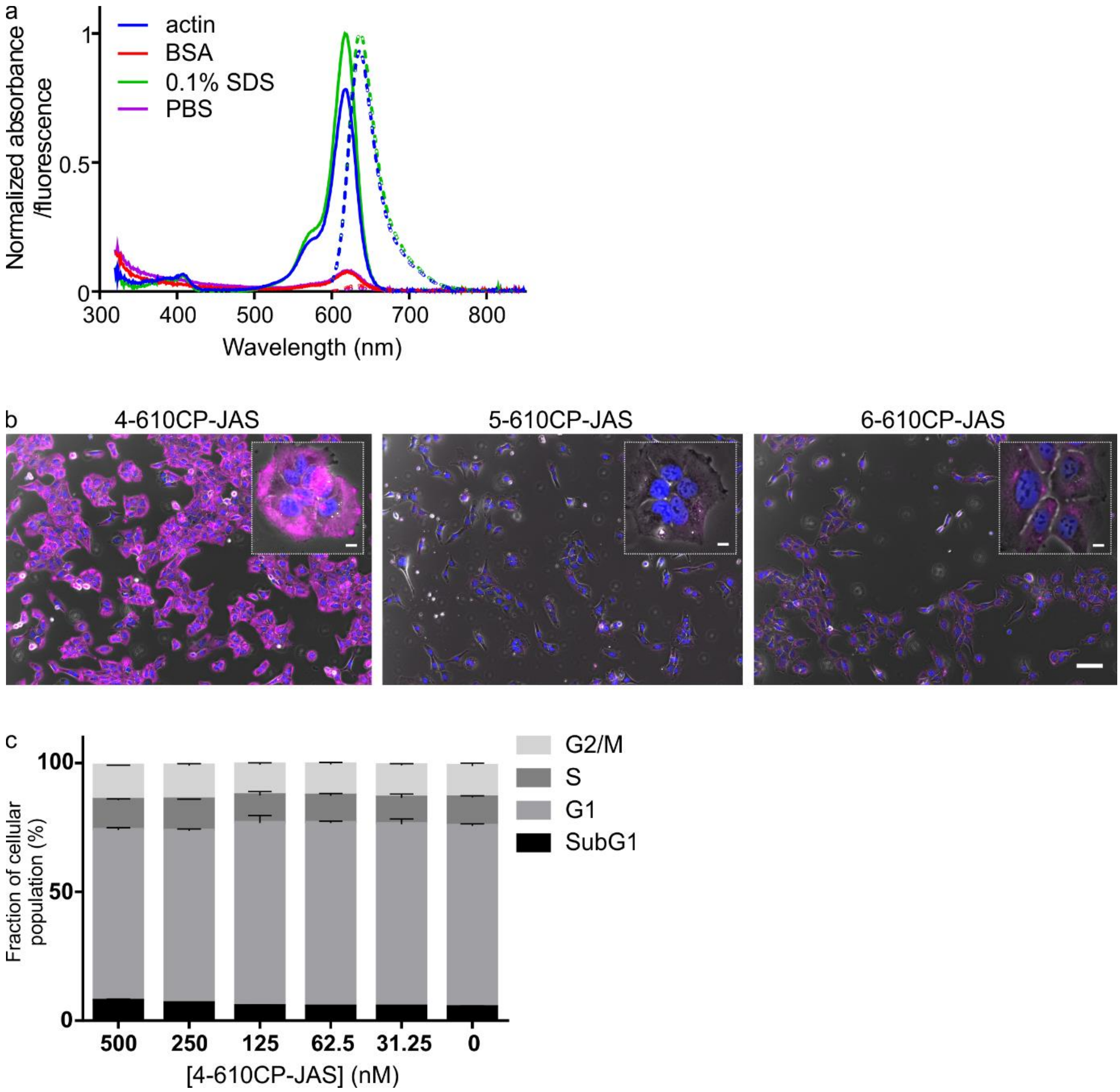

**Supplementary Figure 12.** Properties of the actin probe. **a.** Absorption (solid line) and emission (dashed line) spectra of the actin probe **4-610CP-JAS**. Spectra recorded in PBS or PBS containing 0.1 % SDS. In addition, spectra recorded in the actin polymerization buffer containing 1 mg/ml BSA or actin. **b.** Wide-field microscopy images showing overlay of the light transmission (grey), **4-610CP-JAS** (magenta) and Hoechst 33342 (blue) fluorescence channels. Living U-2 OS cells were stained with 100 nM **4**/**5**/**6-610CP-JAS** and Hoechst 33342 probes for 1h at 37°C. Cells were washed once with HBSS and imaged in growth DMEM media. Inserts show zoomed-in images. Scale bars: large field of view - 100 µm, insert -10 µm. **c.** Cytotoxicity measurements of the **4-610CP-JAS** probe. HeLa cells were incubated with the indicated concentrations of **4-610CP-JAS** at 37 °C for 24 h in a humidified 5% $CO_2$ incubator. Experimental data are averages of three independent experiments (N=3, n ≥ 9000 cells) and presented as means with standard deviations.

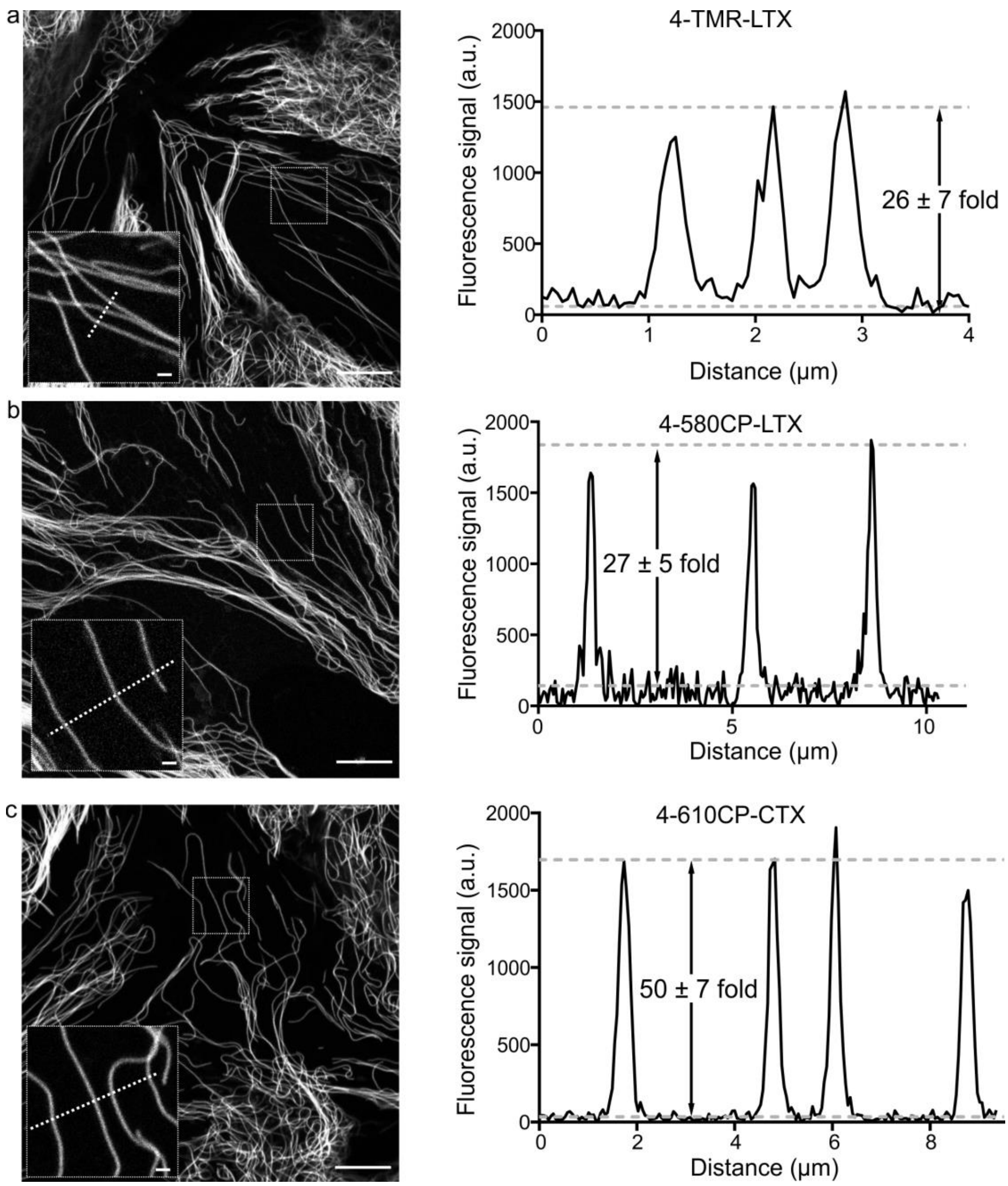

**Supplementary Figure 13.** Confocal imaging of microtubules in living human fibroblasts under no-wash conditions. **a.** Cells stained with 100 nM **4-TMR-LTX** for 1h at 37°C and directly imaged in the growth DMEM medium without probe removal. **b.** Cells stained with 100 nM **4-580CP-LTX** for 1h at 37°C and directly imaged in the growth DMEM medium without probe removal. **c.** Cells stained with 100 nM **4-610CP-CTX** for 1h at 37°C and directly imaged in the growth DMEM medium without probe removal. Images on left show confocal microtubule image, dashed square box shows the position of the zoomed-in insert. Dashed white line in the insert indicates position of the profile graph on the right. Dashed white lines in the graph on right correspond to average signal of baseline and microtubule peak values. Signal to background values are given as mean ± s.d., $N \geq 3$ separate images, $n \geq 20$ individual microtubules. Scale bars: inserts -1 μm, large fields of view - 10 μm.

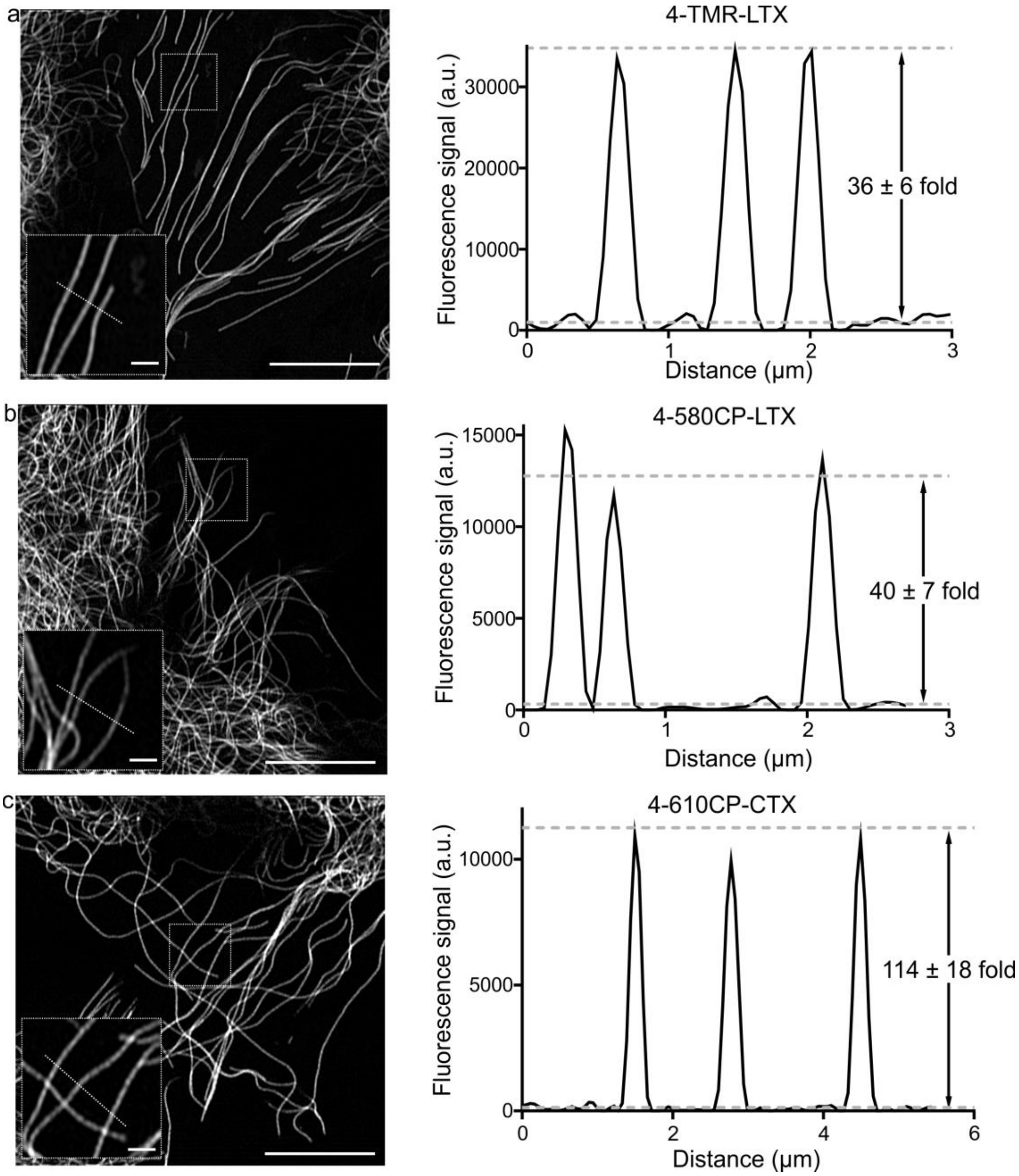

**Supplementary Figure 14.** Airyscan imaging of microtubules in living human fibroblasts under no-wash conditions. **a.** Cells stained with 100 nM **4-TMR-LTX** for 1h at 37°C and directly imaged in the growth DMEM medium without probe removal. **b.** Cells stained with 100 nM **4-580CP-LTX** for 1h at 37°C and directly imaged in the growth DMEM medium without probe removal. **c.** Cells stained with 100 nM **4-610CP-CTX** for 1h at 37°C and directly imaged in the growth DMEM medium without probe removal. Images on left show Airyscan microtubule image, dashed square box shows the position of the zoomed-in insert. Dashed white line in the insert indicates position of the profile graph on the right. Dashed white lines in the graph on right correspond to average signal of baseline and microtubule peak values. Signal to background values are given as mean ± s.d., N ≥ 3 separate images, n ≥ 20 individual microtubules. Scale bars: inserts -1 µm, large fields of view - 10 µm.

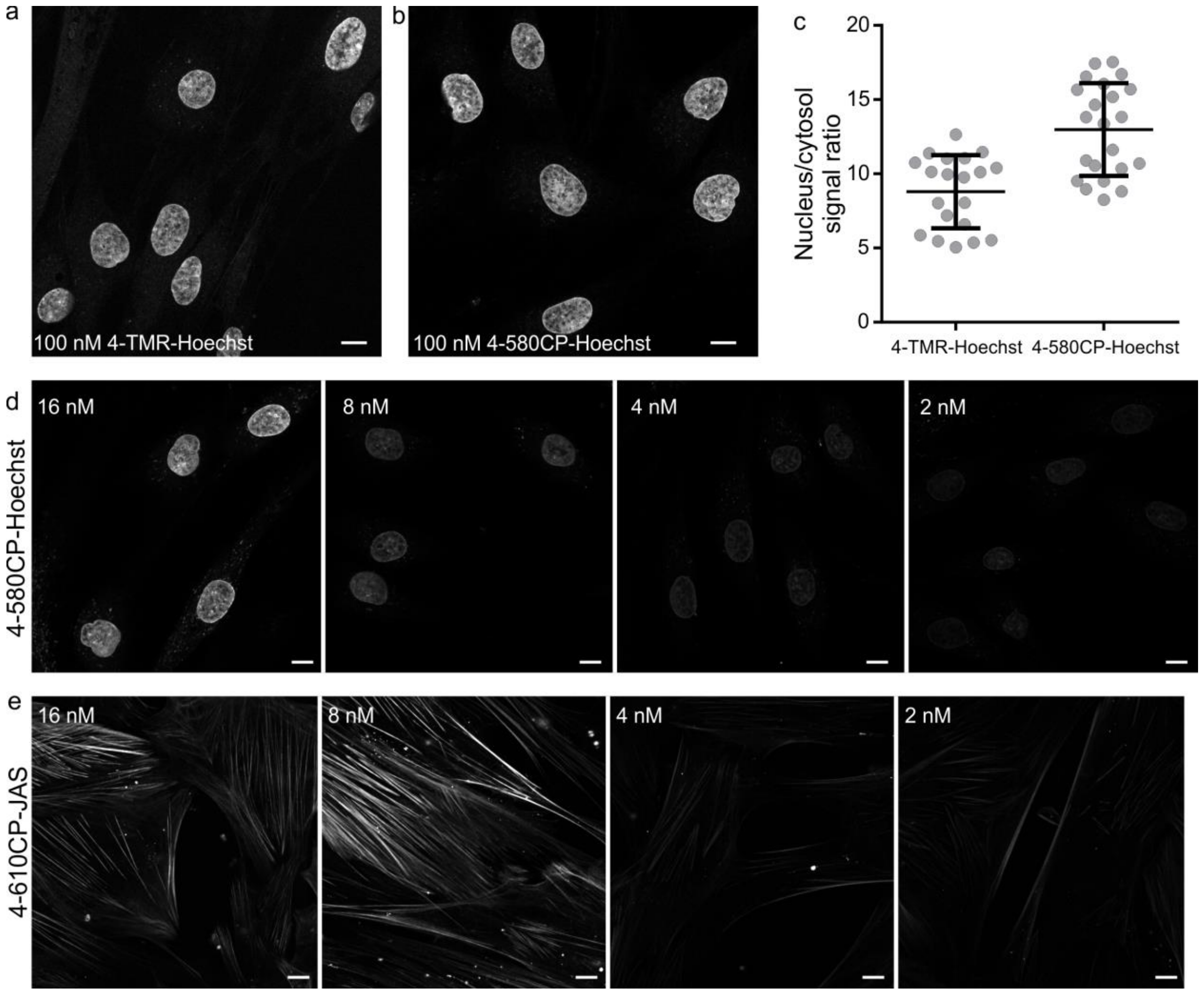

**Supplementary Figure 15.** Confocal imaging of DNA and actin in living human fibroblasts under no-wash conditions. **a.** Cells stained with 100 nM **4-TMR-Hoechst** for 1h at 37°C and directly imaged in the growth DMEM medium without probe removal. Scale bar 10 µm. **b.** Cells stained with 100 nM **4-580CP-Hoechst** for 1h at 37°C and directly imaged in the growth DMEM medium without probe removal. Scale bar 10 µm. **c.** Comparison of fluorescence signal in the nucleus and the cytosol of cells. Each grey dot represents single cell measurement and at least three independent fields of view were examined per condition. **d.** Airyscan images of living cells stained with two-fold serial dilution of **4-580CP-Hoechst** probe for 16 h at 37°C. Scale bars: 10 µm. **e.** Airyscan images of living cells stained with two-fold serial dilution of **4-610CP-JAS** probe for 16 h at 37°C. Scale bars: 10 µm.

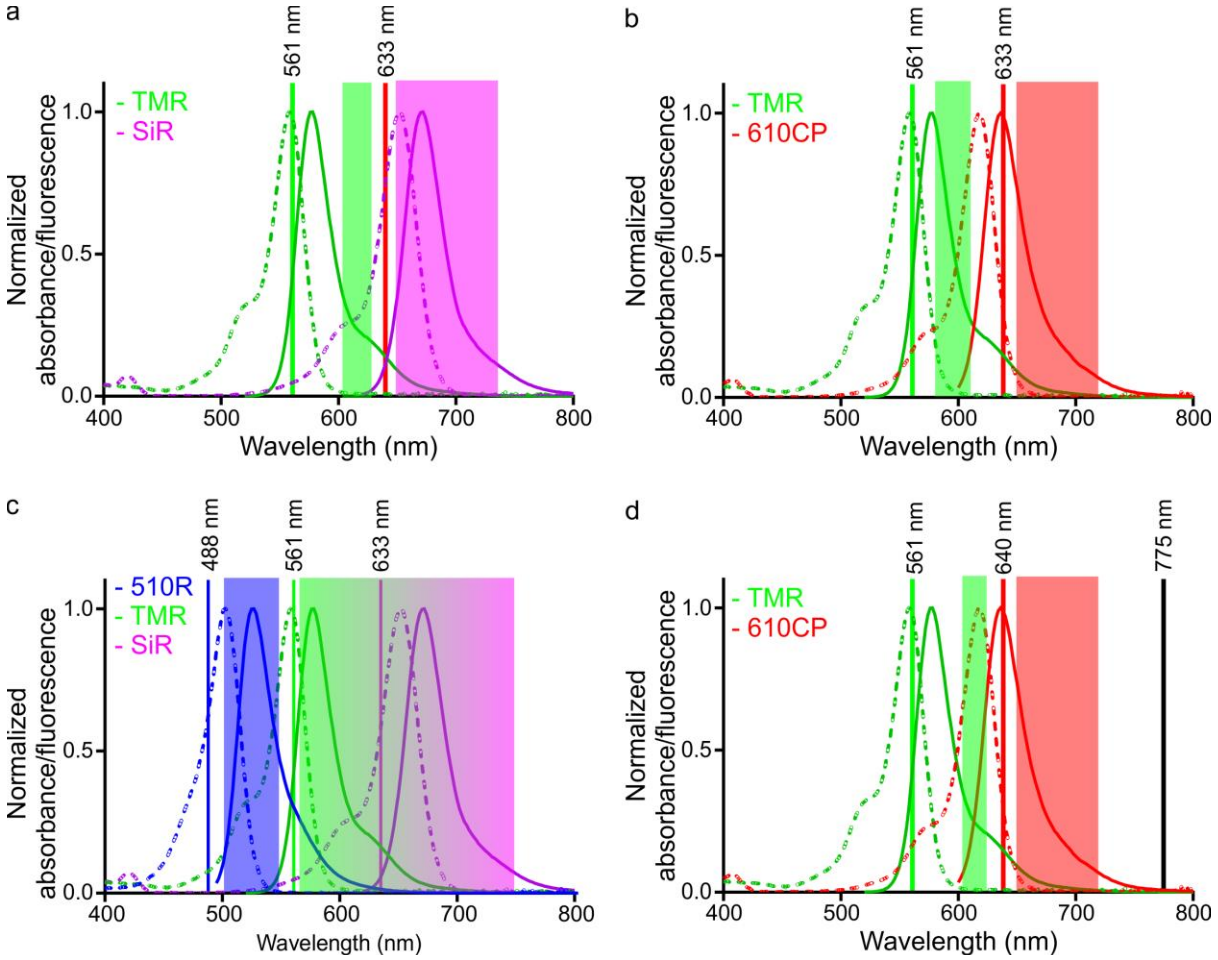

**Supplementary Figure 16.** Excitation and detection schemes used in multicolour microscopy experiments. **a.** Two-colour **TMR** and **SiR** confocal imaging **b.** Two-colour **TMR** and **SiR** confocal and Airyscan imaging. **c.** Three colour Airyscan imaging. **d.** Two-colour confocal and STED imaging. Graphs show normalized absorption and emission spectra of **510R** (blue), **TMR** (green), **610CP** (red), and **SiR** (magenta) dyes, related laser lines (solid vertical lines), and detection windows (transparent rectangles). De-excitation laser (black line) is set at 775 nm.

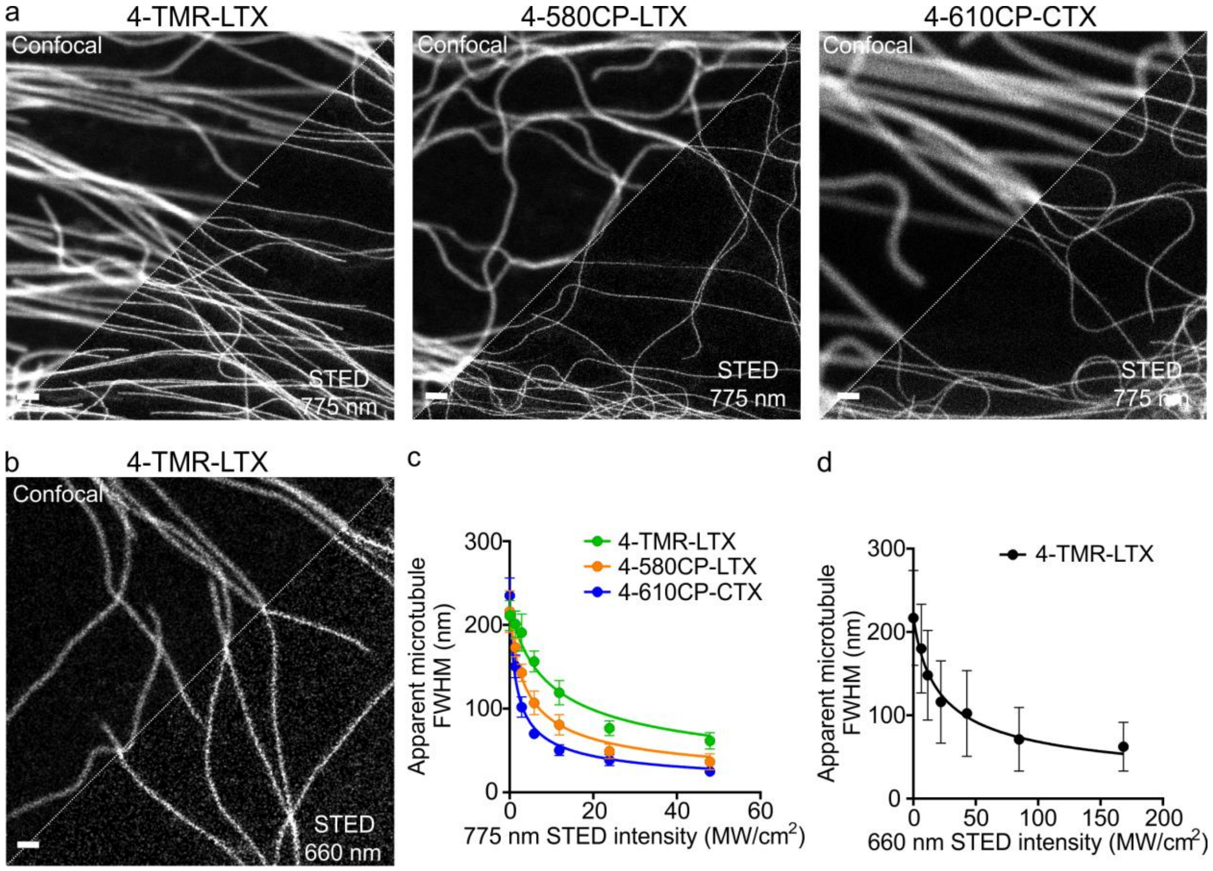

**Supplementary Figure 17.** Estimation of resolution in confocal and STED (660 and 775 nm) images of living human fibroblasts stained with tubulin probes. **a.** STED images acquired with 775 nm depletion laser . Living human fibroblasts were incubated with 100nM of the indicated tubulin probe in growth medium containing 10% FBS at 37 °C for 1 h, washed once with HBSS and imaged. Images were acquired on Abberior STED 775 QUAD scanning microscope equipped with 775 nm STED laser. Scale bars: 1 µm. **b.** STED image acquired with 660 nm depletion laser. Living human fibroblasts were incubated with the 100 nM **4-TMR-LTX** in growth medium containing 10% FBS at 37 °C for 1 h, washed once with HBSS and imaged. Images were acquired on a Leica TCS SP8 STED scanning microscope equipped with 660 nm STED laser. Scale bars: 1 µm. **c.** The apparent microtubule FWHM as a function of the 775 nm STED laser intensity used for imaging of specimens stained with **4-TMR-LTX** (green), **4-580CP-LTX** (blue) and **4-610CP-CTX** (orange). Data are presented as mean values with standard deviations, N ≥ 10 microtubules in at least 3 different fields of view. d. The apparent microtubule FWHM as a function of the 660nm STED laser intensity used for imaging of specimens stained with **4-TMR-LTX**. Data are presented as mean values with standard deviations, N ≥ 10 microtubules in at least 3 different fields of view.

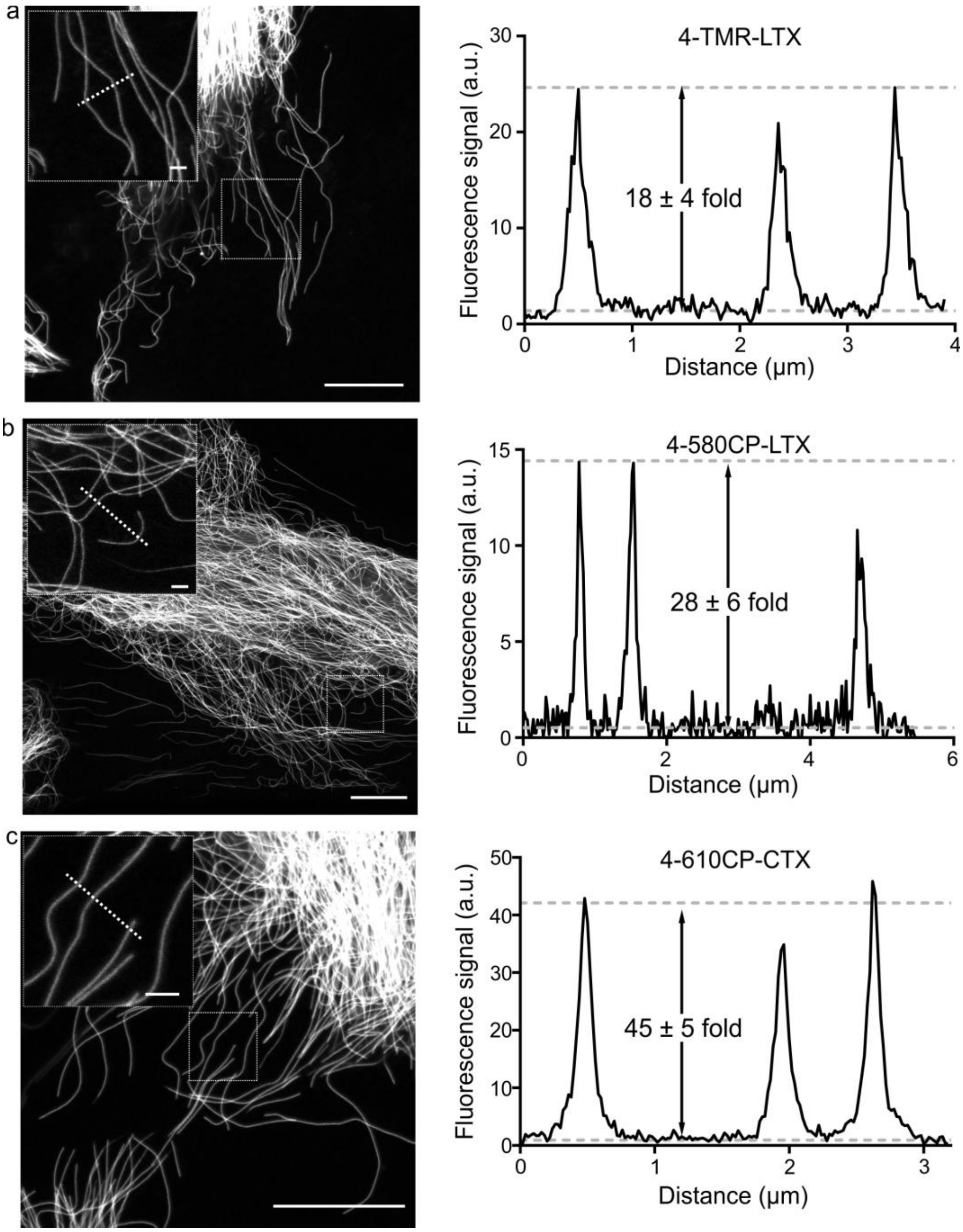

**Supplementary Figure 18.** STED imaging of microtubules in living human fibroblasts under no-wash conditions. **a.** Cells stained with 100 nM **4-TMR-LTX** for 1h at 37°C and directly imaged in the growth DMEM medium without probe removal. **b.** Cells stained with 100 nM **4-580CP-LTX** for 1h at 37°C and directly imaged in the growth DMEM medium without probe removal. **c.** Cells stained with 100 nM **4-610CP-CTX** for 1h at 37°C and directly imaged in the growth DMEM medium without probe removal. Images on left show STED microtubule image, dashed square box shows the position of the zoomed-in insert. Dashed white line in the insert indicates position of the profile graph on the right. Dashed white lines in the graph of right correspond to average signal of baseline and microtubule peak values. Signal to background values are given as mean ± s.d., N ≥ 3 separate images, n ≥ 20 individual microtubules. Scale bars: inserts -1 µm, large fields of view - 10 µm.

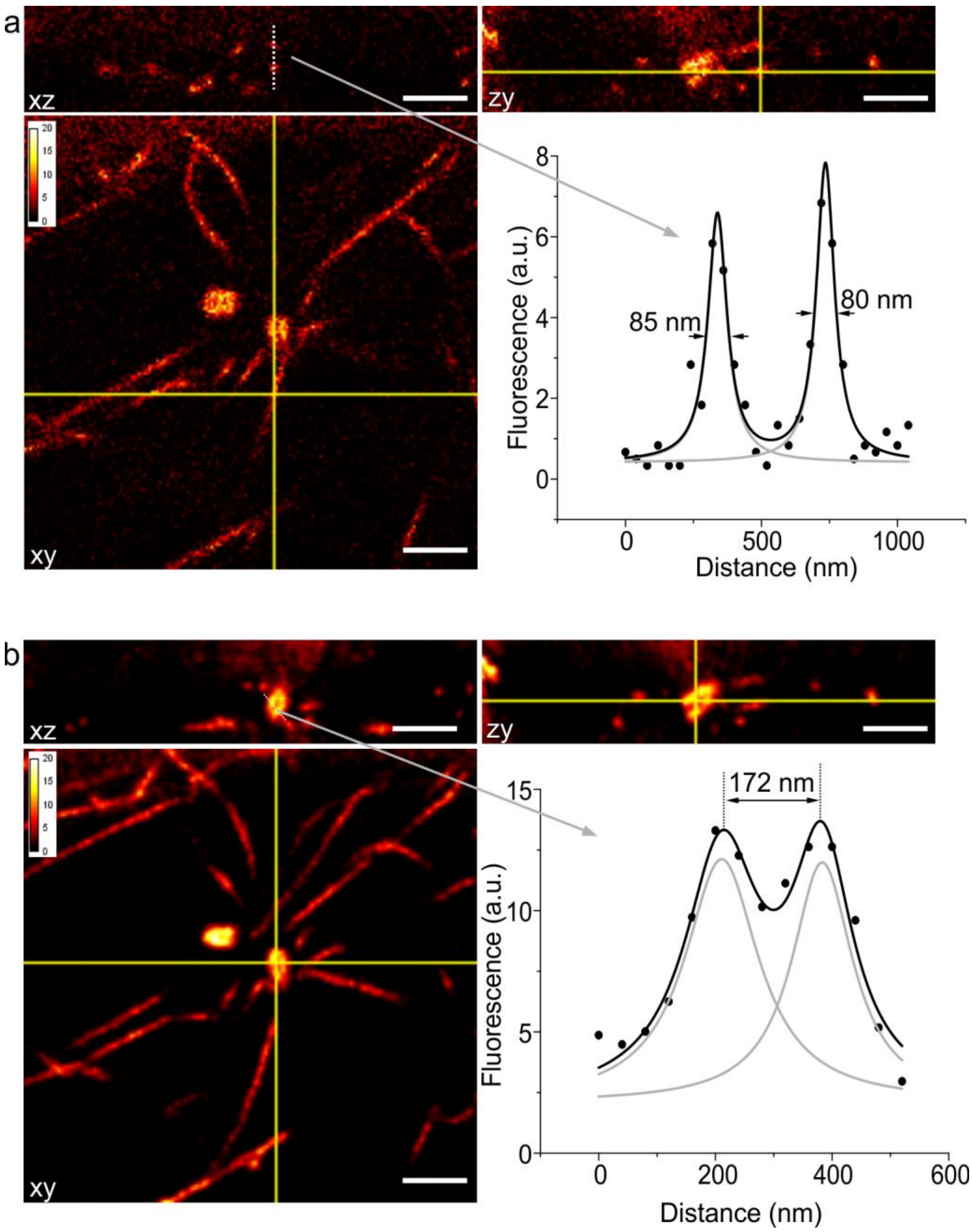

**Supplementary Figure 19**. Isotropic 3D STED images of microtubules in living human fibroblasts under no-wash conditions. **a.** Panel shows raw data of cells stained with 100 nM **4-610CP-CTX** for 1h at 37°C and directly imaged in the growth DMEM medium without probe removal. **b.** Panel shows deconvolved image of cells stained with 100 nM **4-610CP-CTX** for 1h at 37°C and directly imaged in the growth DMEM medium without probe removal. Deconvolution performed using SVI Huygens software. Dashed white line in the insert indicates position of the profile graph. Black line in the graph corresponds to sum of the two Lorentz distributions indicated in gray line. Yellow solid lines indicate position of xz, xy and zy sections. Voxel size 40 x 40 x 40 nm. Scale bars: 1 µm.

#### Supplementary Movies

**Supplementary Movie 1**. Time-lapse Airyscan movie of a living fibroblast stained with 100 nM **4-610CP-JAS**. Imaging performed without washing step after labeling. Scale bar 10 µm.

**Supplementary Movie 2.** Time-lapse Airyscan movie of a living fibroblast stained with 0.06 nM **4-TMR-LTX** and 20 nM **5-SiR-Hoechst**. Imaging performed without washing step after labeling. Scale bar 5 µm.

**Supplementary Movie 3.** Long-term time-lapse movie of a living fibroblast stained with 100nM **4-TMR-Hoechst** and 10nM **6-SiR-CTX**. Imaging performed without washing step after labeling. Scale bar 5 µm.

**Supplementary Movie 4.** Rotating maximum intensity projection of three-color ZEISS Airyscan image of a living HeLa cell at metaphase stained with 3 nM **4-TMR-LTX** (green), 20 nM **5-SiR-Hoechst** (red) and 1000 nM **6-510R-JAS** (yellow). Scale bar 1 µm.

**Supplementary Movie 5.** Confocal and STED comparative timecourse of a living fibroblast stained with 100 nM **4-610CP-CTX**. Imaging performed without washing step after labeling. Scale bar 5 µm.

**Supplementary Movie 6.** Rotating maximum intensity projection of tubulin network 3D STED image. Living fibroblast stained with 100 nM 4-610CP-CTX and imaged without washing off the probe. Centrosome structure is visible at the centre of the image. Tuble-like centrosiole structure is clrealy visible. Scale bar 5 µm.

### Supplementary Tables

**Supplementary Table 1**. Photophysical properties of rhodamine fluorescent dyes used in the study.

| Dye | Solvent | $\lambda_{max}^{abs}$ (nm) | $\lambda_{max}^{em}$ (nm) | ε (m$^{-1}$ cm$^{-1}$) | QY | τ (ns) |
|---|---|---|---|---|---|---|
| **4-TMR-COOH** | PBS | 551 | 573 | 77600 ± 5000 | 0.409 ± 0.01 | 2.04 ±0.04 |
| | PBS + 0.1%SDS | 554 | 575 | 77800 ± 4500 | 0.587 ± 0.005 | 2.99 ±0.01 |
| **5-TMR-COOH** | PBS | 550 | 579 | 83000 ± 1600 | 0.411 ± 0.005 | 2.14 ± 0.02 |
| | PBS + 0.1%SDS | 552 | 580 | 83600 ± 1100 | 0.51 ± 0.02 | 2.80 ± 0.02 |
| **6-TMR-COOH** | PBS | 550 | 576 | 83300 ± 4200 | 0.359 ± 0.003 | 1.97 ± 0.02 |
| | PBS + 0.1%SDS | 550 | 576 | 86600 ± 4800 | 0.399 ± 0.002 | 2.05± 0.02 |
| **4-580CP-COOH** | PBS | 585 | 609 | 108000 ± 2900 | 0.597 ± 0.02 | 3.55 ± 0.01 |
| | PBS + 0.1%SDS | 591 | 613 | 117000 ± 5500 | 0.660 ± 0.01 | 4.2 ± 0.01 |
| **5-580CP-COOH** | PBS | 582 | 610 | 100400 ± 6100 | 0.563 ± 0.004 | 3.47± 0.01 |
| | PBS + 0.1%SDS | 586 | 613 | 104600 ± 5400 | 0.650 ± 0.003 | 4.07 ± 0.01 |
| **6-580CP-COOH** | PBS | 582 | 608 | 100300 ± 2500 | 0.584 ± 0.008 | 3.64 ± 0.03 |
| | PBS + 0.1%SDS | 583 | 609 | 100200 ± 4400 | 0.614 ± 0.006 | 3.63 ± 0.66 |
| **4-610CP-COOH** | PBS | 611 | 636 | 101400 ± 8400 | 0.497 ± 0.01 | 3.12 ± 0.01 |
| | PBS + 0.1%SDS | 616 | 640 | 105500 ± 9400 | 0.668 ± 0.01 | 4.23 ± 0.01 |
| **5-610CP-COOH** | PBS | 608 | 636 | 98200 ± 4000 | 0.471 ± 0.003 | 2.99 ± 0.01 |
| | PBS + 0.1%SDS | 608 | 637 | 106000 ± 5500 | 0.48 ± 0.01 | 2.99 ± 0.02 |
| **6-610CP-COOH** | PBS | 608 | 635 | 102000 ± 2500 | 0.483 ± 0.007 | 2.97 ± 0.01 |
| | PBS + 0.1%SDS | 609 | 636 | 101000 ± 2300 | 0.541 ± 0.005 | 3.11 ± 0.02 |
| **4-SiR-COOH** | MeOH+0.1%TFA | 656 | 677 | 108500 ± 2000 | 0.519 ± 0.004 | 3.46 ± 0.03 |
| | PBS | 649 | 669 | 34200 ± 2700 | 0.433 ±0.024 | 2.77 ± 0.01 |
| | PBS + 0.1%SDS | 655 | 674 | 73300 ± 5100 | 0.619 ± 0.006 | 3.95 ± 0.03 |
| **5-SiR-COOH** | PBS | 645 | 670 | 97000 ± 1900 | 0.399 ± 0.005 | 2.61 ± 0.01 |
| | PBS + 0.1%SDS | 649 | 672 | 101000 ± 2000 | 0.595 ± 0.005 | 3.55 ± 0.01 |
| **6-SiR-COOH** | PBS | 645 | 668 | 99600 ± 5000 | 0.406 ± 0.007 | 2.69 ± 0.01 |
| | PBS + 0.1%SDS | 646 | 668 | 90000 ± 4000 | 0.459 ± 0.003 | 2.82 ± 0.01 |

**Supplementary Table 2**. $^{dye}D_{50}$ values for the fluorescent dyes.

| Fluorophore | TMR-COOH | 580CP-COOH | 610CP-COOH | SiR-COOH |
|---|---|---|---|---|
| **4'-isomer** | 14 ± 1 | 46 ± 2 | 32.1 ± 0.7 | 78.6 ± 0.2 |
| **5'-isomer** | 11.7 ± 0.7 | 39 ± 2 | 33.5 ± 0.6 | 68.8± 0.4 |
| **6'-isomer** | 9.3 ± 0.4 | 33 ± 2 | 30.7 ± 0.6 | 65.4 ± 0.2 |

**Note:** Data presented as fitted value ± standard error. Fitting to dose-response equation performed using GraphPad Prism 6.0.

**Supplementary Table 3**. $^{probe}D_{50}$ and $^{probe}A_{50}$ values for the fluorescent tubulin probes.

| | 1,4-Dioxane-Water | | 1,4-Dioxane-Water with constant 0.3%SDS additive | | | |
|---|---|---|---|---|---|---|
| **Probe** | **TMR-LTX** | | **TMR-LTX** | **580CP-LTX** | **610CP-CTX** | **SiR-CTX** |
| | $D_{50}$ | $A_{50}$ | $D_{50}$ | $D_{50}$ | $D_{50}$ | $D_{50}$ |
| **4'-isomer** | 23.4 ± 0.3 | 82.2 ± 0.7 | 26.5 ± 0.5 | 62 ± 1 | 65 ± 2 | >80 |
| **5'-isomer** | 12.3 ± 0.5 | 79 ± 1 | 13.0 ± 0.4 | 40.0 ± 0.6 | 39.7 ± 0.2 | 66.8 ± 0.3 |
| **6'-isomer** | 13.1 ± 0.7 | 81 ± 2 | 11.2 ± 0.5 | 36.0 ± 0.7 | 36.4 ± 0.4 | 66.5 ± 0.4 |

**Note:** Data presented as fitted value ± standard error. TMR-LTX titration without SDS data points were fitted to bell-shaped dose-response equation performed using GraphPad Prism 6.0.

**Supplementary Table 4.** Calculated total potential energies of spirolactone and zwitterion forms of model isomeric rhodamines in water and 1,4-dioxane environment.

| | Form | E ( in 1,4-Dioxane), Hartrees | E (in Water), Hartrees |
|---|---|---|---|
| **6-TMR-NHMe** | Spirolactone | -1471.396820 | -1471.412378 |
| | Zwitterion | -1471.383652 | -1471.415182 |
| **5-TMR-NHMe** | Spirolactone | -1471.396150 | -1471.412729 |
| | Zwitterion | -1471.382063 | -1471.415307 |
| **4-TMR-NHMe** | Spirolactone | -1471.399407 | -1471.411604 |
| | Zwitterion | -1471.383782 | -1471.412636 |

**Supplementary Table 5**. Chemical shift of amide NH proton in $d_6$-DMSO of isomeric dye-C8-taxane conjugates.

| Probe | Conjugate isomer | | |
|---|---|---|---|
| | 4'-isomer | 5'-isomer | 6'-isomer |
| **TMR-LTX** | 9.07 | 8.76 | 8.62 |
| **580CP-LTX** | 9.14 | 8.72 | 8.69[1] |
| **610CP-CTX** | 9.09 | 8.73 | 8.66 |
| **SiR-CTX** | 8.99 | 8.75 | 8.69[1] |
| **Average** | 9.07 ± 0.06 | 8.74 ± 0.01 | 8.67 ± 0.03 |

**Supplementary Table 6**. Retention times on HPLC $C_{18}$ column of the tubulin probes.

| Probe | TMR-LTX | 580CP-LTX | 610CP-CTX | SiR-CTX |
|---|---|---|---|---|
| **4'-isomer** | 81.1 ± 0.6 s | 132 ± 2 s | 699 ± 4 s | 1403 ± 9 s |
| **5'-isomer** | 53.1 ± 0.5 s | 45.9 ± 0.3 s | 114.3 ± 0.4 s | 683 ± 4 s |
| **6'-isomer** | 39.6 ± 0.5 s | 33.4 ±0.5 s | 76.1 ± 0.5 s | 471 ± 2 s |

**Supplementary Table 7**. Properties of the best performing probes.

| Probe | $\lambda_{max}^{abs}$ (nm) | $\lambda_{max}^{em}$ (nm) | ε•10³ ($M^{-1}cm^{-1}$) | QY | $I_{sat}^{STED}$ ($MWcm^{-1}$) | Brightness ($M^{-1}cm^{-1}$) | $Fl_{target}$ increase | Toxicity $EC_{50}$ (nM) |
|---|---|---|---|---|---|---|---|---|
| | | | | **Tubulin probes** | | | | |
| **4-TMR-LTX** | 557 | 576 | 65 ± 14 | 0.56 ± 0.02[a] | 4.91±0.69 | 36100 ± 8100 | 2.3 ± 0.4 | 6.0 ± 0.6 |
| **4-580CP-LTX** | 591 | 612 | 72 ± 11 | 0.63 ± 0.01[a] | 2.29±0.14 | 45400 ± 6700 | 13 ± 3 | 27 ± 2 |
| **4-610CP-CTX** | 617 | 638 | 99 ± 10 | 0.73 ± 0.02[a] | 2.75±0.28 | 72000 ± 7100 | 8 ± 1 | 14 ± 1 |
| **6-SiR-CTX** | 652 | 670 | 39 ± 3 | 0.74 ± 0.05[a] | 1.0 ± 0.1[c] | 28500 ± 2300 | 178 ± 61 | 235 ± 15 |
| | | | | **DNA probes** | | | | |
| **4-TMR-Hoechst** | 558 | 576 | 34 ± 3 | 0.27 ± 0.01[b] | - | 9100 ± 900 | 87 ± 7 | > 4000 |
| **4-580CP-Hoechst** | 589 | 610 | 67 ± 4 | 0.51 ± 0.01[b] | - | 34400 ± 1800 | 34 ± 1 | > 1000 |
| | | | | **Actin probe** | | | | |
| **4-610CP-JAS** | 618 | 636 | 83 ± 4 | 0.67 ± 0.04[a] | - | 55300 ± 2500 | 38 ± 2 | > 500 |

[a] Relative quantum yield. [b] Absolute quantum yield. [c1].

#### Computation, molecular biology and biochemical methods

*Preparation of hairpin DNA*

For the DNA binding studies hairpin forming oligonucleotide 5'-CGCGAATTCGCGTTTTCGCGAATTCGCG-3' (28 bp) was purchased from Sigma-Aldrich. Previously, this hairpin DNA has been used for structural studies of the interaction of Hoechst 33342 with DNA[2]. Synthetic oligonucleotides were dissolved in PBS (Lonza, pH 7.4) at 1 mM concentration. Hairpin was formed by putting the tube with hpDNA solution into a boiling water bath which was then slowly cooled down to room temperature.

*Measurements of absorbance spectra in 1,4–dioxane-water mixtures*

Measurements of the absorbance changes in 1,4-dioxane-water mixtures were performed by pipetting 2µL stock solutions of dyes or probes in DMSO into a 96 glass bottom well plate (11 wells per sample) made from propylene (Corning 3364). To the wells going from right to left 300 µL of 1,4-dioxane-water mixtures containing 100%, 90%, 80%, 70%, 60%, 50%, 40%, 30%, 20%, 10% or 0% 1,4-dioxane was added (if needed, mixtures with 0.3% SDS are used, with an exception in 100% dioxane due to solubility issues). After incubation for 1 hour at room temperature, absorption of solutions in each well was recorded from 320 nm to 850 nm with wavelength step size of 1 nm on a multiwell plate reader Spark® 20M (Tecan). The background absorption of the glass bottom plate was measured in wells containing only 1,4–dioxane-water mixture with similar amount of DMSO and subtracted from the spectra of the samples. Plots of $\lambda_{max}$ versus dielectric constant (D) of 1,4-dioxane-water mixtures[3]. $D_{50}$ value was obtained by fitting to dose-response equation $EC_{50}$ as implemented in GraphPad 6.0 software:

$$A = A_0 + (A_{max} - A_0)/\left(1 + \left(\frac{D_{50}}{d}\right)^{Hill}\right) \qquad (1)$$

where $A_o$ – absorbance at $\lambda_{max}$ at $\varepsilon_r = 0$, $A_{max}$ – the highest reached absorbance at $\lambda_{max}$. $d$ – dielectric constant of 1,4-dioxane-water mixture at a given point, *Hill* - Hill slope coefficient determining the steepness of a dose-response curve, $D_{50}$ - corresponds to d value that provokes half of the absorbance amplitude ($A_{max}$-$A_0$).

Data points from the **4-TMR-LTX** titration were fitted to bell-shaped dose response curve described by following equation:

$$A = A_0 + \frac{A_{max} - A_0}{1 + \left(\frac{D_{50}}{d}\right)^{Hill_1}} - \frac{A_{max} - A_0}{1 + \left(\frac{A_{50}}{d}\right)^{Hill_2}} \qquad (2)$$

where $A_o$ – absorbance at $\lambda_{max}$ at $\varepsilon_r = 0$, $A_{max}$ – the highest reached absorbance at $\lambda_{max}$ during the titration experiment. $d$ – dielectric constant of 1,4-dioxane-water mixture at a given point, $Hill_1$ - Hill slope coefficient determining the steepness of the ascending dose-response curve part, $Hill_2$ - Hill slope coefficient determining the steepness of the declining dose-response curve part, $D_{50}$ - corresponds to d value that provokes half of the absorbance amplitude ($A_{max}$-$A_0$) in the ascending dose-response curve part, $A_{50}$ - corresponds to d value that provokes half of the absorbance amplitude ($A_{max}$-$A_0$) in the declining dose-response curve part.

*Determination of HPLC retention times*

HPLC retention times of the analysed probes were determined by performing analysis on an Agilent 1260 Infinity II LC/MS system equipped with an autosampler, diode array detector WR, fluorescence detector Spectra and Infinity Lab LC/MSD 6100 series quadruple with API electrospray. Analysis was done under isocratic elution conditions by using SUPELCO Titan C18, 2.1 x 75 mm, 1.9 µm threaded column, elution performed by pumping solvent with one pump with premixed 75% : 25% MeOH: 25 mM $HCOONH_4$ (pH = 3.6) aqueous buffer under the 0.4 mL/min flow, column was thermostated at 45°C.

*Determination of Quantum Yields and Lifetimes*

The fluorescence quantum yields (absolute values) were obtained with a Quantaurus-QY absolute PL quantum yield spectrometer (model C11347-12, Hamamatsu) according to the manufacturer's instructions. Fluorescence lifetimes were measured with a Quantaurus-Tau fluorescence lifetime spectrometer (model C11367-32, Hamamatsu) according to the manufacturer's instructions.

Relative quantum yields of the probes bound to the target were calculated by recording absorbance and fluorescence spectra using a multiwell plate reader Spark® 20M (Tecan) and glass bottom 96-well plates at room temperature (25 °C). To account for background due to light scattering, spectra of the solutions containing no probes, but equivalent amount of DMSO, were acquired and subtracted from the respective probe spectra. Absolute quantum yields of probes were measured for SDS sample as described using Quantaurus-QY absolute PL quantum yield spectrometer. The experiment was repeated three times, and spectra were averaged. Relative quantum yields ($QY_{\mathrm{target}}$) of the probes bound to the target were calculated in a|e - UV-Vis-IR Spectral Software v2.2 (FluorTools) using the following formula:

$$QY_{target} = QY_{SDS} \cdot \frac{A_{SDS} \cdot Fl_{target}}{Fl_{SDS} \cdot A_{target}}. \qquad (3)$$

where $QY_{SDS}$- absolute quantum yield of the probe dissolved in PBS containing 0.2 % SDS, $A_{SDS}$ – absorbance of the probe solution in PBS containing 0.2 % SDS at $\lambda_{\mathrm{max}}$, $A_{target}$ – absorbance of the probe in the solution containing excess of the target, $Fl_{SDS}$ – integrated fluorescence intensity of the probe solution in PBS containing 0.2 % SDS at $\lambda_{\mathrm{max}}$, $Fl_{target}$ – integrated fluorescence intensity of the probe in the solution containing excess of the target.

*Estimation of absorbance and fluorescence increase upon target binding or SDS addition*

Fluorescence increase of 2 µM tubulin probes upon 1mg/ml BSA (Sigma, Cat. No. A7030), tubulin (Cytoskeleton, Inc. Cat. No. T240) binding or 0.1% (w/v) SDS (Acros Organics) addition was measured in the tubulin polymerization buffer (80 mM PIPES pH 6.9, 2 mM MgCl2, 0.5 mM EGTA) supplemented with 1 mM GTP (Thermo Scientific, Cat. No. R0461). Samples prepared in a half-area 96-well black plate (Greiner Bio-One Cat.675076), mixed and incubated at 37ºC for 3-5 h before measurements.

Absorbance and fluorescence increase of 2 µM actin probes upon interaction with 1mg/ml BSA or actin (Cytoskeleton, Inc. Cat. No AKL95) measured in the actin polymerization buffer (Cytoskeleton, Inc. Cat. No. BSA02) supplemented with 0.2 mM ATP (Cytoskeleton, Inc. Cat. No. BSA04). In parallel, fluorescence increase of 2 µM actin probe solution in PBS buffer (Lonza, Cat. No. BE17-516F) with or without 0.1% SDS was estimated. Samples prepared in a glass bottom 96-well plate (MatTek, Cat. No. PBK96G-1.5-5-F), mixed and and incubated at 37ºC for 3-5 h before measurements.

Absorbance and fluorescence increase of Hoechst-based probes binding to hpDNA was estimated using the following procedure: the probe from 1 mM DMSO stock solution was diluted to the final concentration of 2 µM in PBS buffer containing 30 µM of hpDNA (5’-CGCGAATTCGCGTTTTCGCGAATTCGCG-3’).

Absorption and fluorescence were measured on a multiwell plate reader Spark® 20M (Tecan) in glass bottom 96-well plates at room temperature (25 °C). Only fluorescence was measured for tubulin probes. Absorption of solutions was recorded from 320 nm to 850 nm with wavelength step size of 1 nm. The background absorption of the glass bottom plate was measured in wells containing only buffer with similar amount of DMSO and subtracted from the spectra of the samples. Fluorescence emission of the free dyes or final probes was recorded from 520 nm to 850 nm (for **TMR**, 495 nm exc., bandwidth 15 nm), 560 nm to 850 nm (for **580CP**, 530 nm exc., bandwidth 15 nm), 600 nm to 850 nm (for **610CP**, 570 nm exc., bandwidth 15 nm), 620 nm to 850 nm (for **SiR**, 595 nm exc., bandwidth 15 nm) with 5 nm emission bandwidth and 2 nm step size.

All samples were prepared in technical triplicates, which were repeated three times as three independent experiments performed on different days.

*Determination of $K_d$*

$K_d$ measurements were performed by titrating DNA probes based on Hoechst in PBS (Lonza) with increasing concentrations of the 28 bp hpDNA in a 96-well plate and measuring the increase in fluorescence on a plate reader after 1 h incubation at room temperature. **4-TMR-Hoechst** and **4-580CP-Hoechst** were excited at 540 nm and 570 nm while recording emission at 580 nm and 610 nm, respectively. The excitation and emission bandwidth was 15 nm. The $K_d$ values were determined by plotting the emission signal vs hpDNA concentration and fitting the curve in GraphPad Prism 6 to “Single site binding” function:

$$Y = F\text{min} + (F\text{max} - F\text{min}) * \frac{(p+X+K_d^{\text{app}}) - \sqrt{(p+X+K_d)^2 - 4*p*X}}{2*p} \quad (4)$$

where $F_{min}$ – fluorescence of probe without target, $F_{max}$ – fluorescence of probe at saturating concentration, $p$ – probe concentration, X – target hpDNA concentration, $K_d$ – dissociation constant of the probe. All measurements performed 3 times on different days, each time technical triplicates were measured.

*In vitro tubulin polymerization assay*

Measurements of the polymeric tubulin stabilization by the tested probes were performed using a commercial tubulin polymerization fluorescence assay kit from Cytoskeleton, Inc. (cat. BK011P). Taxanes are known to stabilize tubulin in the polymerized state resulting in an increased apparent polymerization rate. Immediately before measurement, 0.5 mg porcine brain tubulin was dissolved in 1 ml buffer (80 mM PIPES pH 6.9, 2 mM $MgCl_2$, 0.5 mM EGTA, 10 µM DAPI), supplemented with 1 mM GTP. Probes were diluted in water to 30 µM, and an aliquots of 5 µl were added into wells of a black half-area 96-well plate. Control samples contained 1:32 dilution of DMSO in water. The polymerization reaction was started by quickly adding 50 µl of tubulin stock using automatic dispenser. The plate was placed into a plate-reader pre-warmed to 37°C and the kinetic fluorescence readings were started immediately. Tubulin polymerization was followed by increase in DAPI fluorescence, using Tecan Spark20M plate reader, set to 350 nm excitation and 450 nm emission with 20 nm bandwidth in both cases. Data points from 3-5 independent kinetic traces were globally fitted into plateau followed by one-phase association function:

$$y=y_0 + (\text{plateau}-y_0)\times(1-e\hat{}(-K \times (t-t_0))) \quad (5),$$

where $t_0$ is the time when the tubulin polymerization begins, $y_0$ is the average y value up to time $t_0$, plateau is the y value at infinite times, and K is the rate constant.

*Cell cycle analysis by imaging flow cytometry and $EC_{50}$ determination*

HeLa cells were grown in 6-well plates (~250,000 cells per well) for 24 h in the presence of the fluorescent probe in variable concentrations. The probes were dissolved in DMSO at 500 - 2000-fold stock concentration and added to the media of cultured cells at 500 – 2000-fold dilution accordingly. In parallel, the appropriate DMSO control samples were prepared by adding corresponding amount of DMSO volume to the separate well. Cells were processed according to the NucleoCounter® NC-3000™ two-step cell cycle analysis protocol for cells attached to T-flasks, cell culture plates or micro-carriers. In particular, the 250 µl lysis solution (Solution 10, Chemometec Cat. No. 910-3010) supplemented with 10 µg/ml DAPI (Solution 12, Chemometec Cat. No. 910-3012) was used per well, incubated at 37 °C for 5 min. Then 250 µl of stabilization solution (Solution 11, Chemometec Cat. No. 910-3011) was added. Cells were counted on a NucleoCounter® NC-3000™ in NC-Slide A2™ slides (Chemometec, Cat. No. 942-0001) loaded with ~30 µl of each of the cell suspensions into the chambers of the slide. Each time, ~10,000 cells in total were measured, and the obtained cell cycle histograms were analysed with ChemoMetec NucleoView NC-3000 software, version 2.1.25.8. All experiments were repeated three times and the results are presented as means with standard deviations. The $EC_{50}$ values were determined by plotting the percentage of cells in subG1 phase and fitting the curve in GraphPad Prism 6 to the following function:

$$Y = Y_{min} + (Y_{max} - Y_{min}) / \left(1 + \left(\frac{EC_{50}}{X}\right)^{Hill}\right) \quad (6)$$

where $Y_{min}$ – cells population percentage in subG1 phase then no probe was added, $Y_{max}$ – highest reachable percentage of cells in subG1 phase and shared value for all data sets equal to 69%. $X$ - cells population percentage in subG1 phase then added probe is at X concentration, *Hill* - Hill slope coefficient determining the steepness of a dose-response curve, $EC_{50}$ - the concentration of probe that provoking halfway of subG1 cells in a population between the baseline ($Y_{min}$) and maximum response ($Y_{max}$).

*Processing and visualization of acquired images*

All acquired or reconstructed images were processed and visualized using Fiji[4]. Line profiles were measured using the “straight line” tool with the line width set to 3 pixels.

For the signal measurements, image files were converted to TIF file using Fiji and analyzed with CellProfiler 3.1.8 (ref.[5]), where the pipeline identified the nuclear region and measured the mean signal in this region. Background signal was measured in the region which is 3 pixels (450 nm) away from the nulear border and 7 pixels (1050 nm) wide. The background subtracted signal was processed with GraphPad Prism 6.

Actin/tubulin cytosolic signal was estimated using CellProfiler v.3.1.8. Briefly, probe channel was smoothed using median filter, nuclei were identified in DAPI channel and were used as seeds to find cell outlines in a smoothed actin channel. Background was defined as a lower quartile of pixel intensity in the area not covered by cells in the original probe channel. The background was subtracted from the original probe channel, and actin/tubulin staining was measured as mean pixel intensity per object in background-corrected probe channel. Statistical analysis performed GraphPad Prism 6

#### General experimental information and synthesis

NMR spectra were recorded at 25 °C with an Agilent 400-MR spectrometer at 400.06 MHz (1H) and 100.60 MHz ($^{13}$C), Bruker Avance III HD 500 spectrometer (av500) at 500.25 MHz ($^{1}$H) and 125.80 MHz ($^{13}$C), Varian Mercury Plus 300 spectrometer at 300.14 MHz ($^{1}$H) and are reported in ppm. All $^{1}$H and $^{13}$C spectra are referenced to tetramethylsilane (δ = 0 ppm) using the residual signals of the solvents according to the values reported in literature[6]. Multiplicities of signals are described as follows: s = singlet, d = doublet, t = triplet, q = quartet, p = pentet, m = multiplet or overlap of non-equivalent resonances; br = broad signal. Coupling constants (J) are given in Hz.

ESI-MS were recorded on a Varian 500-MS spectrometer (Agilent). ESI-HRMS were recorded on a MICROTOF spectrometer (Bruker) equipped with ESI ion source (Apollo) and direct injector with LC autosampler Agilent RR 1200. Liquid chromatography:

Analytical LC–MS analysis was performed on an Agilent 1260 Infinity II LC/MS system equipped with an autosampler, diode array detector WR, fluorescence detector Spectra and Infinity Lab LC/MSD 6100 series quadruple with API electrospray. Analysis was done by using an Agilent Zorbax SB-C18 RRHT, 2.1 x 50 mm, 1.8 µm threaded column and SUPELCO Titan C18, 2.1 x 75 mm, 1.9 µm column with A: 25 mM $HCOONH_4$ (pH = 3.6) aqueous buffer and B: MeOH

Preparative HPLC was performed on an Interchim puriFlash 4250 2X preparative HPLC/Flash hybrid system (Article No. 1I5140, Interchim) with a 2 mL / 5 mL injection loop, a 200-600 nm UV-Vis detector and an integrated ELSD detector (Article No. 1A3640, Interchim). Preparative column: Eurospher II 100-5 C18 5 µm, 250×20.0 mm (Article No.: 25PE181E2J, Knauer), typical flow rate: 25 mL/min, unless specified otherwise. Analytical TLC was performed on Merck Millipore ready-to-use plates with silica gel 60 (F254) (Cat. No. 1.05554.0001). Flash chromatography was performed on Biotage Isolera flash purification system using the indicated type of cartridge and solvent gradient.

Source of important chemical reagents used in the study: Larotaxel was synthesised according to the previously described procedure[7]. Cabazitaxel was bought from Carbosynth. **6-TMR-COOH** and **5-TMR-COOH** were bought from abcr GmbH. **5-580CP-COOH**, **5-610CP-COOH** and **5-SiR-COOH** were synthesised according to previously described procedures[8]. **6-580CP-COOH**, **6-610CP-COOH** and **6-SiR-COOH** synthesised according to[9]. Des-bromo-des-methyl-Lys-jasplakinolide was obtained according to literature procedure [10].

**Di-*tert*-butyl 3-bromophthalate (1):**

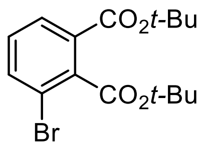

3-Bromophtalic acid (1.0 g, 4.08 mmol) was suspended in DCM (15 mL) in a sealable pressure glass tube. The mixture was cooled in a NaCl/ice bath and ~10 mL of isobutylene gas was condensed into the mixture. Catalytic amount of concentrated sulphuric acid (0.1 mL, 1.87 mmol) was added to the stirred and cooled reaction mixture and the pressure tube was tightly sealed. Reaction mixture was stirred at room temperature for 48 h, during this time the suspension became a clear solution. Then reaction mixture was cooled in ice bath and the tube was carefully opened with vigorous release of pressure. The resulting solution was poured to saturated $NaHCO_3$ solution (50 mL) and extracted with DCM (2 x 30mL). The organic extracts were combined and washed with water and brine, dried over $Na_2SO_4$. The product was isolated by flash column chromatography (Teledyne Isco RediSep Rf 40g, isocratic hold 5% of EtOAc in Hexane), fractions containing the product were evaporated to give 1.02g (70%) of white solid.

$^{1}$H NMR (400 MHz, $CDCl_3$) δ 7.79 (dd, *J* = 7.8, 1.1 Hz, 1H), 7.67 (dd, *J* = 8.0, 1.1 Hz, 1H), 7.23 (t, *J* = 7.9 Hz, 1H), 1.61 (s, 9H), 1.56 (s, 9H).

$^{13}$C NMR (101 MHz, $CDCl_3$) δ 165.9, 163.6, 137.9, 136.1, 131.6, 129.5, 128.5, 120.3, 83.0, 82.1, 28.1, 28.0.

ESI-MS, positive mode: m/z = 357.1, 359.1 $[M+H]^+$.

HRMS (ESI) calcd for $C_{16}H_{22}N_2BrO_4$ $[M+H]^+$ 357.0696, found 357.0698.

**3,6-bis((tert-butyldimethylsilyl)oxy)-9H-xanthen-9-one (2):**

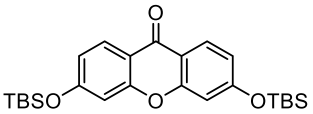

Was synthesised according to previously published procedure [11].

**3,6-bis((tert-butyldimethylsilyl)oxy)-10,10-dimethylanthracen-9(10H)-one (3)**:

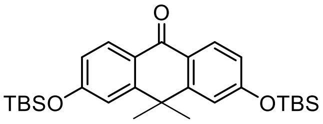

Was synthesised according to previously published procedure [12].

**3,7-bis((tert-butyldimethylsilyl)oxy)-5,5-dimethyldibenzo[b,e]silin-10(5H)-one (4):**

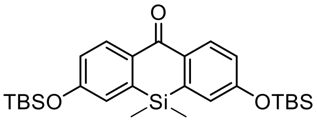

Was synthesised according to previously published procedure.[13]

**General procedure for the synthesis of compounds <u>5-7</u>:**

In a 25 mL round-bottom flask, a degassed solution of **1** (740 mg, 2.07 mmol, 2 eq.) in anhydrous THF (5 mL) and pentane (5 mL) was cooled to ~-116 °C (diethyl ether – liquid $N_2$ cooling bath). n-Butyllithium (1.3 mL of 1.6 M solution in hexanes, 2.07 mmol, 2 eq.) was carefully introduced through a needle. Clear solution quickly turned yellow and then deep orange; it was stirred at ~-116 °C for 10 min, and the solution of corresponding ketone (**2**[11], **3**[12] or **4**[13], 1.04 mmol 1 eq.) in THF (3 mL) was slowly injected into the reaction mixture. Stirred at ~-116°C for 10-15 minutes and then flask was taken out of the cooling bath and left to slowly warm up to rt and stirred for further 30 min. The reaction mixture was quenched with water (10 mL), adjusted to pH ~ 5 with acetic acid, extracted with ethyl acetate (3×30 mL), the combined organic layers were washed with brine and dried over $Na_2SO_4$. The products were isolated by flash column chromatography (Büchi Reveleris HP silica 40 g; gradient 0% to 20% ethyl acetate – hexane.) In some cases additional purification was needed (Teledyne Isco RediSep Rf 40g, gradient 20% to 100% of DCM-Hexane). However, it is advised to proceed to the next step with a semi-pure substance as in further steps impurities are seperated easier.

**Tert-butyl 3',6'-bis((tert-butyldimethylsilyl)oxy)-3-oxo-3H-spiro[isobenzofuran-1,9'-xanthene]-4-carboxylate (5)**:

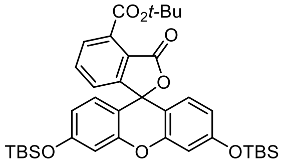

Product yield 433 mg (63%) of off-white solid.

$^{1}$H NMR (400 MHz, $d_6$-acetone) δ 7.86 (t, *J* = 7.6 Hz, 1H), 7.82 (dd, *J* = 7.6, 1.4 Hz, 1H), 7.40 (dd, *J* = 7.6, 1.4 Hz, 1H), 6.81 (d, *J* = 2.4 Hz, 2H), 6.76 (d, *J* = 8.6 Hz, 2H), 6.70 (dd, *J* = 8.6, 2.4 Hz, 2H), 1.66 (s, 9H), 1.00 (s, 18H), 0.27 (s, 12H).

$^{13}$C NMR (101 MHz, $d_6$-acetone) δ 165.9, 165.0, 157.6, 153.9, 152.1, 135.2, 133.1, 129.3, 129.1, 126.0, 123.2, 116.9, 112.4, 107.4, 82.4, 81.3, 27.2, 25.0, 17.9, -5.3.

ESI-MS, positive mode: m/z = 661.3 $[M+H]^+$.

HRMS (ESI) calcd for $C_{37}H_{49}O_7Si_2$ $[M+H]^+$ 661.3011, found 661.3004.

**Tert-butyl 3,6-bis((tert-butyldimethylsilyl)oxy)-10,10-dimethyl-3'-oxo-3'H,10H-spiro[anthracene-9,1'-isobenzofuran]-4'-carboxylate (6)**:

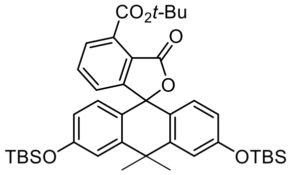

Product yield 471 mg (66%) of off-white solid.

$^{1}$H NMR (400 MHz, $CDCl_3$) δ 7.71 (dd, J = 7.5, 1.0 Hz, 1H), 7.57 (t, J = 7.6 Hz, 1H), 7.06 – 7.03 (m, 3H), 6.66 – 6.63 (m, 2H), 6.60 (dd, J = 8.6, 2.4 Hz, 2H), 1.78 (s, 3H), 1.70 (s, 3H), 1.69 (s, 9H), 0.97 (s, 18H), 0.20 (s, 12H).

$^{13}$C NMR (101 MHz, $CDCl_3$) δ 167.4, 165.5, 156.5, 156.2, 146.7, 134.2, 133.1, 129.3, 129.0, 125.8, 124.2, 123.2, 119.0, 117.5, 85.0, 83.3, 38.0, 34.9, 33.0, 28.0, 25.7, 18.2, -4.4.

ESI-MS, positive mode: m/z = 687.4 $[M+H]^+$.

HRMS (ESI) calcd for $C_{40}H_{55}O_6Si_2$ $[M+H]^+$ 687.3532, found 687.3523.

**Tert-butyl 3,7-bis((tert-butyldimethylsilyl)oxy)-5,5-dimethyl-3'-oxo-3'H,5H-spiro[dibenzo[b,e]siline-10,1'-isobenzofuran]-4'-carboxylate (7)**:

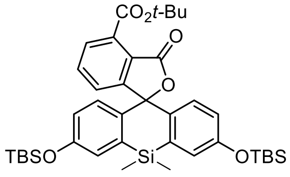

Product yield 446 mg (61%) of off-white solid.

$^{1}$H NMR (400 MHz, $CDCl_3$) δ 7.69 (dd, *J* = 7.6, 1.0 Hz, 1H), 7.62 (t, *J* = 7.6 Hz, 1H), 7.31 (dd, *J* = 7.6, 1.0 Hz, 1H), 7.09 (d, *J* = 2.7 Hz, 2H), 6.87 (dd, *J* = 8.7, 0.4 Hz, 2H), 6.67 (dd, *J* = 8.7, 2.7 Hz, 2H), 1.65 (s, 9H), 0.96 (s, 18H), 0.60 (s, 3H), 0.56 (s, 3H), 0.17 (s, 12H).

$^{13}C$ NMR (101 MHz, $CDCl_3$) δ 167.5, 165.6, 155.2, 155.2, 137.2, 136.9, 134.0, 133.5, 128.9, 128.6, 126.3, 125.0, 123.1, 121.2, 89.1, 83.4, 28.1, 25.7, 18.3, -1.3, -2.9, -4.3, -4.

ESI-MS, positive mode: m/z = 703.3 $[M+H]^+$.

HRMS (ESI) calcd for $C_{39}H_{55}O_6Si_3$ $[M+H]^+$ 703.3301, found 703.3304.

**General procedure for the synthesis of compounds <u>8-10</u>:**

To a cooled (ice-water bath) solution of corresponding compound **5-7** (1 eq.) in THF (15 mL) tetrabutylammonium fluoride trihydrate (4 eq.) solution in THF (5 mL) was added. The resulting intensively coloured solution was stirred at 0 °C for 1 h. Sat. aq. $NH_4Cl$ (20 mL) was added followed by minimal amount of water necessary to dissolve the solids, the mixture was extracted with ethyl acetate (3×30 mL), the combined organic layers were washed with brine and dried over $Na_2SO_4$. The products were isolated by flash column chromatography (Teledyne Isco RediSep Rf 24 g; gradient 2% to 30% ethyl acetate – $CH_2Cl_2$) and evaporated to obtain viscous oils which solidifies overtime.

**Tert-butyl 3',6'-dihydroxy-3-oxo-3H-spiro[isobenzofuran-1,9'-xanthene]-4-carboxylate (8)**: Reaction was carried starting from compound **5** (420 mg, 0.635 mmol). Product yield 214 mg (78%) of orange solid material.

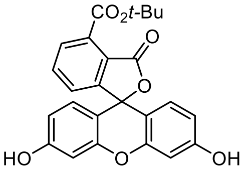

$^1H$ NMR (400 MHz, $d_6$-dmso) δ 10.15 (s, 2H), 7.87 – 7.76 (m, 2H), 7.39 (dd, $J$ = 7.3, 1.4 Hz, 1H), 6.69 (d, $J$ = 2.0 Hz, 2H), 6.63 – 6.47 (m, 4H), 1.60 (s, 9H).

$^{13}C$ NMR (101 MHz, $d_6$-dmso) δ 166.1, 164.9, 159.5, 153.4, 151.9, 135.6, 132.2, 129.2, 128.9, 126.3, 122.6, 112.7, 109.3, 102.3, 82.6, 82.2, 27.7.

ESI-MS, positive mode: m/z = 433.1 $[M+H]^+$.

HRMS (ESI) calcd for $C_{25}H_{21}O_7$ $[M+H]^+$ 433.1282, found 433.1279.

**Tert-butyl 3,6-dihydroxy-10,10-dimethyl-3'-oxo-3'H,10H-spiro[anthracene-9,1'-isobenzofuran]-4'-carboxylate (9)**:

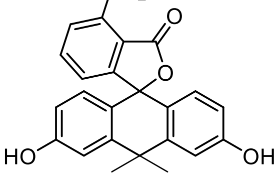

Reaction was carried starting from compound **6** (445 mg, 0.647 mmol). Product yield 264 mg (89%) of orange solid material.

$^1H$ NMR (300 MHz, $CDCl_3$) δ 7.73 (dd, $J$ = 7.6, 1.0 Hz, 1H), 7.60 (t, $J$ = 7.6 Hz, 1H), 7.14 (s, 2H), 7.05 (dd, $J$ = 7.6, 1.0 Hz, 1H), 6.98 (d, $J$ = 2.4 Hz, 2H), 6.51 (dd, $J$ = 8.6, 2.4 Hz, 2H), 6.44 (d, $J$ = 8.6 Hz, 2H), 1.68 (s, 9H).

[13]C NMR (126 MHz, $CDCl_3$) δ 169.0, 165.9, 156.7, 156.2, 147.3, 134.7, 132.7, 129.2, 129.1, 126.1, 123.2, 122.4, 114.8, 112.9, 87.0, 84.1, 38.1, 34.8, 32.3, 28.1.

ESI-MS, positive mode: m/z = 459.2 $[M+H]^+$.

HRMS (ESI) calcd for $C_{28}H_{27}O_6$ $[M+H]^+$ 459.1802, found 459.1805.

**Tert-butyl 3,7-dihydroxy-5,5-dimethyl-3'-oxo-3'H,5H-spiro[dibenzo[b,e]siline-10,1'-isobenzofuran]-4'-carboxylate (10)**:

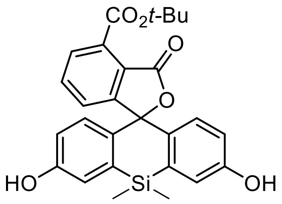

Reaction was carried starting from compound **7** (420 mg, 0.575 mmol). Product yield 254 mg (93%) of orange solid material.

[1]H NMR (400 MHz, $d_6$-dmso) δ 9.74 (s, 2H), 7.83 (t, *J* = 7.6 Hz, 1H), 7.72 (dd, *J* = 7.6, 0.9 Hz, 1H), 7.46 (dd, *J* = 7.6, 0.9 Hz, 1H), 7.16 – 7.08 (m, 2H), 6.74 – 6.60 (m, 4H), 1.55 (s, 9H), 0.56 (s, 3H), 0.48 (s, 3H).

[13]C NMR (101 MHz, $d_6$-dmso) δ 167.4, 165.4, 157.3, 154.6, 137.5, 135.2, 134.5, 133.5, 129.0, 128.6, 127.1, 122.3, 120.7, 117.3, 90.0, 83.0, 28.0, 0.4, -1.4.

ESI-MS, positive mode: m/z = 475.2 $[M+H]^+$.

HRMS (ESI) calcd for $C_{27}H_{27}O_6Si$ $[M+H]^+$ 475.1571, found 475.1573.

**General procedure for the synthesis of compounds <u>11-13</u>:**

Trifluoromethanesulfonic anhydride 1M solution in DCM (4 eq.) was slowly added dropwise to a solution of corresponding compound **8-10** (1 eq.) and pyridine (8 eq.) in dry DCM (10 mL), cooled in ice-water bath. The flask was then removed from the cooling bath, and the mixture was stirred at rt for 1 h. Afterwards, the mixture was diluted with water (30 mL), extracted with $CH_2Cl_2$ (3×20 mL), the combined extracts were washed with water, brine and dried over $Na_2SO_4$. The products were isolated by flash column chromatography (Teledyne Isco RediSep Rf 24 g; gradient 5% to 40% ethyl acetate – hexane).

**Tert-butyl 3-oxo-3',6'-bis(((trifluoromethyl)sulfonyl)oxy)-3H-spiro[isobenzofuran-1,9'-xanthene]-4-carboxylate (11)**:

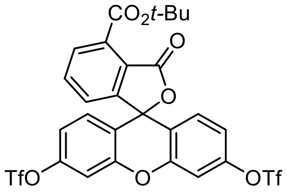

Reaction was carried starting from compound **8** (200 mg, 0.463 mmol). Product yield 255 mg (79%) of white solid material.

[1]H NMR (400 MHz, $CDCl_3$) δ 7.87 (d, *J* = 7.4 Hz, 1H), 7.73 (t, *J* = 7.4 Hz, 1H), 7.30 (d, *J* = 2.2 Hz, 2H), 7.22 (d, *J* = 7.4 Hz, 1H), 7.08 – 6.99 (m, 4H), 1.70 (s, 9H).

$^{13}C$ NMR (101 MHz, $CDCl_3$) δ 165.4, 164.6, 153.3, 151.2, 150.2, 135.4, 133.7, 130.7, 130.0, 125.7, 122.4, 119.1, 118.6 (q, $^1J_{C-F}$ = 319 Hz, -$CF_3$), 117.7, 110.7, 83.9, 78.8, 28.0.

ESI-MS, positive mode: m/z = 697.0 $[M+H]^+$.

HRMS (ESI) calcd for $C_{27}H_{19}F_6O_{11}S_2$ $[M+H]^+$ 697.0267, found 697.0249.

**Tert-butyl 10,10-dimethyl-3'-oxo-3,6-bis(((trifluoromethyl)sulfonyl)oxy)-3'H,10H-spiro[anthracene-9,1'-isobenzofuran]-4'-carboxylate (12)**:

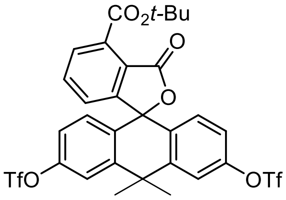

Reaction was carried starting from compound **9** (245 mg, 0.535 mmol). Product yield 271 mg (70%) of white solid material.

$^1H$ NMR (400 MHz, $CDCl_3$) δ 7.83 (dd, *J* = 7.6, 0.9 Hz, 1H), 7.68 (t, *J* = 7.6 Hz, 1H), 7.54 (d, *J* = 2.5 Hz, 2H), 7.11 (dd, *J* = 8.8, 2.5 Hz, 2H), 7.07 (dd, *J* = 7.6, 0.9 Hz, 1H), 6.94 (d, *J* = 8.8 Hz, 2H), 1.89 (s, 3H), 1.79 (s, 3H), 1.70 (s, 9H).

$^{13}C$ NMR (101 MHz, $CDCl_3$) δ 166.4, 164.8, 154.8, 150.1, 146.9, 135.1, 133.8, 131.3, 130.3, 130.2, 125.6, 122.6, 120.3, 119.6, 118.9 (q, $^1J_{C-F}$ = 319 Hz,-$CF_3$), 83.8, 82.6, 38.8, 34.7, 33.0, 28.0.

ESI-MS, positive mode: m/z = 745.1 $[M+Na]^+$.

HRMS (ESI) calcd for $C_{30}H_{25}O_{10}S_2F_6$ $[M+H]^+$ 723.0788, found 723.0800.

**Tert-butyl 5,5-dimethyl-3'-oxo-3,7-bis(((trifluoromethyl)sulfonyl)oxy)-3'H,5H-spiro[dibenzo[b,e]siline-10,1'-isobenzofuran]-4'-carboxylate (13)**:

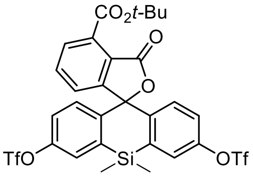

Reaction was carried starting from compound **10** (230 mg, 0.484 mmol). Product yield 258 mg (72%) of white solid material.

$^1H$ NMR (400 MHz, $CDCl_3$) δ 7.81 (dd, *J* = 7.6, 1.1 Hz, 1H), 7.75 (t, *J* = 7.6 Hz, 1H), 7.56 (dd, *J* = 2.3, 0.8 Hz, 2H), 7.39 (dd, *J* = 7.6, 1.1 Hz, 1H), 7.24 – 7.17 (m, 4H), 1.67 (s, 9H), 0.74 (s, 3H), 0.70 (s, 3H).

$^{13}C$ NMR (101 MHz, $CDCl_3$) δ 166.2, 164.8, 152.8, 149.3, 144.0, 138.7, 134.8, 134.3, 130.0, 129.2, 126.3, 126.1, 122.8, 122.6, 118.7 (q, $^1J_{C-F}$ = 319 Hz,-$CF_3$), 87.3, 83.8, 28.0,-0.2, -1.7.

ESI-MS, positive mode: m/z = 761.0 $[M+Na]^+$.

HRMS (ESI) calcd for $C_{29}H_{25}O_{10}S_2SiF_6$ $[M+H]^+$ 739.0557, found 739.0562.

**General procedure for the synthesis of compounds 14-17:**

A mixture of $Pd_2(dba)_3$ (0.1 eq.), Xantphos (0.3 eq.), $Cs_2CO_3$ (3 eq.) and corresponding triflate (1 eq., **11**, **12** or **13**) in dry 1,4-dioxane (2.5 mL) was degassed on a Schlenk line. Then solution of *tert*-butyl N-methylcarbamate in 1,4-dioxane (2.5 eq. for compound **15**) or 2M solution of dimethylamine in THF (2.5 eq. for compounds **14**, **16-17**) were introduced. Reaction mixture was stirred in a septa sealed tube at 100°C under argon for 18h, except compound **17**, which was stirred at 80°C for 5h. Upon cooling, the resulting brown mixture was diluted with water (30 mL), extracted with ethyl acetate (3×30 mL), the combined organic layers were washed with brine and dried over $Na_2SO_4$. The filtrate was evaporated and the products were isolated by flash chromatography.

**Tert-butyl 3',6'-bis(dimethylamino)-3-oxo-3H-spiro[isobenzofuran-1,9'-xanthene]-5-carboxylate (14)**:

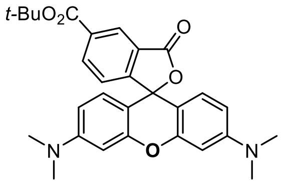

Reaction was carried starting from compound **11** (180 mg, 0.258 mmol). Product was isolated by flash column chromatography Büchi Reveleris HP silica 24 g; gradient 2% to 30% MeOH – DCM. The fractions containing the product were evaporated, the residue was redissolved in acetonitrile – water (1:1), microfiltered through a 0.45 μm PTFE membrane filter and lyophilized to obtain 67 mg (53%) of pink solid.

$^1$H NMR (400 MHz, $CD_3OD$) δ 8.00 (dd, J = 7.8, 1.3 Hz, 1H), 7.57 (t, J = 7.8 Hz, 1H), 7.39 (dd, J = 7.8, 1.3 Hz, 1H), 7.31 (d, J = 9.5 Hz, 2H), 7.00 (dd, J = 9.5, 2.5 Hz, 2H), 6.85 (d, J = 2.5 Hz, 2H), 3.26 (s, 12H), 1.61 (s, 9H).

$^{13}$C NMR (101 MHz, $CD_3OD$) δ 173.7, 167.8, 159.1, 158.9, 158.8, 144.0, 133.6, 133.3, 131.9, 131.8, 131.6, 128.1, 115.3, 114.9, 97.3, 83.3, 40.9, 28.3.

ESI-MS, positive mode: m/z = 487.2 $[M+H]^+$.

HRMS (ESI) calcd for $C_{29}H_{31}N_2O_5$ $[M+H]^+$ 487.2227, found 487.2232

**Tert-butyl 10,10-dimethyl-3,6-bis(methylamino)-3'-oxo-3'H,10H-spiro[anthracene-9,1'-isobenzofuran]-5'-carboxylate (15)**:

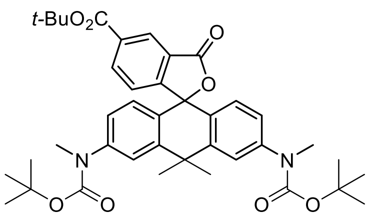

Reaction was carried starting from compound **12** (240 mg, 0.332 mmol). Product was isolated by flash column chromatography Büchi Reveleris HP silica 24 g; gradient 5% to 50% EtOAc – Hexane. The fractions containing the product were evaporated, the residue was redissolved in acetonitrile – water (1:1), microfiltered through a 0.45 μm PTFE membrane filter and lyophilized to obtain 173 mg (76%) of violet powder.

$^1$H NMR (400 MHz, $CDCl_3$) δ 7.75 (dd, J = 7.6, 0.9 Hz, 1H), 7.59 (t, J = 7.6 Hz, 1H), 7.52 (d, J = 2.2 Hz, 2H), 7.06 (dd, J = 7.6, 0.9 Hz, 1H), 7.01 (dd, J = 8.6, 2.2 Hz, 2H), 6.76 (d, J = 8.6 Hz, 2H), 3.27 (s, 6H), 1.85 (s, 3H), 1.76 (s, 3H), 1.70 (s, 9H), 1.46 (s, 18H).

$^{13}$C NMR (101 MHz, $CDCl_3$) δ 167.4, 165.5, 156.2, 154.6, 145.3, 144.7, 134.5, 133.4, 129.5, 128.4, 127.9, 126.0, 123.6, 123.6, 123.2, 84.4, 83.6, 80.8, 38.3, 37.3, 35.0, 33.3, 28.5, 28.2.

ESI-MS, positive mode: m/z = 685.4 $[M+H]^+$.

HRMS (ESI) calcd for $C_{40}H_{49}N_2O_8$ $[M+H]^+$ 685.3483, found 685.3474.

**Tert-butyl 3,6-bis(dimethylamino)-10,10-dimethyl-3'-oxo-3'H,10H-spiro[anthracene-9,1'-isobenzofuran]-5'-carboxylate (16)**:

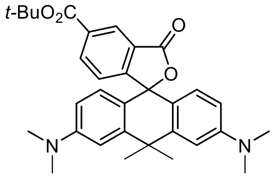

Reaction was carried starting from compound **12** (250 mg, 0.36 mmol). Product was isolated by flash column chromatography Büchi Reveleris HP silica 24 g; gradient 20% to 80% EtOAc – Hexane. The fractions containing the product were evaporated, the residue was redissolved in 1,4-dioxane – water (1:1), microfiltered through a 0.45 μm PTFE membrane filter and lyophilized to obtain 131 mg (71%) of violet powder.

$^{1}$H NMR (400 MHz, $CDCl_3$) δ 7.69 (dd, J = 7.6, 1.0 Hz, 1H), 7.55 (t, J = 7.6 Hz, 1H), 7.09 (dd, J = 7.6, 1.0 Hz, 1H), 6.88 (d, J = 2.6 Hz, 2H), 6.65 (d, J = 8.8 Hz, 2H), 6.53 (dd, J = 8.8, 2.6 Hz, 2H), 2.99 (s, 12H), 1.87 (s, 3H), 1.78 (s, 3H), 1.70 (s, 9H).

$^{13}$C NMR (101 MHz, $CDCl_3$) δ 167.9, 166.0, 157.0, 150.8, 146.7, 134.0, 133.0, 129.1, 128.8, 126.1, 124.0, 119.5, 111.8, 109.2, 86.5, 83.3, 40.6, 38.6, 35.6, 33.1, 28.2.

ESI-MS, positive mode: m/z = 513.6 $[M+H]^+$.

HRMS (ESI) calcd for $C_{32}H_{37}N_2O_4$ $[M+H]^+$ 513.2748, found 513.2748.

**Tert-butyl 3,7-bis(dimethylamino)-5,5-dimethyl-3'-oxo-3'H,5H-spiro[dibenzo[b,e]siline-10,1'-isobenzofuran]-5'-carboxylate (17)**:

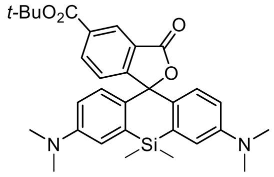

Reaction was carried starting from compound **13** (210 mg, 0.284 mmol). Product was isolated by flash column chromatography Büchi Reveleris HP silica 24 g; gradient 5% to 50% EtOAc – Hexane. The fractions containing the product were evaporated, the residue was redissolved in 1,4-dioxane – water (1:1), microfiltered through a 0.45 μm PTFE membrane filter and lyophilized to obtain 73 mg (48%) of light-blue powder.

$^{1}$H NMR (400 MHz, $CDCl_3$) δ 7.69 (dd, J = 7.6, 1.0 Hz, 1H), 7.61 (t, J = 7.6 Hz, 1H), 7.32 (dd, J = 7.6, 1.0 Hz, 1H), 6.96 (d, J = 2.9 Hz, 2H), 6.82 (d, J = 8.9 Hz, 2H), 6.57 (dd, J = 8.9, 2.9 Hz, 2H), 2.96 (s, 12H), 1.67 (s, 9H), 0.64 (s, 3H), 0.60 (s, 3H).

$^{13}$C NMR (101 MHz, $CDCl_3$) δ 167.8, 166.0, 155.8, 149.4, 136.7, 133.8, 133.3, 131.8, 128.7, 128.3, 126.5, 123.8, 116.6, 113.6, 90.3, 83.3, 40.4, 28.1, 0.4, -1.1.

ESI-MS, positive mode: m/z = 529.3 $[M+H]^+$.

HRMS (ESI) calcd for $C_{31}H_{37}N_2O_4Si$ $[M+H]^+$ 529.2517, found 529.2521.

**General procedure for the synthesis of compounds 4-TMR-COOH (18), 4-580CP-COOH (19), 4-610CP-COOH (20) and 4-SiR-COOH (21):**

Trifluoroacetic acid (2 mL) was added dropwise to a solution of corresponding compound **14-17** in DCM (8 mL). The resulting coloured solution was stirred at room temperature overnight. The reaction mixture was then evaporated to dryness, the residue was re-evaporated three times with toluene to remove excess of trifluoroacetic acid. The residue was lyophilized from 1,4-dioxane – water (1:1). Products were obtained as trifluoroacetic acid salts.

**4-TMR-COOH (18):**

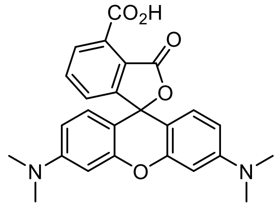

Reaction was performed starting from compound **14** (67 mg, 0.138 mmol). Obtained 74 mg (98%) of pink powder as trifluoroacetic acid salt.

$^{1}H$ NMR (400 MHz, $CD_3OD$) δ 8.25 (dd, J = 7.8, 1.2 Hz, 1H), 7.82 (t, J = 7.8 Hz, 1H), 7.60 (dd, J = 7.8, 1.2 Hz, 1H), 7.21 (d, J = 9.5 Hz, 2H), 7.08 (dd, J = 9.5, 2.4 Hz, 2H), 6.87 (d, J = 2.4 Hz, 2H), 3.29 (s, 12H).

$^{13}C$ NMR (101 MHz, $CD_3OD$) δ 170.5, 168.6, 159.0, 159.0, 157.0, 136.8, 134.1, 132.8, 132.5, 132.2, 132.0, 131.0, 115.5, 115.2, 97.4, 41.0.

ESI-MS, positive mode: m/z = 431.2 $[M+H]^+$.

HRMS (ESI) calcd for $C_{25}H_{23}N_2O_5$ $[M+H]^+$ 431.1601, found 431.1606.

**4-580CP-COOH (19):**

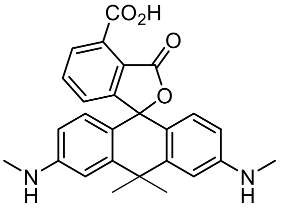

Reaction was performed starting from compound **15** (220 mg, 0.321 mmol). Obtained 170 mg (98%) of violet powder as trifluoroacetic acid salt.

$^{1}H$ NMR (400 MHz, $CD_3OD$) δ 8.17 (dd, J = 7.7, 1.1 Hz, 1H), 7.76 (t, J = 7.7 Hz, 1H), 7.52 (dd, J = 7.7, 1.1 Hz, 1H), 7.11 (d, J = 2.2 Hz, 2H), 7.03 (d, J = 9.2 Hz, 2H), 6.64 (dd, J = 9.2, 2.2 Hz, 2H), 3.06 (s, 6H), 1.81 (s, 3H), 1.68 (s, 3H).

$^{13}C$ NMR (101 MHz, $CD_3OD$) δ 170.6, 168.9, 162.5, 159.4, 139.6, 139.5, 136.6, 136.0, 134.2, 131.72, 131.68, 130.57, 122.1, 112.7 (visible in HSQC spectra), 42.7, 35.8, 31.9, 30.1.

ESI-MS, positive mode: m/z = 429.2 $[M+H]^+$.

HRMS (ESI) calcd for $C_{26}H_{25}N_2O_4$ $[M+H]^+$ 429.1809, found 429.1807.

**4-610CP-COOH (20):**

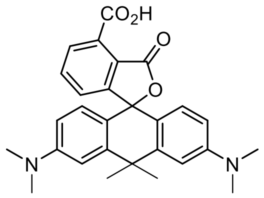

Reaction was performed starting from compound **16** (230 mg, 0.448 mmol). Obtained 251 mg (98%) of dark-violet powder as trifluoroacetic acid salt.

$^{1}H$ NMR (400 MHz, $CD_3OD$) δ 8.17 (dd, J = 7.8, 1.2 Hz, 1H), 7.77 (t, J = 7.8 Hz, 1H), 7.53 (dd, J = 7.8, 1.2 Hz, 1H), 7.23 (d, J = 2.5 Hz, 2H), 7.09 (d, J = 9.4 Hz, 2H), 6.84 (dd, J = 9.4, 2.5 Hz, 2H), 3.34 (s, 12H), 1.86 (s, 3H), 1.72 (s, 3H).

$^{13}C$ NMR (101 MHz, $CD_3OD$) δ 170.6, 168.9, 158.3, 157.9, 138.9, 136.6, 135.9, 134.2, 131.8, 131.7, 130.6, 122.0, 114.0, 112.1, 43.2, 41.0, 36.2, 32.3.

ESI-MS, positive mode: m/z = 457.2 $[M+H]^+$.

HRMS (ESI) calcd for $C_{28}H_{29}N_2O_4$ $[M+H]^+$ 457.2122, found 457.2119.

**4-SiR-COOH (21):**

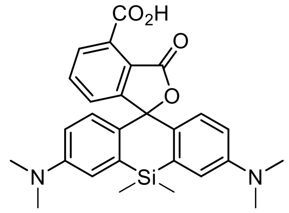

Reaction was performed starting from compound **17** (50 mg, 0.095 mmol). Obtained 54 mg (98%) of blue powder as trifluoroacetic acid salt.

$^1$H NMR (400 MHz,$d_5$-pyridine) δ 8.17 (dd, J = 7.7, 0.9 Hz, 1H), 7.77 (t, J = 7.7 Hz, 1H), 7.54 (dd, J = 7.7, 0.9 Hz, 1H), 7.18 (d, J = 2.9 Hz, 2H), 7.04 (d, J = 9.0 Hz, 2H), 6.54 (dd, J = 9.0, 2.9 Hz, 2H), 2.85 (s, 12H), 0.73 (s, 3H), 0.65 (s, 3H).

$^{13}$C NMR (101 MHz, $d_5$-pyridine) δ 170.1, 169.4, 150.4 (overlapped with pyridine, visible in HMBC spectra), 157.1, 137.4, 135.6, 134.9, 132.3, 129.7, 129.2, 127.0, 124.2 (overlapped with pyridine, visible in HMBC spectra), 117.4, 114.6, 92.3, 40.4, 0.8, -0.8.

ESI-MS, positive mode: m/z = 473.2 $[M+H]^+$.

HRMS (ESI) calcd for $C_{27}H_{29}N_2O_4Si$ $[M+H]^+$ 473.1891, found 473.1890.

**General procedure for the synthesis of compounds CTX-C8NH-Boc (SI-1) and LTX-C8-NHBoc (SI-2):**

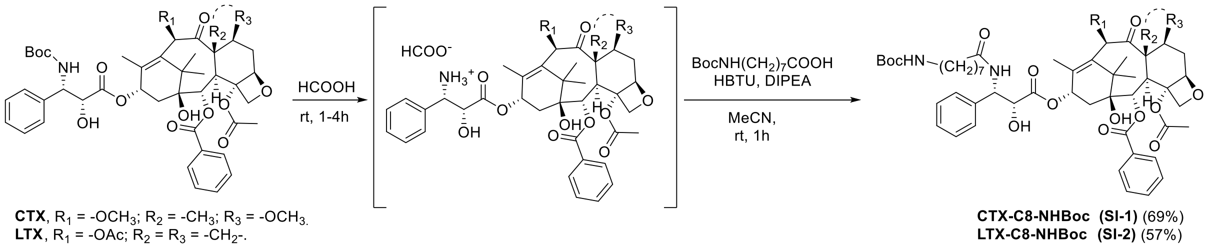

A solution of corresponding taxane derivative (0.24 mmol) in 95% formic acid (2 mL) was stirred at room temperature for 1 - 4h. Reaction progress was monitored by HPLC analysis. Once reaction was complete, formic acid was evaporated on rotary evaporator and residue was dissolved in water and lyophilised to obtain white powder.

Into a solution of Boc protected 8-aminooctanoic acid (1.4 eq., 0.336 mmol, 91.7 mg) in MeCN (2 mL) was added HBTU (1.2 eq., 0.288 mmol, 109 mg), followed by DIPEA (4.3 eq., 1 mmol, 100µL). Mixture was stirred at rt for 5 min and previously obtained corresponding taxol-amine (0.24 mmol) was added. Mixture was stirred for 1 hour then solvent was removed by rotary evaporator. The residue was re-dissolved in 70% MeCN-$H_2O$ mixture, microfiltered through a 0.45 µm PTFE membrane filter and purified by the preperative HPLC (preparative column: Knauer 100 C18, 5 µm, 250 × 30 mm; solvent A: acetonitrile, solvent B: $H_2O$ + 0.2% v/v HCOOH; temperature 25 °C, gradient A:B - 5 min 50:50 isocratic, 5-30 min 50:50 to 100:0 gradient). Fractions containing the product were collected, solvent was removed and obtained residue was lyophilised from 50:50 MeCN : $H_2O$ mixture to obtain products as white solids.

**CTX-C8-NHBoc (SI-1):**

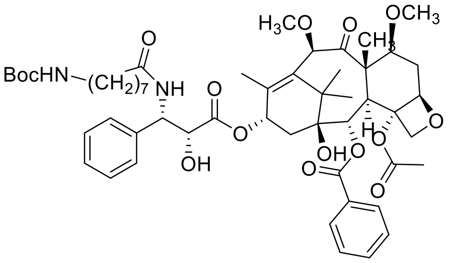

Obtained 69% (162 mg) of white fluffy solid.

$^{1}H$ NMR (400 MHz, $d_6$-dmso) δ 8.37 (d, J = 9.1 Hz, 1H), 8.00 (dt, J = 7.1, 1.4 Hz, 2H), 7.70 (tt, J = 7.4, 2.2 Hz, 1H), 7.61 (t, J = 7.4 Hz, 2H), 7.40 – 7.30 (m, 4H), 7.23 (tt, J = 7.4, 1.4 Hz, 1H), 6.74 (t, J = 5.7 Hz, 1H), 5.99 – 5.91 (m, 2H), 5.40 (d, J = 7.1 Hz, 1H), 5.30 (dd, J = 9.2, 5.8 Hz, 1H), 4.97 (dd, J = 9.7, 2.0 Hz, 1H), 4.72 (s, 1H), 4.66 (s, 1H), 4.44 (dd, J = 6.8, 5.8 Hz, 1H), 4.04 (s, 2H), 3.77 (dd, J = 10.6, 6.5 Hz, 1H), 3.65 (d, J = 7.1 Hz, 1H), 3.32 (s, 3H), 3.23 (s, 3H), 2.87 (q, J = 6.6 Hz, 2H), 2.72 – 2.63 (m, 1H), 2.26 (s, 3H), 2.17 (t, J = 7.2 Hz, 2H), 1.99 (dd, J = 15.4, 9.1 Hz, 1H), 1.92 – 1.86 (m, 1H), 1.85 (s, 3H), 1.53 (s, 3H), 1.52 – 1.43 (m, 3H), 1.37 (s, 9H), 1.35 – 1.30 (m, 2H), 1.24 – 1.16 (m, 6H), 1.04 (s, 3H), 0.99 (s, 3H).

13C NMR (101 MHz, $d_6$-dmso) δ 204.7, 172.6, 171.9, 169.9, 165.2, 155.5, 139.6, 138.4, 134.9, 133.3, 129.9, 129.6, 128.7, 128.1, 127.1, 83.2, 82.1, 80.3, 80.2, 77.3, 76.9, 75.3, 74.4, 73.6, 70.0, 56.6, 56.6, 56.0, 54.9, 46.4, 43.0, 39.8, 35.4, 34.9, 31.7, 29.5, 28.6, 28.5, 28.3, 26.7, 26.2, 25.4, 22.4, 21.2, 14.0, 10.2.

ESI-MS, positive mode: m/z = 977.5 $[M+H]^+$.

HRMS (ESI) calcd for $C_{53}H_{73}N_2O_{15}$ $[M+H]^+$ 977.5005, found 977.4995.

**LTX-C8-NHBoc: (SI-2)**

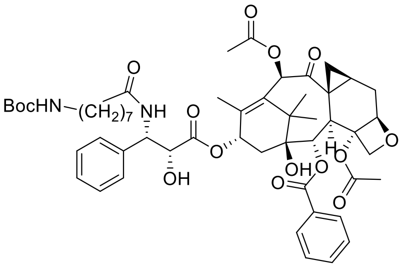

Obtained 57% (133 mg) of white fluffy solid.

$^{1}H$ NMR (400 MHz, $d_6$-dmso) δ 8.34 (d, J = 9.2 Hz, 1H), 8.08 – 8.00 (m, 2H), 7.69 (tt, J = 7.4, 1.4 Hz, 1H), 7.61 (t, J = 7.5 Hz, 2H), 7.41 – 7.29 (m, 4H), 7.19 (tt, J = 7.4, 1.7 Hz, 1H), 6.73 (t, J = 5.7 Hz, 1H), 6.12 (s, 1H), 6.02 – 5.91 (m, 1H), 5.87 (d, J = 6.8 Hz, 1H), 5.44 (d, J = 7.7 Hz, 1H), 5.34 (dd, J = 9.2, 5.2 Hz, 1H), 4.77 (s, 1H), 4.72 (d, J = 3.0 Hz, 1H), 4.49 (dd, J = 6.8, 5.2 Hz, 1H), 4.10 – 3.94 (m, 2H), 3.91 (d, J = 7.6 Hz, 1H), 2.86 (q, J = 6.5 Hz, 2H), 2.36 – 2.23 (m, 4H), 2.17 – 1.87 (m, 9H), 1.74 (d, J = 1.4 Hz, 3H), 1.54 (dd, J = 7.3, 4.8 Hz, 1H), 1.50 – 1.41 (m, 2H), 1.36 (s, 9H), 1.35 – 1.28 (m, 2H), 1.19 – 1.14 (m, 6H), 1.12 (s, 3H), 1.08 (s, 3H).

$^{13}C$ NMR (101 MHz, $d_6$-dmso) δ 202.0, 172.8, 172.4, 170.2, 169.5, 165.8, 156.0, 140.5, 140.3, 134.0, 133.7, 130.4, 130.1, 129.2, 128.6, 127.5, 127.5, 84.2, 80.0, 79.0, 77.8, 77.7, 75.7, 74.9, 74.0, 70.0, 55.1, 43.0, 40.3, 38.0, 36.1, 35.8, 34.9, 31.7, 29.9, 29.1, 28.9, 28.7, 26.7, 26.4, 26.0, 25.8, 22.4, 21.8, 21.0, 15.2, 14.4.

ESI-MS, positive mode: m/z = 973.6 $[M+H]^+$.

HRMS (ESI) calcd for $C_{53}H_{69}N_2O_{15}$ $[M+H]^+$ 973.4692, found 973.4687.

**General synthesis procedure for Rhodamine-C8-Taxane conjugates:**

A solution of corresponding taxane (**CTX-C8-NHBoc (SI-1)** or **LTX-C8-NHBoc (SI-2)** derivative (0.015 mmol) in 95% formic acid (1 mL) was stirred at room temperature for 1 h. Reaction progress was monitored by HPLC analysis. Once reaction was complete, formic acid was evaporated on rotary evaporator and residue was dissolved in water and lyophilised to obtain white powder, which was used further without any additional purifications.

The corresponding rhodamine dye (0.01 mmol, 1eq), DIPEA (52 µL, 0.3 mmol, 30 eq.) and HBTU (0.012 mmol, 4.5 mg, 1.2 eq.) were dissolved in 400 µL of dry MeCN and stirred at room temperature for 5 min. A solution of previously obtained deprotected taxane (**CTX-C8-NH$_2$ (SI-3)** or **LTX-C8-NH$_2$ (SI-4)** derivative (0.015 mmol, 1.5 eq) in MeCN and 10 µL of DIPEA were added to the reaction mixture and stirring continued for 1 hour. Reaction was monitored by HPLC analysis. Obtained products were purified by preparative HPLC (preparative column: Eurospher 100 C18, 5 µm, 250 × 20 mm; solvent A: acetonitrile, solvent B: $H_2O$ + 0.2% v/v HCOOH; temperature 25 °C) and lyophilised from acetonitrile: water mixture. In some cases additional flash chromatography purification was performed.

**4-TMR-LTX (22):**

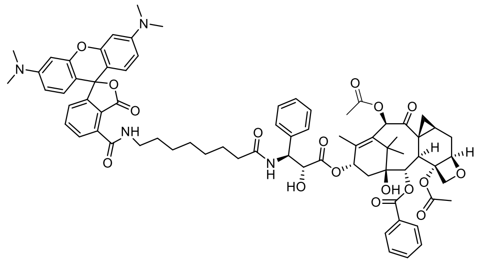

Purified by preparative HPLC (preparative column: Eurospher 100 C18, 5 µm, 250 × 20 mm; solvent A: acetonitrile, solvent B: $H_2O$ + 0.2% v/v HCOOH; temperature 25 °C, gradient A:B - 5 min 20:80 isocratic, 5-30 min 20:80 to 70:30 gradient). Additionally purified by flash column chromatography (Interchim Puriflash 12g, 15µm column, gradient 3% to 30% DCM – MeOH) and lyophilised from acetonitrile: water mixture. Yield 44% (5.6 mg) of light purple fluffy solid.

$^1$H NMR (400 MHz, d$_6$-dmso) δ 9.07 (t, J = 5.5 Hz, 1H), 8.50 (d, J = 9.1 Hz, 1H), 8.02 (d, J = 7.2 Hz, 2H), 7.84 (d, J = 7.5 Hz, 1H), 7.76 (t, J = 7.6 Hz, 1H), 7.68 (dd, J = 8.4, 6.1 Hz, 1H), 7.60 (t, J = 7.6 Hz, 2H), 7.37 – 7.30 (m, 4H), 7.23 (d, J = 7.5 Hz, 1H), 7.17 (tt, J = 7.3, 1.9 Hz, 1H), 6.61 (d, J = 9.5 Hz, 2H), 6.50 – 6.46 (m, 4H), 6.12 (s, 1H), 5.94 (t, J = 9.0 Hz, 1H), 5.50 – 5.33 (m, 2H), 5.32 (dd, J = 9.1, 5.4 Hz, 1H), 4.77 (s, 1H), 4.71 (d, J = 3.1 Hz, 1H), 4.49 (d, J = 5.5 Hz, 1H), 4.05 – 3.96 (m, 2H), 3.90 (d, J = 7.6 Hz, 1H), 3.30 – 3.27 (m, 2H), 2.94 (s, 12H), 2.32 – 2.25 (m, 4H), 2.17 – 1.86 (m, 9H), 1.74 (s, 3H), 1.59 – 1.44 (m, 5H), 1.35 – 1.20 (m, 7H), 1.11 (s, 3H), 1.08 (s, 3H).

$^1$H-$^{13}$C NMR ((400, 101) MHz, d$_6$-dmso) δ (8.00 130.09), (7.81 130.12), (7.74 135.60), (7.65 133.75), (7.57 129.19), (7.31 128.62), (7.30 127.50), (7.20 125.77), (7.15 127.38), (6.57 129.09),

(6.47 109.30), (6.46 98.37), (6.09 75.67), (5.92 69.90), (5.40 79.98), (5.29 55.23), (4.69 84.16), (4.46 73.96), (3.96 74.90), (3.88 37.97), (3.29 39.77), (2.91 38.96), (2.91 40.22), (2.26 22.36), (2.25 26.03), (2.13 35.83), (2.09 20.99), (1.81 14.82), (1.71 14.39), (1.52 29.37), (1.45 25.88), (1.30 26.91), (1.27 29.01), (1.21 29.10), (1.15 31.63), (1.08 21.82), (1.05 26.34).

ESI-MS, positive mode: m/z = 1285.7 $[M+H]^+$.

HRMS (ESI) calcd for $C_{73}H_{81}N_4O_{17}$ $[M+H]^+$ 1285.5591, found 1285.5591.

**5-TMR-LTX (23):**

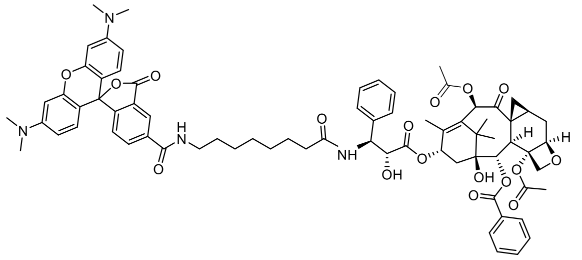

Purified by preparative HPLC (preparative column: Eurospher 100 C18, 5 µm, 250 × 20 mm; solvent A: acetonitrile, solvent B: $H_2O$ + 0.2% v/v HCOOH; temperature 25 °C, gradient A:B - 5 min 20:80 isocratic, 5-30 min 20:80 to 70:30 gradient), lyophilised from acetonitrile: water mixture. Yield 55% (7.1 mg) of deep purple fluffy solid.

$^1$H NMR (400 MHz, $d_6$-dmso) δ 8.76 (t, J = 5.6 Hz, 1H), 8.42 (dd, J = 1.7, 0.8 Hz, 1H), 8.33 (d, J = 9.2 Hz, 1H), 8.20 (dd, J = 8.0, 1.6 Hz, 1H), 8.05 – 7.97 (m, 2H), 7.66 (tt, J = 7.2, 1.4 Hz, 1H), 7.57 (t, J = 7.4 Hz, 2H), 7.38 – 7.22 (m, 5H), 7.21 – 7.11 (m, 1H), 6.52 – 6.41 (m, 6H), 6.10 (s, 1H), 5.93 (t, J = 9.1 Hz, 1H), 5.86 (d, J = 6.8 Hz, 1H), 5.41 (d, J = 7.6 Hz, 1H), 5.36 – 5.27 (m, 1H), 4.75 (s, 1H), 4.69 (d, J = 4.0 Hz, 1H), 4.47 (dd, J = 6.8, 5.2 Hz, 1H), 4.04 – 3.92 (m, 2H), 3.88 (d, J = 7.5 Hz, 1H), 3.26 (q, J = 6.7 Hz, 2H), 2.92 (s, 12H), 2.28 (s, 4H), 2.16 – 1.88 (m, 9H), 1.72 (s, 3H), 1.53 – 1.43 (m, 5H), 1.30 – 1.21 (m, 7H), 1.09 (s, 3H), 1.05 (s, 3H).

$^1$H-$^{13}$C NMR ((400, 101) MHz, $d_6$-dmso) δ (8.41 123.48), (8.20 134.91), (8.01 130.10), (7.65 133.75), (7.57 129.19), (7.32 128.53), (7.31 127.51), (7.27 124.57), (7.15 127.48), (6.48 98.42), (6.47 128.84), (6.47 109.42), (6.09 75.67), (5.93 70.01), (5.41 80.00), (5.32 55.15), (4.69 84.19), (4.47 73.98), (3.99 74.86), (3.88 37.99), (3.26 39.84), (2.92 40.22), (2.29, 1.92 26.02), (2.28 22.38), (2.13, 1.88 35.83), (2.10 21.01), (2.00, 1.51 15.24), (1.98 35.54), (1.72 14.41), (1.49 29.43), (1.45 25.86), (1.27 29.52), (1.24 26.90), (1.21 29.16), (1.15 31.66), (1.09 21.84), (1.05 26.35).

ESI-MS, positive mode: m/z = 1285.5 $[M+H]^+$.

HRMS (ESI) calcd for $C_{73}H_{81}N_4O_{17}$ $[M+H]^+$ 1285.5591, found 1285.5571.

**6-TMR-LTX (24):**

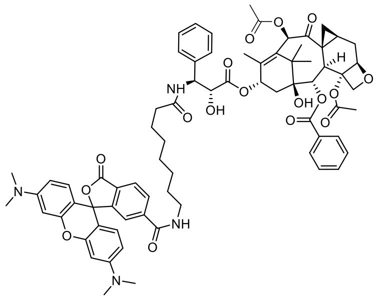

Purified by preparative HPLC (preparative column: Eurospher 100 C18, 5 µm, 250 × 20 mm; solvent A: acetonitrile, solvent B: $H_2O$ + 0.2% v/v HCOOH; temperature 25 °C, gradient A:B - 5 min 20:80 isocratic, 5-30 min 20:80 to 70:30 gradient), lyophilised from acetonitrile: water mixture. Yield 52% (6.7 mg) of deep purple fluffy solid.

$^1$H NMR (400 MHz, d$_6$-dmso) δ 8.62 (t, *J* = 5.6 Hz, 1H), 8.32 (d, *J* = 9.1 Hz, 1H), 8.12 (dd, *J* = 8.0, 1.3 Hz, 1H), 8.04 – 7.97 (m, 3H), 7.64 (t, *J* = 7.3 Hz, 1H), 7.60 – 7.52 (m, 3H), 7.32 – 7.26 (m, 4H), 7.12 (tt, *J* = 6.8, 1.9 Hz, 1H), 6.50 – 6.43 (m, 6H), 6.09 (s, 1H), 5.97 – 5.84 (m, 2H), 5.40 (d, *J* = 7.7 Hz, 1H), 5.30 (dd, *J* = 9.2, 5.2 Hz, 1H), 4.73 (s, 1H), 4.69 (d, *J* = 3.2 Hz, 1H), 4.45 (t, *J* = 4.1 Hz, 1H), 4.03 – 3.93 (m, 2H), 3.87 (d, *J* = 7.6 Hz, 1H), 3.13 (q, *J* = 6.7 Hz, 3H), 2.91 (s, 12H), 2.29 – 2.23 (m, 4H), 2.10 – 1.84 (m, 9H), 1.70 (s, 3H), 1.51 (dd, *J* = 7.2, 4.7 Hz, 1H), 1.42 – 1.34 (m, 4H), 1.21 – 1.12 (m, 7H), 1.08 (s, 3H), 1.04 (s, 3H).

$^1$H-$^{13}$C NMR ((400, 101) MHz, d$_6$-dmso) δ (8.12 129.57), (8.03 125.12), (7.99 130.07), (7.64 133.75), (7.60 122.59), (7.56 129.17), (7.30 128.56), (7.29 127.45), (7.13 127.44), (6.48 98.38), (6.48 128.90), (6.47 109.49), (6.09 75.65), (5.92 70.00), (5.40 79.98), (5.30 55.11), (4.69 84.20), (4.45 73.94), (3.97 74.88), (3.88 37.97), (3.13 39.84), (2.91 40.21), (2.28, 1.96 26.06), (2.26 22.36), (2.09 20.99), (2.09 35.82), (1.99, 1.51 15.11), (1.87 36.09), (1.70 14.40), (1.39 25.82), (1.38 29.27), (1.21 29.28), (1.15 31.66), (1.14 26.82), (1.13 28.93), (1.08 21.83), (1.04 26.33).

ESI-MS, positive mode: m/z = 1285.5 [M+H]$^+$.

HRMS (ESI) calcd for $C_{73}H_{81}N_4O_{17}$ [M+H]$^+$ 1285.5591, found 1285.5564.

**4-580CP-LTX (25):**

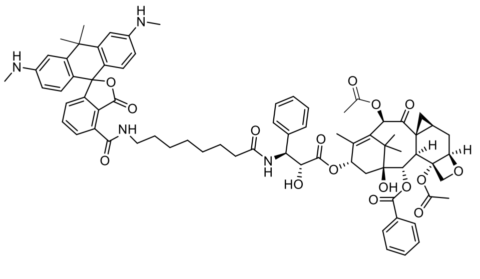

Purified by preparative HPLC (preparative column: Eurospher 100 C18, 5 µm, 250 × 20 mm; solvent A: acetonitrile, solvent B: $H_2O$ + 0.2% v/v HCOOH; temperature 25 °C, gradient A:B - 5 min 30:70 isocratic, 5-30 min 30:70 to 70:30 gradient). Additionally purified by flash column chromatography (Interchim Puriflash 12g, 15µm column, gradient 2% to 15% DCM – MeOH) and lyophilised from acetonitrile: water mixture. Yield 38% (4.9 mg) of violet fluffy solid.

$^1$H NMR (500 MHz, d$_6$-dmso) δ 9.14 (t, *J* = 5.5 Hz, 1H), 8.37 (d, *J* = 9.2 Hz, 1H), 8.03 (d, *J* = 7.1 Hz, 2H), 7.78 (dd, *J* = 7.5, 1.0 Hz, 1H), 7.71 – 7.65 (m, 2H), 7.60 (t, *J* = 7.6 Hz, 2H), 7.38 – 7.31 (m, 4H), 7.17 (tt, *J* = 7.0, 1.6 Hz, 1H), 7.05 (dd, *J* = 7.7, 1.0 Hz, 1H), 6.76 (d, *J* = 2.4 Hz, 2H), 6.45 (d, *J* = 8.6 Hz, 2H), 6.41 – 6.36 (m, 2H), 6.12 (s, 1H), 5.96 (t, *J* = 8.9 Hz, 1H), 5.91 – 5.80 (m, 3H), 5.43 (d, *J* = 7.7 Hz, 1H), 5.34 (dd, *J* = 9.2, 5.2 Hz, 1H), 4.78 (s, 1H), 4.72 (d, *J* = 4.1 Hz, 1H), 4.49 (dd, *J* =

6.8, 5.3 Hz, 1H), 4.05 – 3.97 (m, 2H), 3.91 (d, $J$ = 7.6 Hz, 1H), 3.30 – 3.16 (m, 2H), 2.70 (s, 3H), 2.69 (s, 3H), 2.31 – 2.25 (m, 4H), 2.18 – 1.88 (m, 9H), 1.75 (s, 3H), 1.74 (s, 3H), 1.65 (s, 3H), 1.57 – 1.45 (m, 5H), 1.37 – 1.23 (m, 7H), 1.11 (s, 3H), 1.08 (s, 3H).

$^{1}$H-$^{13}$C NMR ((500, 126) MHz, d$_6$-dmso) δ (8.03 129.69), (7.77 129.15), (7.69 134.93), (7.68 133.36), (7.60 128.78), (7.35 128.15), (7.33 127.07), (7.17 127.04), (7.06 125.16), (6.75 107.95), (6.46 128.47), (6.38 111.40), (6.11 75.23), (5.95 69.56), (5.44 79.53), (5.34 54.71), (4.71 83.75), (4.49 73.54), (4.02, 3.99 74.38), (3.91 37.53), (3.32 39.34), (2.69 29.56), (2.31,1.95 25.56), (2.30 21.95), (2.16 35.41), (2.12 20.57), (2.09, 1.89 35.55), (2.03, 1.53 14.88), (1.75 33.25), (1.73 13.98), (1.65 34.56), (1.55 28.92), (1.47 25.46), (1.36 28.30), (1.33 26.48), (1.23 28.88), (1.23 28.57), (1.11 21.41), (1.07 25.91).

ESI-MS, positive mode: m/z = 1283.7 [M+H]$^+$.

HRMS (ESI) calcd for $C_{74}H_{83}N_4O_{16}$ [M+H]$^+$ 1283.5799, found 1283.5796

**5-580CP-LTX (26):**

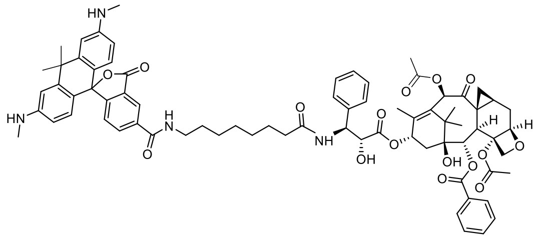

Purified by preparative HPLC (preparative column: Eurospher 100 C18, 5 μm, 250 × 20 mm; solvent A: acetonitrile, solvent B: $H_2O$ + 0.2% v/v HCOOH; temperature 25 °C, gradient A:B - 5 min 30:70 isocratic, 5-30 min 30:70 to 70:30 gradient). Lyophilised from acetonitrile: water mixture. Yield 48% (6.1 mg) of violet fluffy solid.

$^{1}$H NMR (400 MHz, d$_6$-dmso) δ 8.72 (t, $J$ = 5.6 Hz, 1H), 8.37 (s, 1H), 8.32 (d, $J$ = 9.2 Hz, 1H), 8.12 (d, $J$ = 8.2 Hz, 1H), 8.06 – 7.98 (m, 2H), 7.69 – 7.61 (m, 1H), 7.57 (t, $J$ = 7.4 Hz, 2H), 7.36 – 7.27 (m, 4H), 7.19 – 7.04 (m, 2H), 6.83 – 6.67 (m, 2H), 6.47 – 6.17 (m, 4H), 6.09 (s, 1H), 5.94 (d, $J$ = 8.7 Hz, 1H), 5.91 – 5.73 (m, 3H), 5.44 – 5.39 (m, 1H), 5.32 (dd, $J$ = 9.1, 5.1 Hz, 1H), 4.74 (s, 1H), 4.69 (d, $J$ = 4.1 Hz, 1H), 4.47 (dd, $J$ = 6.7, 5.2 Hz, 1H), 4.04 – 3.92 (m, 2H), 3.88 (d, $J$ = 7.6 Hz, 1H), 3.23 (d, $J$ = 6.5 Hz, 2H), 2.67 (s, 6H), 2.28 (s, 4H), 2.15 – 1.84 (m, 9H), 1.73 (s, 3H), 1.71 (s, 3H), 1.62 (s, 3H), 1.52 – 1.40 (m, 5H), 1.26 – 1.15 (m, 7H), 1.08 (s, 3H), 1.05 (s, 3H).

$^{1}$H-$^{13}$C NMR ((400, 101) MHz, d$_6$-dmso) δ (8.38 123.42), (8.12 134.62), (8.00 130.10), (7.64 133.74), (7.57 129.17), (7.30 128.57), (7.30 127.68), (7.14 127.46), (7.10 124.17), (6.75 108.49), (6.33 111.71), (6.32 128.63), (6.09 75.63), (5.93 70.03), (5.41 79.99), (5.32 55.13), (4.68 84.17), (4.46 73.96), (3.99 74.85), (3.88 37.92), (3.24 39.83), (2.67 29.98), (2.27 22.37), (2.24, 1.92 26.00), (2.12, 1.88 35.84), (2.10 20.98), (2.08, 35.82) (2.01, 1.51 15.48), (1.73 33.14), (1.71 14.39), (1.62 35.21), (1.48 29.49), (1.44 25.87), (1.23 26.89), (1.20 29.04), (1.14 31.70), (1.12 29.08), (1.08 21.83), (1.05 26.34).

ESI-MS, positive mode: m/z = 1283.7 [M+H]$^+$.

HRMS (ESI) calcd for $C_{74}H_{83}N_4O_{16}$ [M+H]$^+$ 1283.5799, found 1283.5792.

**6-580CP-LTX (27):**

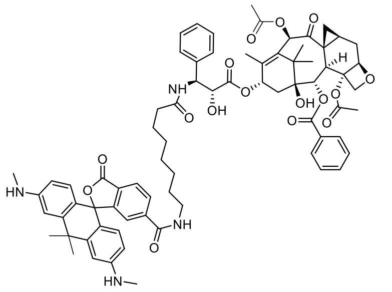

Was synthesised according to previously published procedure [1].

**4-610CP-CTX (28):**

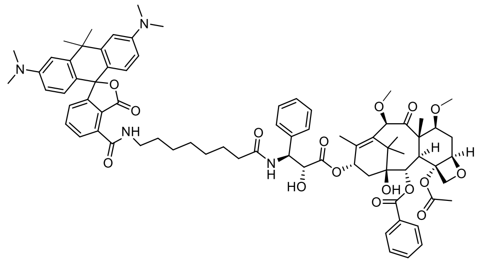

Purified by preparative HPLC (preparative column: Eurospher 100 C18, 5 µm, 250 × 20 mm; solvent A: acetonitrile, solvent B: $H_2O$ + 0.2% v/v HCOOH; temperature 25 °C, gradient A:B - 5 min 30:70 isocratic, 5-30 min 30:70 to 100:0 gradient). Lyophilised from acetonitrile: water mixture. Yield 46% (6.0 mg) of light blue fluffy solid.

$^1$H NMR (400 MHz, $d_6$-dmso) δ 9.09 (t, J = 5.5 Hz, 1H), 8.37 (d, J = 9.0 Hz, 1H), 7.98 – 7.91 (m, 2H), 7.75 (dd, J = 7.5, 1.0 Hz, 1H), 7.68 – 7.61 (m, 2H), 7.61 – 7.51 (m, 2H), 7.36 – 7.26 (m, 4H), 7.18 (tt, J = 7.1, 1.6 Hz, 1H), 7.01 (dd, J = 7.7, 1.0 Hz, 1H), 6.88 (d, J = 2.0 Hz, 2H), 6.60 – 6.49 (m, 4H), 6.00 – 5.87 (m, 2H), 5.36 (d, J = 7.1 Hz, 1H), 5.26 (dd, J = 9.1, 5.8 Hz, 1H), 4.92 (dd, J = 9.6, 2.1 Hz, 1H), 4.68 (s, 1H), 4.62 (s, 1H), 4.40 (t, J = 6.2 Hz, 1H), 3.99 (s, 2H), 3.72 (dd, J = 10.6, 6.6 Hz, 1H), 3.60 (d, J = 7.1 Hz, 1H), 3.34 – 3.31 (m, 2H), 3.27 (s, 3H), 3.18 (s, 3H), 2.91 (s, 12H), 2.69 – 2.57 (m, 1H), 2.21 (s, 3H), 2.15 (t, J = 7.3 Hz, 2H), 1.97 – 1.91 (m, 1H), 1.88 – 1.82 (m, 1H), 1.80 (s, 3H), 1.79 (s, 3H), 1.69 (s, 3H), 1.56 – 1.43 (m, 8H), 1.37 – 1.23 (m, 6H), 1.00 (s, 3H), 0.94 (s, 3H).

$^1$H-$^{13}$C NMR ((400, 101) MHz, $d_6$-dmso) δ (7.95 130.01), (7.75 129.60), (7.66 135.42), (7.64 133.80), (7.56 129.10), (7.33 128.58), (7.30 127.57), (7.18 127.54), (7.01 125.47), (6.88 109.55), (6.53 112.18), (6.53 128.79), (5.92 70.41), (5.36 74.79), (5.26 55.41), (4.91 83.63), (4.67 82.52), (4.40 74.01), (3.99 75.70), (3.72 80.67), (3.60 46.85), (3.30 39.78), (3.27 57.03), (3.18 57.07), (2.90 40.41), (2.62, 1.46 32.14), (2.21 22.81), (2.15 35.84), (1.94, 1.84 35.26), (1.80 14.44), (1.79 34.01), (1.68 34.94), (1.54 29.33), (1.49 10.58), (1.48 25.84), (1.32 26.90), (1.24 29.05), (1.22 29.13), (1.00 27.12), (0.94 21.63).

ESI-MS, positive mode: m/z = 1315.6 $[M+H]^+$.

HRMS (ESI) calcd for $C_{76}H_{91}N_4O_{16}$ $[M+H]^+$ 1315.6425, found 1315.6409.

**5-610CP-CTX (29):**

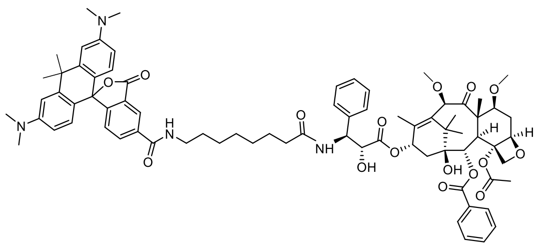

Purified by preparative HPLC (preparative column: Eurospher 100 C18, 5 µm, 250 × 20 mm; solvent A: acetonitrile, solvent B: $H_2O$ + 0.2% v/v HCOOH; temperature 25 °C, gradient A:B - 5 min 30:70 isocratic, 5-30 min 30:70 to 100:0 gradient). Lyophilised from acetonitrile: water mixture. Yield 44% (5.8 mg) of blue fluffy solid.

$^{1}$H NMR (400 MHz, d$_6$-dmso) δ 8.73 (t, *J* = 5.6 Hz, 1H), 8.41 – 8.38 (m, 1H), 8.36 (d, *J* = 9.0 Hz, 1H), 8.11 (dd, *J* = 8.0, 1.6 Hz, 1H), 7.97 – 7.92 (m, 2H), 7.67 – 7.62 (m, 1H), 7.59 – 7.54 (m, 2H), 7.35 – 7.28 (m, 4H), 7.17 (tt, *J* = 7.2, 1.6 Hz, 1H), 7.08 (d, *J* = 8.4 Hz, 1H), 6.89 (d, *J* = 2.6 Hz, 2H), 6.54 (dd, *J* = 9.0, 2.6 Hz, 2H), 6.41 (d, *J* = 8.8 Hz, 2H), 5.96 – 5.89 (m, 2H), 5.36 (d, *J* = 7.1 Hz, 1H), 5.26 (dd, *J* = 9.0, 5.8 Hz, 1H), 4.92 (dd, *J* = 9.5, 2.2 Hz, 2H), 4.67 (s, 1H), 4.62 (s, 1H), 4.43 – 4.35 (m, 2H), 3.99 (s, 2H), 3.75 – 3.70 (m, 1H), 3.60 (d, *J* = 7.1 Hz, 1H), 3.27 (s, 3H), 3.26 – 3.22 (m, 2H), 3.18 (s, 3H), 2.91 (s, 12H), 2.68 – 2.56 (m, 2H), 2.22 (s, 3H), 2.14 (t, *J* = 7.7 Hz, 2H), 1.97 – 1.91 (m, 1H), 1.87 – 1.82 (m, 1H), 1.80 (s, 3H), 1.79 (s, 3H), 1.68 (s, 3H), 1.48 (d, *J* = 8.8 Hz, 8H), 1.28 – 1.18 (m, 6H), 1.00 (s, 3H), 0.94 (s, 3H).

$^{1}$H-$^{13}$C NMR ((400, 101) MHz, d$_6$-dmso) δ (8.38 123.45), (8.10 134.77), (7.95 130.02), (7.65 133.78), (7.57 129.11), (7.33 128.58), (7.30 127.56), (7.18 127.53), (7.07 124.18), (6.89 109.58), (6.53 112.22), (6.42 128.53), (5.91 70.42), (5.36 74.80), (5.26 55.39), (4.92 83.65), (4.67 82.53), (4.40 73.99), (3.99 75.71), (3.73 80.68), (3.60 46.85), (3.27 57.03), (3.24 39.82), (3.18 57.08), (2.90 40.41), (2.62, 1.46 32.15), (2.22 22.82), (2.14 35.82), (1.94, 1.83 35.27), (1.80 14.44), (1.79 33.64), (1.68 35.09), (1.49 10.58), (1.49 29.43), (1.46 25.83), (1.25 26.89), (1.22 29.02), (1.18 29.00), (1.00 27.12), (0.94 21.63).

ESI-MS, positive mode: m/z = 1314.6 $[M+H]^+$.

HRMS (ESI) calcd for $C_{76}H_{91}N_4O_{16}$ $[M+H]^+$ 1315.6425, found 1315.6455.

**6-610CP-CTX (30):**

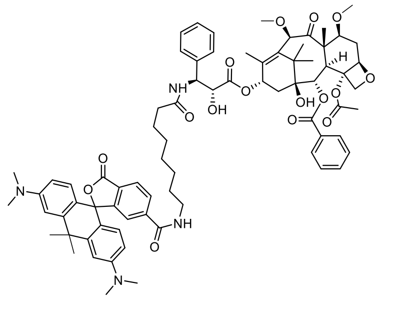

Purified by preparative HPLC (preparative column: Eurospher 100 C18, 5 µm, 250 × 20 mm; solvent A: acetonitrile, solvent B: $H_2O$ + 0.2% v/v HCOOH; temperature 25 °C, gradient A:B - 5 min 30:70 isocratic, 5-30 min 30:70 to 100:0 gradient). Lyophilised from acetonitrile: water mixture. Yield 48% (6.3 mg) of blue fluffy solid.

$^{1}$H NMR (400 MHz, d$_6$-dmso) δ 8.66 (t, *J* = 5.6 Hz, 1H), 8.34 (d, *J* = 9.1 Hz, 1H), 8.08 (dd, *J* = 8.1, 1.4 Hz, 1H), 8.03 (dd, *J* = 8.0, 0.7 Hz, 1H), 8.00 – 7.94 (m, 2H), 7.67 (tt, *J* = 7.2, 1.9 Hz, 1H), 7.59 (t, *J* = 7.7, 7.2 Hz, 2H), 7.41 (s, 1H), 7.38 – 7.28 (m, 4H), 7.19 (tt, *J*

= 7.2, 1.5 Hz, 1H), 6.93 (d, *J* = 2.6 Hz, 2H), 6.58 (dd, *J* = 8.9, 2.5 Hz, 2H), 6.43 (dd, *J* = 8.8, 0.9 Hz, 2H), 5.98 – 5.89 (m, 2H), 5.39 (d, *J* = 7.1 Hz, 1H), 5.28 (dd, *J* = 9.1, 5.6 Hz, 1H), 4.96 (dd, *J* = 10.1, 1.4 Hz, 1H), 4.70 (s, 1H), 4.64 (s, 1H), 4.42 (dd, *J* = 6.9, 5.7 Hz, 1H), 4.02 (s, 2H), 3.75 (dd, *J* = 10.6, 6.6 Hz, 1H), 3.63 (d, *J* = 7.1 Hz, 1H), 3.30 (s, 3H), 3.21 (s, 3H), 3.14 (q, *J* = 6.4 Hz, 2H), 2.94 (s, 12H), 2.71 – 2.61 (m, 1H), 2.24 (s, 3H), 2.13 (t, *J* = 7.1 Hz, 2H), 2.01 – 1.93 (m, 1H), 1.90 – 1.85 (m, 1H), 1.84 (s, 3H), 1.83 (s, 3H), 1.71 (s, 3H), 1.55 – 1.49 (m, 4H), 1.47 – 1.34 (m, 4H), 1.21 – 1.14 (m, 6H), 1.02 (s, 3H), 0.97 (s, 3H).

$^{1}H$-$^{13}C$ NMR ((400, 101) MHz, $d_6$-dmso) δ (8.08 128.13), (8.04 124.28), (7.98 129.27), (7.68 133.02), (7.59 128.36), (7.42 121.68), (7.35 127.83), (7.32 126.80), (7.19 126.77), (6.94 108.83), (6.57 111.52), (6.43 127.86), (5.94 69.68), (5.39 74.05), (5.28 54.58), (4.97 82.91), (4.71 81.78), (4.42 73.23), (4.03 74.96), (3.76 79.89), (3.63 46.11), (3.30 56.29), (3.21 56.32), (3.15 39.07), (2.94 39.65), (2.66, 1.50 31.36), (2.25 22.06), (2.14 35.04), (1.97, 1.86 34.54), (1.84 32.66), (1.83 13.68), (1.72 34.52), (1.53 9.84), (1.45 25.01), (1.41 28.60), (1.24 28.50), (1.19 26.09), (1.19 28.19), (1.02 26.36), (0.97 20.88).

ESI-MS, positive mode: m/z = 1315.7 $[M+H]^+$.

HRMS (ESI) calcd for $C_{76}H_{91}N_4O_{16}$ $[M+H]^+$ 1315.6425, found 1315.6410.

**4-SiR-CTX (31):**

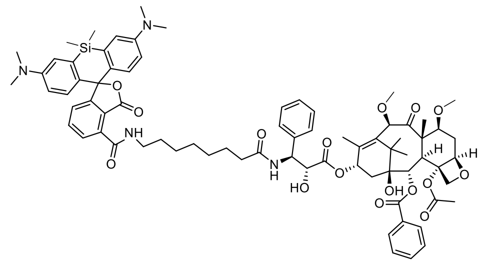

Purified by preparative HPLC (preparative column: Eurospher 100 C18, 5 µm, 250 × 20 mm; solvent A: acetonitrile, solvent B: $H_2O$ + 0.2% v/v HCOOH; temperature 25 °C, gradient A:B - 5 min 40:60 isocratic, 5-30 min 40:60 to 100:0 gradient) and lyophilised from acetonitrile and water mixture. Yield 51% (6.8 mg) of slightly blue fluffy solid.

$^{1}H$ NMR (400 MHz, $d_6$-dmso) δ 8.99 (t, *J* = 5.5 Hz, 1H), 8.34 (d, *J* = 9.0 Hz, 1H), 7.95 (d, *J* = 6.9 Hz, 2H), 7.74 – 7.71 (m, 2H), 7.66 – 7.61 (m, 1H), 7.56 (t, *J* = 7.5 Hz, 2H), 7.32 (ddd, *J* = 15.2, 8.2, 6.8 Hz, 4H), 7.21 (h, *J* = 2.9, 1.9 Hz, 2H), 6.97 (d, *J* = 2.8 Hz, 2H), 6.67 (dd, *J* = 9.0, 1.2 Hz, 2H), 6.63 – 6.59 (m, 2H), 5.92 (dd, *J* = 8.5, 6.4 Hz, 2H), 5.36 (d, *J* = 7.1 Hz, 1H), 5.26 (dd, *J* = 9.1, 5.8 Hz, 1H), 4.92 (d, *J* = 9.5 Hz, 1H), 4.67 (s, 1H), 4.62 (s, 1H), 4.40 (dd, *J* = 6.9, 5.8 Hz, 1H), 3.99 (s, 2H), 3.72 (dd, *J* = 10.6, 6.5 Hz, 1H), 3.60 (d, *J* = 7.1 Hz, 1H), 3.27 (s, 3H), 3.27 – 3.24 (m, 2H), 3.18 (s, 3H), 2.89 (s, 12H), 2.69 – 2.56 (m, 1H), 2.22 (s, 3H), 2.15 (t, *J* = 7.4 Hz, 2H), 1.94 (dd, *J* = 15.3, 8.9 Hz, 1H), 1.87 – 1.81 (m, 1H), 1.80 (s, 3H), 1.54 – 1.43 (m, 8H), 1.34 – 1.17 (m, 6H), 1.00 (s, 3H), 0.94 (s, 3H), 0.59 (s, 3H), 0.49 (s, 3H).

$^{1}H$-$^{13}C$ NMR ((400, 101) MHz, $d_6$-dmso) δ (7.95 129.99), (7.73 129.43), (7.73 134.94), (7.64 133.81), (7.56 129.12), (7.33 128.58), (7.30 127.56), (7.21 125.83), (7.18 127.50), (6.97 116.66), (6.66 128.36), (6.61 114.05), (5.92 70.41), (5.36 74.79), (5.26 55.36), (4.92 83.65), (4.67 82.53), (4.39 74.02),

(3.99 75.71), (3.73 80.65), (3.60 46.85), (3.27 57.03), (3.27 39.77), (3.18 57.09), (2.88 40.22), (2.62, 1.47 32.13), (2.22 22.83), (2.14 35.85), (1.94, 1.85 35.32), (1.80 14.45), (1.51 29.28), (1.49 10.58), (1.47 25.86), (1.30 26.87), (1.22 29.10), (1.21 29.15), (1.00 27.13), (0.94 21.64), (0.59 -0.75), (0.49 0.41).

ESI-MS, positive mode: m/z = 1331.6 [M+H]$^+$.

HRMS (ESI) calcd for $C_{75}H_{91}N_4O_{16}Si$ [M+H]$^+$ 1331.6194, found 1331.6184.

**5-SiR-CTX (32):**

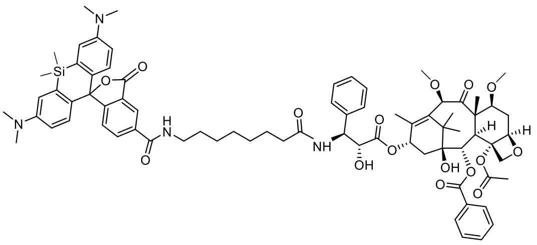

Purified by preparative HPLC (preparative column: Eurospher 100 C18, 5 µm, 250 × 20 mm; solvent A: acetonitrile, solvent B: $H_2O$ + 0.2% v/v HCOOH; temperature 25 °C, gradient A:B - 5 min 40:60 isocratic, 5-30 min 40:60 to 100:0 gradient) and lyophilised from acetonitrile and water mixture. Yield 49% (6.5 mg) of slightly blue fluffy solid.

$^1$H NMR (500 MHz, $d_6$-dmso) δ 8.75 (t, *J* = 5.6 Hz, 1H), 8.42 – 8.35 (m, 2H), 8.20 (dd, *J* = 8.0, 1.6 Hz, 1H), 7.97 (dd, *J* = 7.0, 1.2 Hz, 2H), 7.67 (tt, *J* = 7.3, 1.1 Hz, 1H), 7.58 (t, *J* = 7.6 Hz, 2H), 7.37 – 7.29 (m, 5H), 7.19 (tt, *J* = 7.3, 1.0 Hz, 1H), 7.00 (d, *J* = 2.0 Hz, 2H), 6.63 – 6.57 (m, 4H), 6.06 – 5.83 (m, 2H), 5.38 (d, *J* = 7.1 Hz, 1H), 5.28 (dd, *J* = 9.0, 5.8 Hz, 1H), 4.94 (dd, *J* = 9.7, 2.7 Hz, 1H), 4.69 (s, 1H), 4.65 (s, 1H), 4.42 (t, *J* = 5.9 Hz, 1H), 4.01 (s, 2H), 3.74 (dd, *J* = 10.6, 6.7 Hz, 1H), 3.62 (d, *J* = 7.1 Hz, 1H), 3.29 (s, 3H), 3.28 – 3.24 (m, 2H), 3.20 (s, 3H), 2.90 (s, 12H), 2.69 – 2.60 (m, 1H), 2.24 (s, 3H), 2.19 – 2.13 (m, 2H), 1.96 (dd, *J* = 15.3, 9.2 Hz, 1H), 1.88 – 1.83 (m, 1H), 1.82 (s, 3H), 1.50 (d, *J* = 11.4 Hz, 8H), 1.25 (dd, *J* = 14.6, 8.8 Hz, 6H), 1.02 (s, 3H), 0.96 (s, 3H), 0.62 (s, 3H), 0.51 (s, 3H).

$^1$H-$^{13}$C NMR ((500, 126) MHz, $d_6$-dmso) δ (8.40 123.74), (8.20 133.96), (7.97 129.66), (7.68 133.43), (7.59 128.75), (7.36 128.21), (7.33 124.51), (7.32 127.21), (7.20 127.17), (7.01 116.47), (6.61 113.59), (6.59 127.72), (5.94 70.02), (5.38 74.39), (5.29 55.00), (4.94 83.23), (4.70 82.16), (4.43 73.63), (4.02 75.32), (3.75 80.30), (3.63 46.49), (3.30 56.66), (3.27 39.43), (3.20 56.70), (2.91 39.83), (2.65, 1.49 31.76), (2.25 22.46), (2.17 35.45), (1.97, 1.85 34.91), (1.83 14.08), (1.51 10.21), (1.51 29.05), (1.49 25.47), (1.27 26.50), (1.26 28.69), (1.24 28.70), (1.03 26.75), (0.97 21.26), (0.62 -1.40), (0.52 0.16).

ESI-MS, positive mode: m/z = 1331.6 [M+H]$^+$.

HRMS (ESI) calcd for $C_{75}H_{91}N_4O_{16}$ [M+H]$^+$ 1331.6194, found 1331.6209.

**6-SiR-CTX (33)** [1]**:**

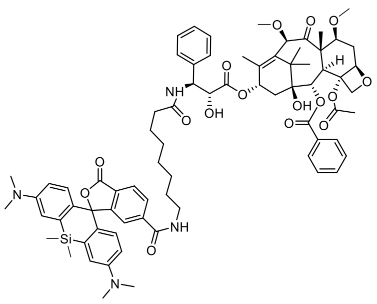

Purified by preparative HPLC (preparative column: Eurospher 100 C18, 5 µm, 250 × 20 mm; solvent A: acetonitrile, solvent B: $H_2O$ + 0.2% v/v HCOOH; temperature 25 °C, gradient A:B - 5 min 40:60 isocratic, 5-30 min 40:60 to 100:0 gradient) and lyophilised from acetonitrile and water mixture. Yield 49% (6.2 mg) of slightly blue fluffy solid. ESI-MS, positive mode: m/z = 1331.6 $[M+H]^+$.

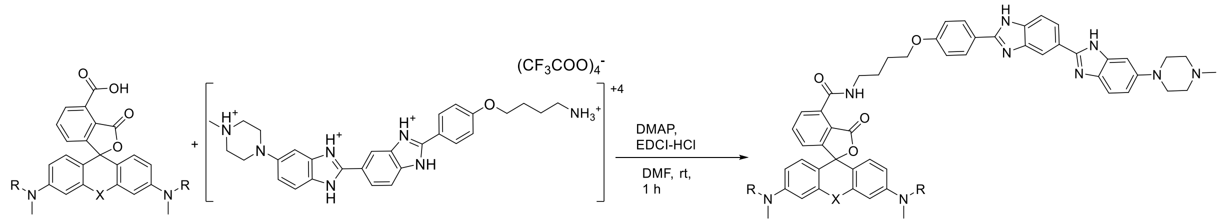

**4-TMR-Hoechst (34):**

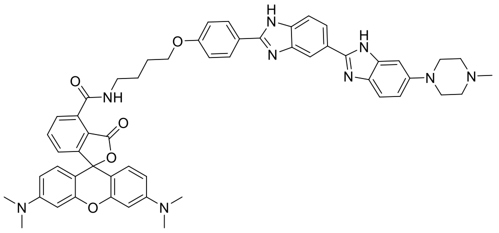

**4-TMR-COOH-**TFA salt (0.0184 mmol, 10 mg, 1 eq.), EDCI-HCl (0.0276 mmol, 8.4mg, 1.5 eq.) DMAP (0.11 mmol, 13.4 mg, 6 eq.) and Hoechst-C4-$NH_2$ [8] (0.0276, 26 mg, 1.5 eq.) were dissolved in 1 mL of dry DMF and stirred for 1 hour. Then DMF was removed at rt under reduced pressure. The product was purified by preparative HPLC (preparative column: Eurospher 100 C18, 5 µm, 250 × 20 mm; solvent A: acetonitrile, solvent B: $H_2O$ + 0.2% v/v HCOOH; temperature 25 °C, gradient A:B - 5 min 20:80 isocratic, 5-30 min 20:80 to 70:30 gradient). Water acetonitrile mixture was removed by rotary evaporator and product was purified one more time by flash chromatography (silica-gel cartridge: Interchim Puriflash 12g, 15µm column, gradient 20% to 80% $CH_2Cl_2$ – $CH_2Cl_2$:MeOH: $NH_{3(aq)}$ [85:15:2]). Solvents were removed and a product was lyophilised from water acetonitrile mixture to obtain 9 mg (54%) of purple solid.

$^1$H NMR (400 MHz, $CD_3OD$ + $CF_3COOD$) δ 8.58 (dd, J = 1.7, 0.7 Hz, 1H), 8.25 (dd, J = 8.7, 1.7 Hz, 1H), 8.18 (d, J = 9.0 Hz, 2H), 8.07 (dd, J = 8.7, 0.7 Hz, 1H), 7.83 – 7.79 (m, 2H), 7.76 (d, J = 9.1 Hz, 1H), 7.51 (dd, J = 6.2, 2.7 Hz, 1H), 7.44 (dd, J = 9.2, 2.2 Hz, 1H), 7.36 (d, J = 2.2 Hz, 1H), 7.29 (d, J = 9.0 Hz, 2H), 7.23 (d, J = 9.5 Hz, 2H), 7.07 (dd, J = 9.5, 2.5 Hz, 2H), 6.97 (d, J = 2.4 Hz, 2H), 4.22 (t, J = 6.2 Hz, 2H), 3.97 (d, J = 13.2 Hz, 2H), 3.68 (d, J = 12.1 Hz, 2H), 3.49 (t, J = 6.9 Hz, 2H), 3.36 (d, J = 11.3 Hz, 2H), 3.30 (s, 12H), 3.19 (d, J = 11.9 Hz, 2H), 3.00 (s, 3H), 2.03 – 1.94 (m, 2H), 1.87 (dd, J = 8.7, 6.0 Hz, 2H).

$^{13}C$ NMR (101 MHz, $CD_3OD$ + $CF_3COOD$) δ 171.1, 169.2, 165.6, 159.1, 158.8, 154.3, 151.1, 149.5, 139.2, 134.6, 133.9, 133.7, 132.5, 131.9, 131.5, 130.2, 128.0, 126.3, 121.7, 120.9, 120.0, 118.1, 117.5, 117.2, 116.4, 115.7, 115.5, 115.2, 115.0, 114.6, 112.4, 101.0, 97.5, 69.5, 54.6, 48.4 (visible in HSQC), 43.6, 41.0, 40.7, 27.5, 26.9.

ESI-MS, positive mode: m/z = 908.5 $[M+H]^+$.

HRMS (ESI) calcd for $C_{54}H_{54}N_9O_5$ $[M+H]^+$ 908.4242, found 908.4230.

**5-TMR-Hoechst (35):**

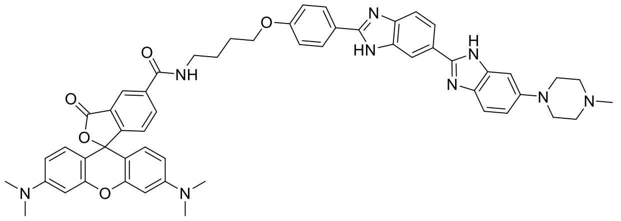

Was synthesised according to previously published procedure. [8]

**6-TMR-Hoechst (36):**

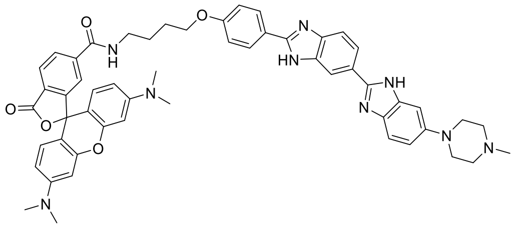

Was synthesised according to previously published procedure. [8]

**4-580CP-Hoechst (37):**

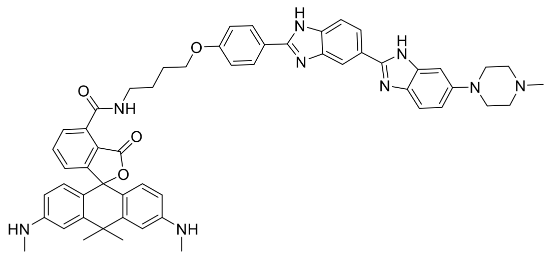

**4-580CP-COOH-**TFA salt (0.0184 mmol, 10 mg, 1 eq.), EDCI-HCl (0.0276 mmol, 8.4mg, 1.5 eq.) DMAP (0.11 mmol, 13.4 mg, 6 eq.) and Hoechst-C4-$NH_2$ [8] (0.0276, 26 mg, 1.5 eq.) were dissolved in 1 mL of dry DMF and stirred for 1 hour. Then DMF was removed at rt under reduced pressure. The product was purified by preparative HPLC (preparative column: Eurospher 100 C18, 5 μm, 250 × 20 mm; solvent A: acetonitrile, solvent B: $H_2O$ + 0.2% v/v HCOOH; temperature 25 °C, gradient A:B - 5 min 20:80 isocratic, 5-30 min 20:80 to 70:30 gradient). Water acetonitrile mixture was removed by rotary evaporator and product was purified one more time by flash chromatography (silica-gel cartridge: Interchim Puriflash 12g, 15μm column, gradient 20% to 80% $CH_2Cl_2$ – $CH_2Cl_2$:MeOH: $NH_{3(aq)}$ [85:15:2]). Solvents were removed and a product was lyophilised from water acetonitrile mixture to obtain 6 mg (36%) of dark-violet solid.

$^1H$ NMR (400 MHz, $CD_3OD$ + $CF_3COOD$) δ 8.39 (s, 1H), 8.17 – 8.04 (m, 3H), 7.95 (d, J = 8.6 Hz, 1H), 7.86 – 7.78 (m, 2H), 7.76 (d, J = 8.9 Hz, 1H), 7.48 – 7.39 (m, 2H), 7.36 (s, 1H), 7.21 – 7.12

(m, 4H), 7.09 (d, J = 9.1 Hz, 2H), 6.68 (dd, J = 9.1, 2.3 Hz, 2H), 4.13 (t, J = 5.7 Hz, 2H), 3.98 (d, J = 13.1 Hz, 2H), 3.77 – 3.71 (m, 2H), 3.52 (t, J = 6.0 Hz, 2H), 3.35 (d, J = 10.6 Hz, 2H), 3.27 (d, J = 13.3 Hz, 2H), 3.07 (s, 6H), 3.03 (s, 3H), 1.99 – 1.85 (m, 4H), 1.83 (s, 3H), 1.70 (s, 3H).

$^{13}$C NMR (101 MHz, $CD_3OD$ + $CF_3COOD$) δ 169.7, 168.6, 162.7, 161.7, 161.4, 157.3, 156.7, 153.7, 149.2, 148.7, 137.4, 136.8, 136.7, 133.2, 132.0, 131.1, 130.3, 129.2, 128.1, 126.9, 124.7, 123.2, 120.7, 118.6, 118.1, 115.2, 115.2, 114.3, 113.7, 112.7, 111.0, 99.7, 67.8, 53.2, 47.1, 42.3, 41.1, 39.4, 34.2, 30.8, 28.9, 26.2, 25.5.

ESI-MS, positive mode: m/z = 906.4 $[M+H]^+$.

HRMS (ESI) calcd for $C_{55}H_{56}N_9O_4$ $[M+H]^+$ 906.4450, found 906.4455.

**5-580CP-Hoechst (38):**

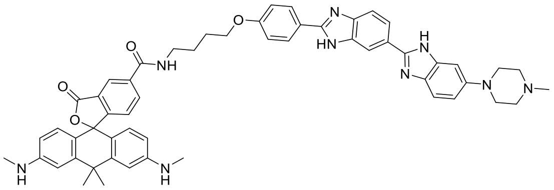

Was synthesised according to previously published procedure. [8]

**6-580CP-Hoechst (39):**

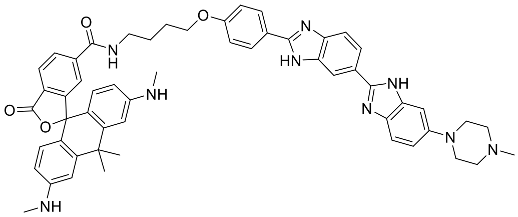

Was synthesised according to previously published procedure. [8]

**4-610CP-C5-COOH (SI-5):**

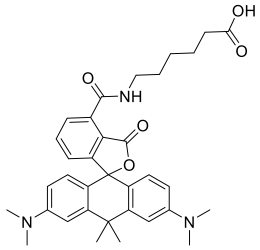

**4-610CP-COOH**-TFA salt (0.0175 mmol, 10 mg, 1 eq.), DIPEA (15 µL, 0.0875 mmol, 5 eq.) and HBTU (0.021 mmol, 8.0 mg, 1.2 eq.) were dissolved in 400 µL of dry DMSO and stirred at room temperature for 15 min. A solution of 6-aminocaproic acid (0.0175 mmol, 2.3 mg, 1 eq.) in 200 µL of DMSO:$H_2O$ mixture (1:1) was added to the reaction mixture and stirring continued for 1 hour. Reaction mixture was neutralised with formic acid and product was purified by preparative HPLC (preparative column: Eurospher 100 C18, 5 µm, 250 × 20 mm; solvent A: acetonitrile, solvent B: H2O + 0.2% v/v HCOOH; temperature 25 °C, gradient A:B - 5 min 40:60 isocratic, 5-30 min 40:60 to 80:20 gradient). Product lyophilised from acetonitrile: water mixture to obtain 7 mg (70%) of blue powder.

$^1$H NMR (400 MHz, $CD_3CN$) δ 9.74 (t, J = 5.1 Hz, 1H), 8.28 (dd, J = 7.7, 1.0 Hz, 1H), 7.72 (t, J = 7.7 Hz, 1H), 7.08 (dd, J = 7.7, 1.0 Hz, 1H), 6.96 (d, J = 2.3 Hz, 2H), 6.64 – 6.53 (m, 4H), 3.49 (q, J

= 7.0 Hz, 2H), 2.97 (s, 13H), 2.31 (t, J = 7.4 Hz, 2H), 2.20 (s, 1H), 1.86 (s, 3H), 1.75 (s, 3H), 1.73 – 1.61 (m, 4H), 1.56 – 1.44 (m, 2H).

$^{13}$C NMR (101 MHz, $CD_3CN$) δ 175.08, 172.55, 164.64, 158.41, 152.15, 147.76, 136.10, 134.66, 132.21, 129.55, 127.07, 123.70, 119.05, 112.78, 110.31, 40.67, 40.59, 39.27, 35.31, 34.14, 33.58, 29.72, 27.25, 25.32.

ESI-MS, positive mode: m/z = 570.3 $[M+H]^+$.

HRMS (ESI) calcd for $C_{34}H_{40}N_3O_5$ $[M+H]^+$ 570.2962, found 570.2965.

**4-610CP-JAS (40):**

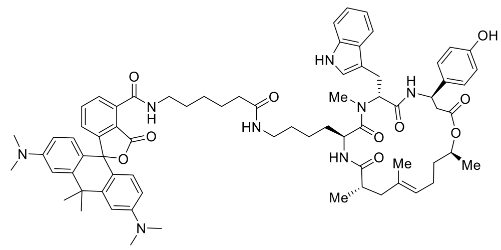

**4-610CP-C5-COOH (SI-5)** (0.00702 mmol, 4.0 mg), TSTU (0.00913 mmol, 2.7 mg) and DIPEA (12 µL, 0.0688 mmol) were dissolved in 500 µL of MeCN and stirred for 1 hour. Then MeCN was removed by rotary evaporator and obtained product was purified by flash chromatography (silica-gel cartridge: Interchim Puriflash 12g, 15µm column, gradient 20% to 100% DCM – EtOAc). The solvents were removed and the obtained NHS ester was redissolved in 400 µL of MeCN followed by the addition of deprotected des-bromo-des-methyl-Lys-jasplakinolide[10] (0.00342 mmol, 2.3 mg) and DIPEA (12 µL, 0.0688 mmol). The reaction mixture was stirred for 1 hour and purified by preparative HPLC (preparative column: Eurospher 100 C18, 5 µm, 250 × 20 mm; solvent A: acetonitrile, solvent B: H2O + 0.2% v/v HCOOH; temperature 25 °C, gradient A:B - 5 min 30:70 isocratic, 5-30 min 30:70 to 100:0 gradient). Product lyophilised from acetonitrile: water mixture to obtain 1.8 mg (48%) of light blue powder.

$^1$H NMR (400 MHz, $d_6$-dmso) δ 10.79 (d, *J* = 1.8 Hz, 1H), 9.27 (s, 1H), 9.11 (t, *J* = 5.5 Hz, 1H), 8.61 (d, *J* = 8.8 Hz, 1H), 7.75 (d, *J* = 7.6 Hz, 1H), 7.70 – 7.60 (m, 4H), 7.26 (d, *J* = 8.1 Hz, 1H), 7.10 (d, *J* = 8.6 Hz, 2H), 7.03 (d, *J* = 2.3 Hz, 1H), 7.02 – 6.97 (m, 2H), 6.92 (t, *J* = 7.2 Hz, 1H), 6.88 (d, *J* = 2.1 Hz, 2H), 6.67 (d, *J* = 8.5 Hz, 2H), 6.58 – 6.50 (m, 4H), 5.49 (dd, *J* = 11.3, 5.1 Hz, 1H), 5.22 – 5.11 (m, 1H), 4.89 (t, *J* = 7.1 Hz, 1H), 4.64 (h, *J* = 6.4 Hz, 1H), 4.58 – 4.44 (m, 1H), 3.35 – 3.31 (m, 2H), 3.08 – 2.96 (m, 4H), 2.91 (s, 12H), 2.89 – 2.71 (m, 3H), 2.65 (dd, *J* = 14.7, 11.3 Hz, 1H), 2.55 (dd, *J* = 14.7, 3.2 Hz, 1H), 2.53 – 2.48 (m, 2H), 2.14 (dd, *J* = 14.6, 11.5 Hz, 1H), 2.05 (t, *J* = 7.5 Hz, 2H), 1.86 – 1.75 (m, 5H), 1.74 – 1.65 (m, 4H), 1.61 – 1.49 (m, 4H), 1.45 (s, 3H), 1.38 – 1.33 (m, 2H), 1.22 – 1.20 (m, 2H), 1.13 (d, *J* = 6.3 Hz, 3H), 1.10 – 1.00 (m, 2H), 0.90 (d, *J* = 6.8 Hz, 3H), 0.85 – 0.71 (m, 4H).

ESI-MS, positive mode: m/z = 1247.6 $[M+Na]^+$.

HRMS (ESI) calcd for $C_{72}H_{89}N_8O_{10}$ $[M+H]^+$ 1225.6696, found 1225.6689.

**5-610CP-JAS (41):**

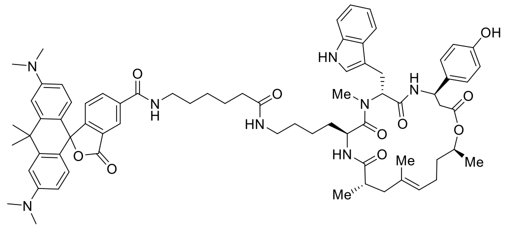

Was synthesised according to previously published procedure[14].

**6-610CP-JAS (42):**

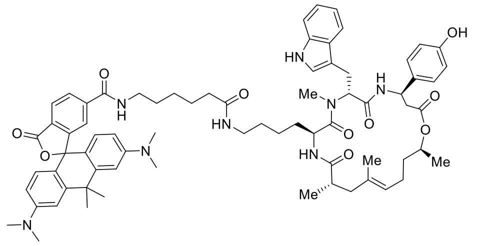

Was synthesised according to previously published procedure[14].

#### Supplementary references

1. Lukinavičius G, Mitronova GY, Schnorrenberg S, Butkevich AN, Barthel H, Belov VN*, et al.* Fluorescent dyes and probes for super-resolution microscopy of microtubules and tracheoles in living cells and tissues. *Chem Sci* 2018, **9**(13)**:** 3324-3334.
2. Breusegem SY, Clegg RM, Loontiens FG. Base-sequence specificity of Hoechst 33258 and DAPI binding to five (A/T)4 DNA sites with kinetic evidence for more than one high-affinity Hoechst 33258-AATT complex. *J Mol Biol* 2002, **315**(5)**:** 1049-1061.
3. Åkerlöf G, Short AO. The Dielectric Constant of Dioxane—Water Mixtures between 0 and 80°. *J Am Chem Soc* 1936, **58**(7)**:** 1241–1243.
4. Schindelin J, Arganda-Carreras I, Frise E, Kaynig V, Longair M, Pietzsch T*, et al.* Fiji: an open-source platform for biological-image analysis. *Nat Methods* 2012, **9**(7)**:** 676-682.
5. Carpenter AE, Jones TR, Lamprecht MR, Clarke C, Kang IH, Friman O*, et al.* CellProfiler: image analysis software for identifying and quantifying cell phenotypes. *Genome Biol* 2006, **7**(10)**:** R100.
6. Gottlieb HE, Kotlyar V, Nudelman A. NMR Chemical Shifts of Common Laboratory Solvents as Trace Impurities. *J Org Chem* 1997, **62**(21)**:** 7512-7515.
7. Ren S, Wang Y, Wang J, Gao D, Zhang M, Ding N*, et al.* Synthesis and biological evaluation of novel larotaxel analogues. *Eur J Med Chem* 2018, **156:** 692-710.
8. Bucevičius J, Keller-Findeisen J, Gilat T, Hell SW, Lukinavičius G. Rhodamine-Hoechst positional isomers for highly efficient staining of heterochromatin. *Chem Sci* 2019, **10**(7)**:** 1962-1970.
9. Butkevich AN, Mitronova GY, Sidenstein SC, Klocke JL, Kamin D, Meineke DN*, et al.* Fluorescent Rhodamines and Fluorogenic Carbopyronines for Super-Resolution STED Microscopy in Living Cells. *Angew Chem Int Ed Engl* 2016, **55**(10)**:** 3290-3294.
10. Tannert R, Milroy LG, Ellinger B, Hu TS, Arndt HD, Waldmann H. Synthesis and structure-activity correlation of natural-product inspired cyclodepsipeptides stabilizing F-actin. *J Am Chem Soc* 2010, **132**(9)**:** 3063-3077.
11. Martinez-Peragon A, Miguel D, Jurado R, Justicia J, Alvarez-Pez JM, Cuerva JM*, et al.* Synthesis and photophysics of a new family of fluorescent 9-alkyl-substituted xanthenones. *Chemistry* 2014, **20**(2)**:** 447-455.
12. Grimm JB, Sung AJ, Legant WR, Hulamm P, Matlosz SM, Betzig E*, et al.* Carbofluoresceins and carborhodamines as scaffolds for high-contrast fluorogenic probes. *ACS Chem Biol* 2013, **8**(6)**:** 1303-1310.
13. Butkevich AN, Belov VN, Kolmakov K, Sokolov VV, Shojaei H, Sidenstein SC*, et al.* Hydroxylated Fluorescent Dyes for Live-Cell Labeling: Synthesis, Spectra and Super-Resolution STED. *Chemistry – A European Journal* 2017, **23**(50)**:** 12114-12119.
14. Gerasimaitė R, Seikowski J, Schimpfhauser J, Kostiuk G, Gilat T, D'Este E*, et al.* Overcoming efflux of fluorescent probes for actin imaging in living cells. *bioRxiv* 2020**:** 2020.2002.2017.951525.

### Copies of NMR spectra

#### Di-tert-butyl 3-bromophthalate (1)

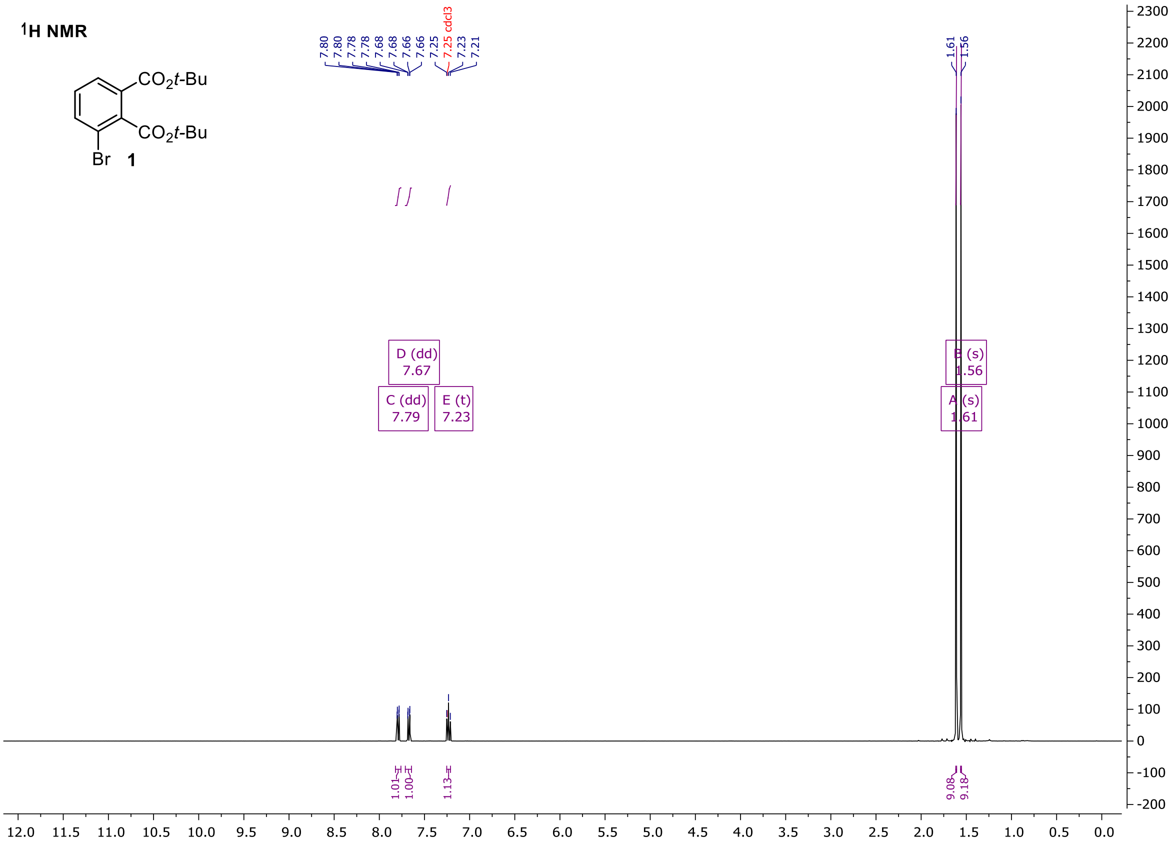

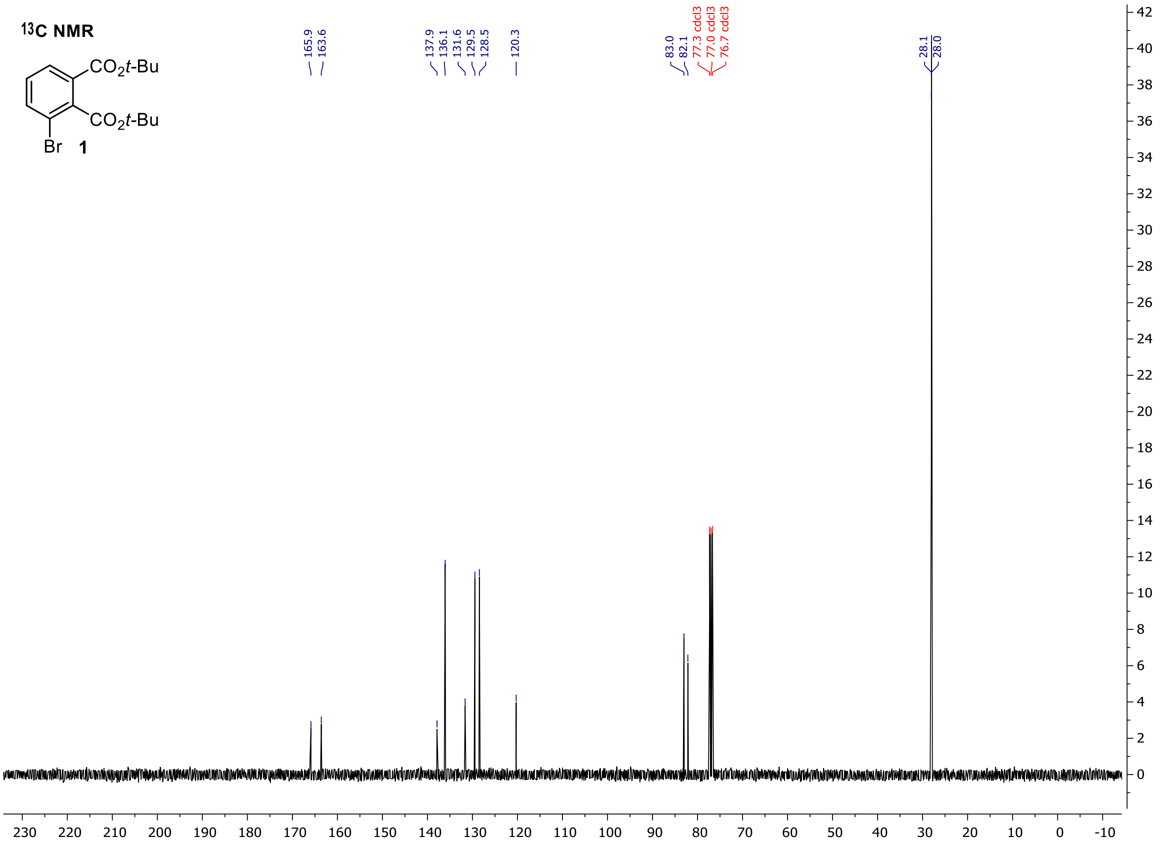

**Tert-butyl 3',6'-bis((tert-butyldimethylsilyl)oxy)-3-oxo-3H-spiro[isobenzofuran-1,9'-xanthene]-4-carboxylate (5)**

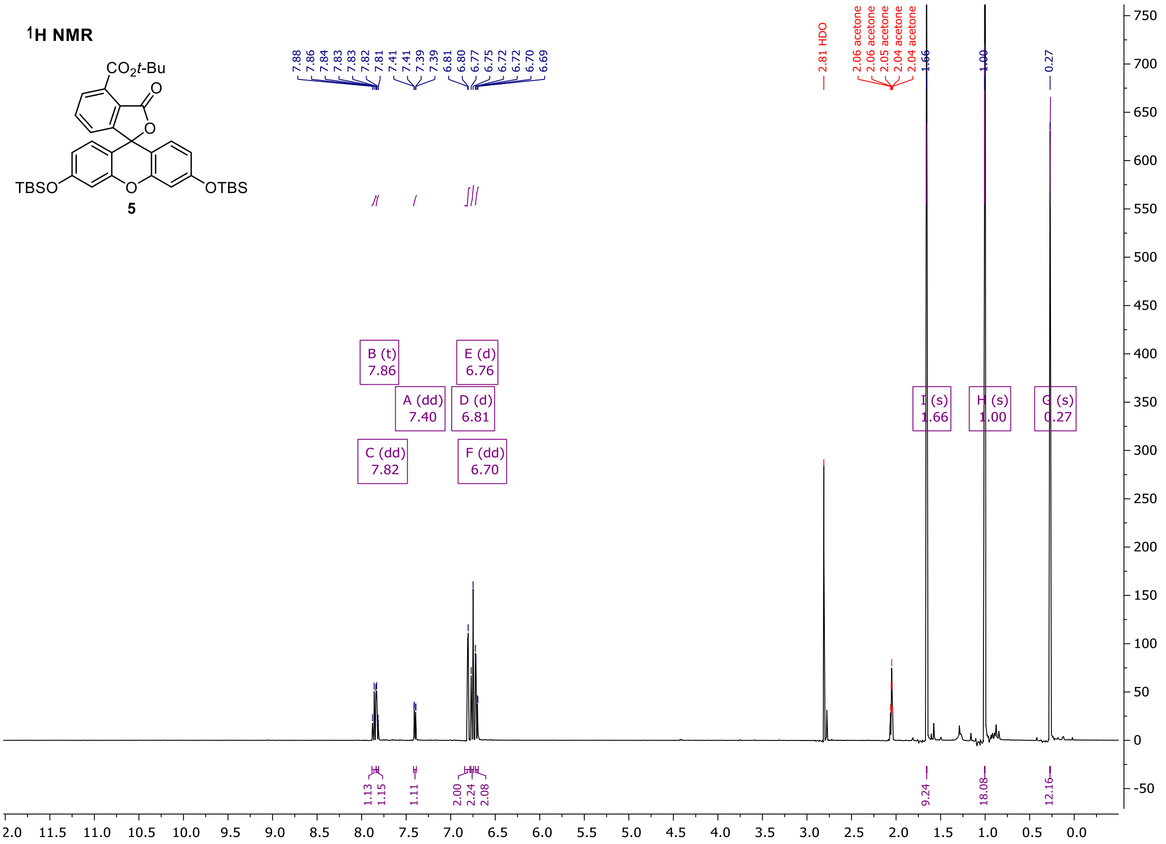

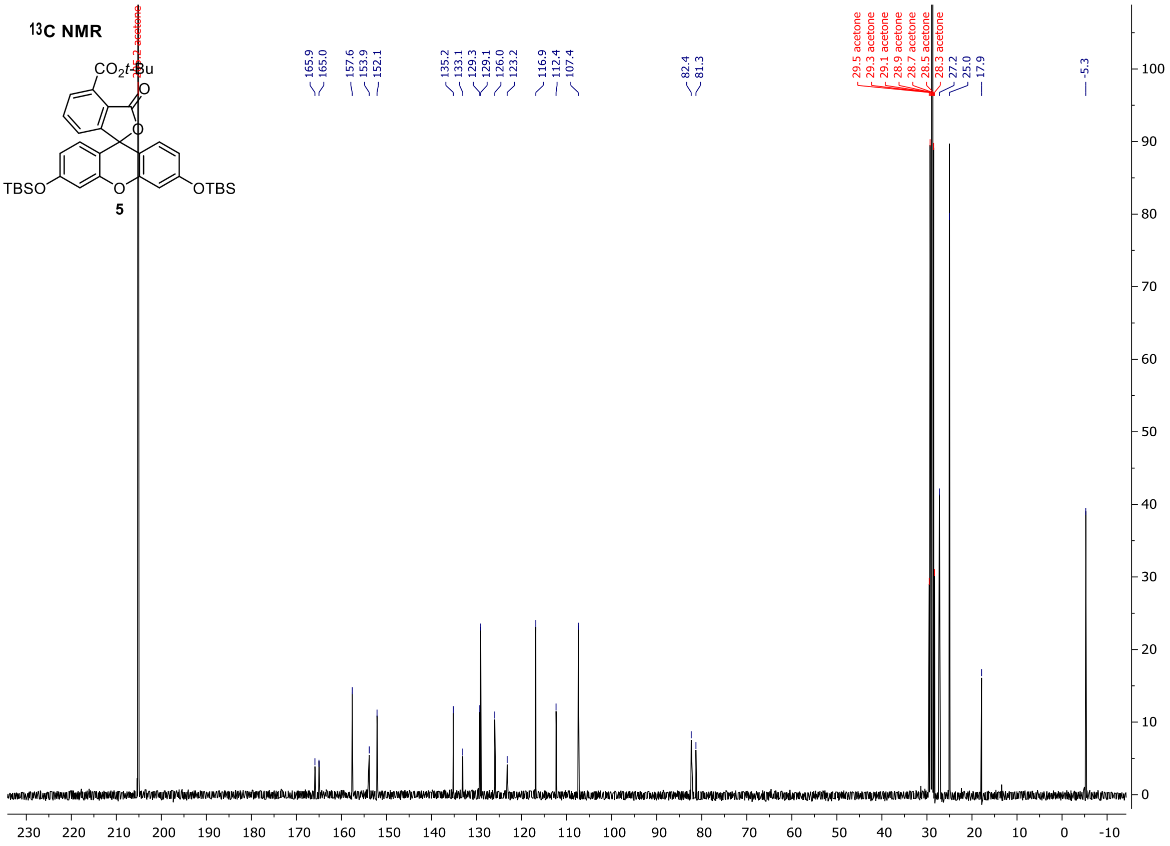

**Tert-butyl 3,6-bis((tert-butyldimethylsilyl)oxy)-10,10-dimethyl-3'-oxo-3'H,10H-spiro[anthracene-9,1'-isobenzofuran]-4'-carboxylate (6)**

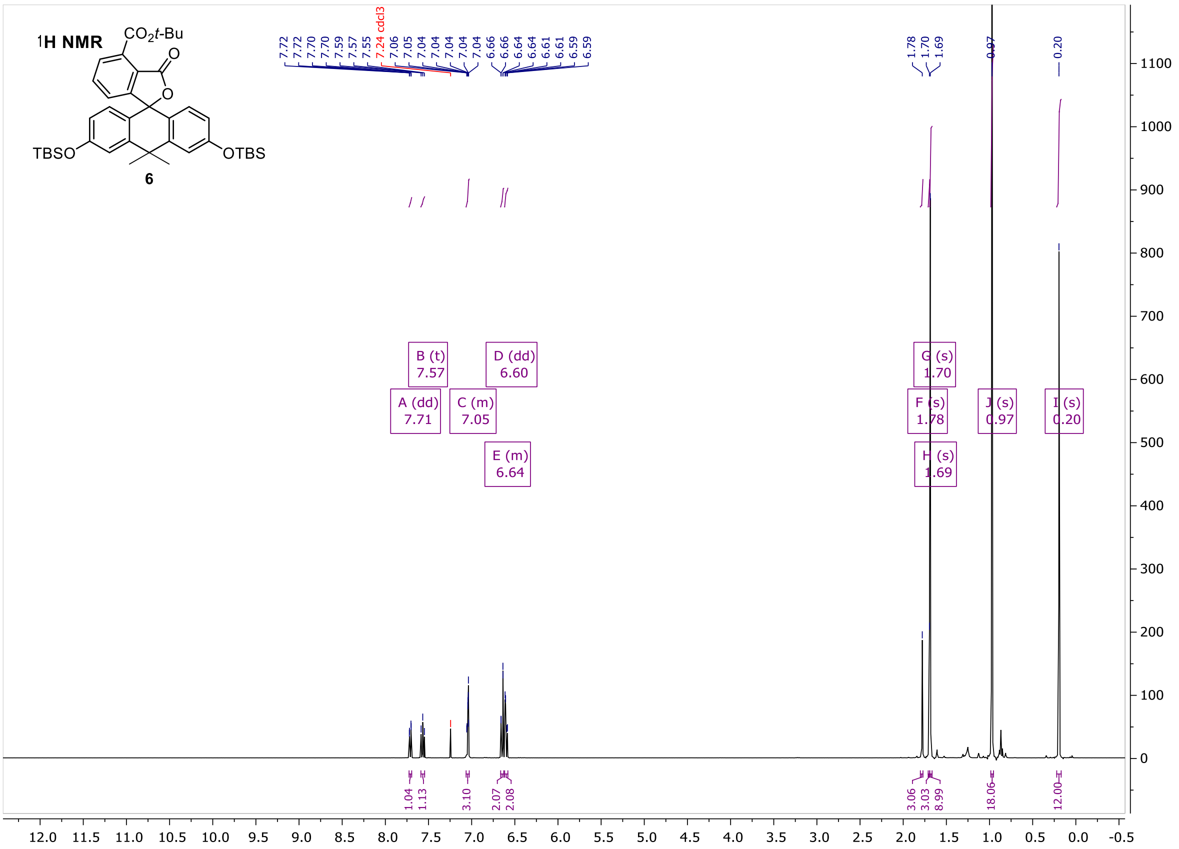

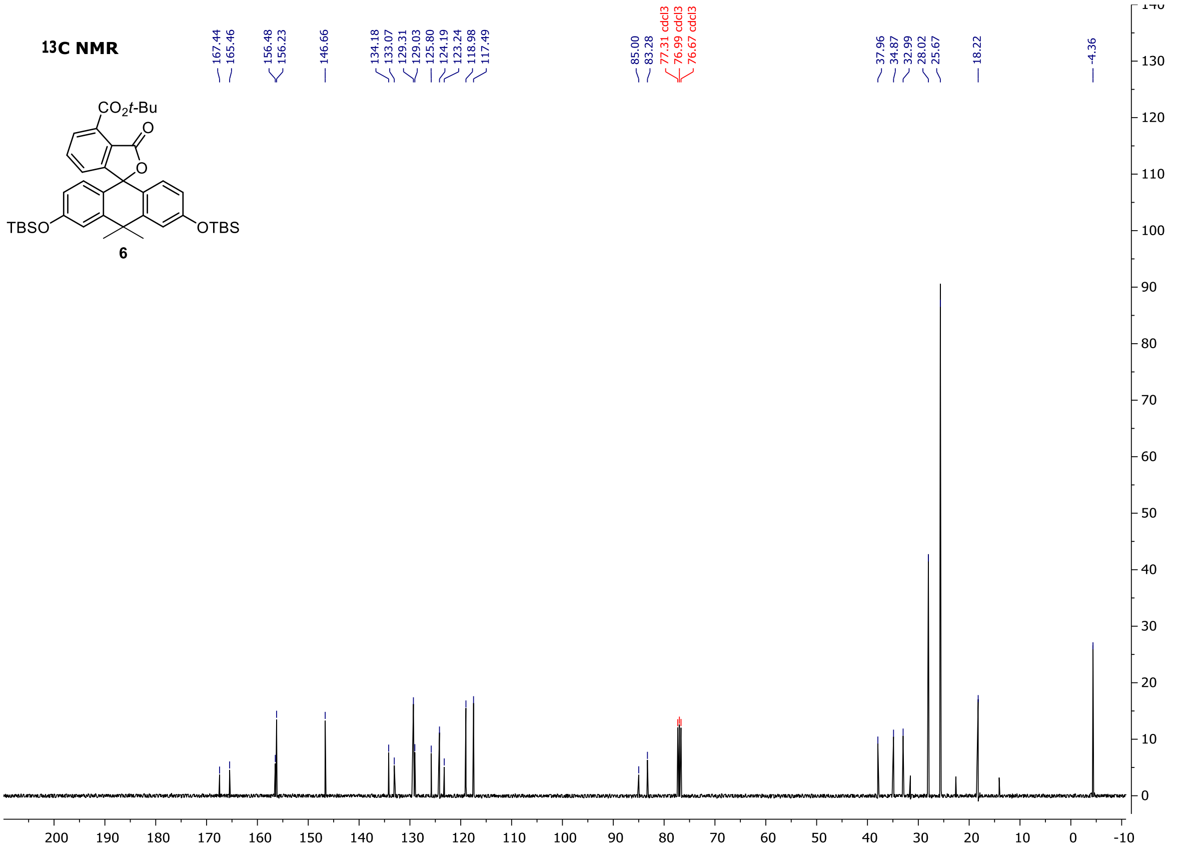

**Tert-butyl 3,7-bis((tert-butyldimethylsilyl)oxy)-5,5-dimethyl-3'-oxo-3'H,5H-spiro[dibenzo[b,e]siline-10,1'-isobenzofuran]-4'-carboxylate (7)**

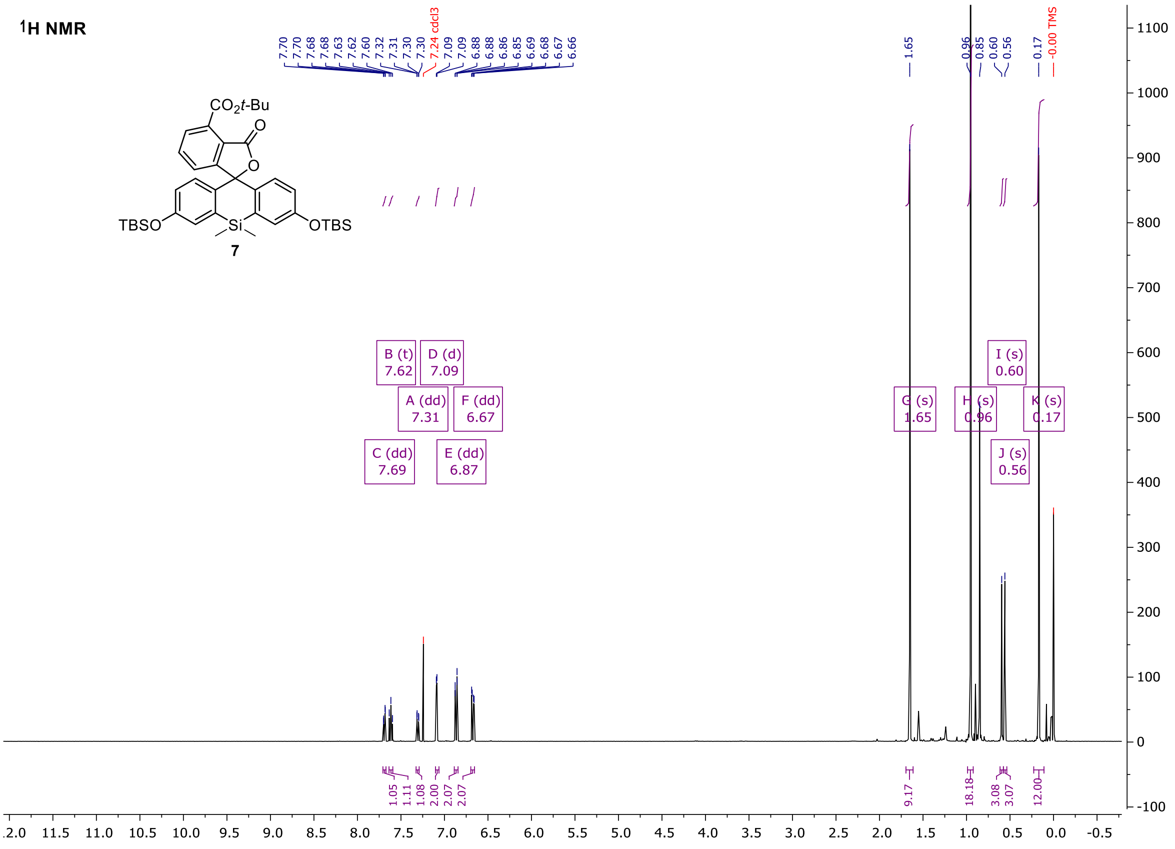

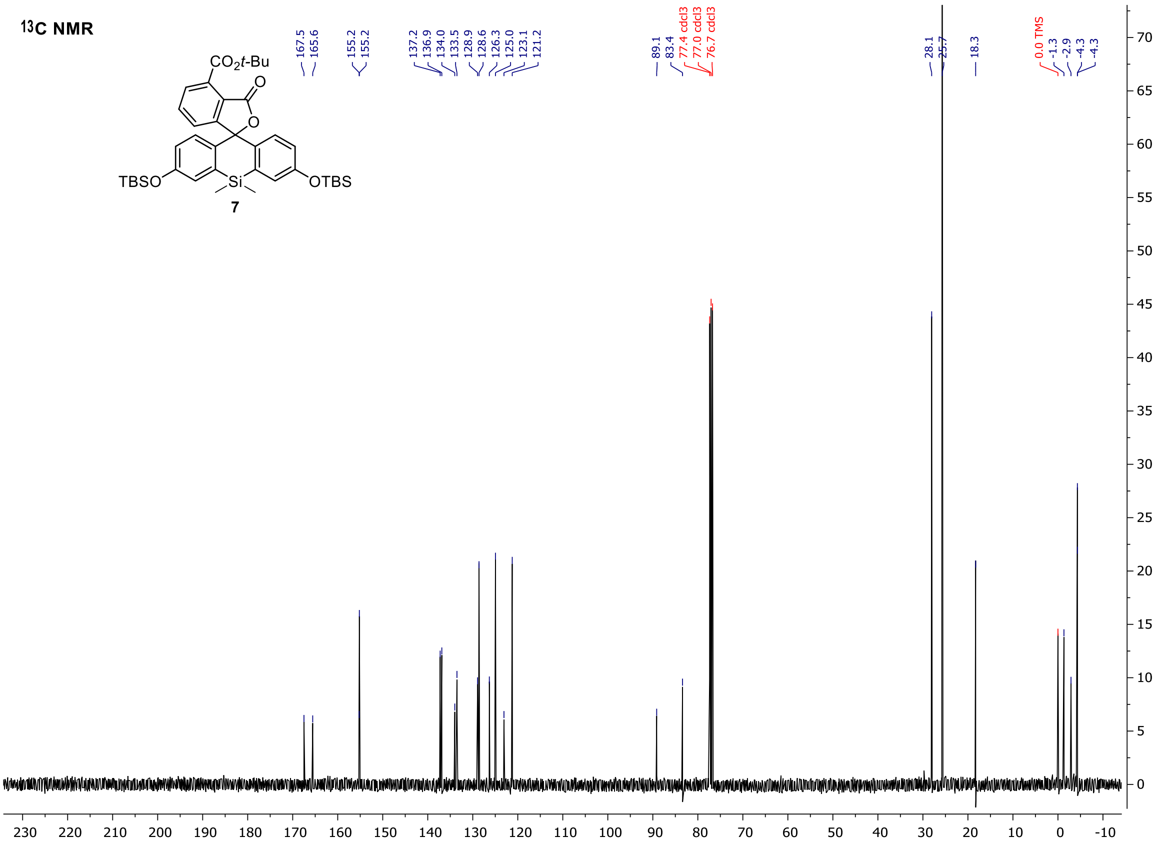

#### Tert-butyl 3',6'-dihydroxy-3-oxo-3H-spiro[isobenzofuran-1,9'-xanthene]-4-carboxylate (8)

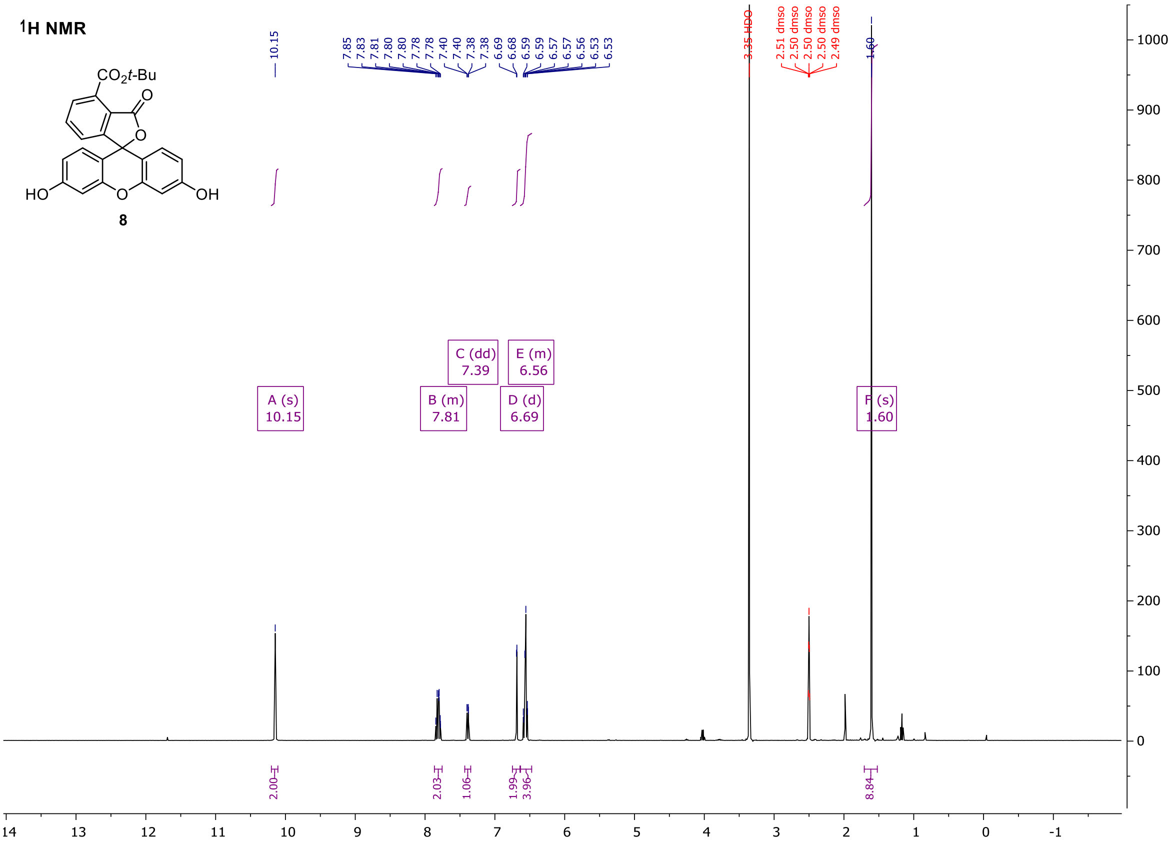

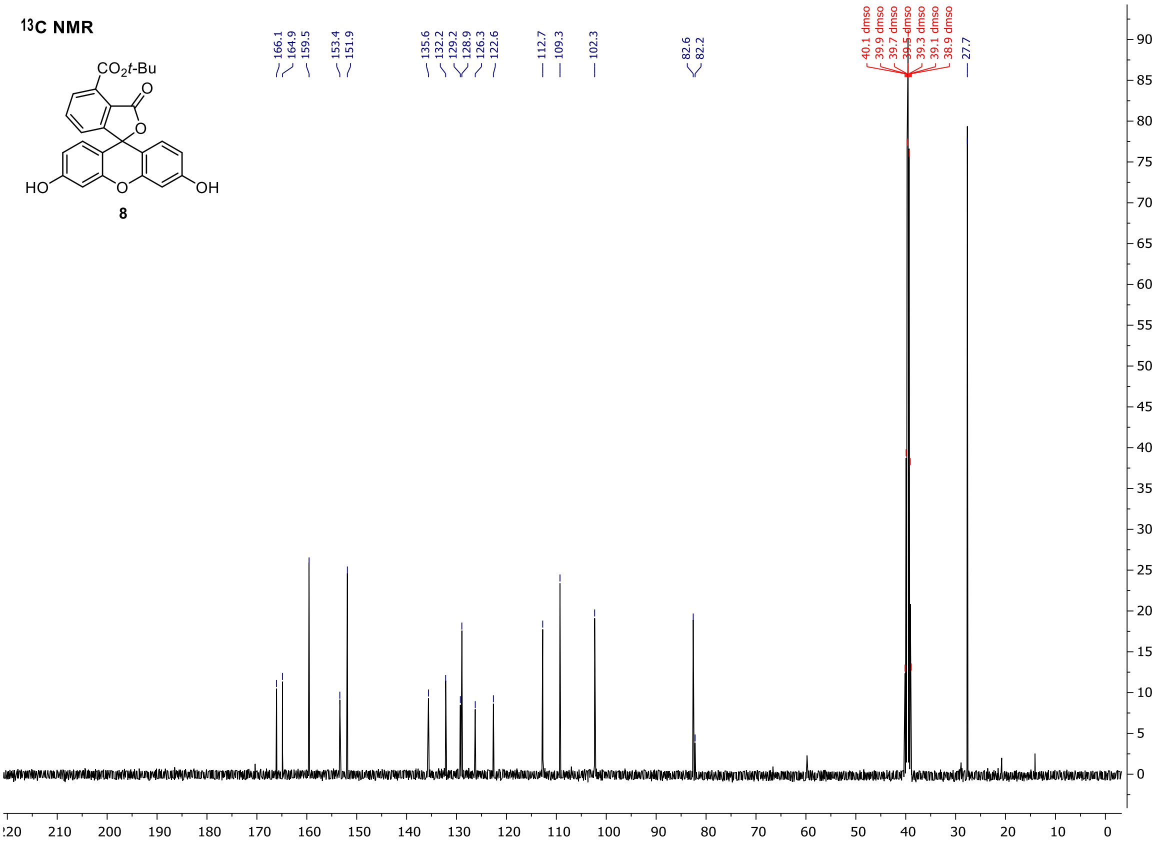

Tert-butyl 3,6-dihydroxy-10,10-dimethyl-3'-oxo-3'H,10H-spiro[anthracene-9,1'-isobenzofuran]-4'-carboxylate (9)

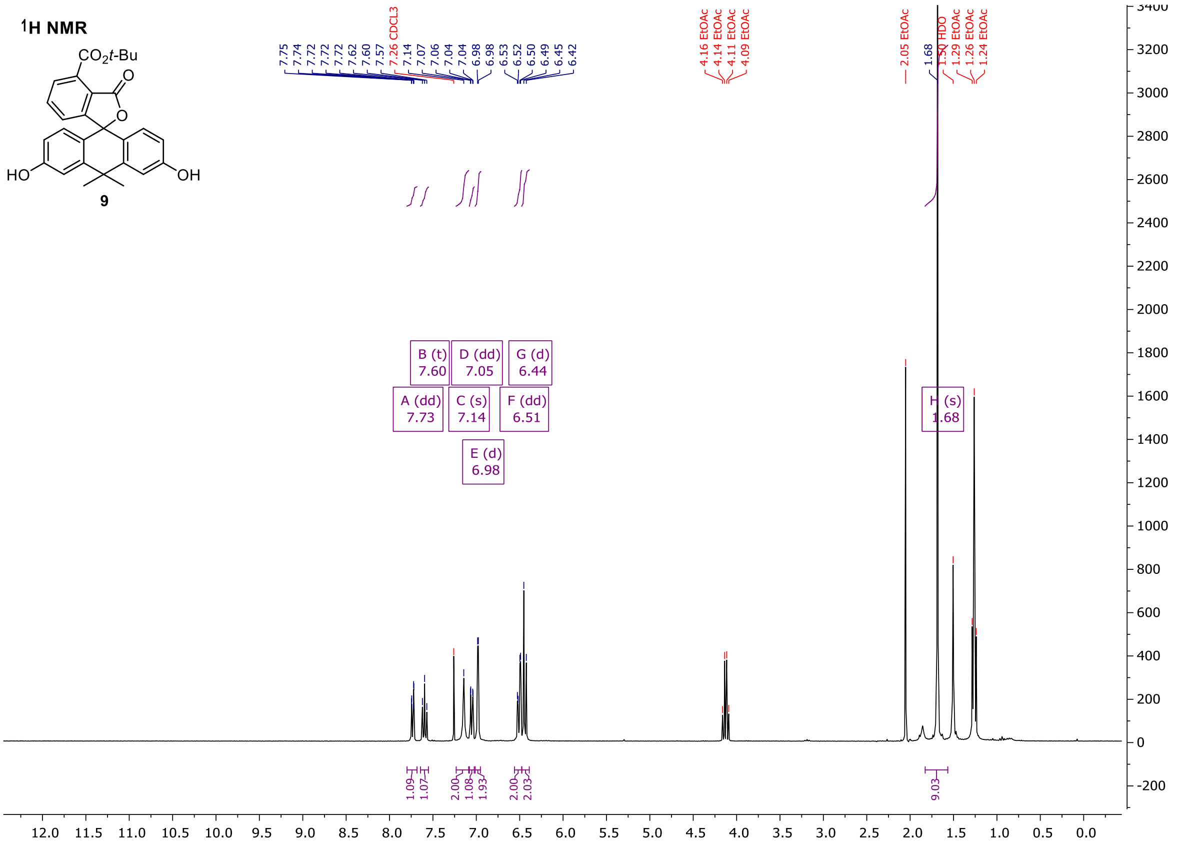

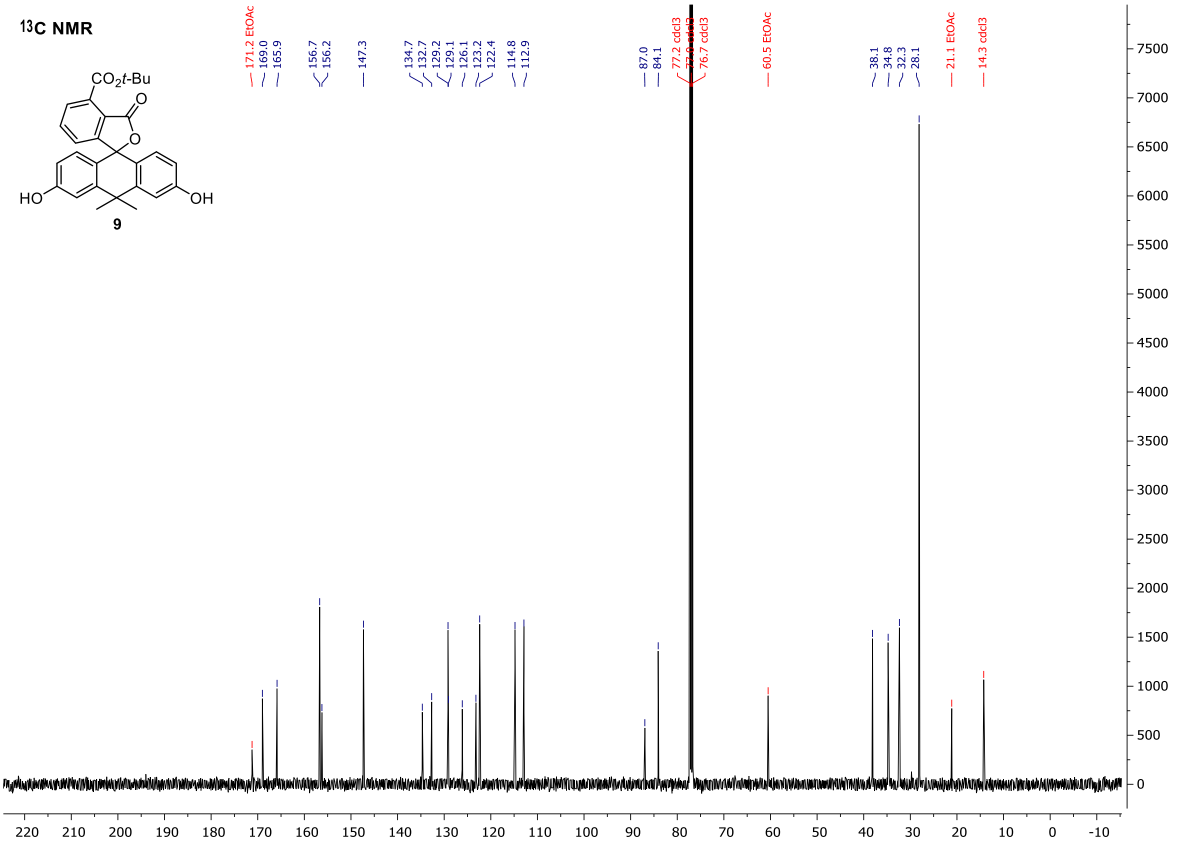

**Tert-butyl 3,7-dihydroxy-5,5-dimethyl-3'-oxo-3'H,5H-spiro[dibenzo[b,e]siline-10,1'-isobenzofuran]-4'-carboxylate (10)**

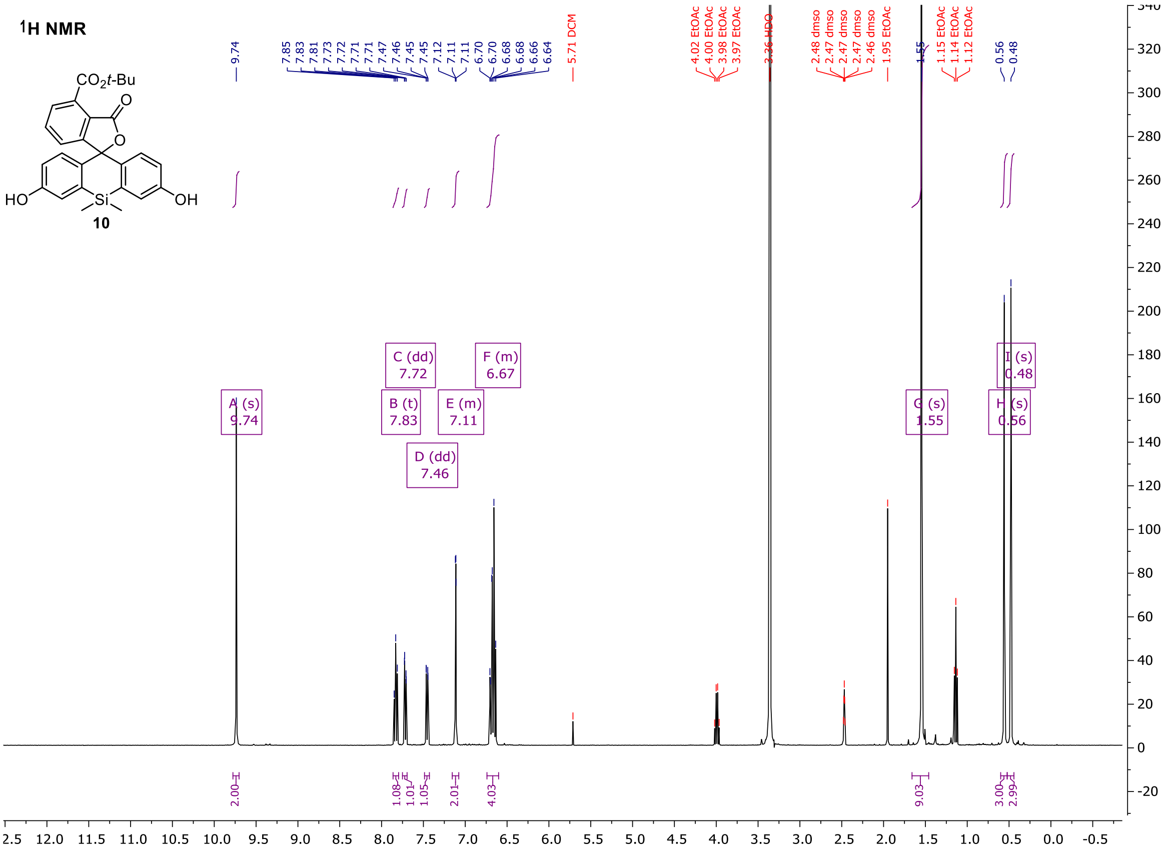

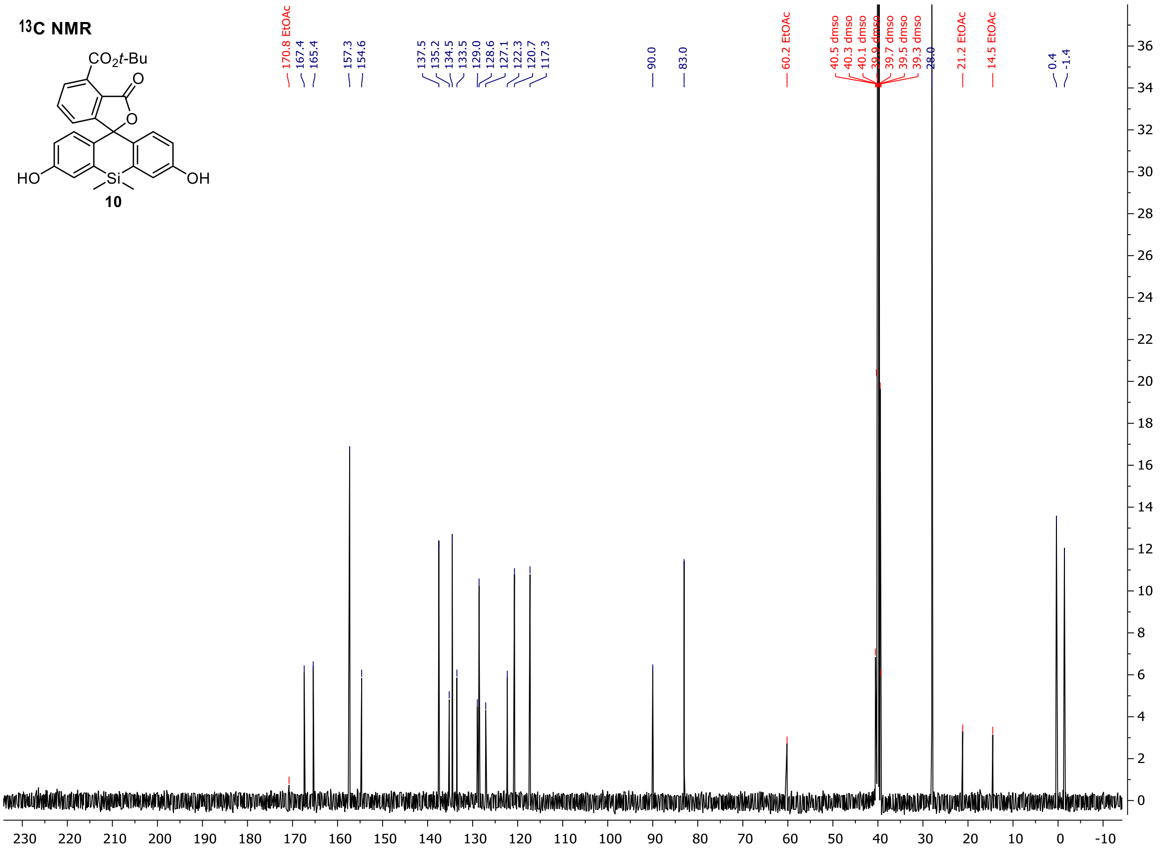

**Tert-butyl 3-oxo-3',6'-bis(((trifluoromethyl)sulfonyl)oxy)-3H-spiro[isobenzofuran-1,9'-xanthene]-4-carboxylate (11)**

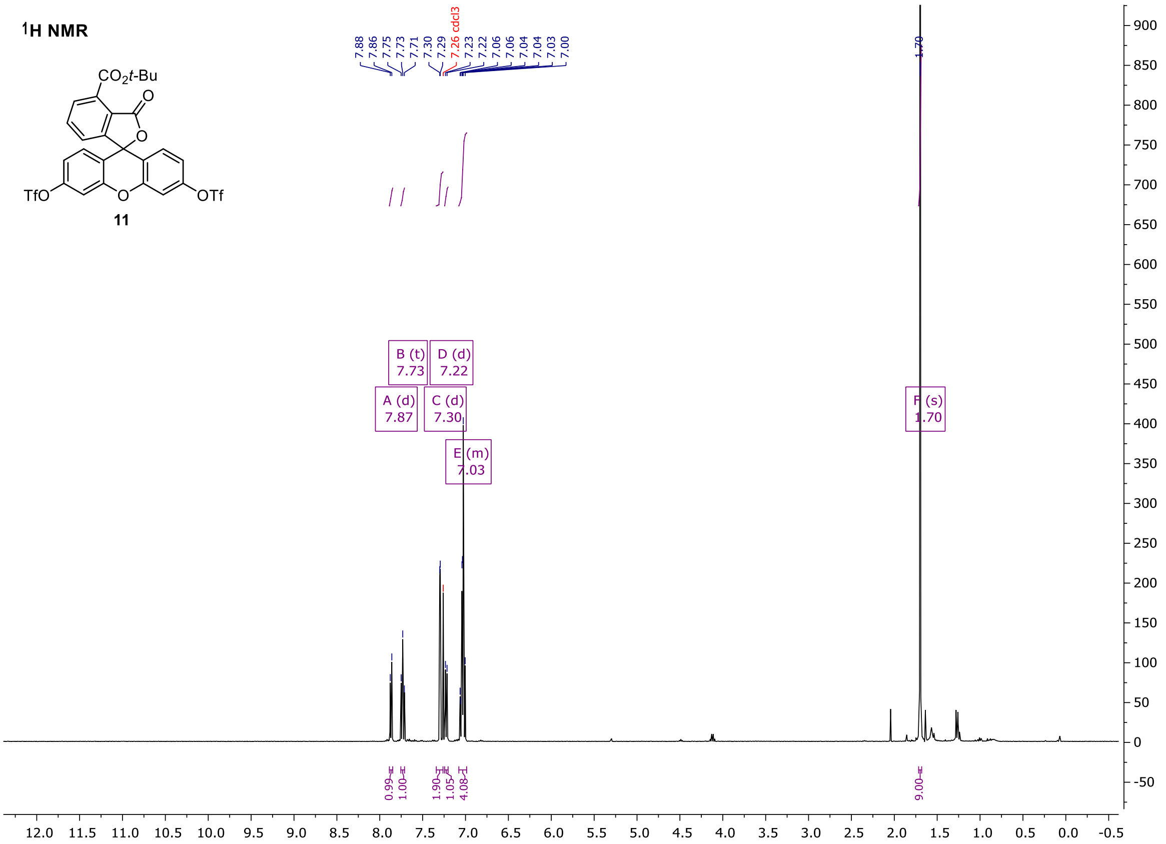

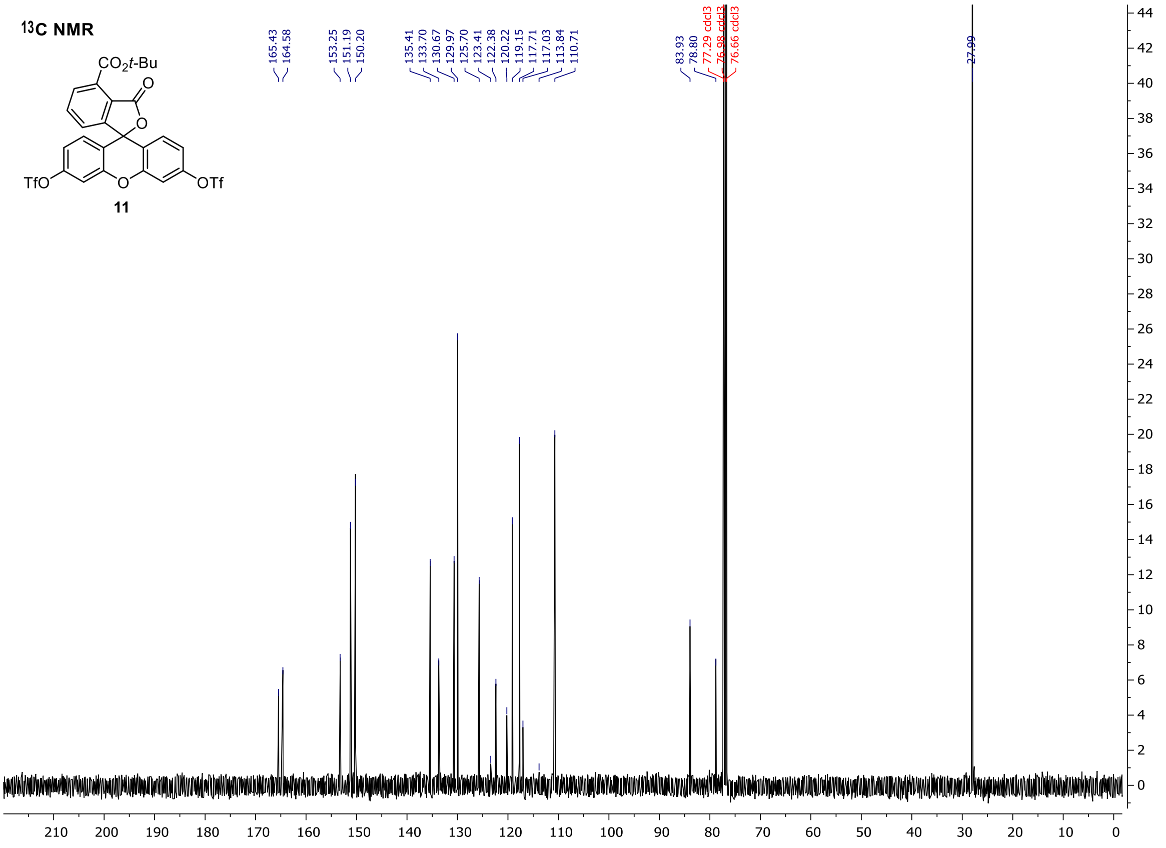

**Tert-butyl 10,10-dimethyl-3'-oxo-3,6-bis(((trifluoromethyl)sulfonyl)oxy)-3'H,10H-spiro[anthracene-9,1'-isobenzofuran]-4'-carboxylate (12)**

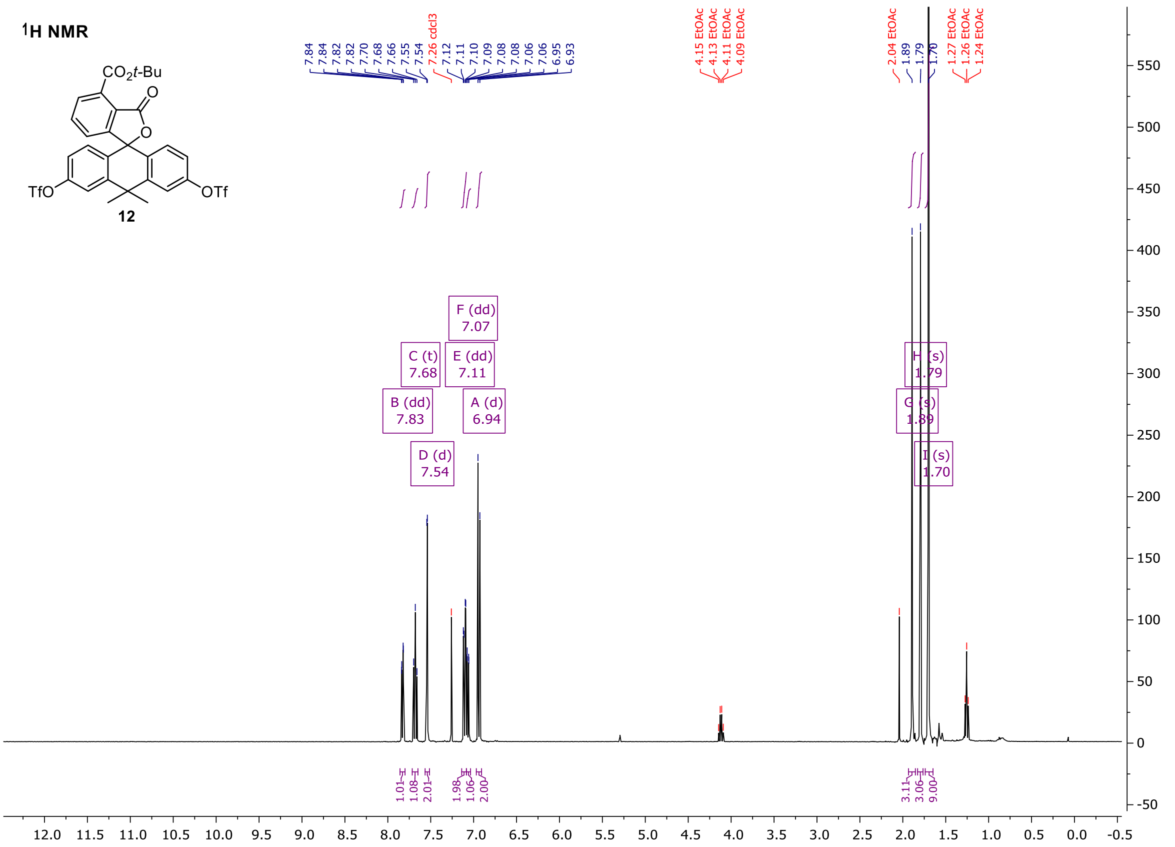

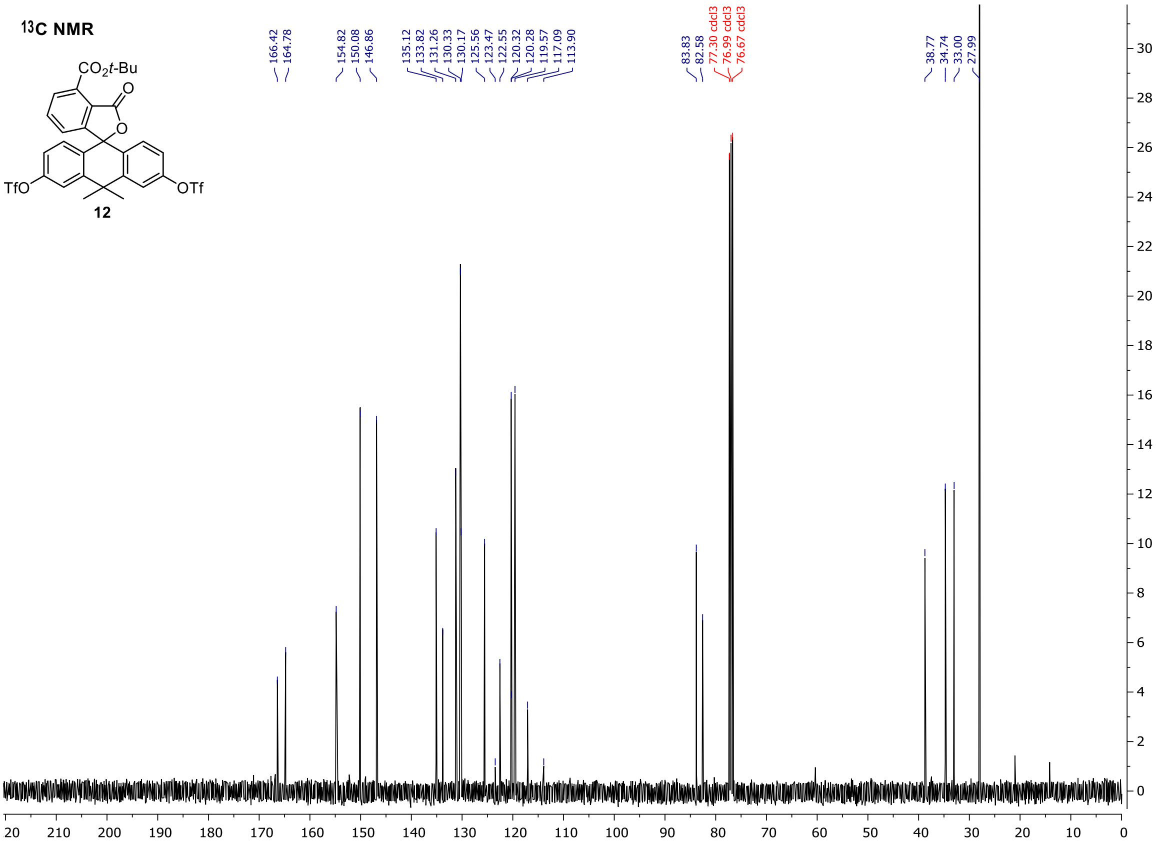

**Tert-butyl 5,5-dimethyl-3'-oxo-3,7-bis(((trifluoromethyl)sulfonyl)oxy)-3'H,5H-spiro[dibenzo[b,e]siline-10,1'-isobenzofuran]-4'-carboxylate (13)**

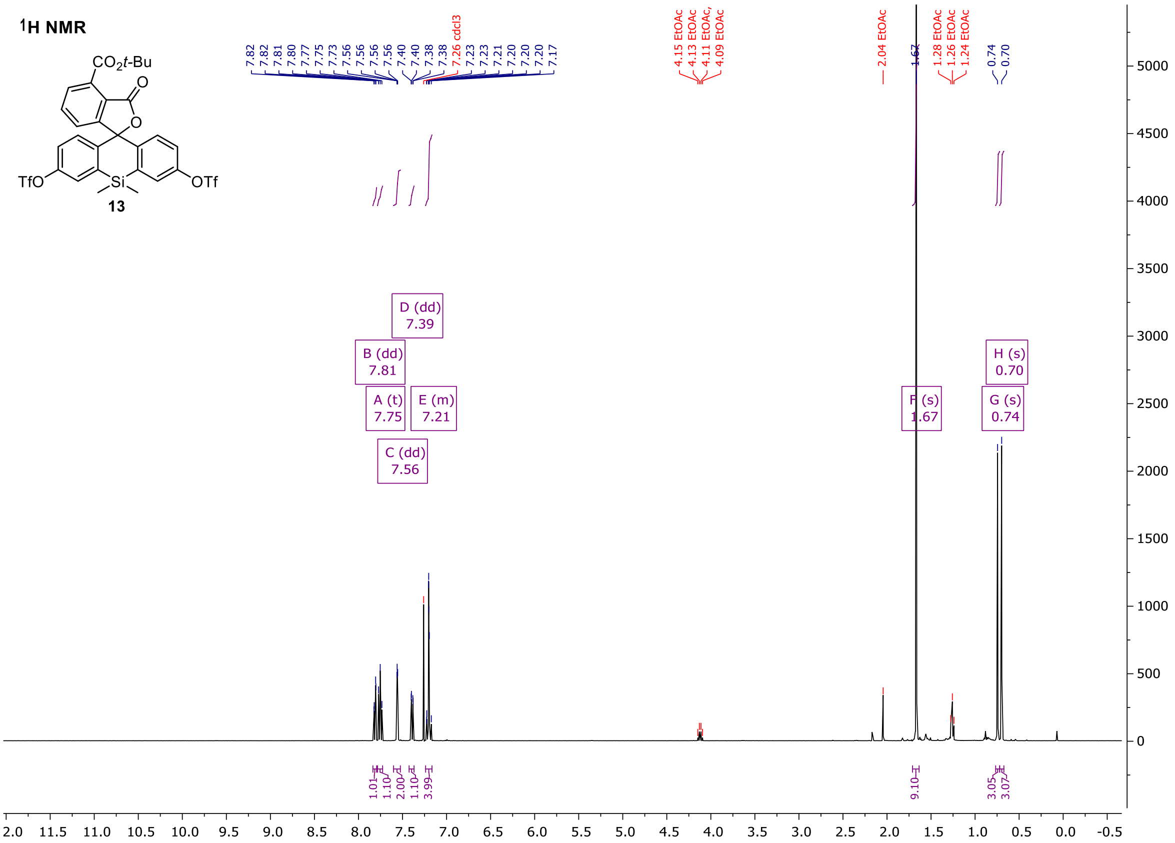

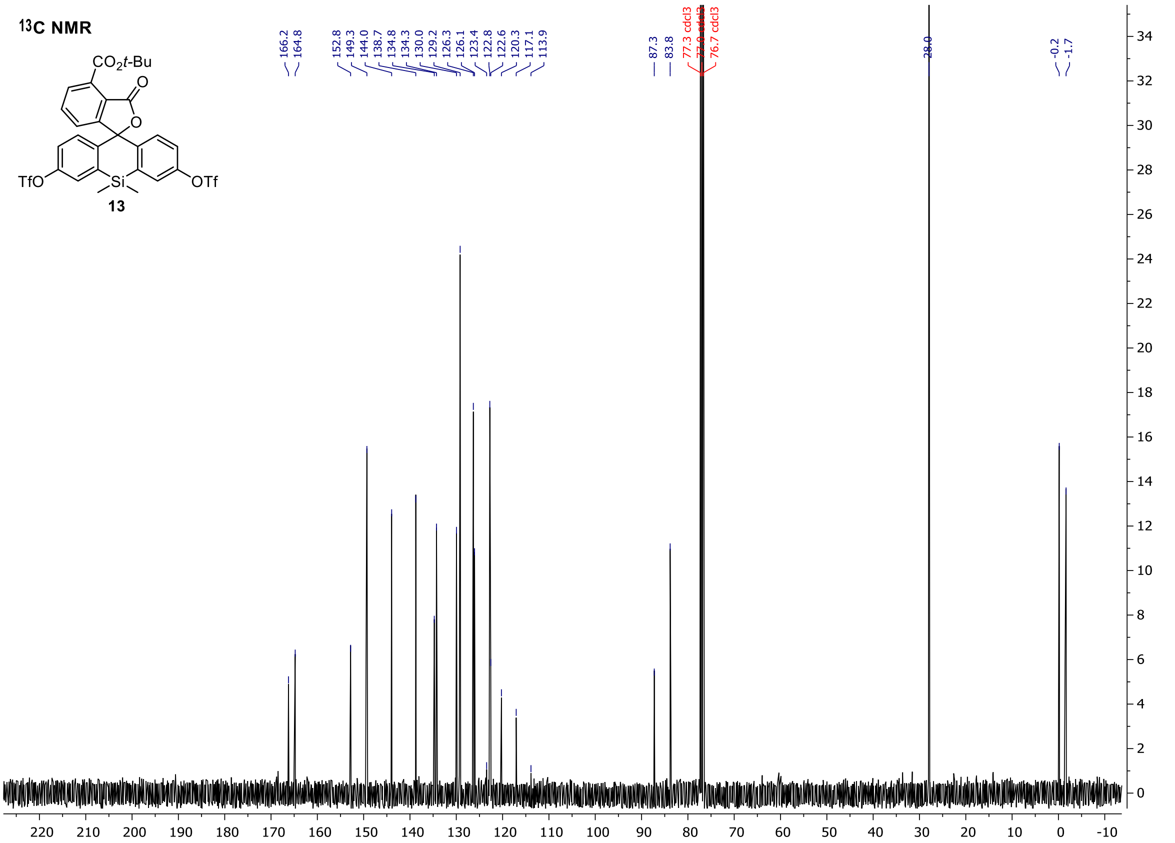

**Tert-butyl 3',6'-bis(dimethylamino)-3-oxo-3H-spiro[isobenzofuran-1,9'-xanthene]-4-carboxylate (14)**

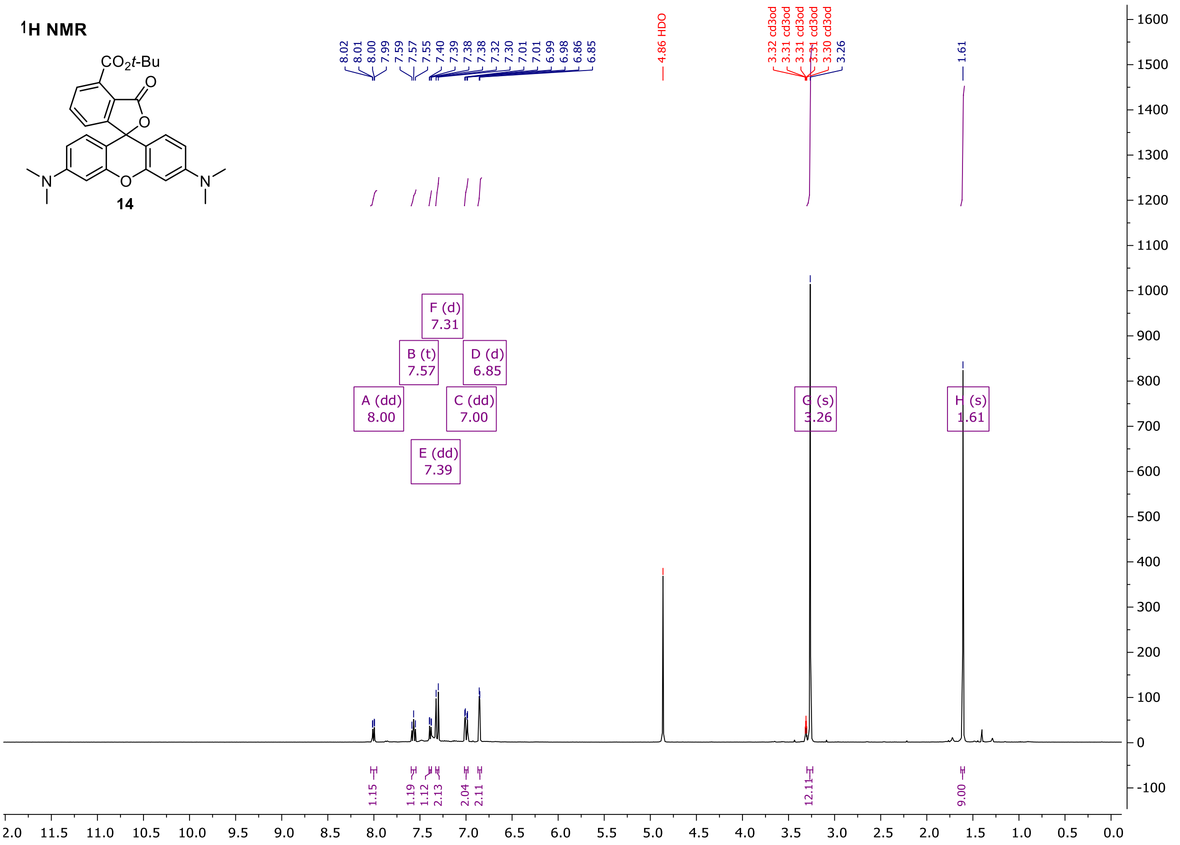

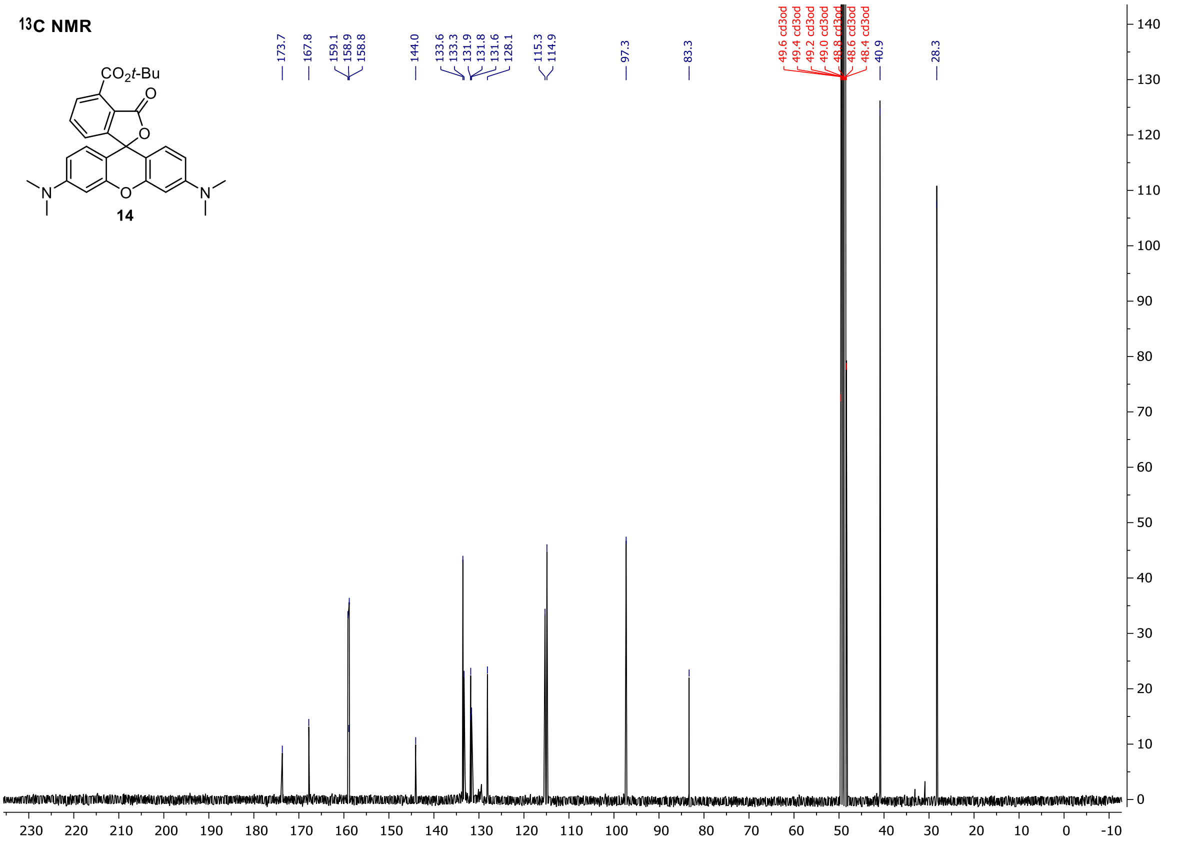

**Tert-butyl 10,10-dimethyl-3,6-bis(methylamino)-3'-oxo-3'H,10H-spiro[anthracene-9,1'-isobenzofuran]-4'-carboxylate (15)**

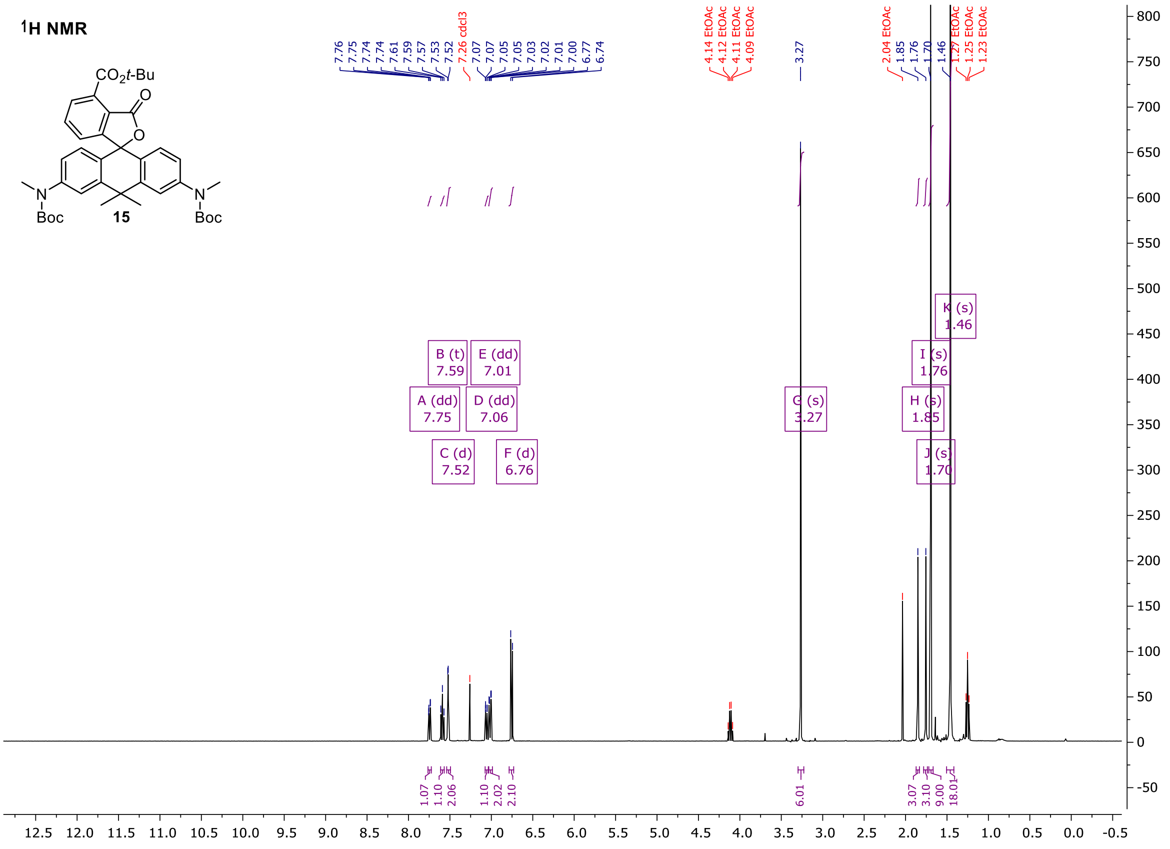

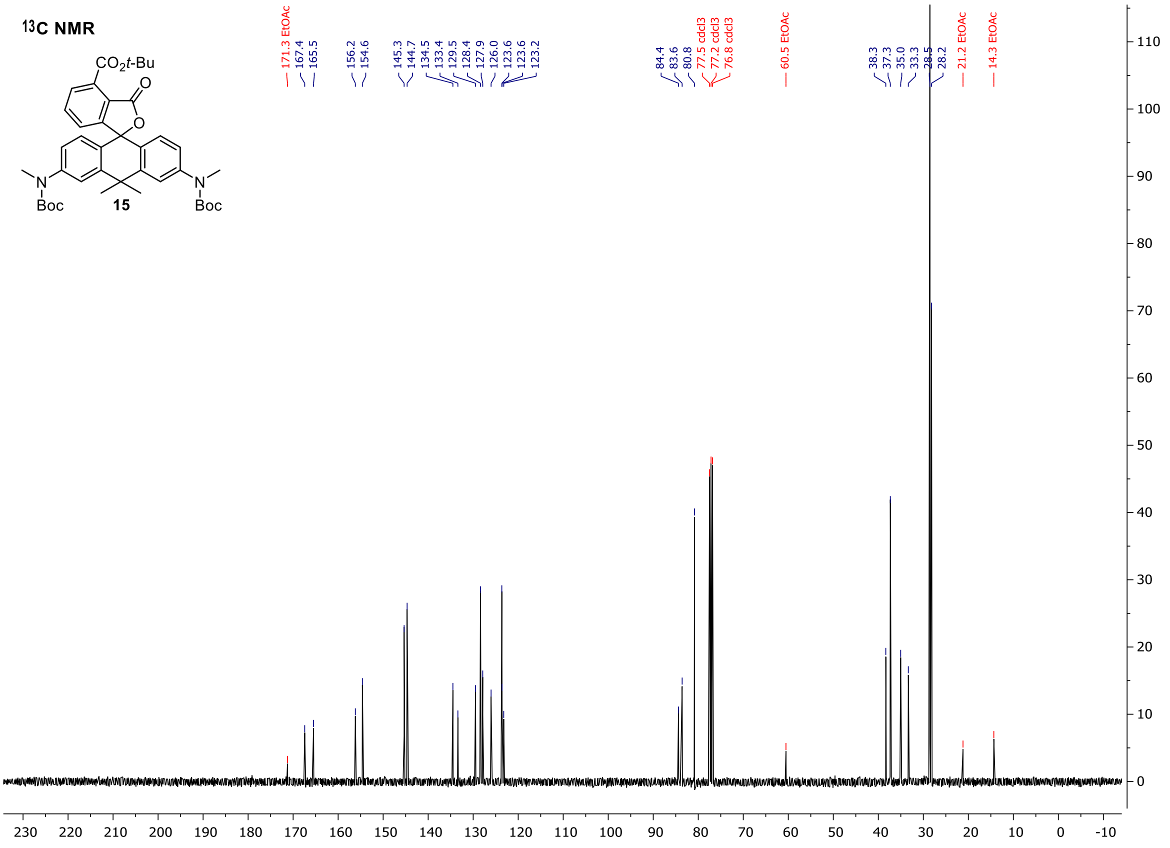

**Tert-butyl 3,6-bis(dimethylamino)-10,10-dimethyl-3'-oxo-3'H,10H-spiro[anthracene-9,1'-isobenzofuran]-4'-carboxylate (16)**

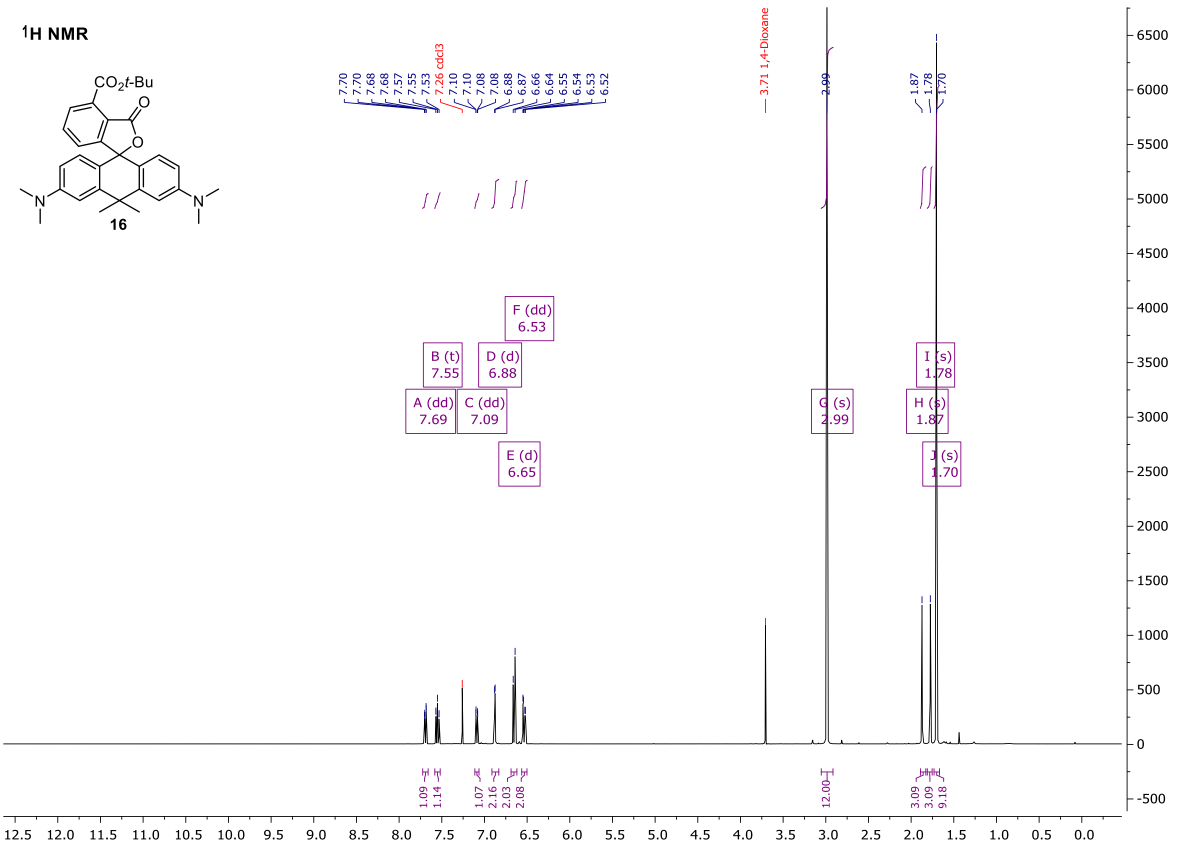

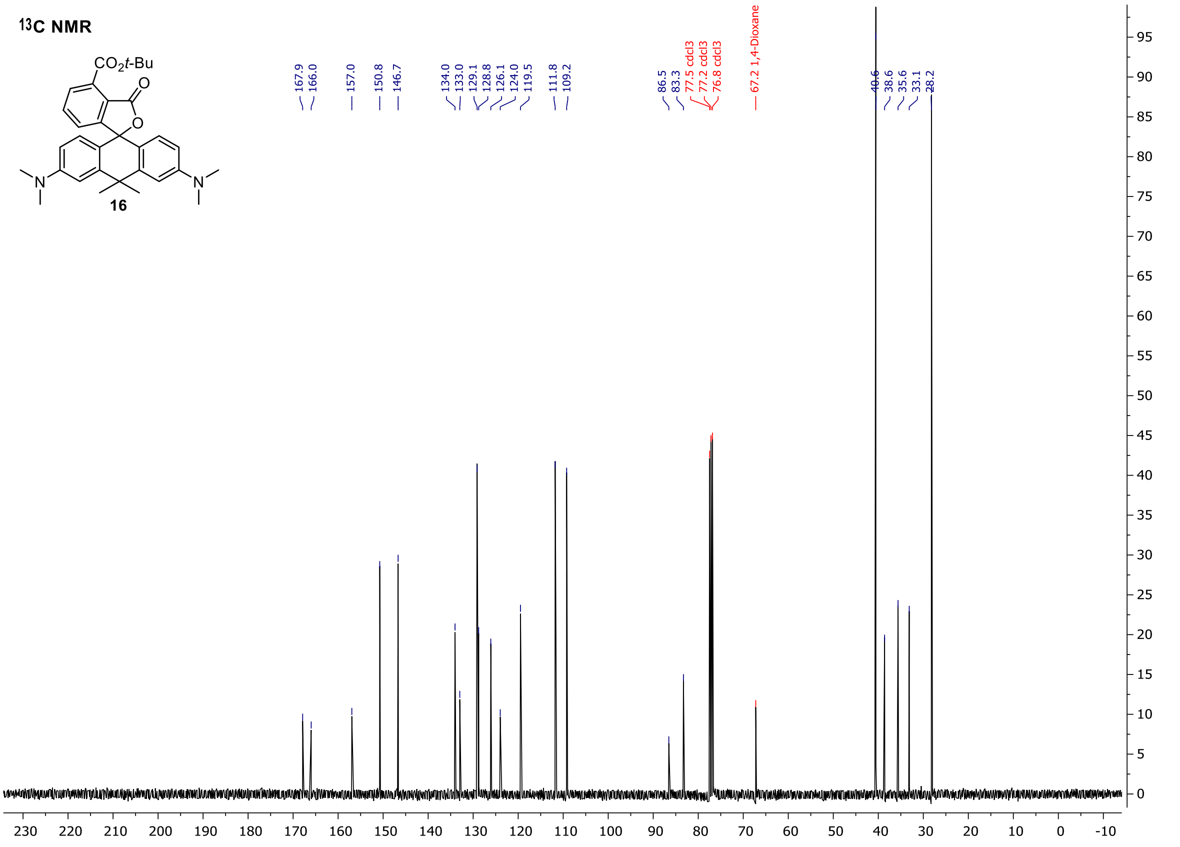

**Tert-butyl 3,7-bis(dimethylamino)-5,5-dimethyl-3'-oxo-3'H,5H-spiro[dibenzo[b,e]siline-10,1'-isobenzofuran]-4'-carboxylate (17)**

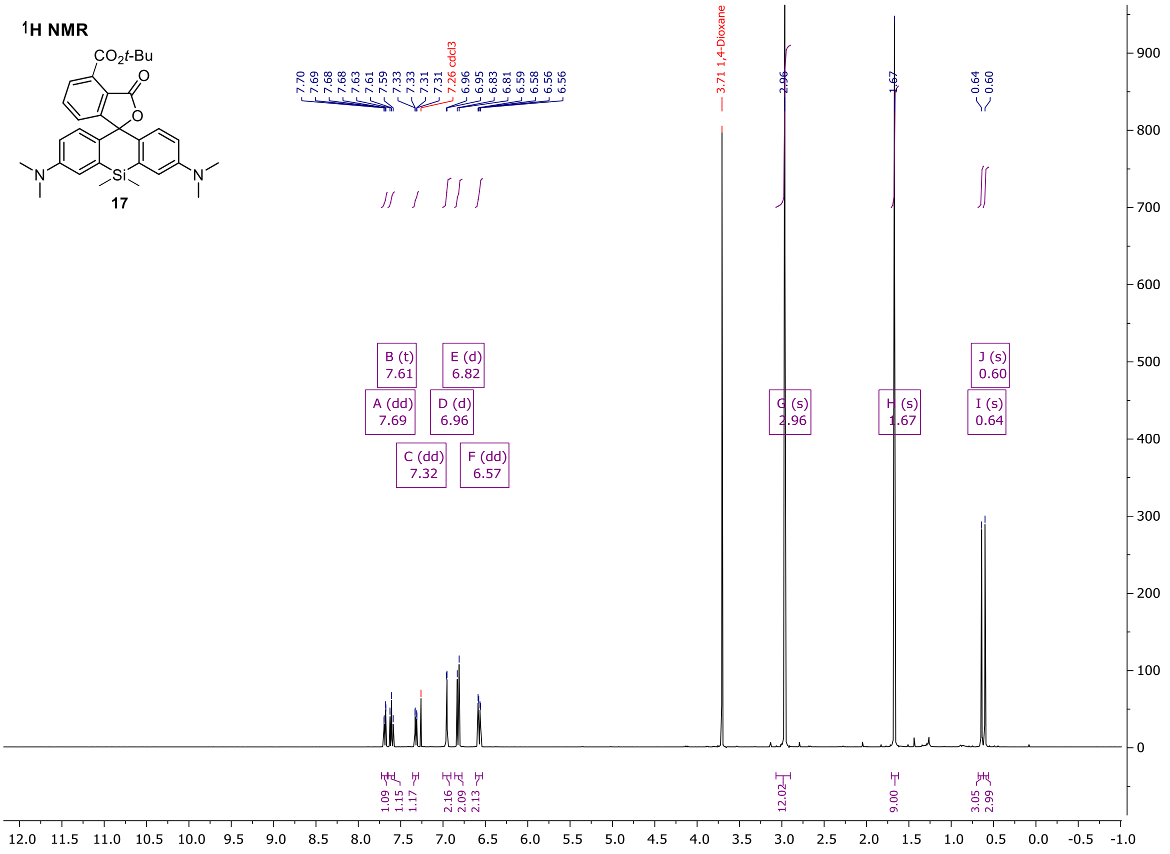

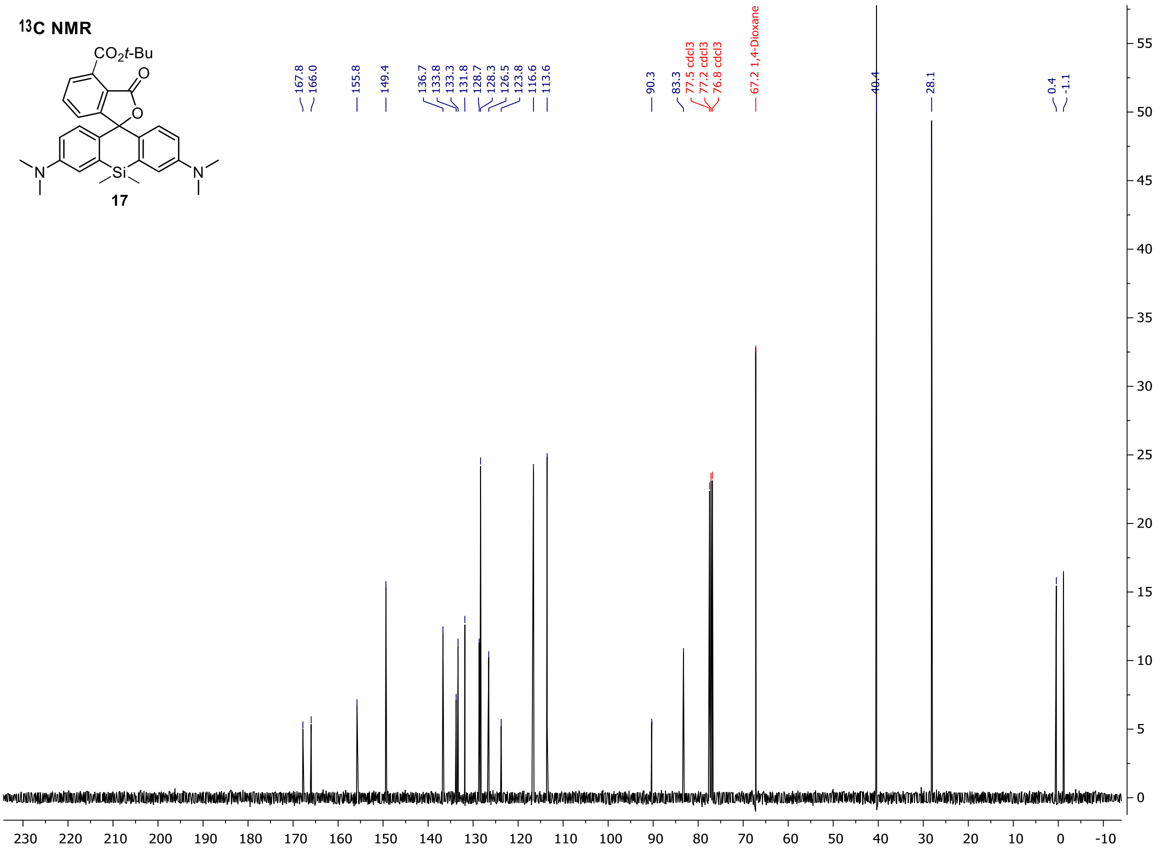

#### 4-TMR-COOH (18)

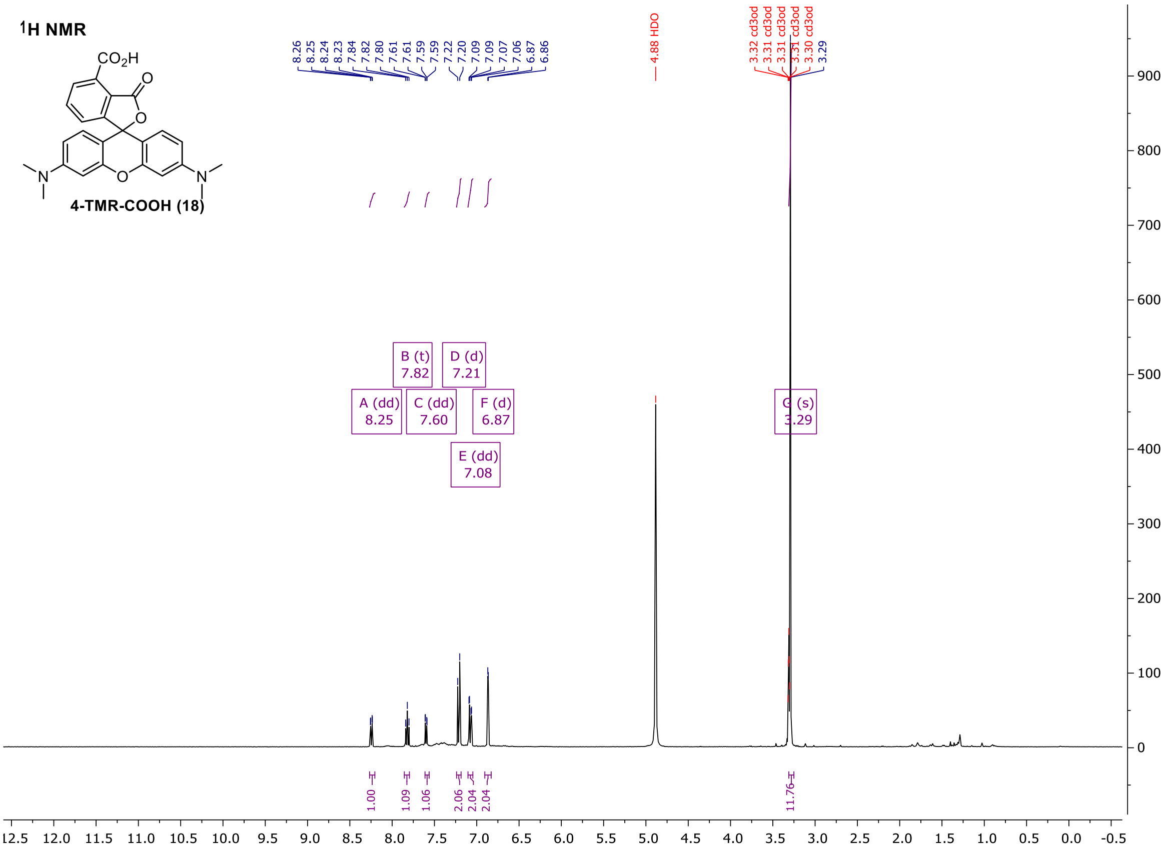
1H NMR
4-TMR-COOH (18)
8.26 8.25 8.24 8.23 7.84 7.82 7.80 7.61 7.61 7.59 7.59 7.22 7.20 7.09 7.09 7.07 7.06 6.87 6.86
4.88 HDO
3.32 cd3od 3.31 cd3od 3.31 cd3od 3.31 cd3od 3.30 cd3od 3.29
A (dd) 8.25
B (t) 7.82
C (dd) 7.60
D (d) 7.21
E (dd) 7.08
F (d) 6.87
G (s) 3.29
1.00 1.09 1.06 2.06 2.04 2.04 11.76

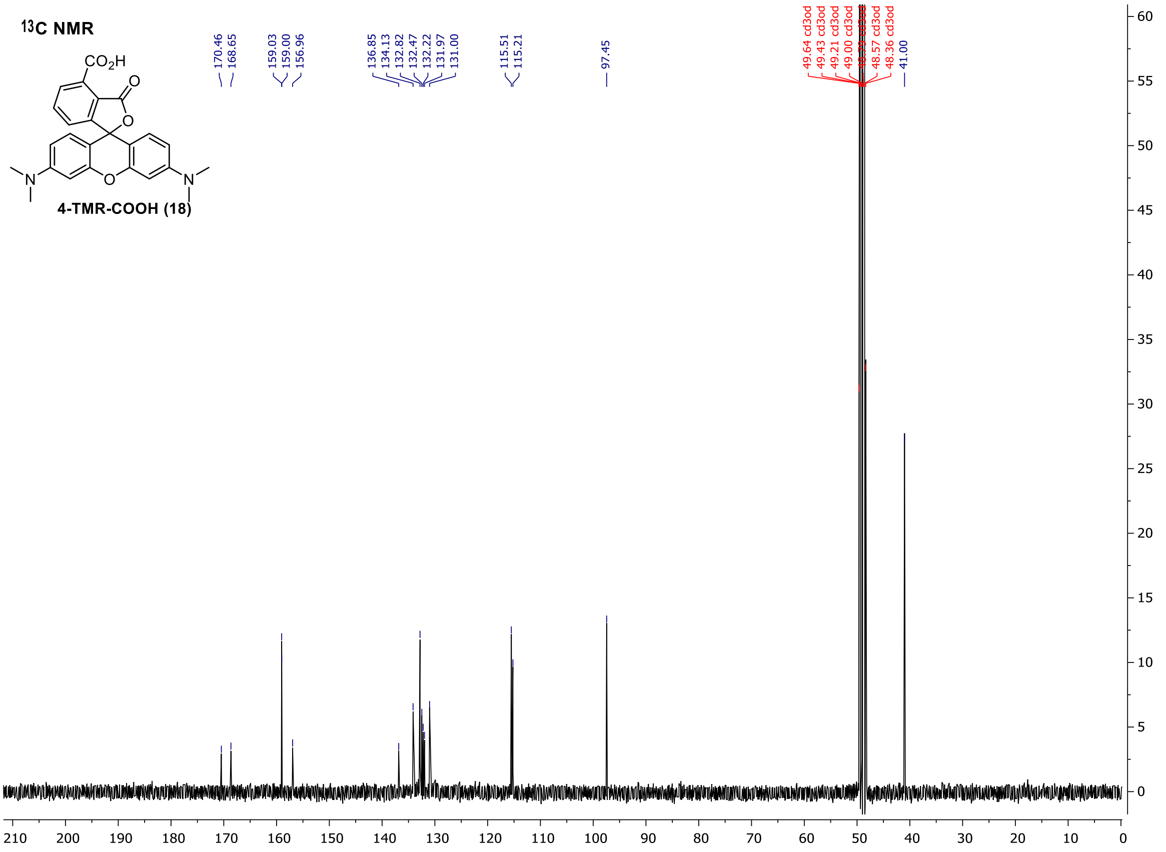
13C NMR
4-TMR-COOH (18)
170.46 168.65 159.03 159.00 156.96 136.85 134.13 132.82 132.47 132.22 131.97 131.00 115.51 115.21 97.45
49.64 cd3od 49.43 cd3od 49.21 cd3od 49.00 cd3od 48.57 cd3od 48.36 cd3od 41.00

#### 4-580CP-COOH (19)

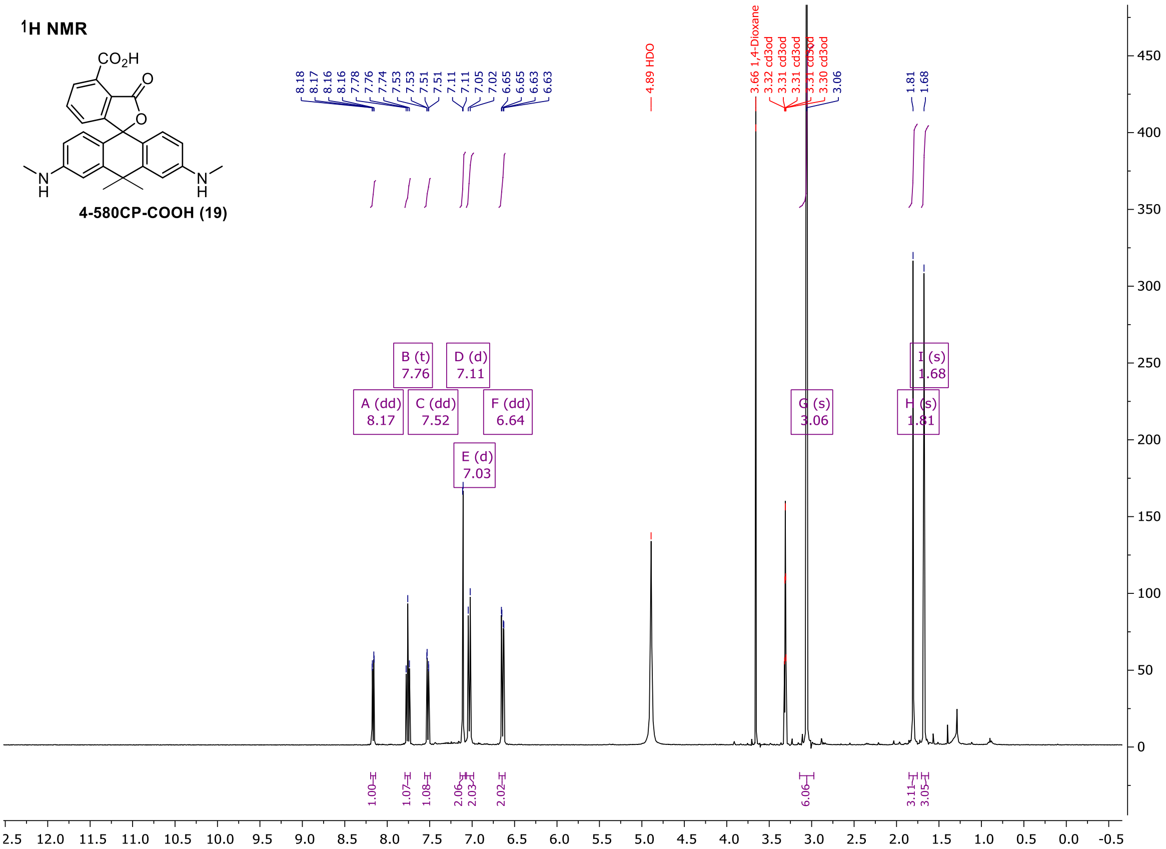

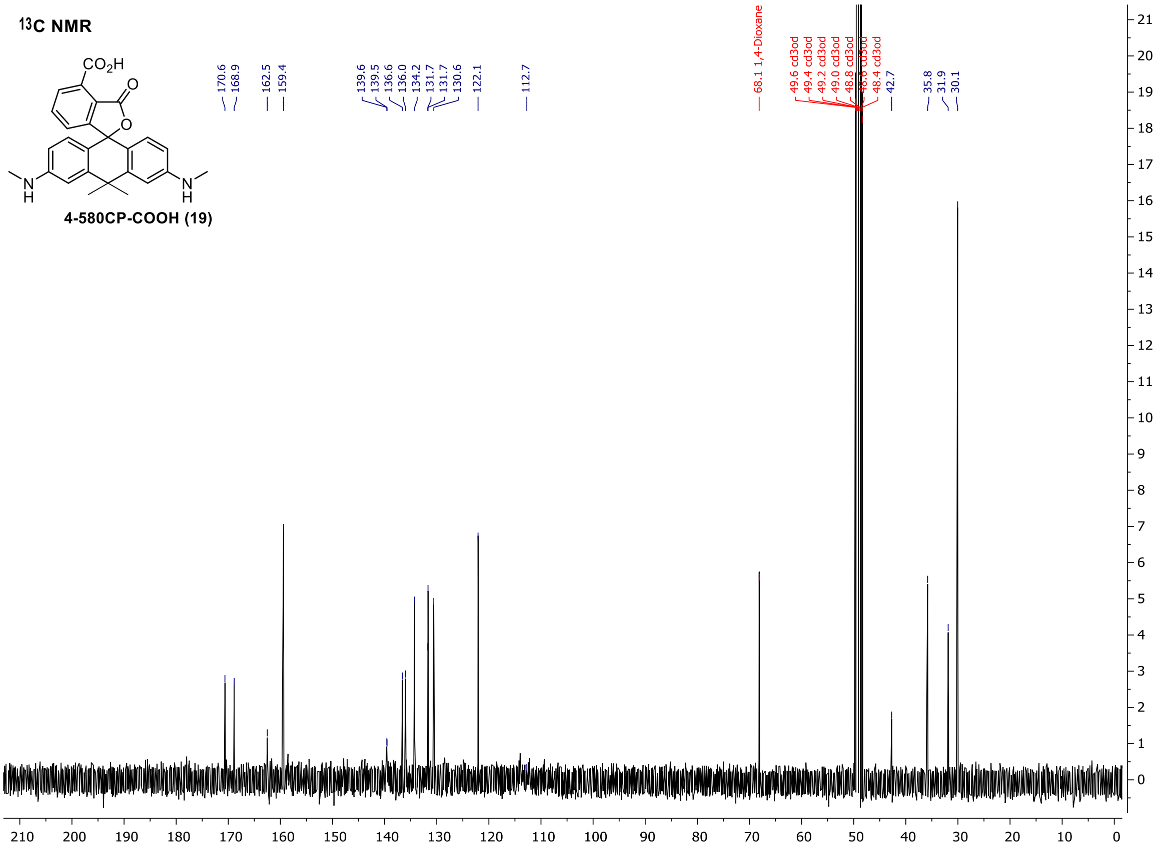

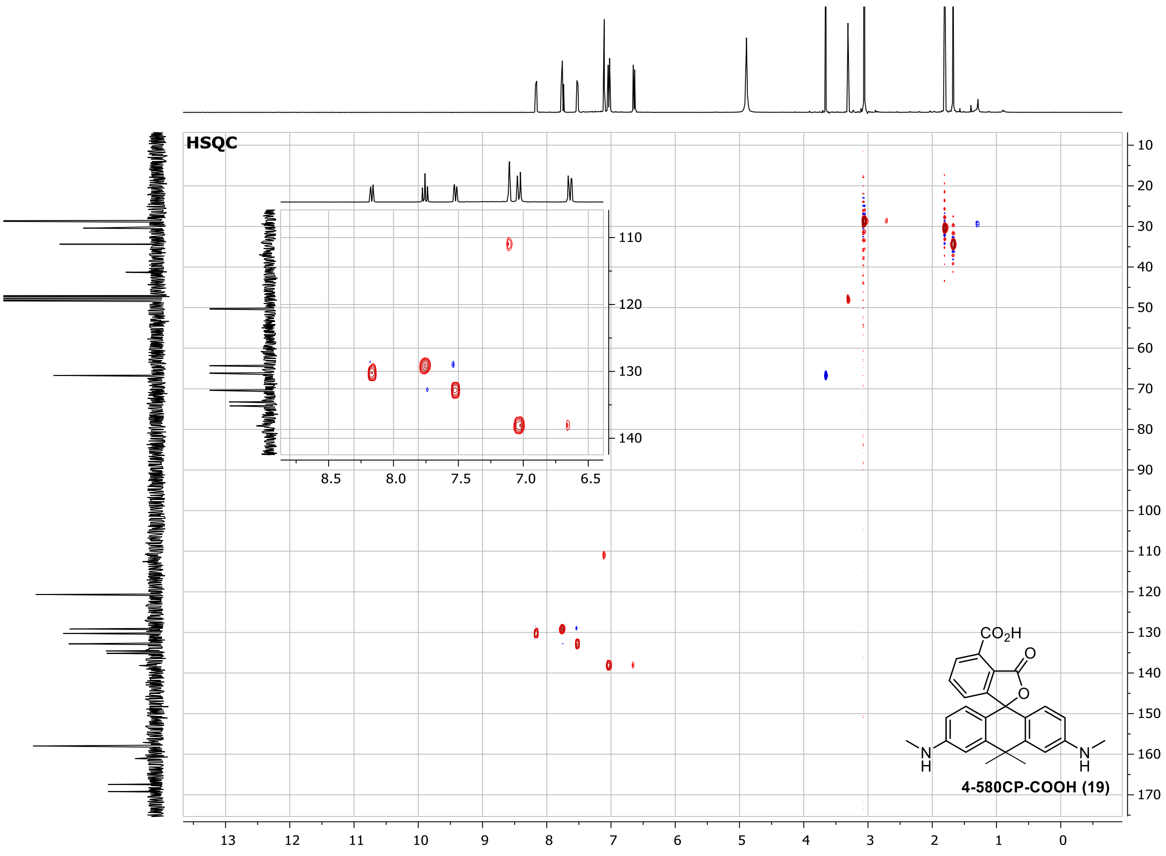
HSQC
4-580CP-COOH (19)

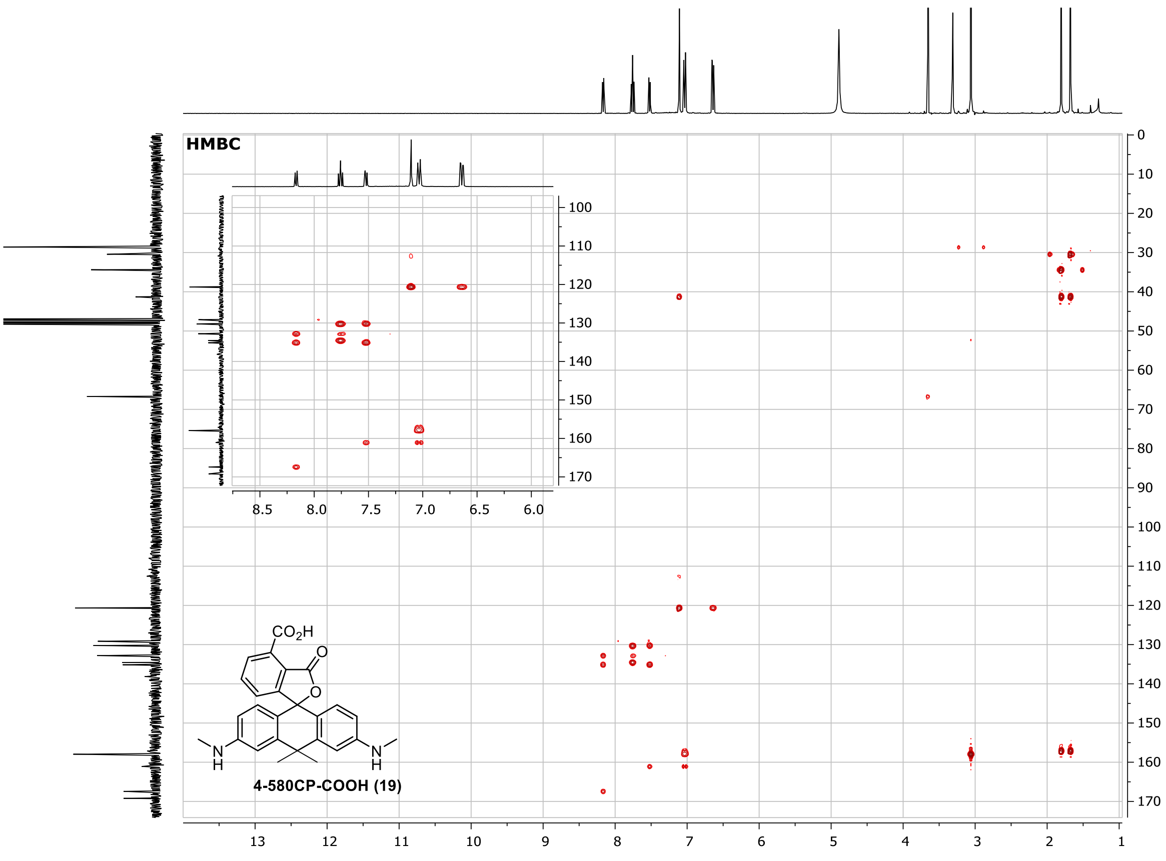
HMBC
4-580CP-COOH (19)

#### 4-610CP-COOH (20)

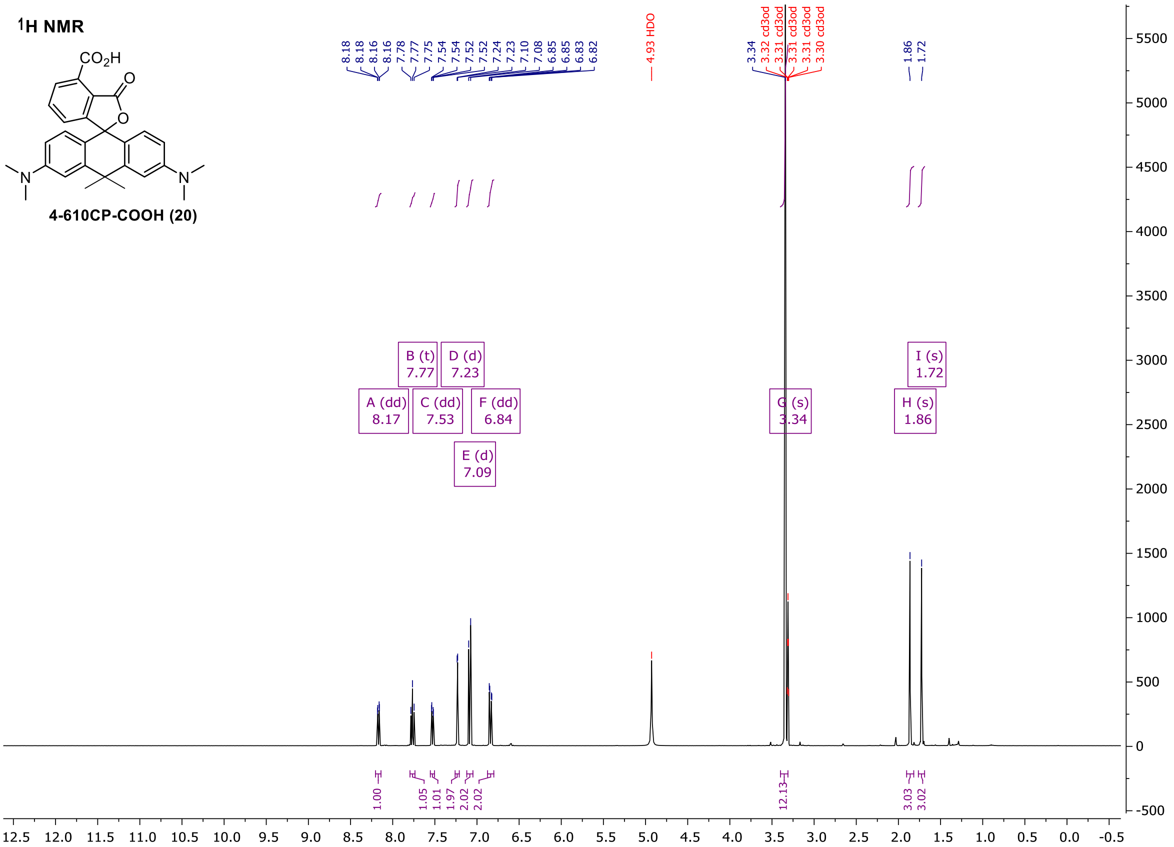

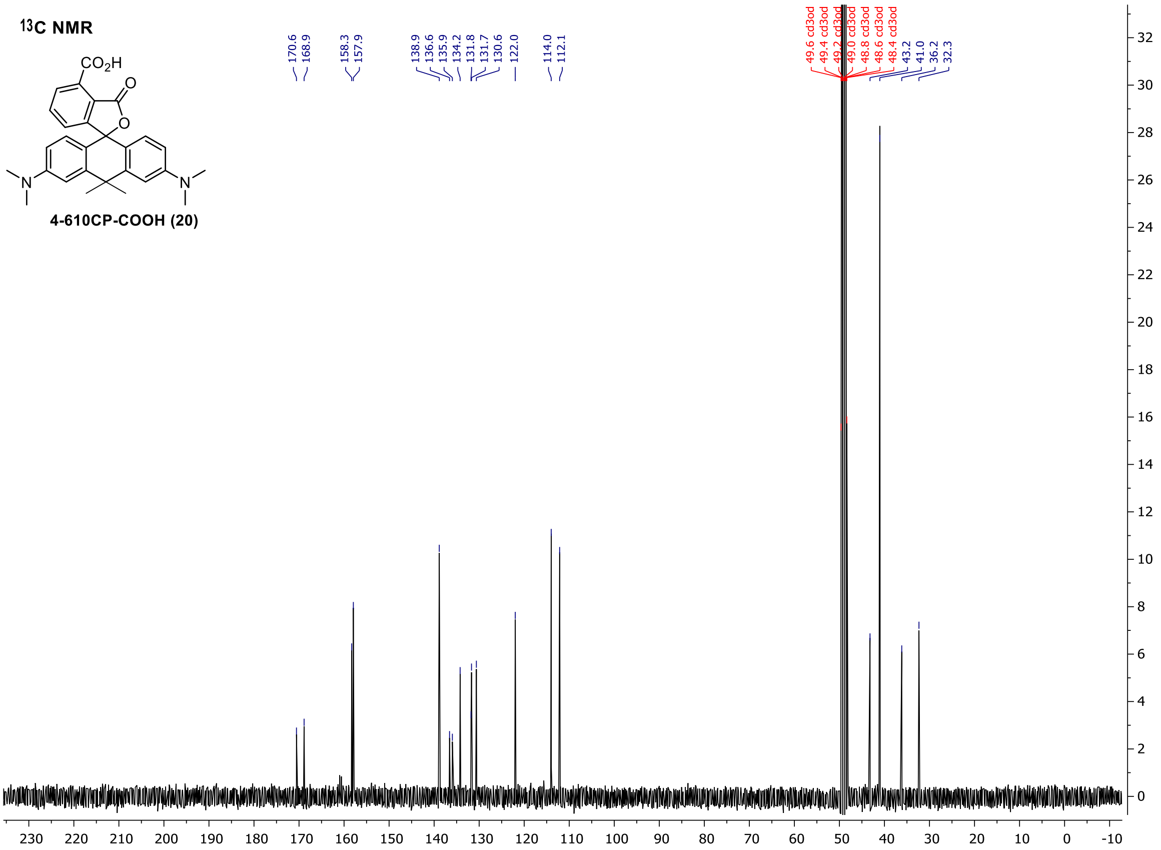

#### 4-SiR-COOH (21)

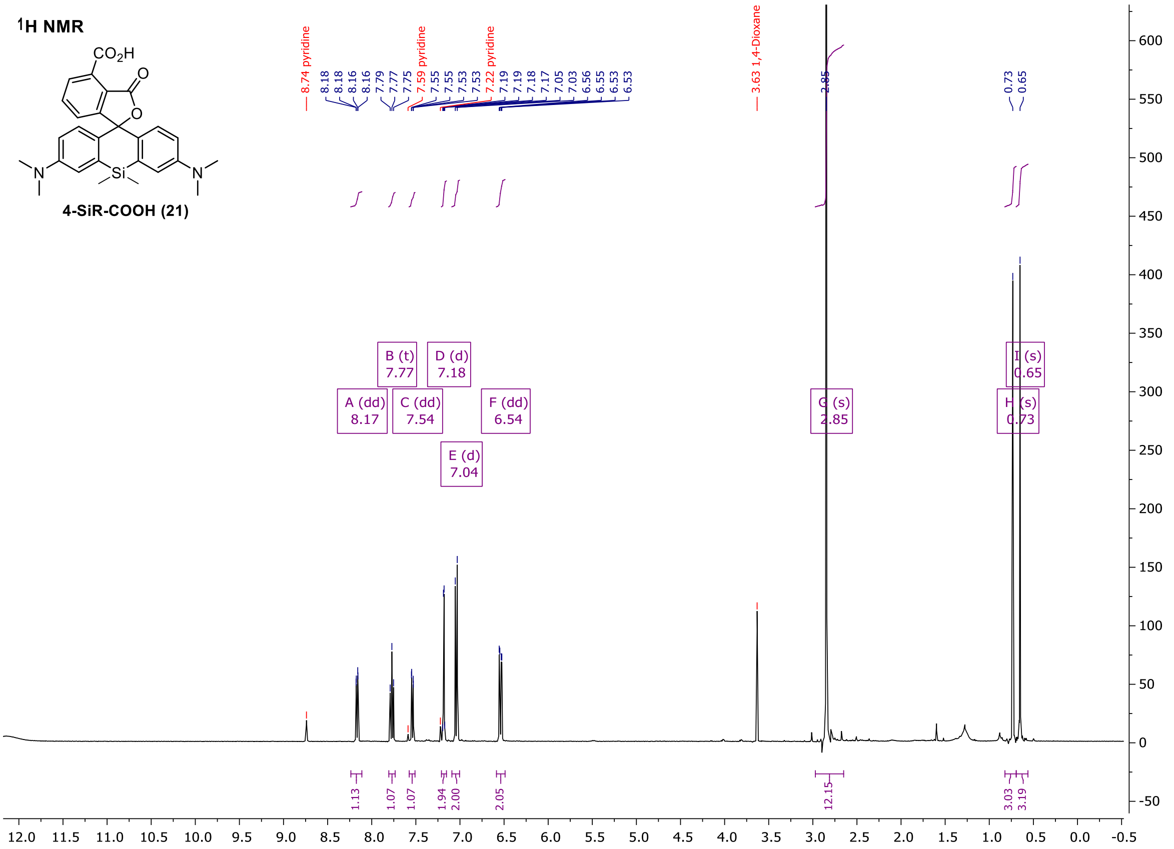
1H NMR
CO2H
4-SiR-COOH (21)
8.74 pyridine
8.18
8.18
8.16
8.16
7.79
7.77
7.75
7.59 pyridine
7.55
7.55
7.53
7.53
7.22 pyridine
7.19
7.19
7.18
7.17
7.05
7.03
6.56
6.55
6.53
6.53
3.63 1,4-Dioxane
2.85
0.73
0.65
A (dd) 8.17
B (t) 7.77
C (dd) 7.54
D (d) 7.18
E (d) 7.04
F (dd) 6.54
G (s) 2.85
H (s) 0.73
I (s) 0.65
1.13
1.07
1.07
1.94
2.00
2.05
12.15
3.03
3.19

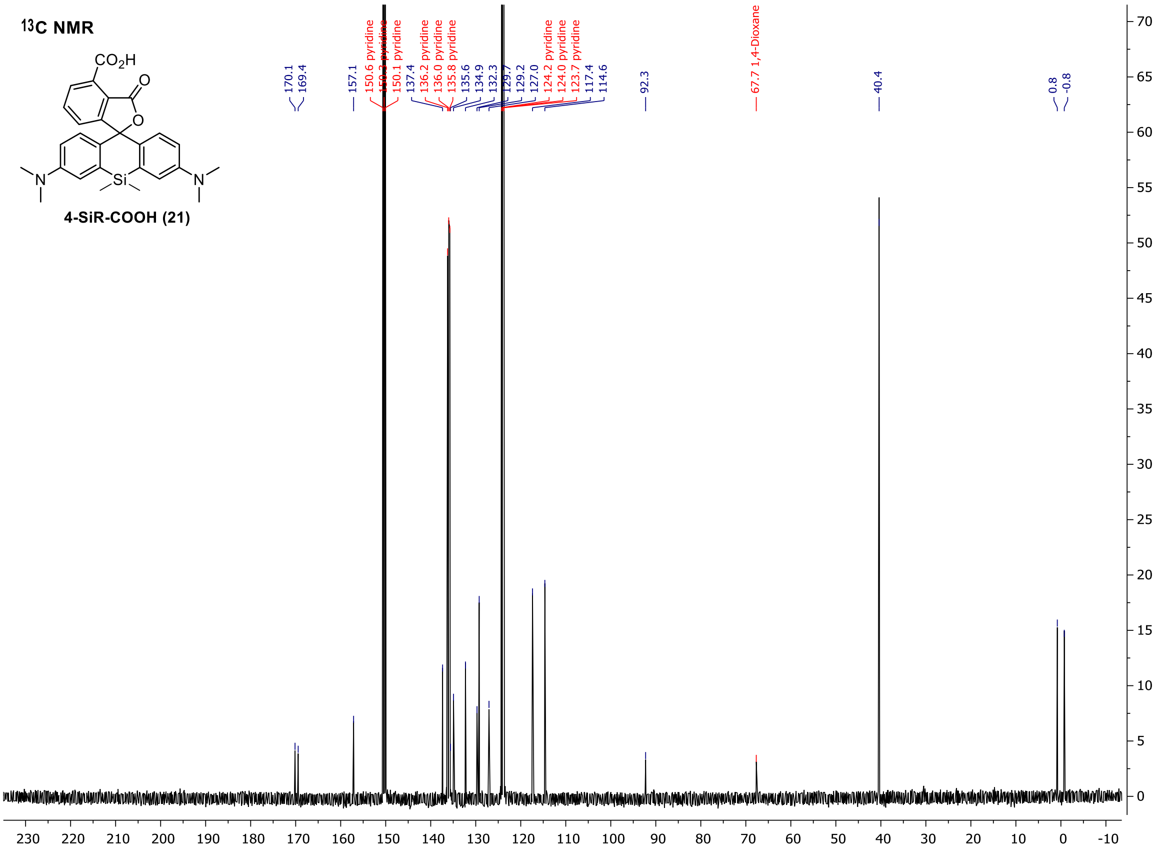
13C NMR
CO2H
4-SiR-COOH (21)
170.1
169.4
157.1
150.6 pyridine
150.3 pyridine
150.1 pyridine
137.4
136.2 pyridine
136.0 pyridine
135.8 pyridine
135.6
134.9
132.3
129.7
129.2
127.0
124.2 pyridine
124.0 pyridine
123.7 pyridine
117.4
114.6
92.3
67.7 1,4-Dioxane
40.4
0.8
-0.8

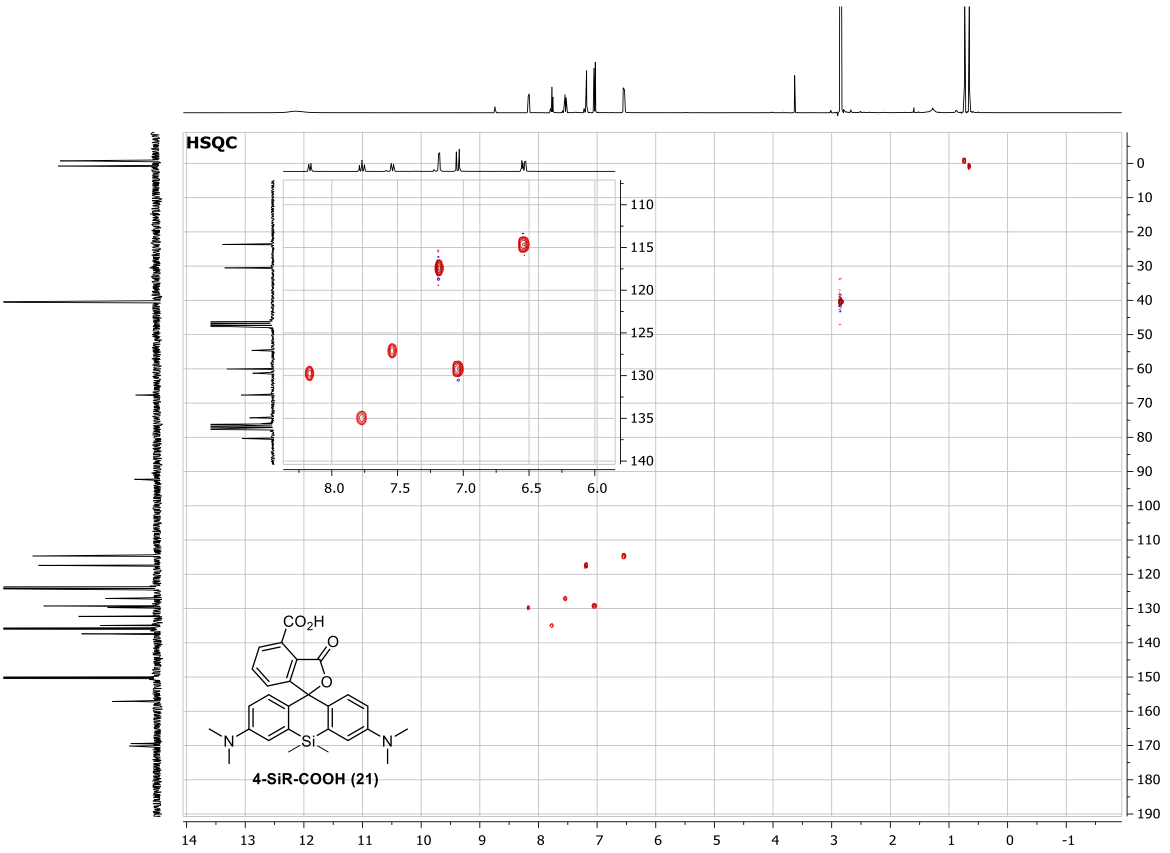
HSQC
$CO_2H$
O
O
N
Si
N
4-SiR-COOH (21)

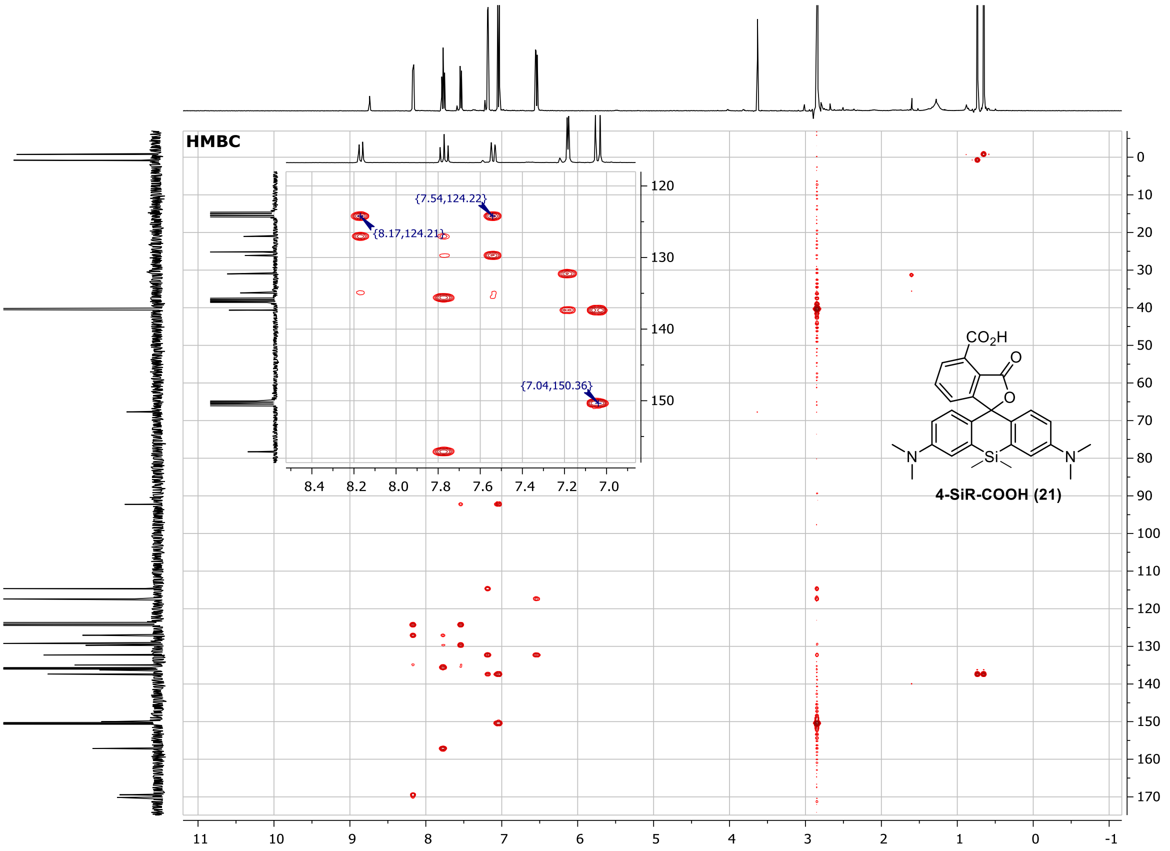
HMBC
{7.54,124.22}
{8.17,124.21}
{7.04,150.36}
$CO_2H$
O
O
N
Si
N
4-SiR-COOH (21)

#### CTX-C8-NHBoc (SI-1)

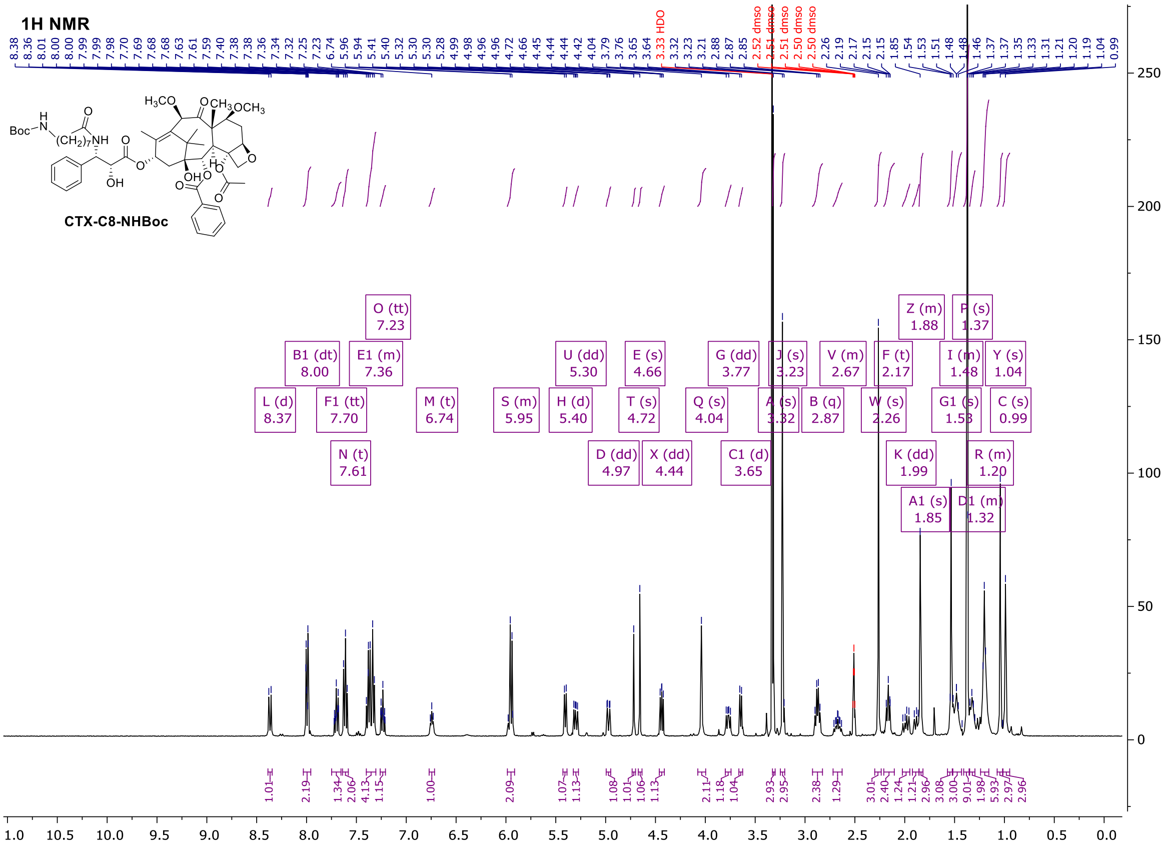

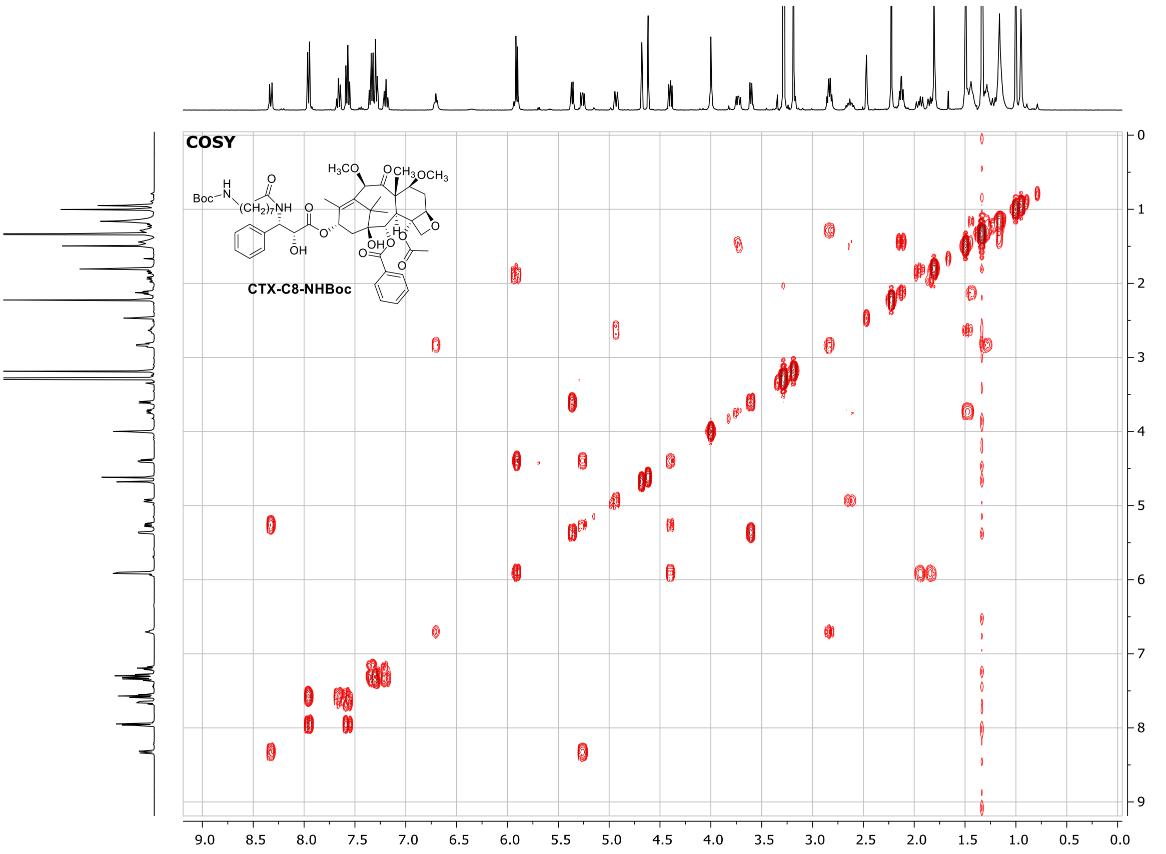

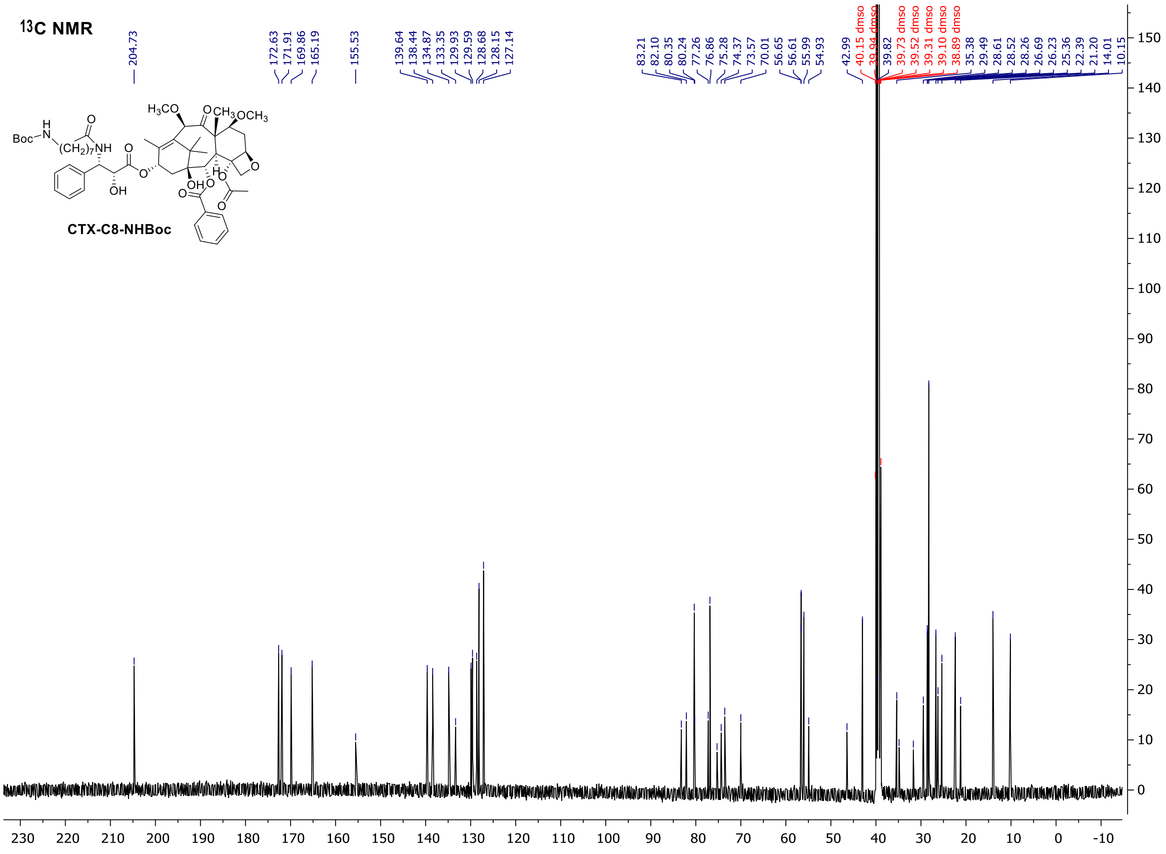

#### LTX-C8-NHBoc (SI-2)

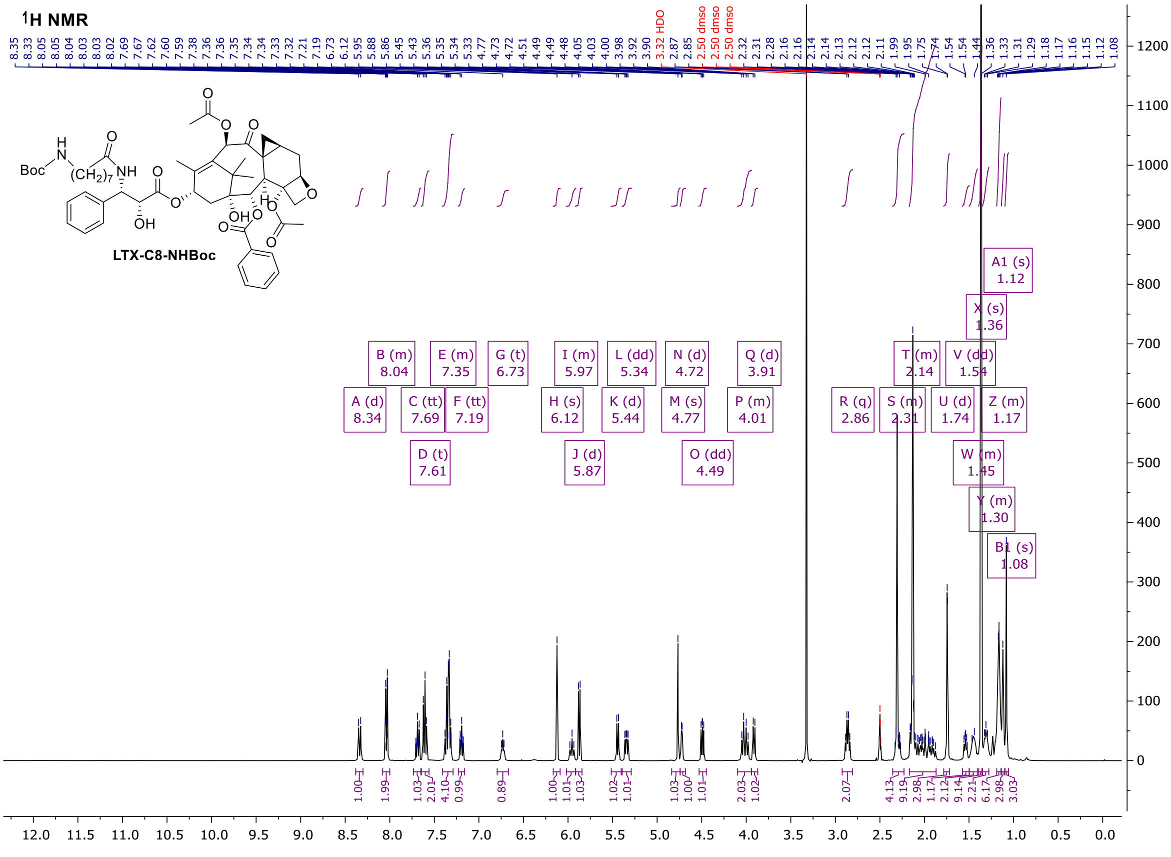

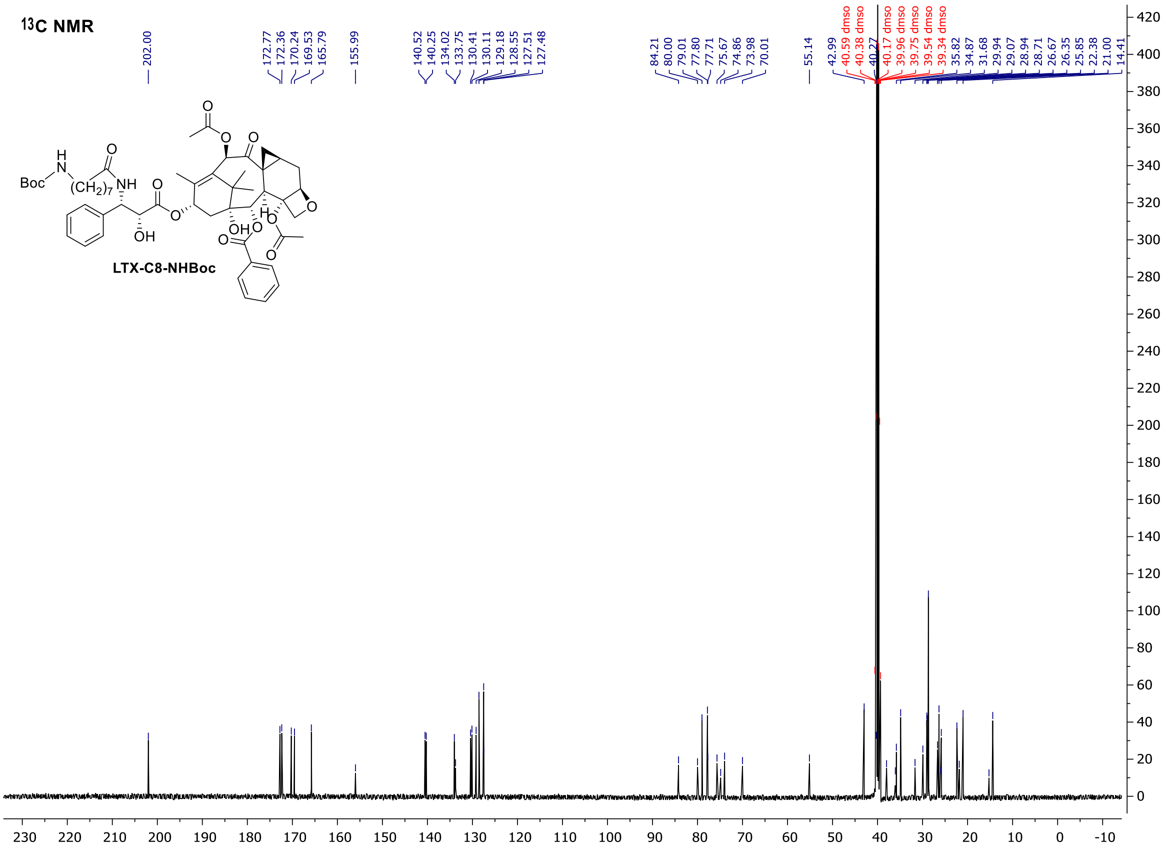

#### 4-TMR-LTX (22)

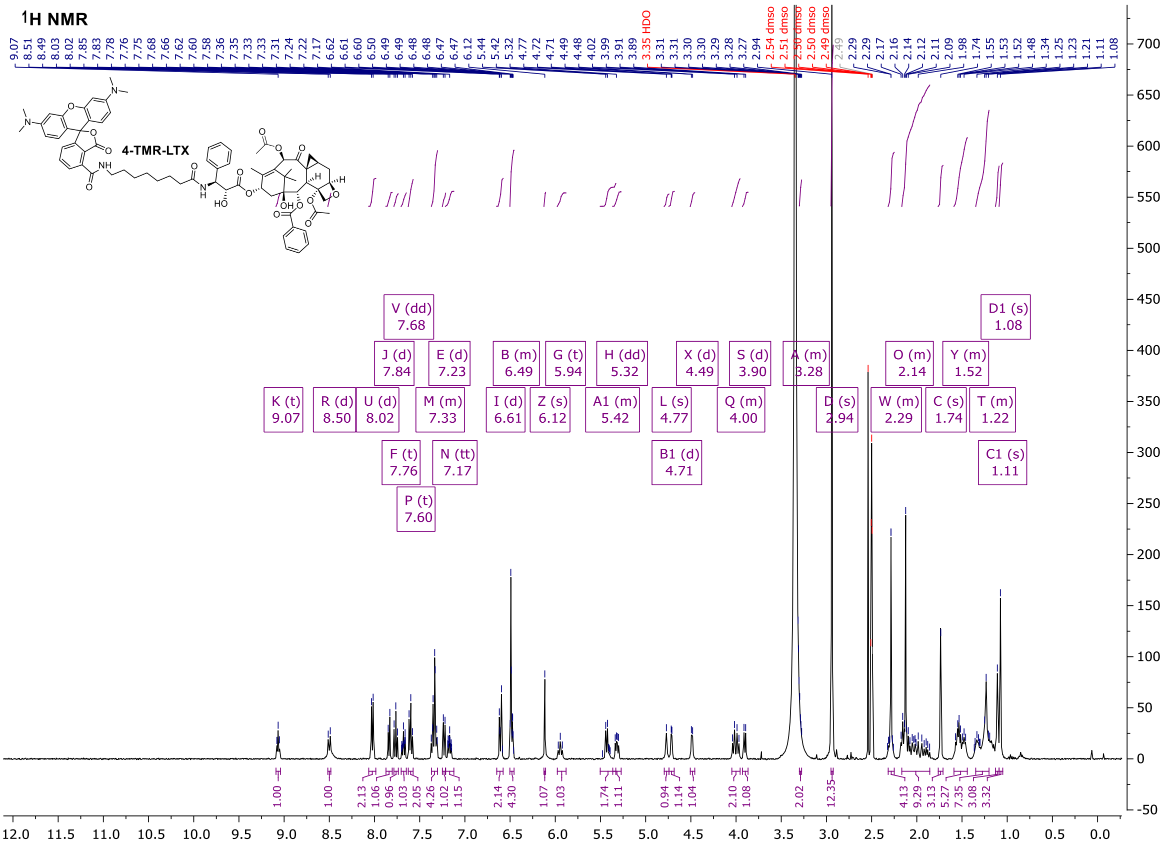

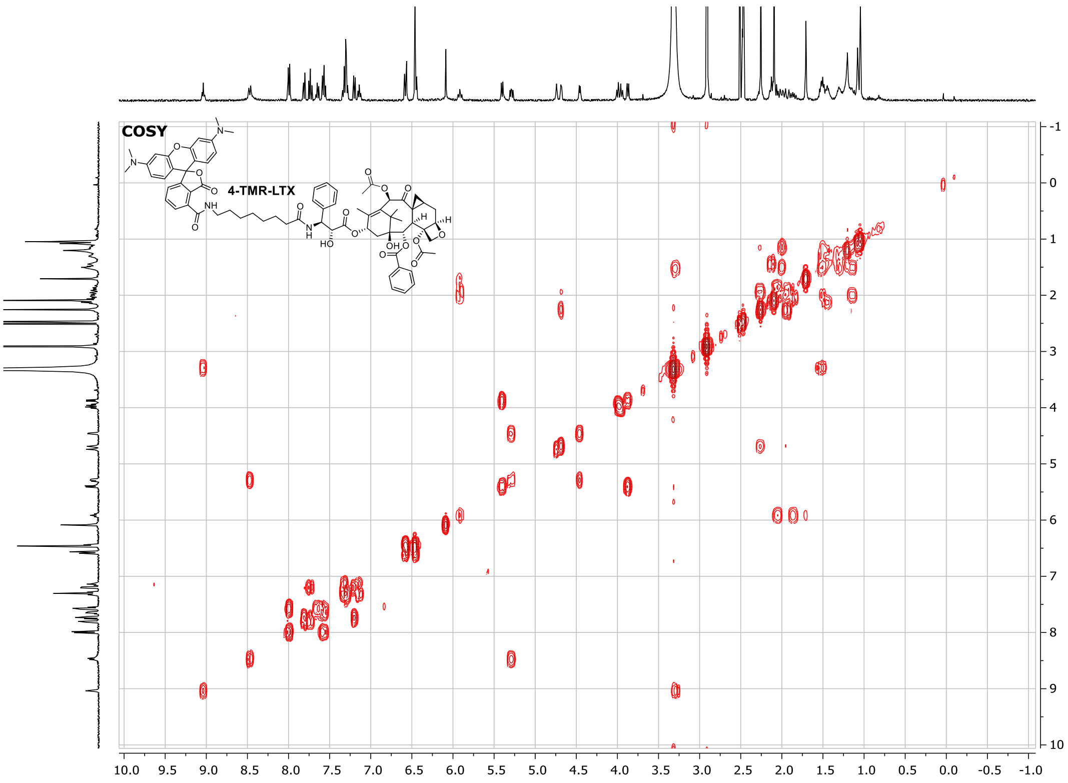
COSY
4-TMR-LTX

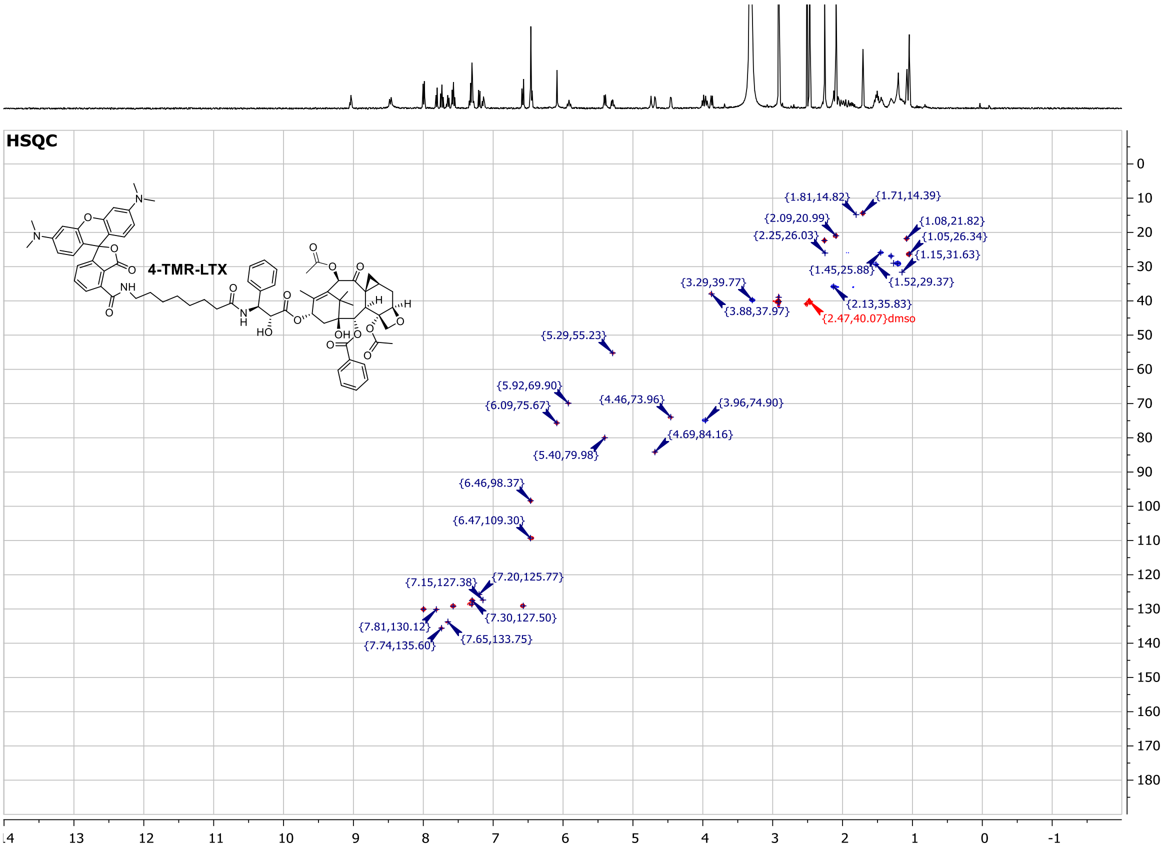
HSQC
4-TMR-LTX
{1.81,14.82}
{1.71,14.39}
{2.09,20.99}
{1.08,21.82}
{2.25,26.03}
{1.05,26.34}
{1.15,31.63}
{1.45,25.88}
{1.52,29.37}
{3.29,39.77}
{3.88,37.97}
{2.13,35.83}
{2.47,40.07}dmso
{5.29,55.23}
{5.92,69.90}
{6.09,75.67}
{4.46,73.96}
{3.96,74.90}
{4.69,84.16}
{5.40,79.98}
{6.46,98.37}
{6.47,109.30}
{7.15,127.38}
{7.20,125.77}
{7.30,127.50}
{7.81,130.12}
{7.74,135.60}
{7.65,133.75}

### 5-TMR-LTX (23)

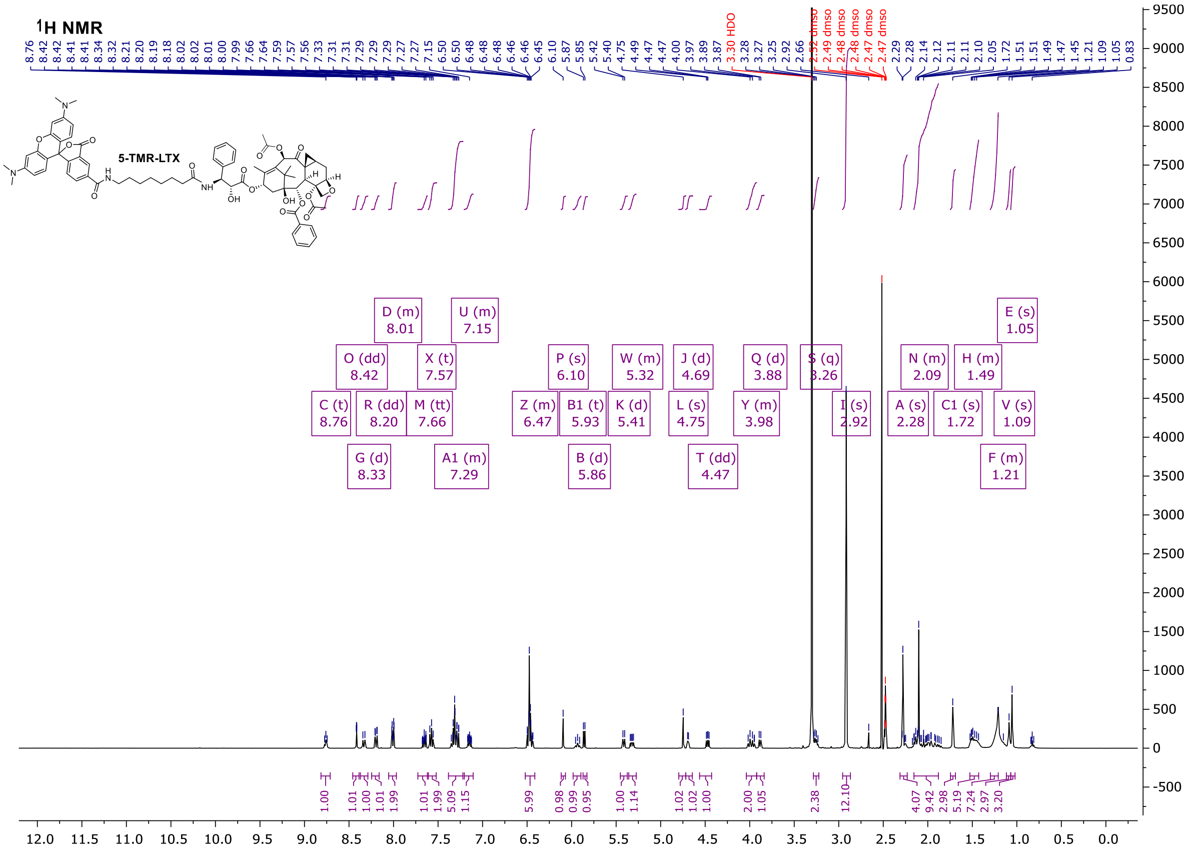

1H NMR
5-TMR-LTX
D (m) 8.01
U (m) 7.15
O (dd) 8.42
X (t) 7.57
P (s) 6.10
W (m) 5.32
J (d) 4.69
Q (d) 3.88
S (q) 3.26
N (m) 2.09
H (m) 1.49
E (s) 1.05
C (t) 8.76
R (dd) 8.20
M (tt) 7.66
Z (m) 6.47
B1 (t) 5.93
K (d) 5.41
L (s) 4.75
Y (m) 3.98
I (s) 2.92
A (s) 2.28
C1 (s) 1.72
V (s) 1.09
G (d) 8.33
A1 (m) 7.29
B (d) 5.86
T (dd) 4.47
F (m) 1.21

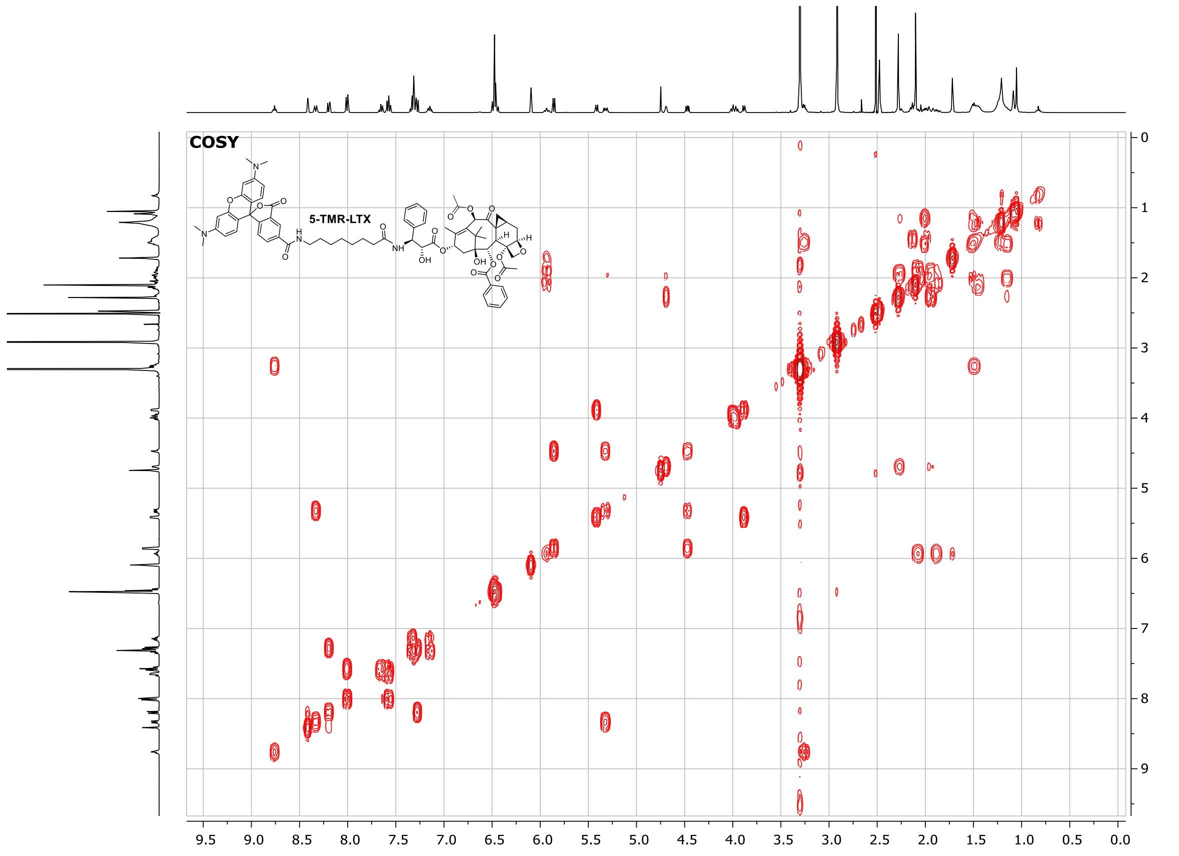

COSY
5-TMR-LTX

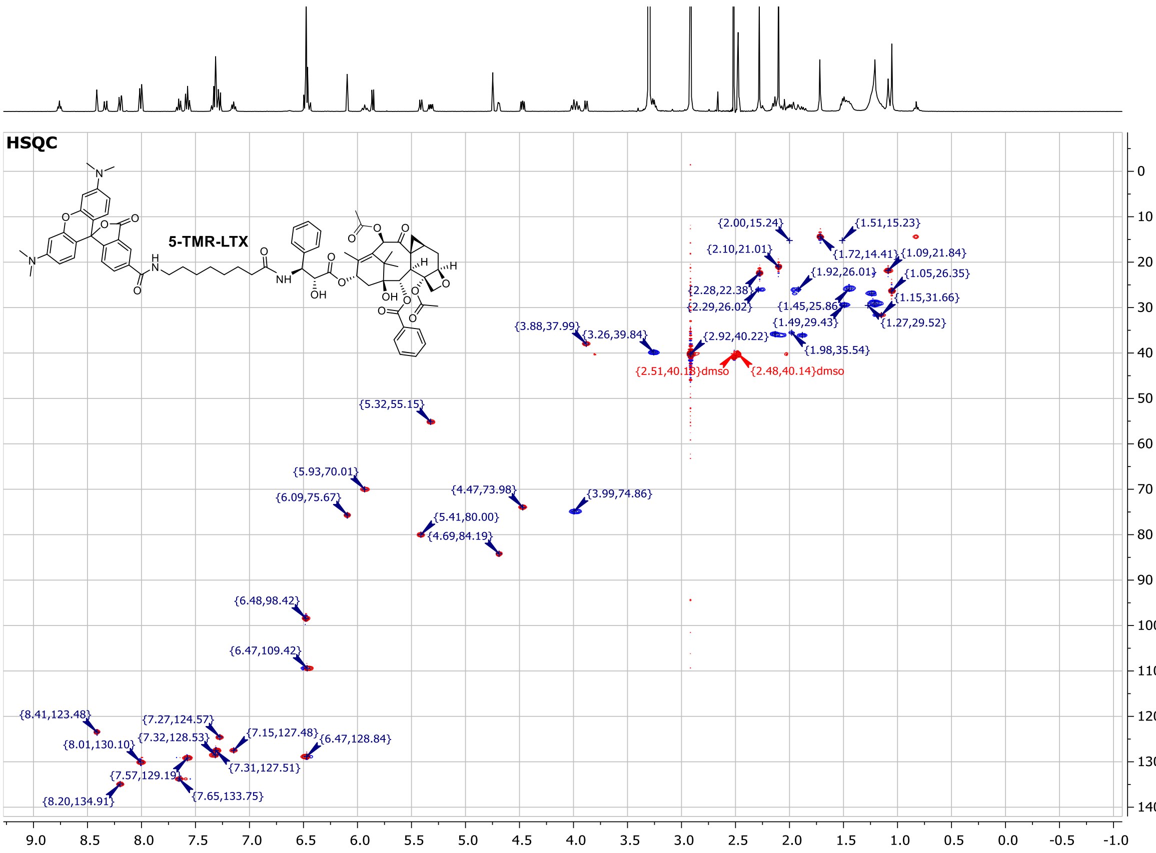

#### 6-TMR-LTX (24)

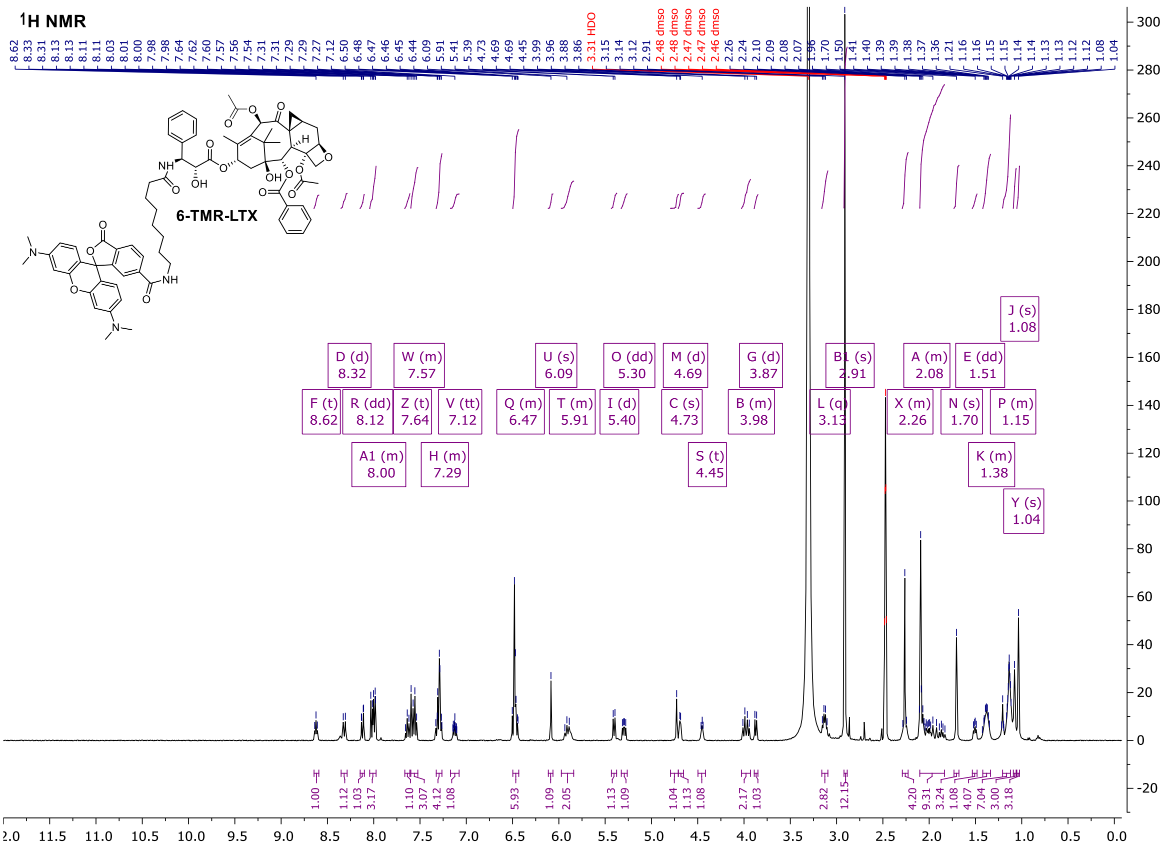

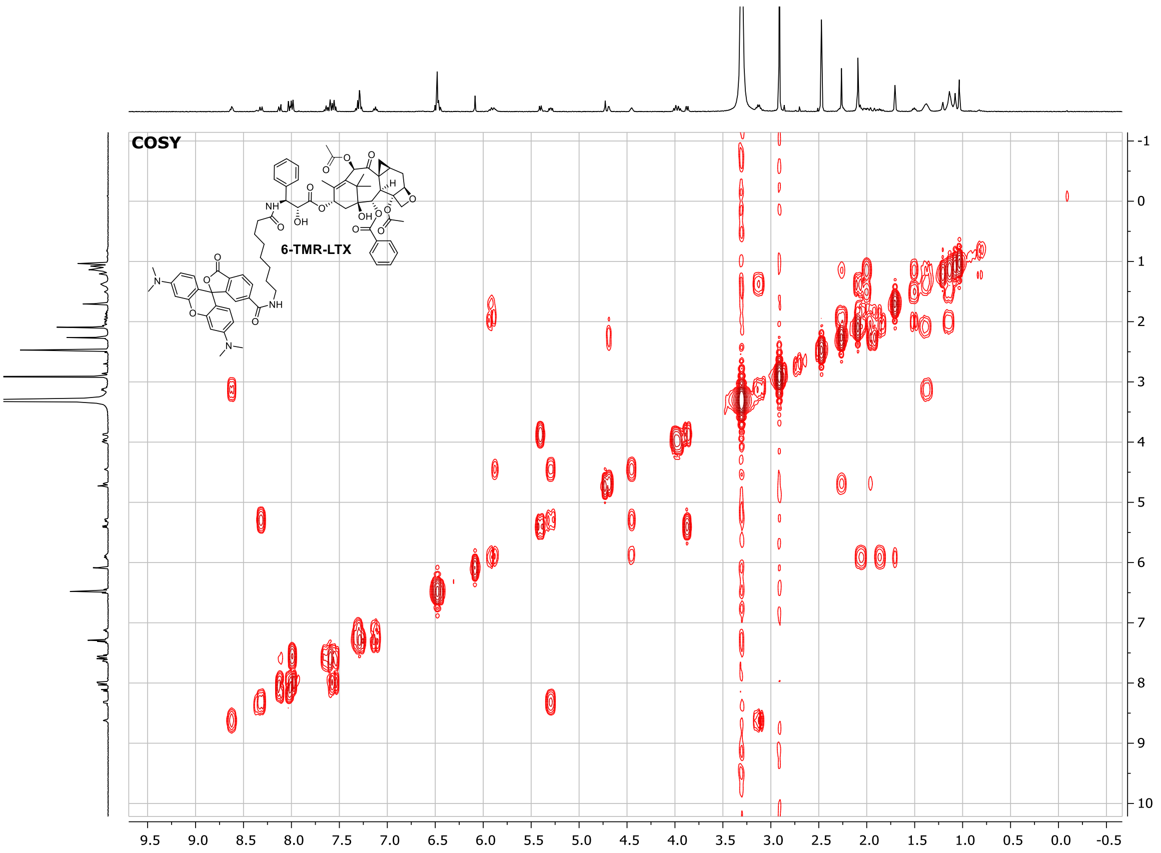
COSY
6-TMR-LTX

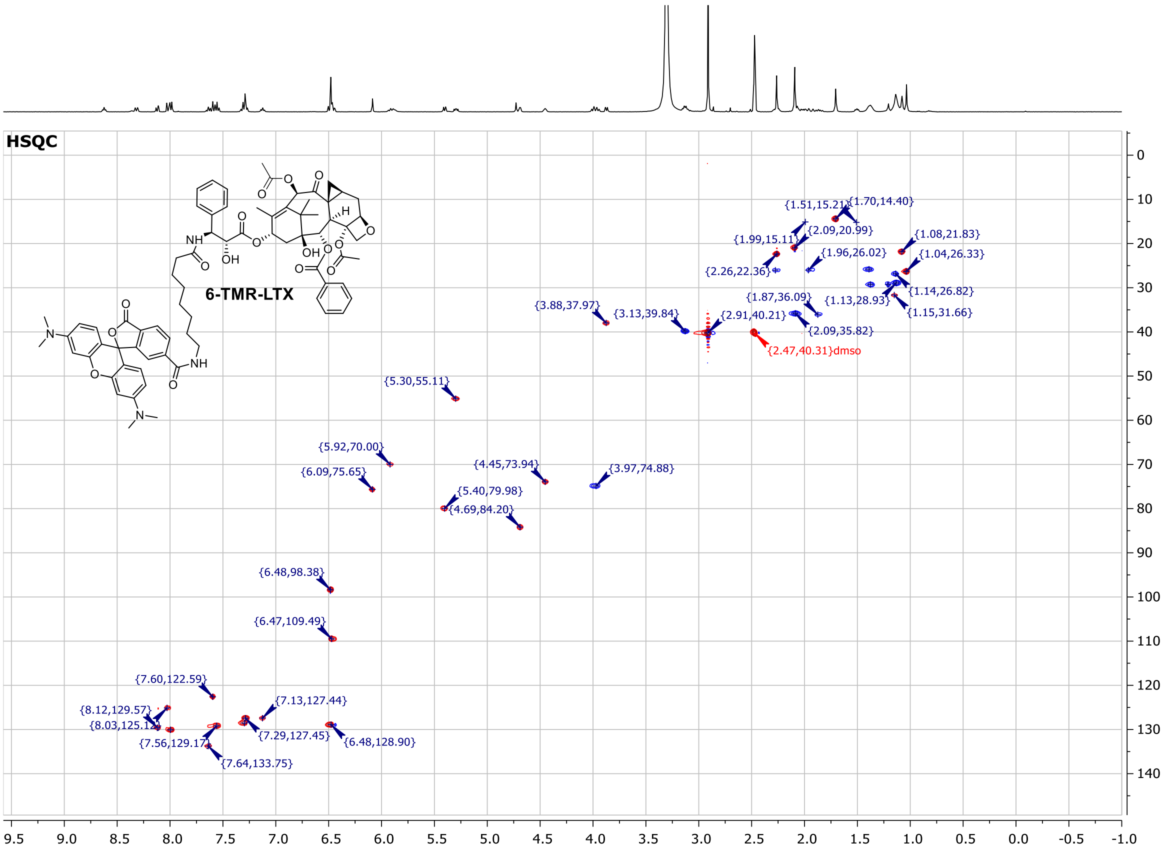
HSQC
6-TMR-LTX
{1.51,15.21}
{1.70,14.40}
{2.09,20.99}
{1.99,15.11}
{1.08,21.83}
{1.96,26.02}
{1.04,26.33}
{2.26,22.36}
{1.14,26.82}
{1.87,36.09}
{1.13,28.93}
{3.88,37.97}
{3.13,39.84}
{2.91,40.21}
{1.15,31.66}
{2.09,35.82}
{2.47,40.31}dmso
{5.30,55.11}
{5.92,70.00}
{6.09,75.65}
{4.45,73.94}
{3.97,74.88}
{5.40,79.98}
{4.69,84.20}
{6.48,98.38}
{6.47,109.49}
{7.60,122.59}
{7.13,127.44}
{8.12,129.57}
{8.03,125.12}
{7.29,127.45}
{6.48,128.90}
{7.56,129.17}
{7.64,133.75}

#### 4-580CP-LTX (25)

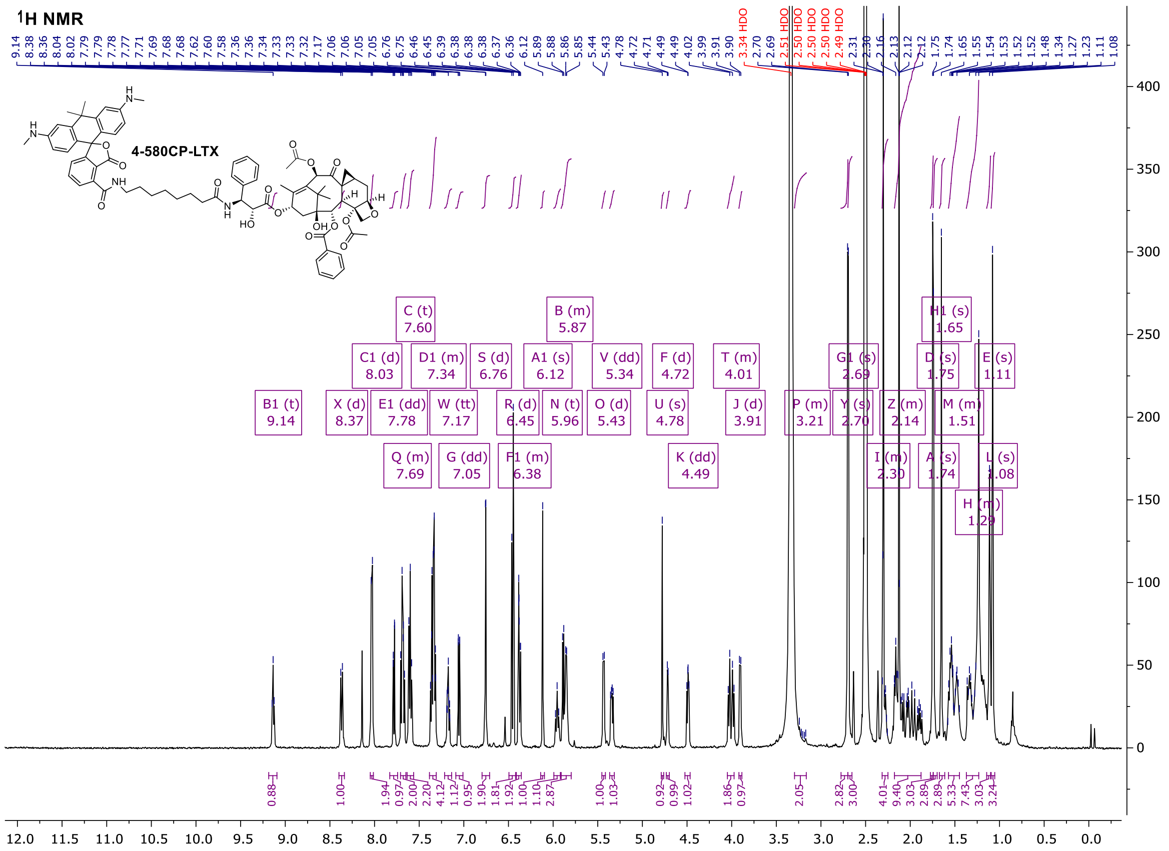

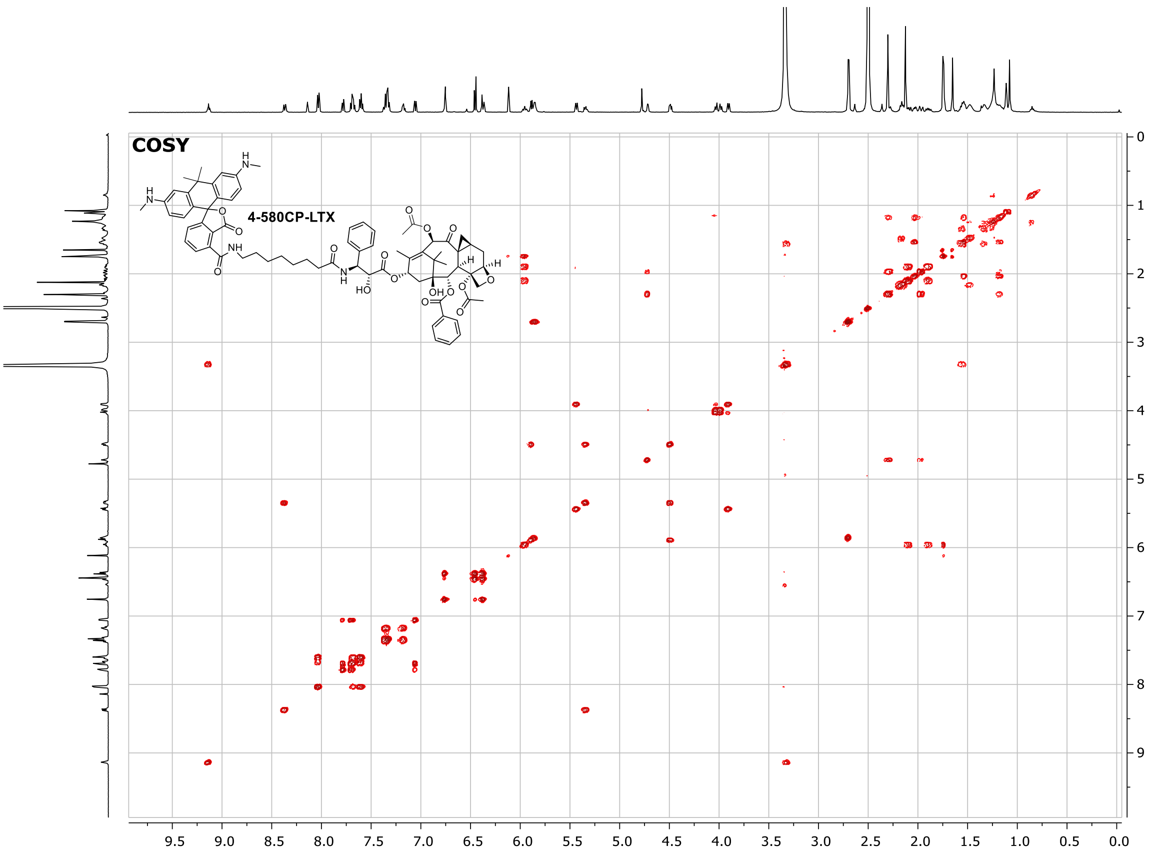

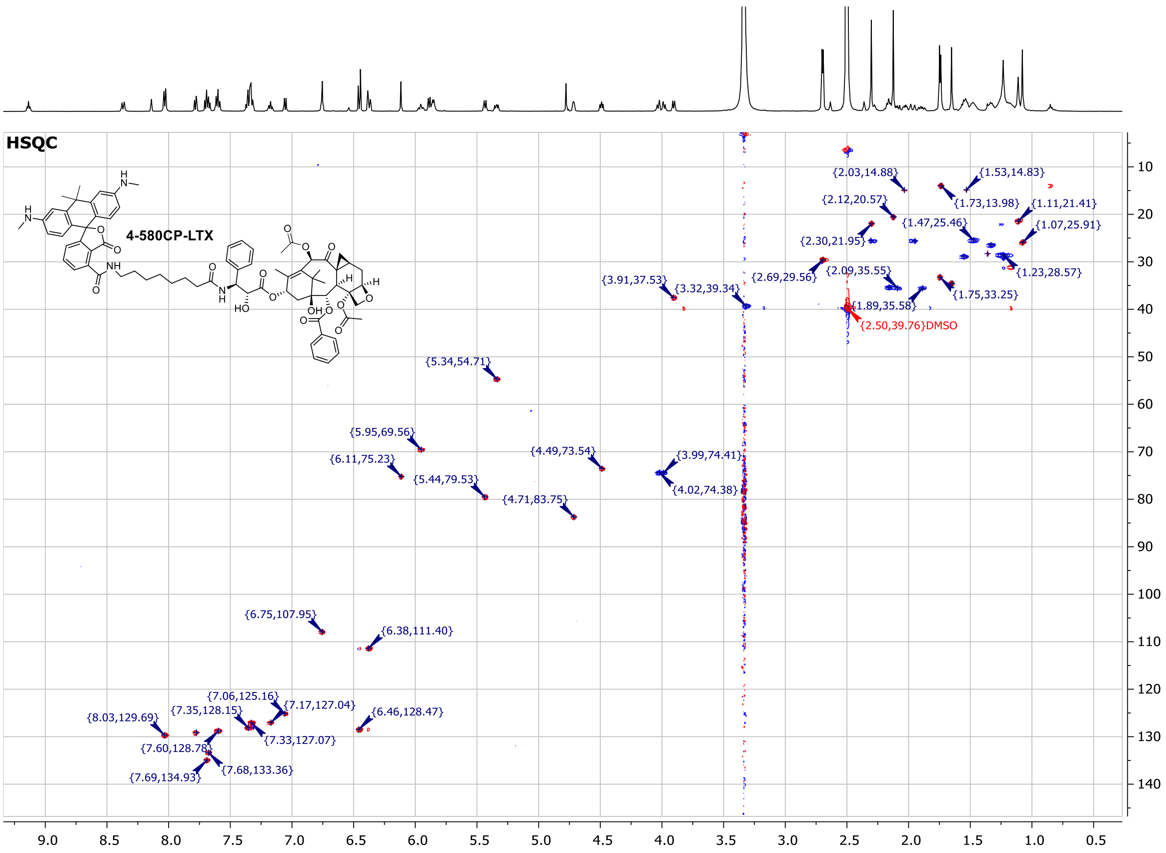

#### 5-580CP-LTX (26)

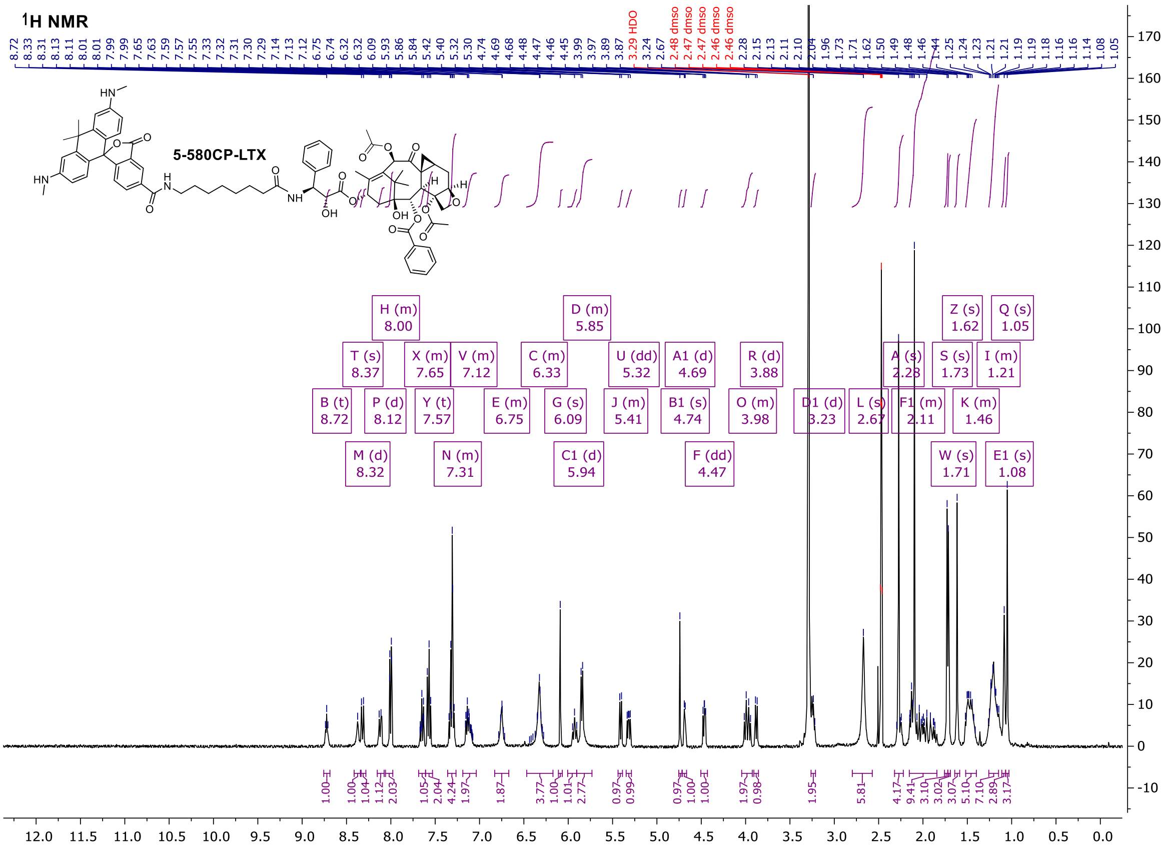

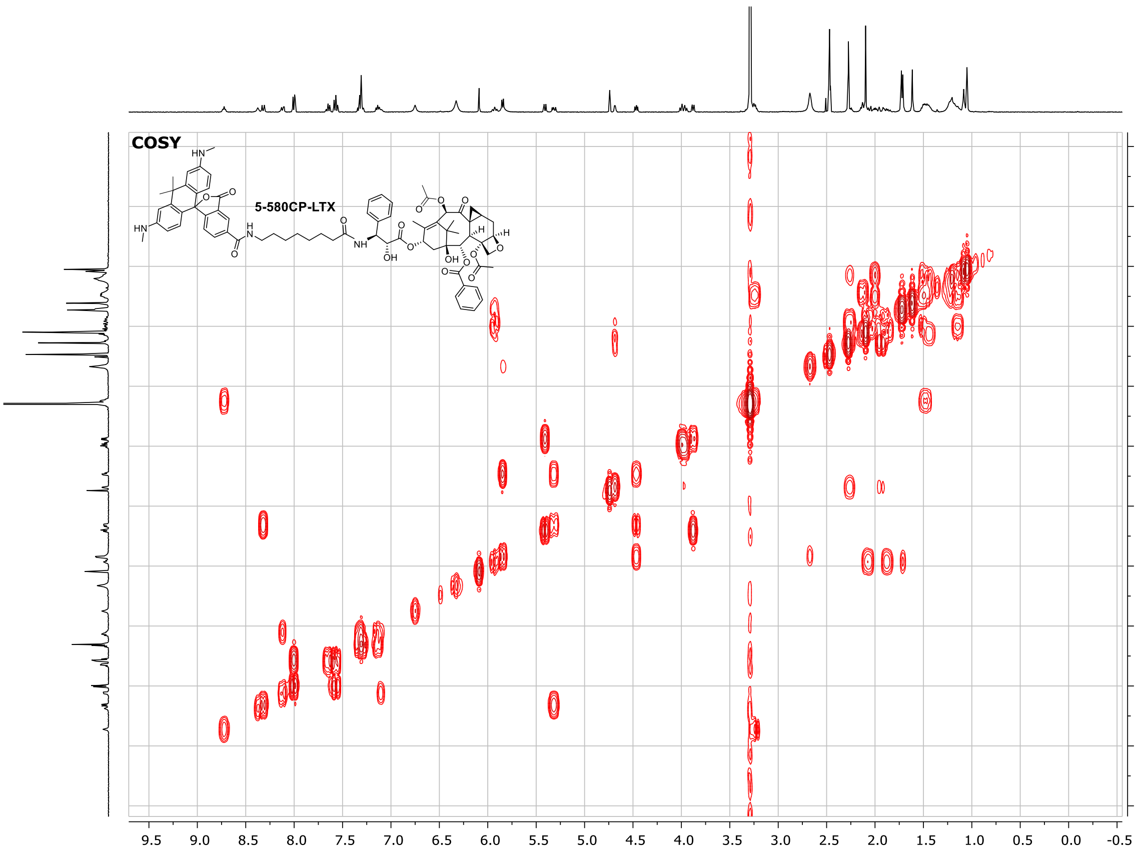
COSY
5-580CP-LTX
HN
OH
O
NH
H
9.5 9.0 8.5 8.0 7.5 7.0 6.5 6.0 5.5 5.0 4.5 4.0 3.5 3.0 2.5 2.0 1.5 1.0 0.5 0.0 -0.5
-1 0 1 2 3 4 5 6 7 8 9 10

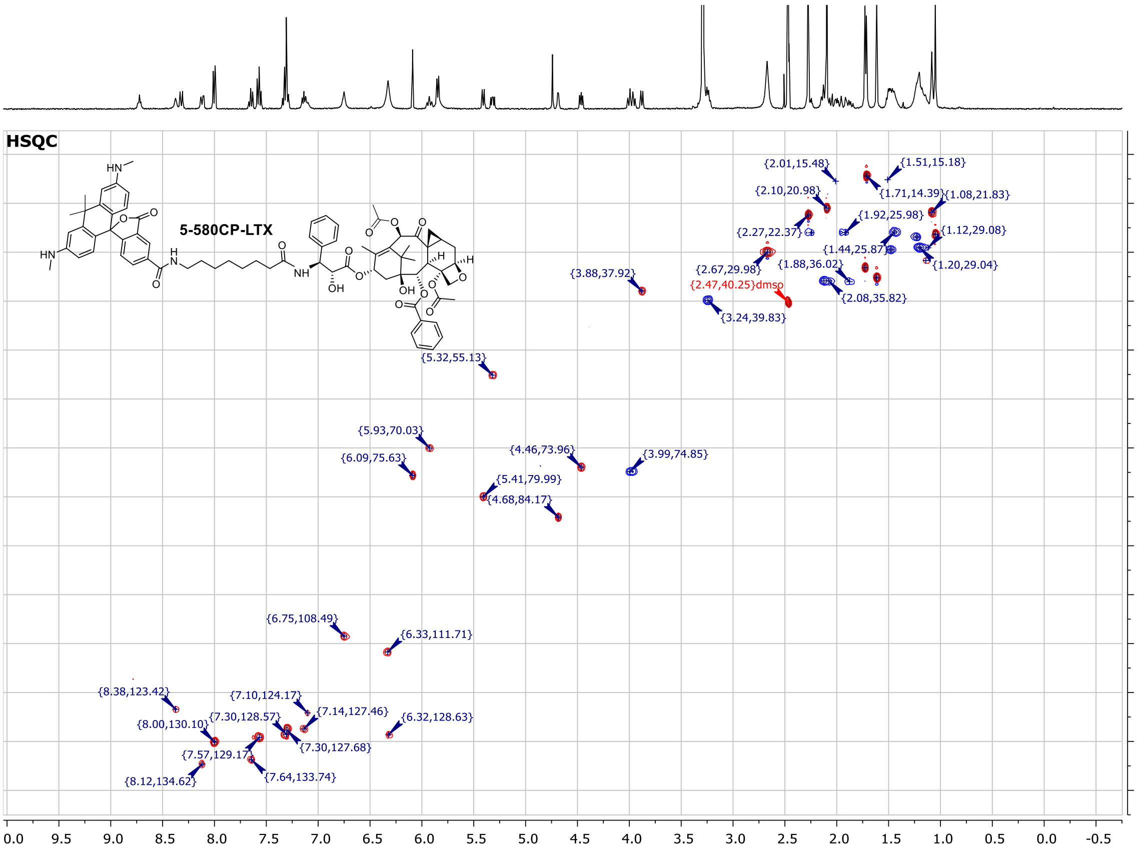
HSQC
5-580CP-LTX
{2.01,15.48}
{1.51,15.18}
{2.10,20.98}
{1.71,14.39}
{1.08,21.83}
{1.92,25.98}
{2.27,22.37}
{1.12,29.08}
{1.44,25.87}
{2.67,29.98}
{1.88,36.02}
{1.20,29.04}
{3.88,37.92}
{2.47,40.25}dmso
{2.08,35.82}
{3.24,39.83}
{5.32,55.13}
{5.93,70.03}
{4.46,73.96}
{3.99,74.85}
{6.09,75.63}
{5.41,79.99}
{4.68,84.17}
{6.75,108.49}
{6.33,111.71}
{8.38,123.42}
{7.10,124.17}
{7.14,127.46}
{7.30,128.57}
{6.32,128.63}
{8.00,130.10}
{7.30,127.68}
{7.57,129.17}
{8.12,134.62}
{7.64,133.74}
0.0 9.5 9.0 8.5 8.0 7.5 7.0 6.5 6.0 5.5 5.0 4.5 4.0 3.5 3.0 2.5 2.0 1.5 1.0 0.5 0.0 -0.5
10 20 30 40 50 60 70 80 90 100 110 120 130 140

### 4-610CP-CTX (28)

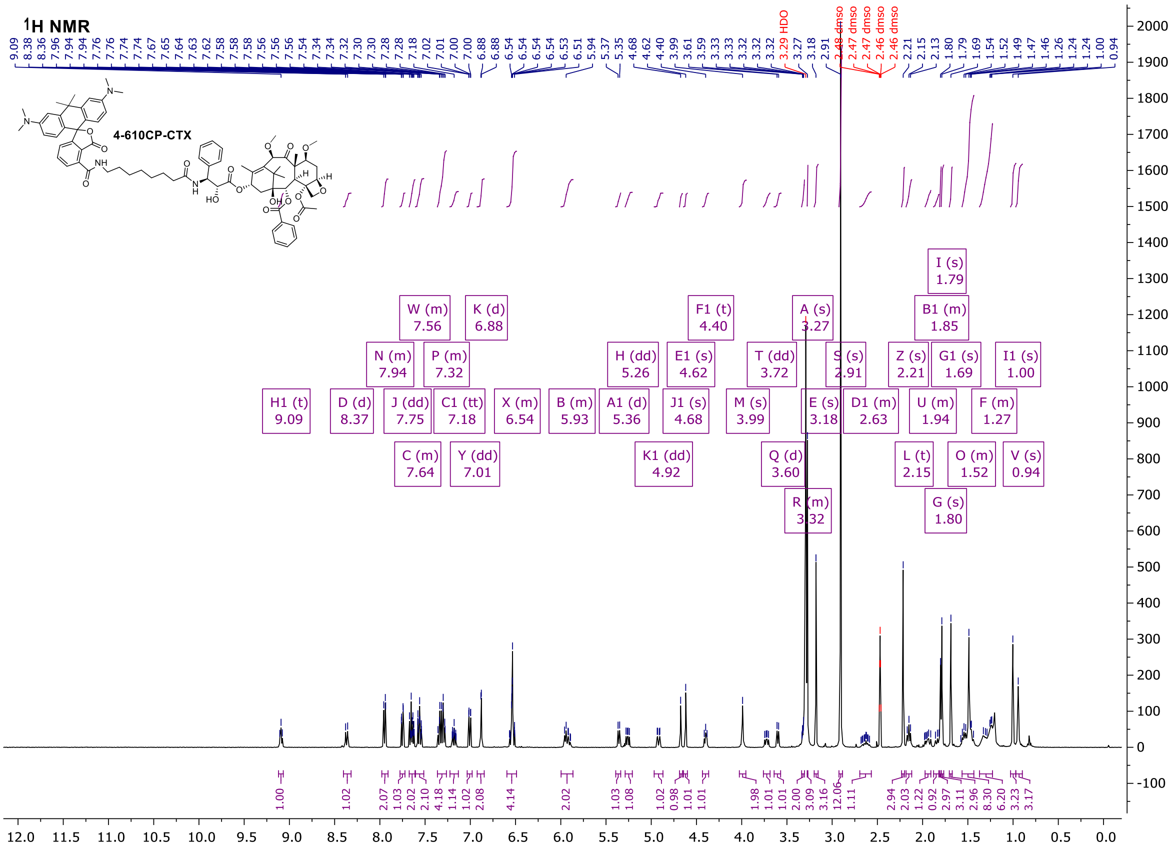

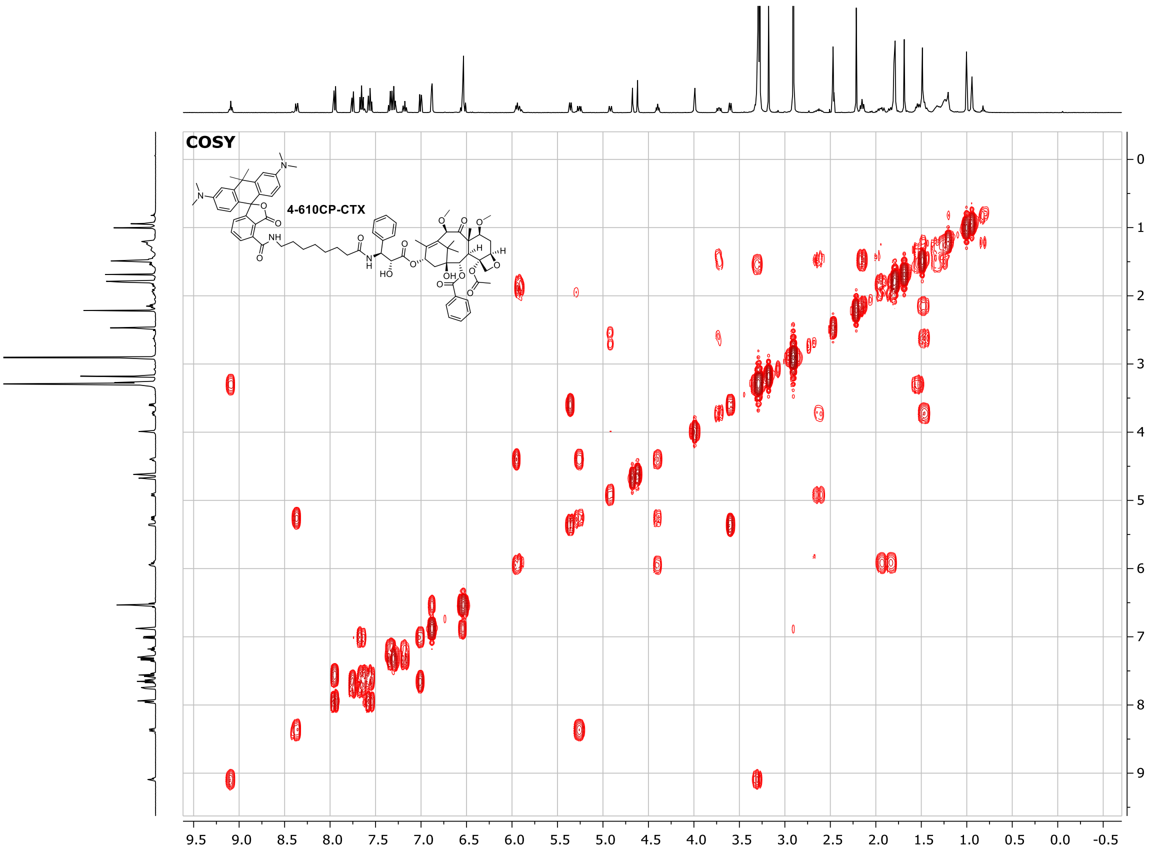

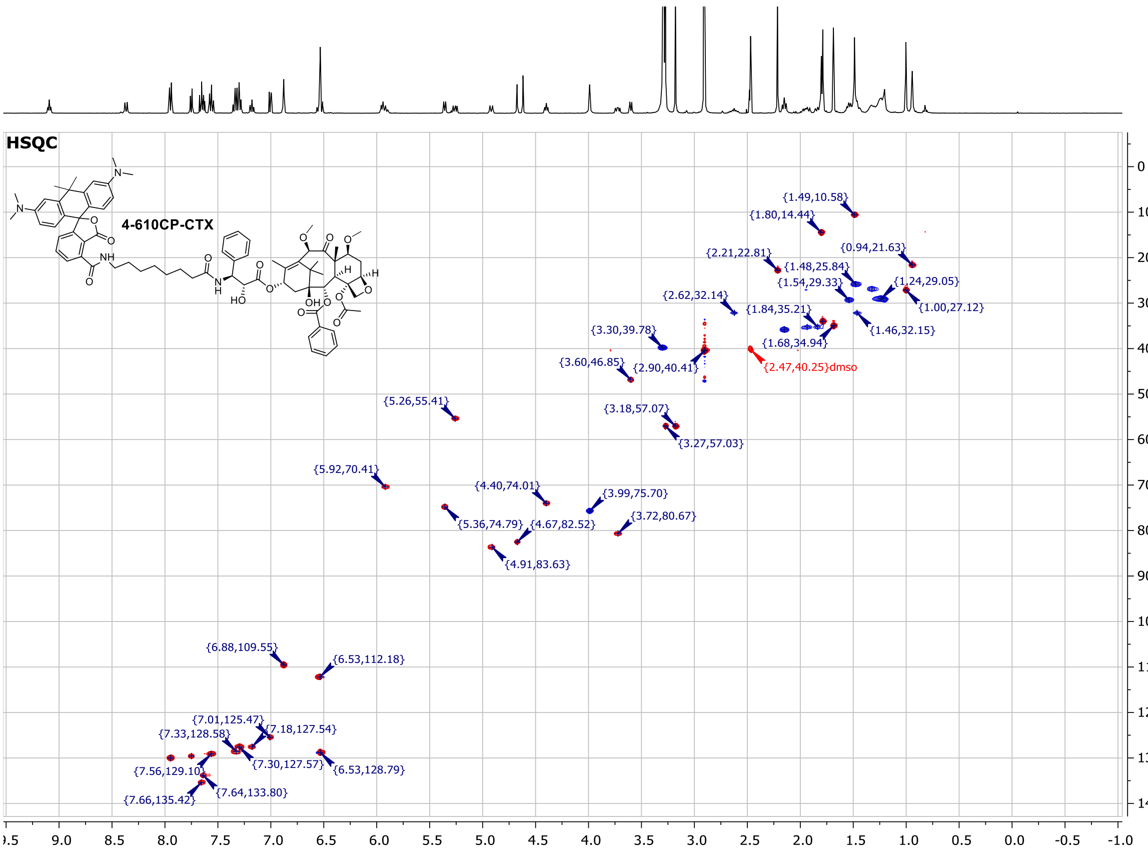

#### 5-610CP-CTX (29)

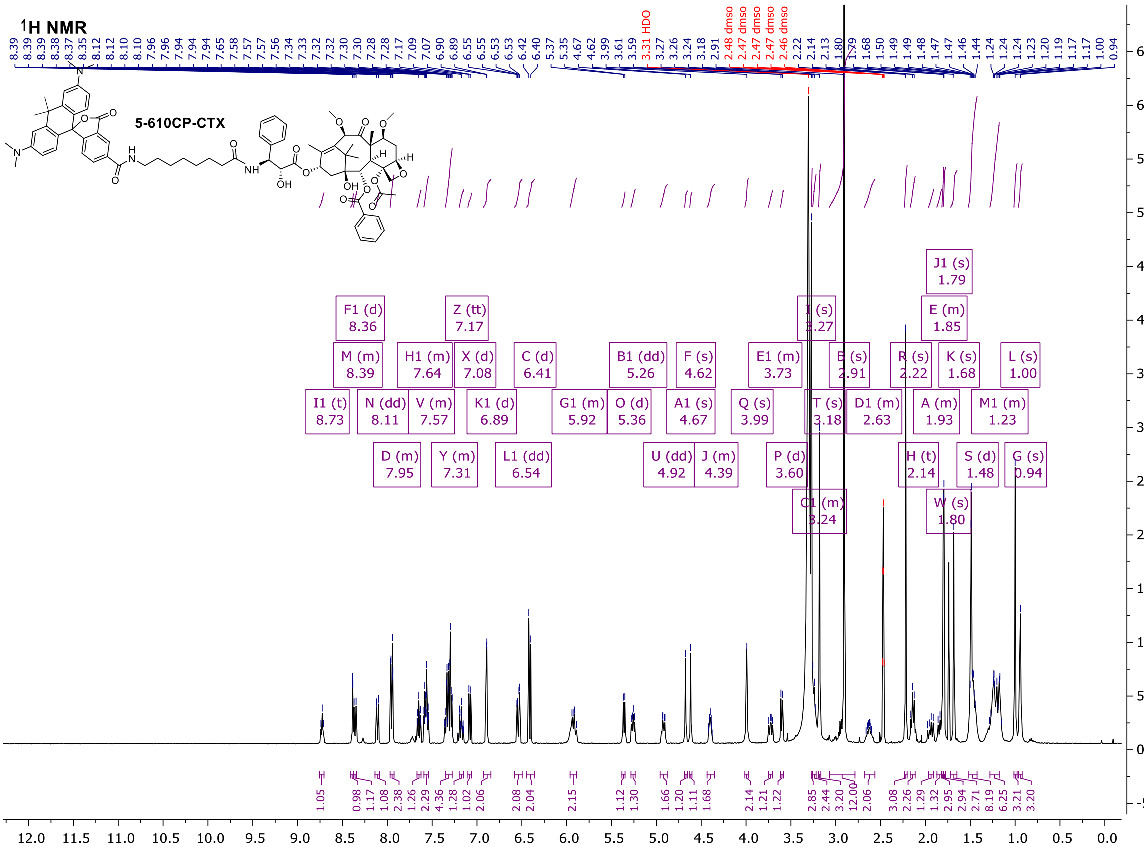

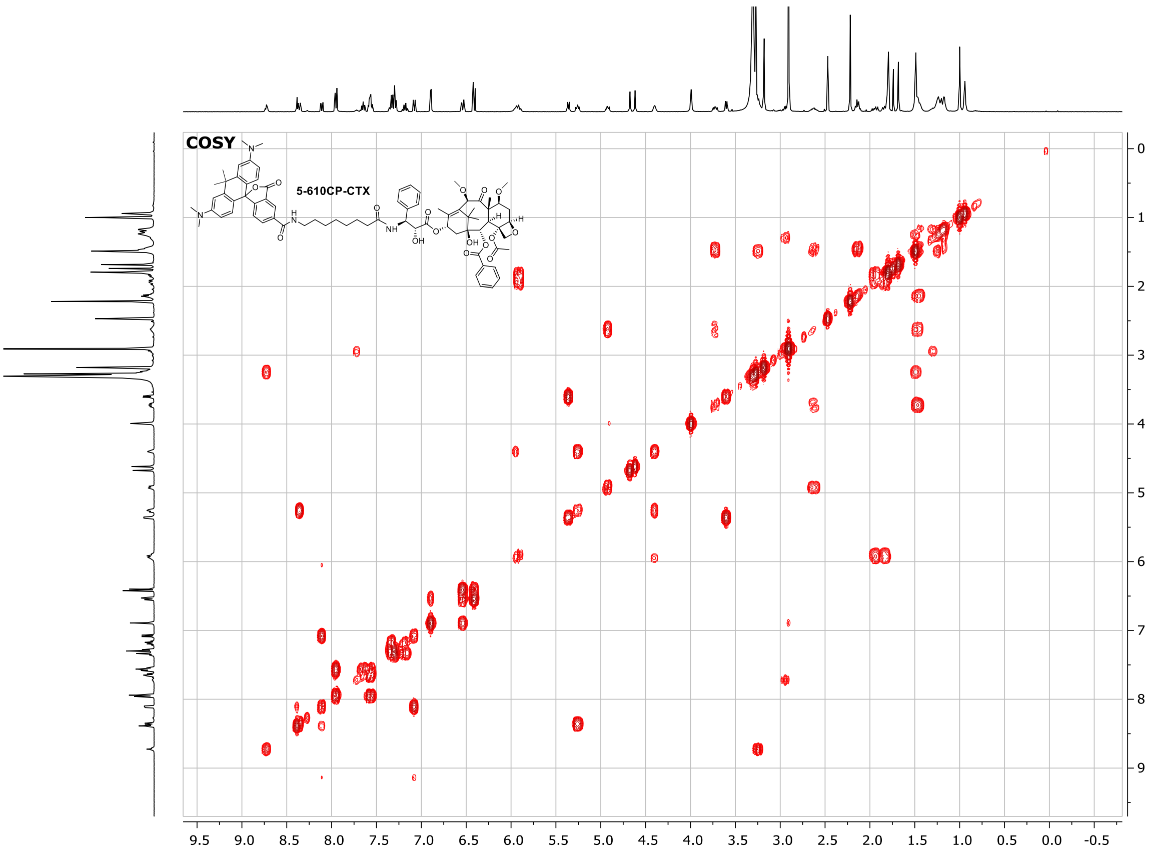

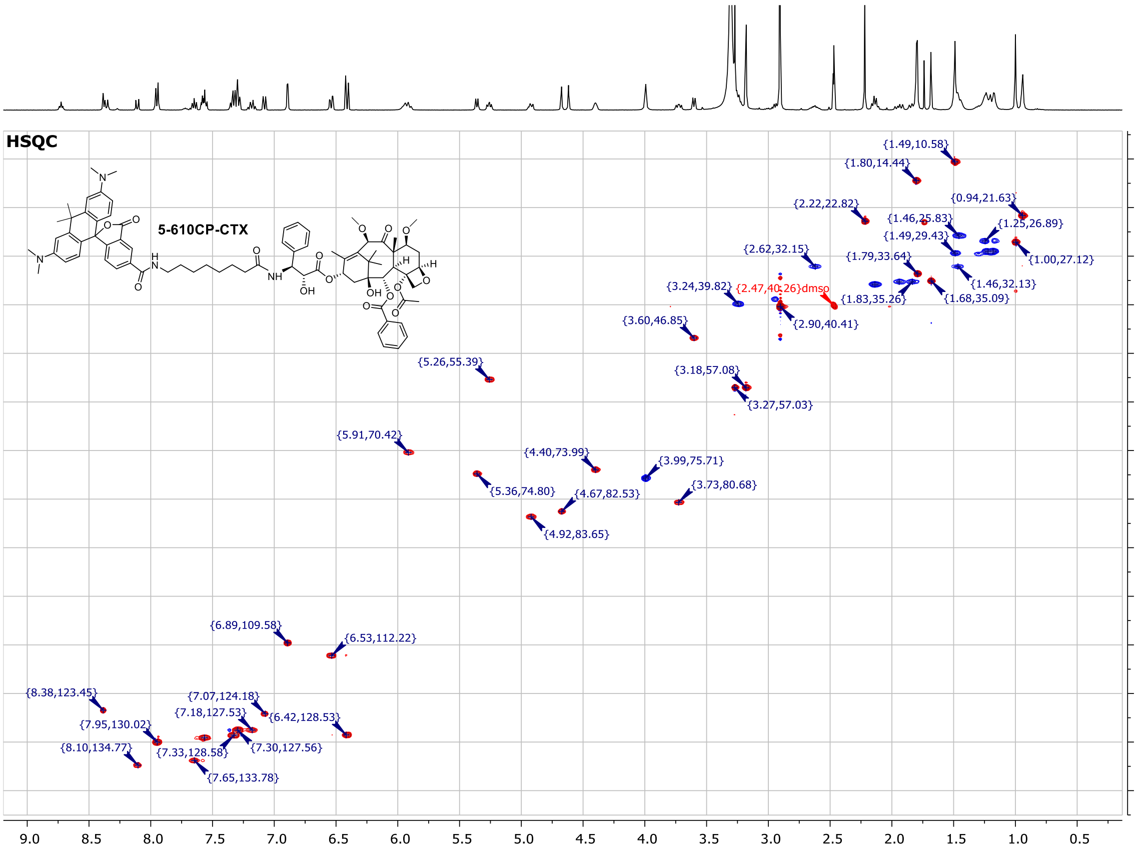

#### 6-610CP-CTX (30)

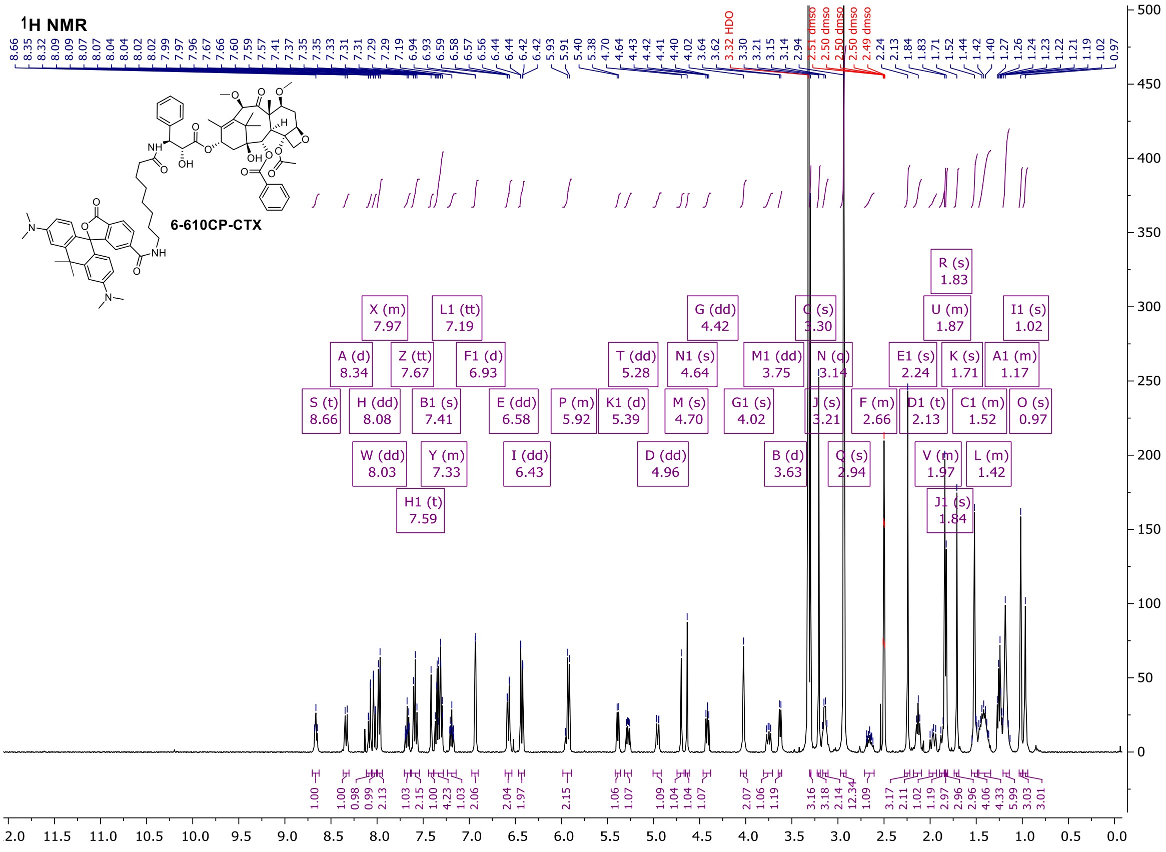
1H NMR
6-610CP-CTX

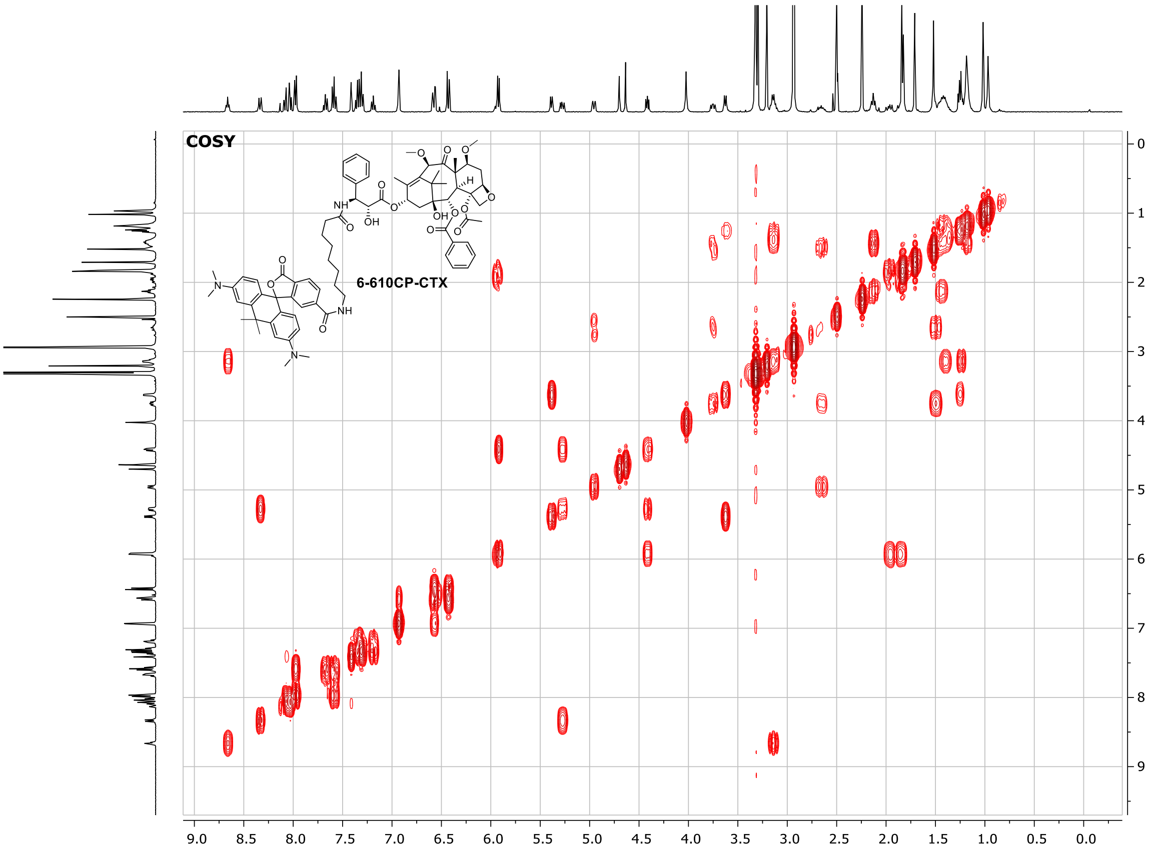
COSY
6-610CP-CTX

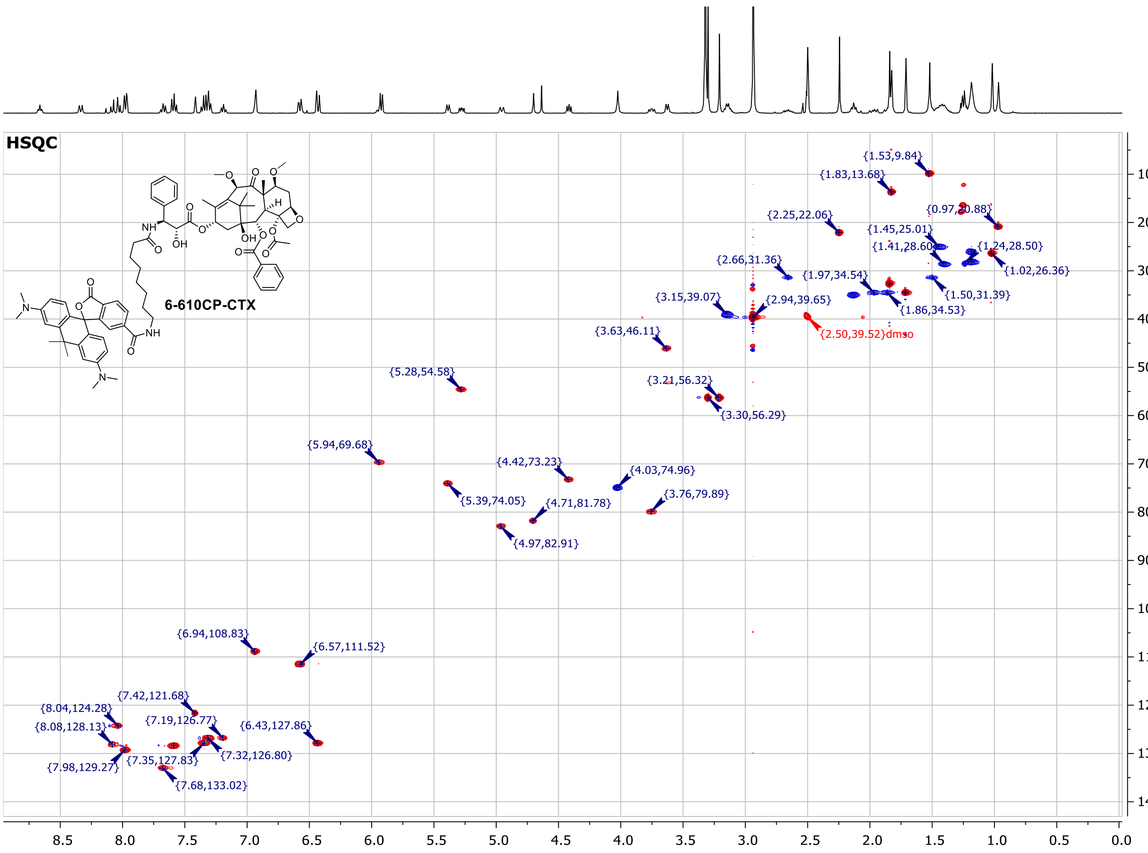

#### 4-SiR-CTX (31)

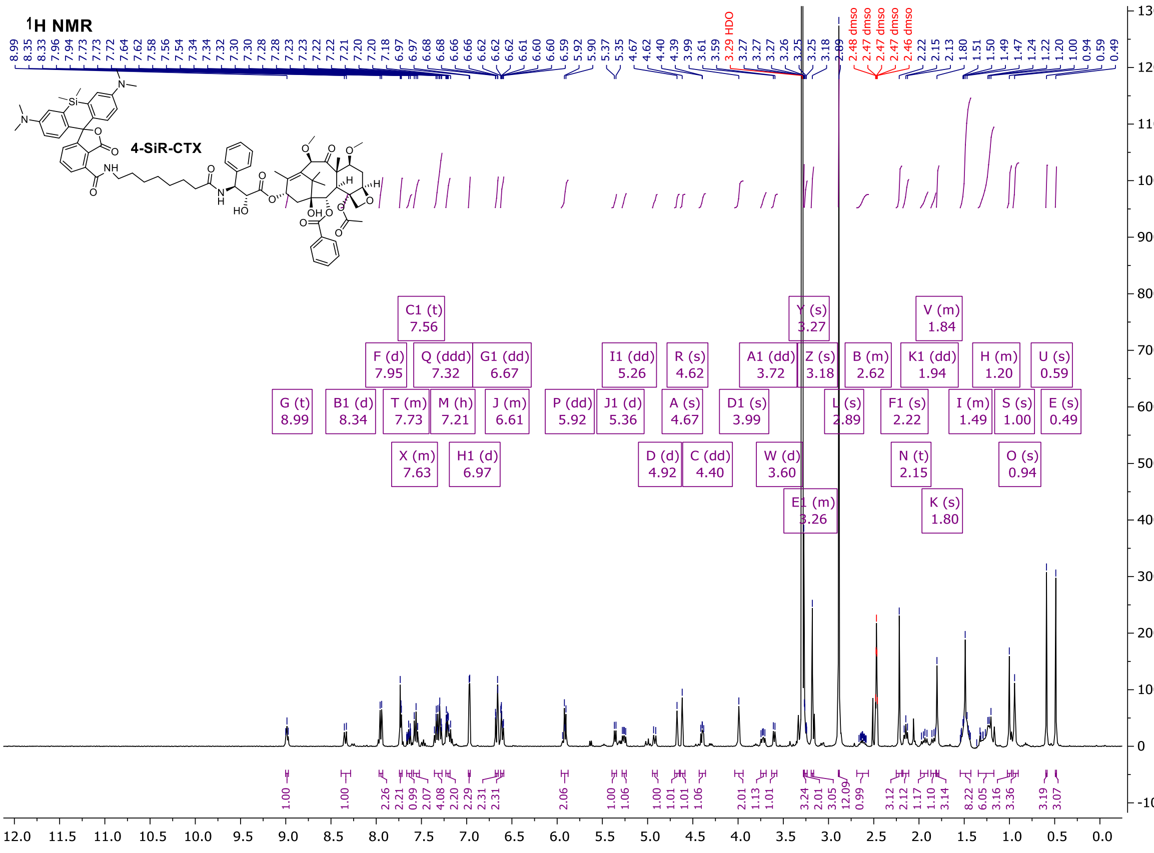

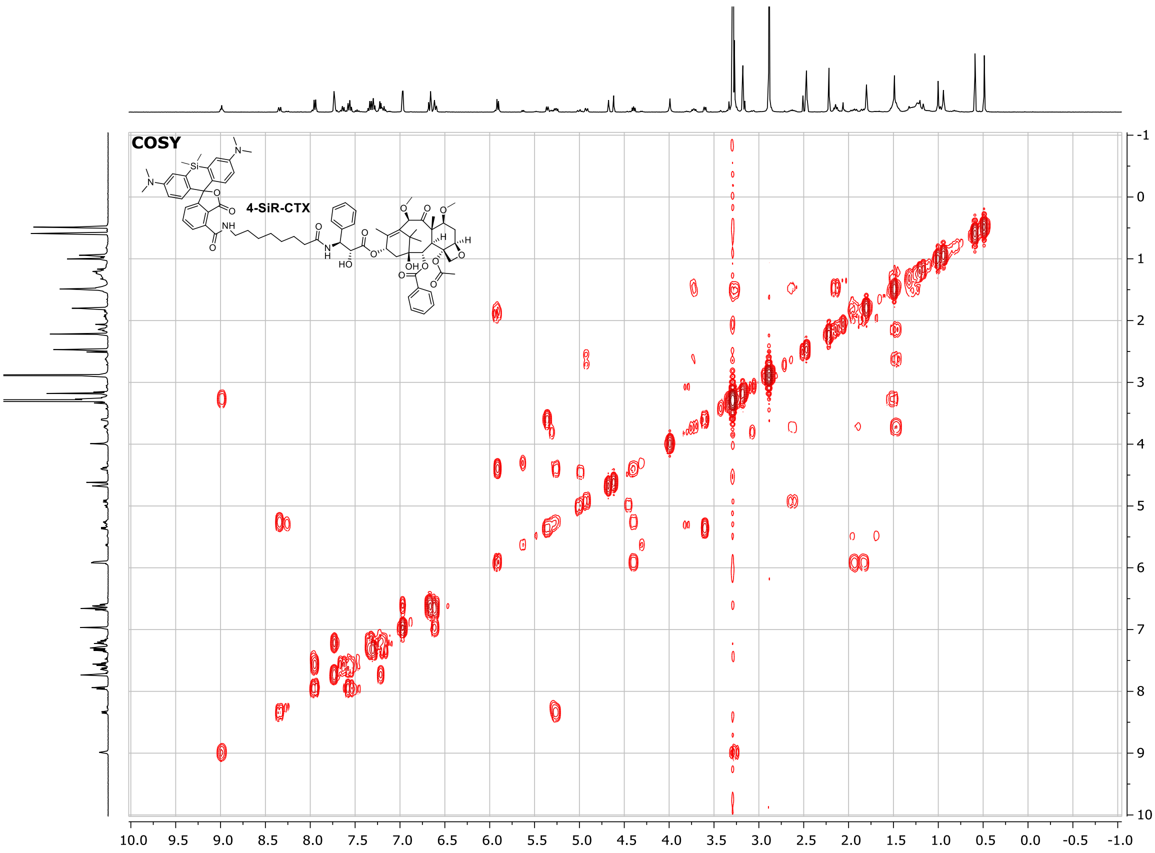
COSY
4-SiR-CTX

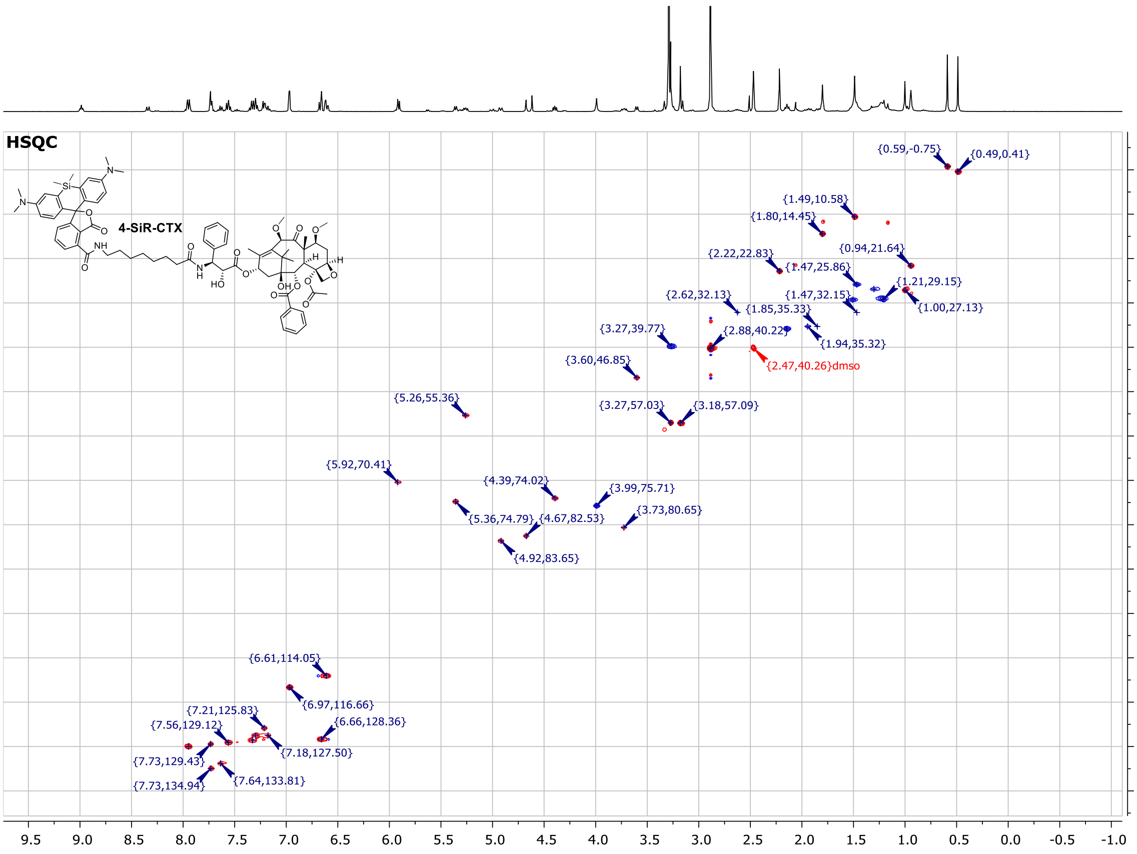
HSQC
4-SiR-CTX
{0.59,-0.75}
{0.49,0.41}
{1.49,10.58}
{1.80,14.45}
{0.94,21.64}
{2.22,22.83}
{1.47,25.86}
{1.21,29.15}
{1.47,32.15}
{1.00,27.13}
{2.62,32.13}
{1.85,35.33}
{3.27,39.77}
{2.88,40.22}
{1.94,35.32}
{2.47,40.26}dmso
{3.60,46.85}
{5.26,55.36}
{3.27,57.03}
{3.18,57.09}
{5.92,70.41}
{4.39,74.02}
{3.99,75.71}
{5.36,74.79}
{4.67,82.53}
{3.73,80.65}
{4.92,83.65}
{6.61,114.05}
{6.97,116.66}
{7.21,125.83}
{6.66,128.36}
{7.56,129.12}
{7.18,127.50}
{7.73,129.43}
{7.73,134.94}
{7.64,133.81}

#### 5-SiR-CTX (32)

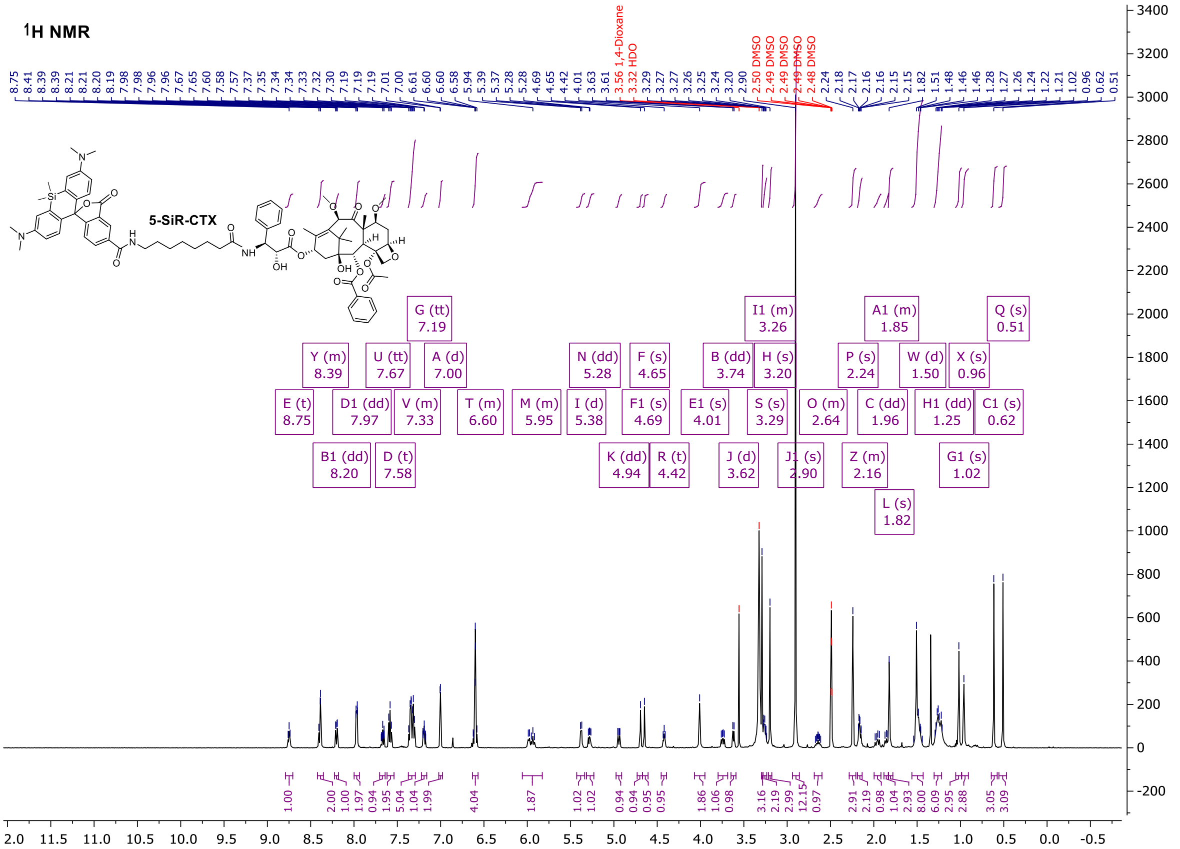

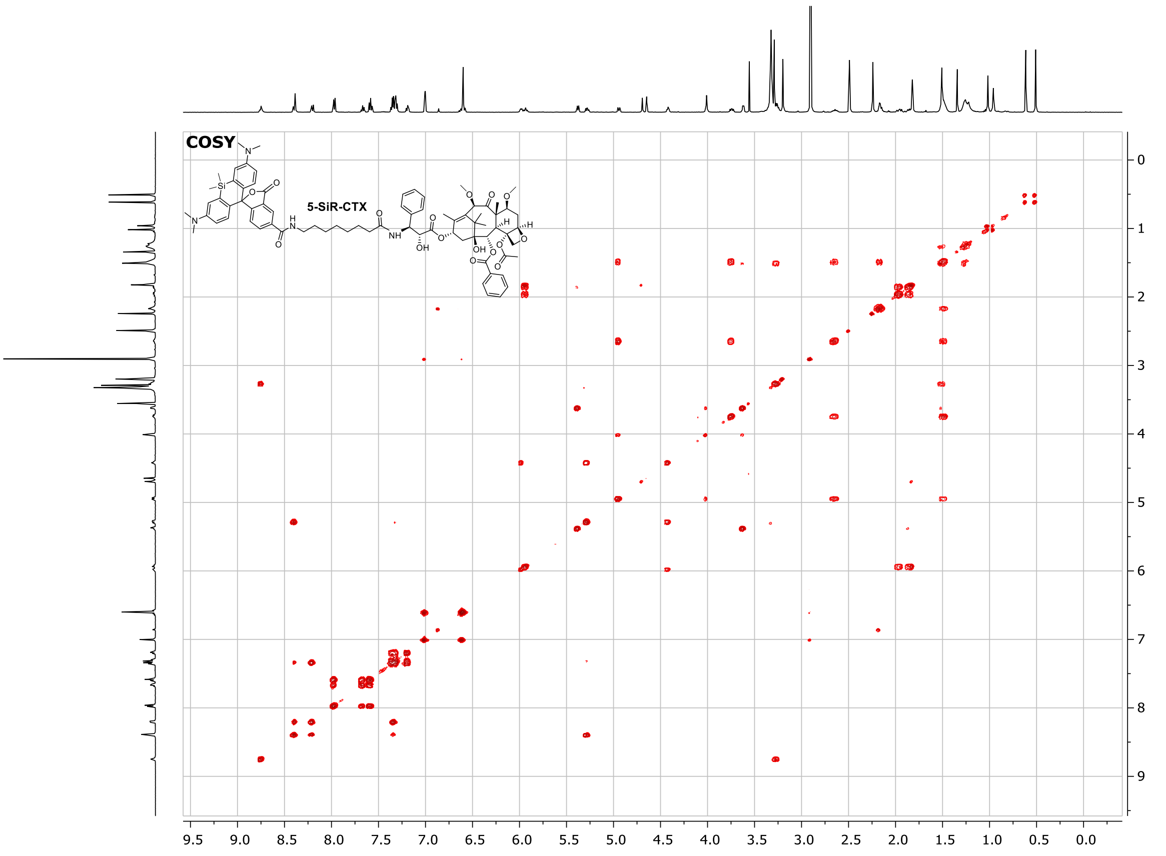

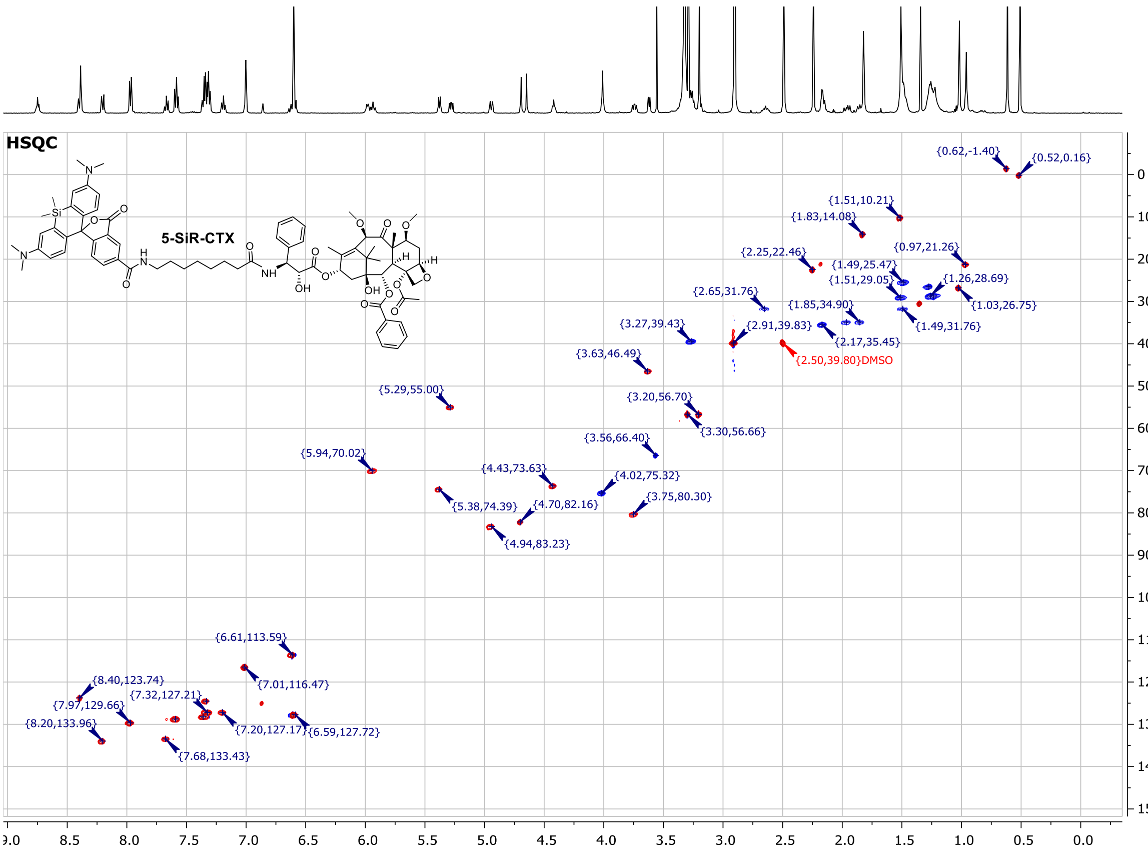

#### 4-610CP-C5-COOH (SI-5)

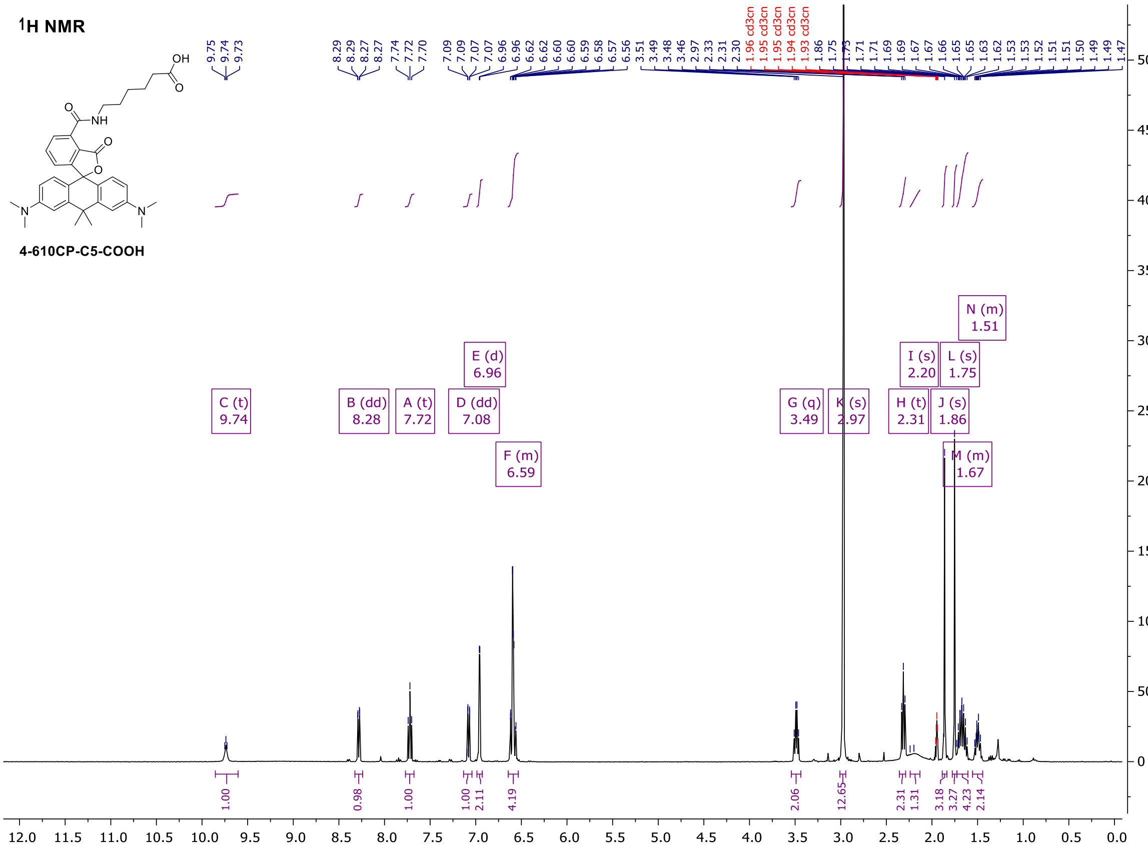

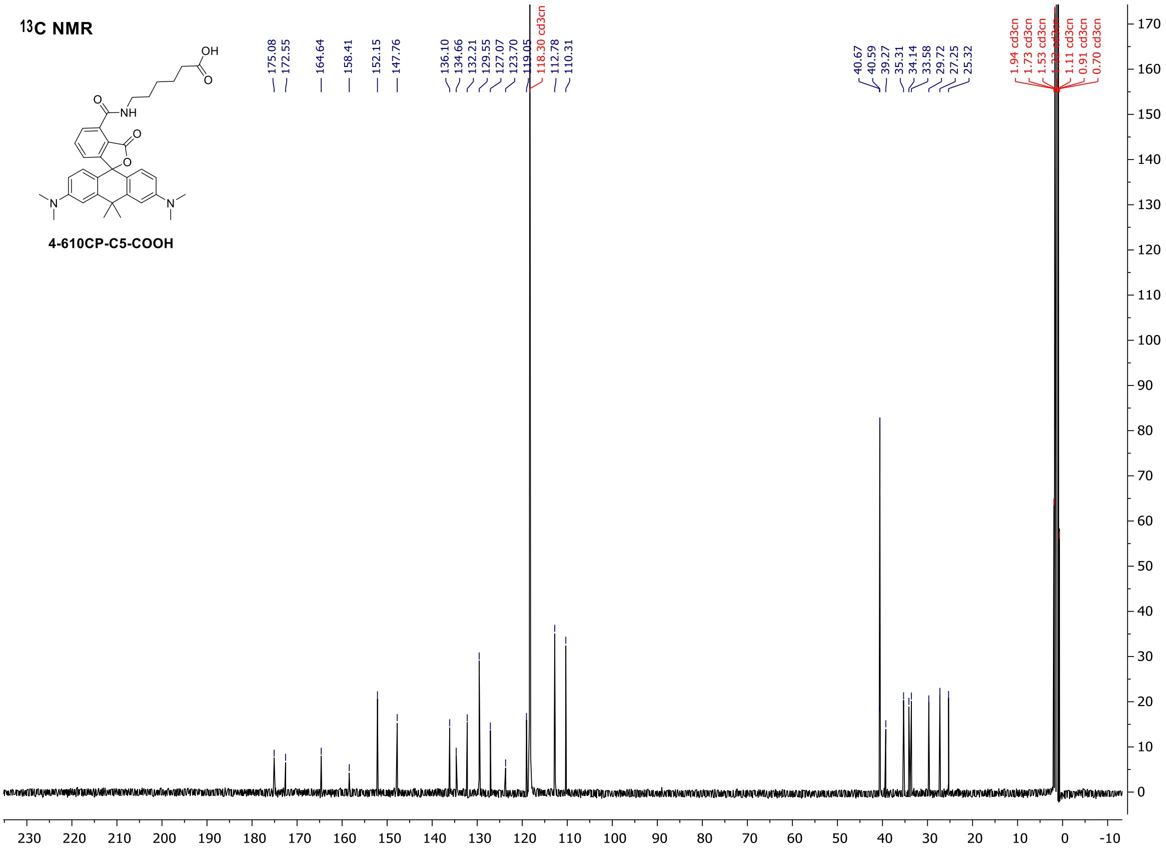

#### 4-TMR-Hoechst (34)

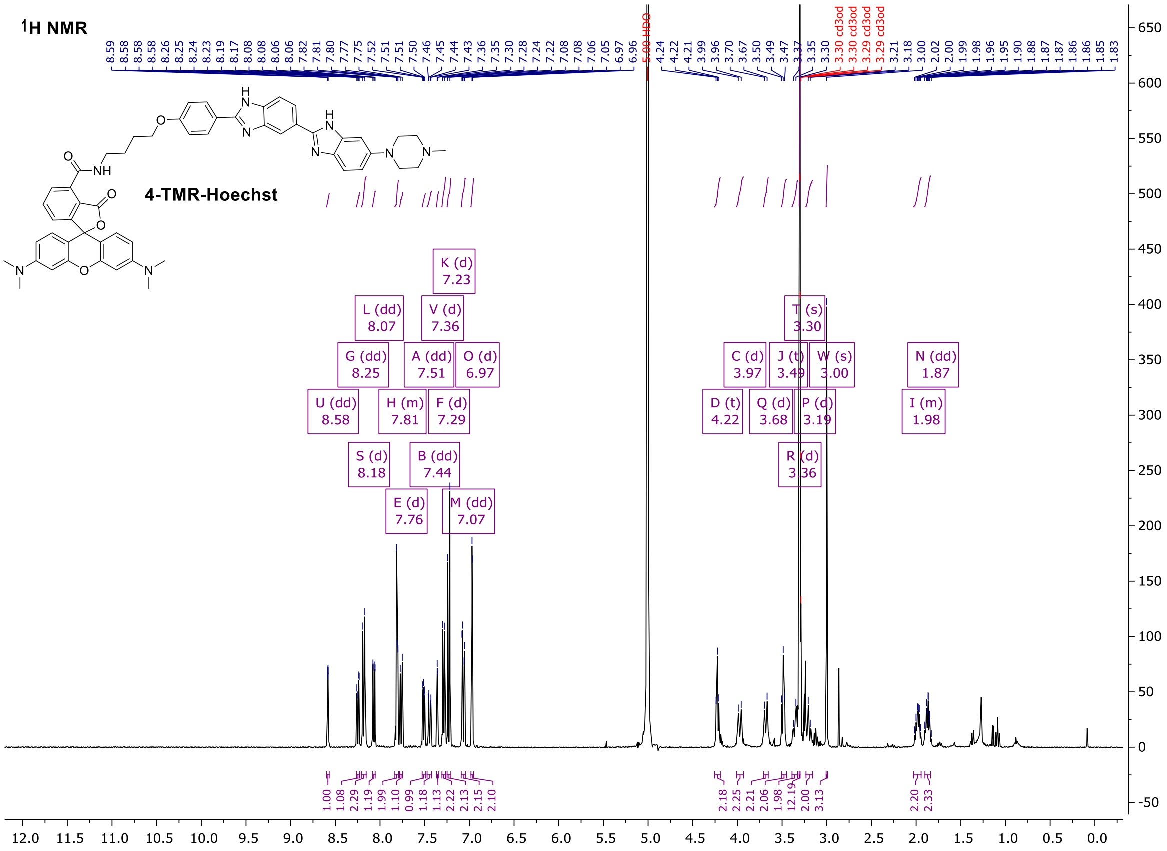

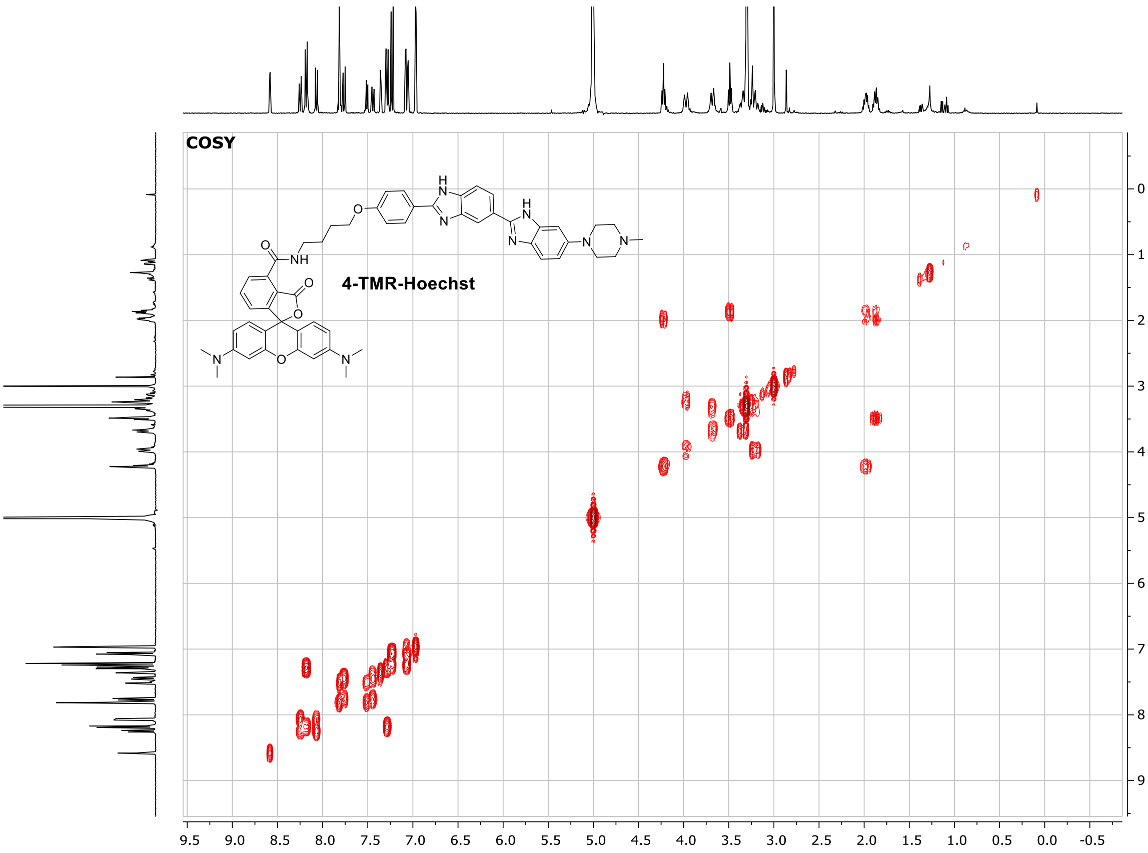
COSY
4-TMR-Hoechst

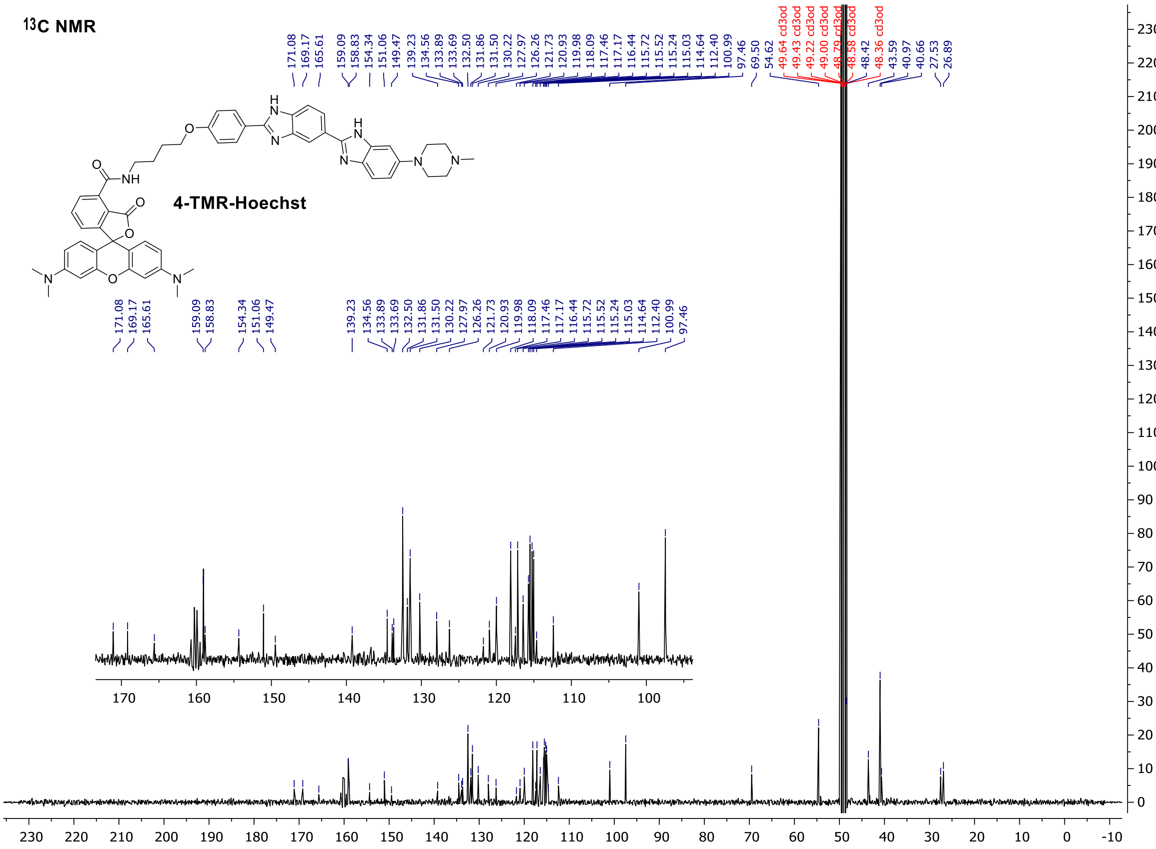
13C NMR
171.08 169.17 165.61 159.09 158.83 154.34 151.06 149.47 139.23 134.56 133.89 133.69 132.50 131.86 131.50 130.22 127.97 126.26 121.73 120.93 119.98 118.09 117.46 117.17 116.44 115.72 115.52 115.24 115.03 114.64 112.40 100.99 97.46 69.50 54.62 49.64 cd3od 49.43 cd3od 49.22 cd3od 49.00 cd3od 48.79 cd3od 48.58 cd3od 48.42 48.36 cd3od 43.59 40.97 40.66 27.53 26.89
4-TMR-Hoechst

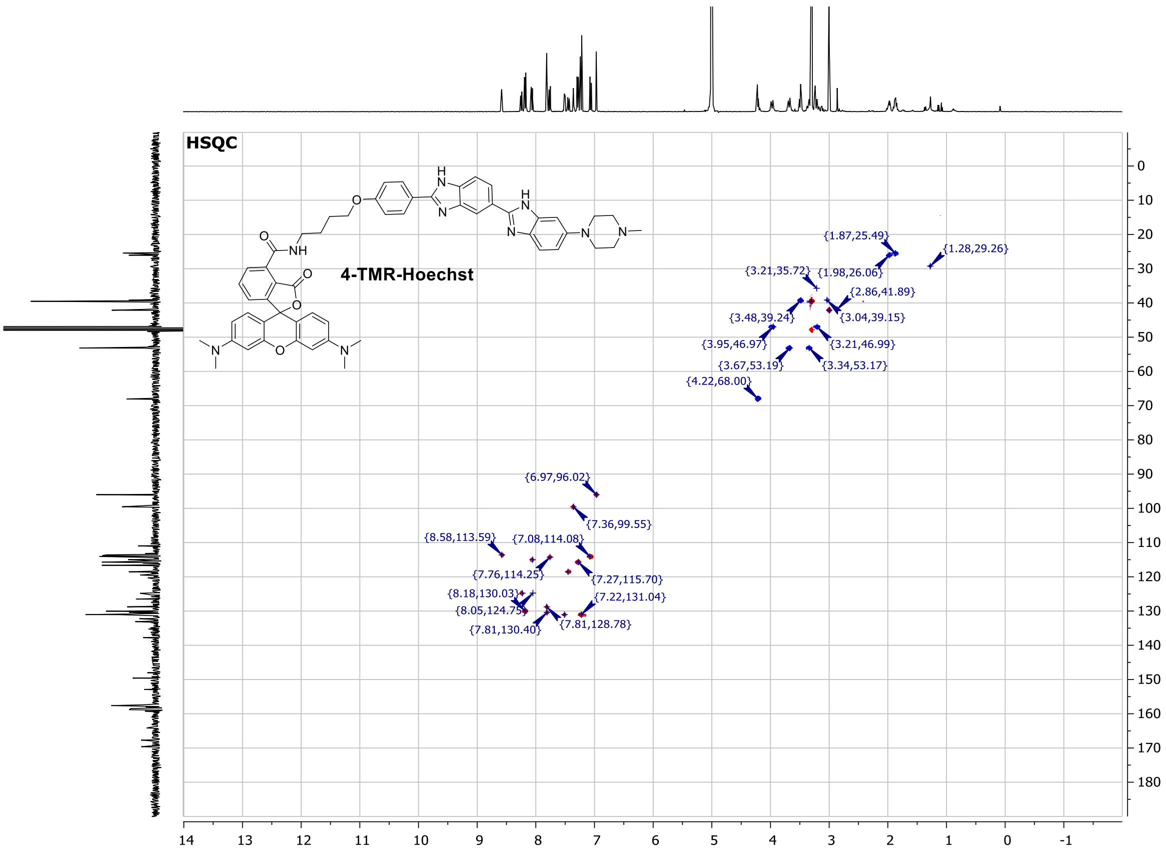

#### 4-580CP-Hoechst (37)

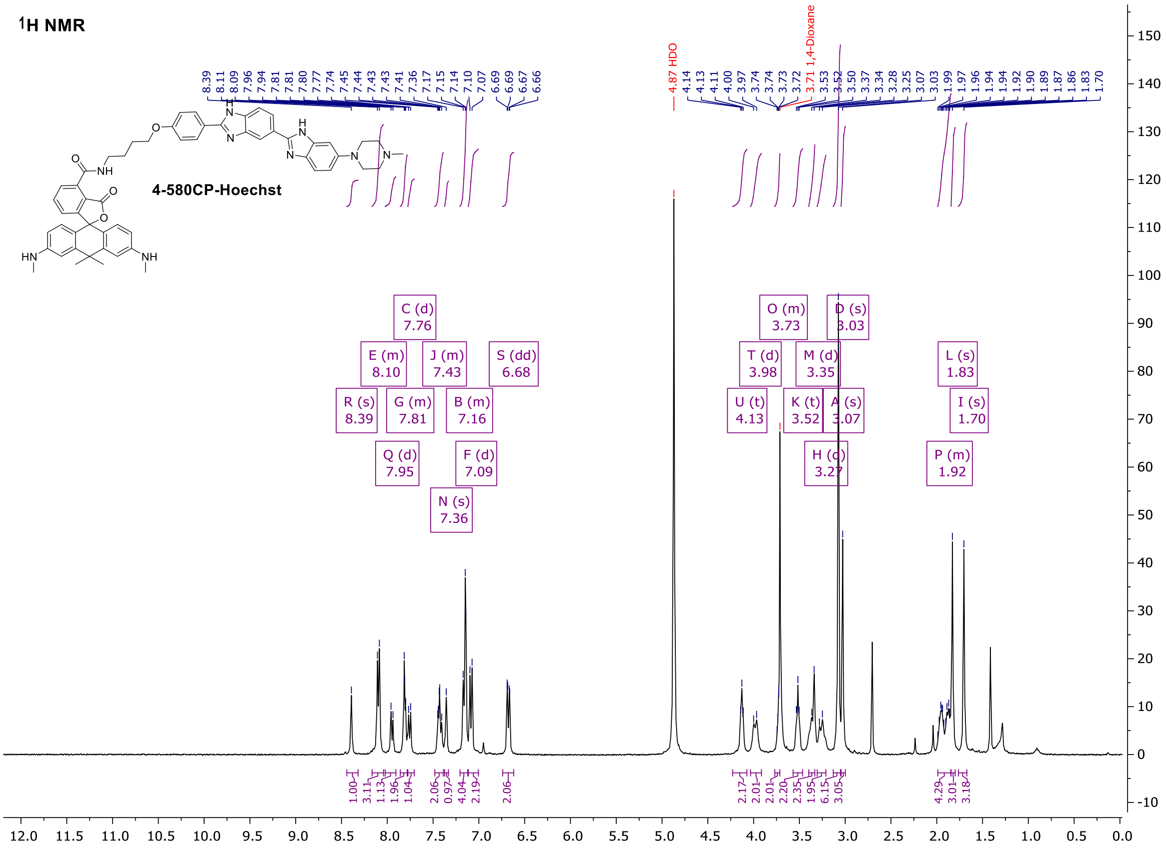

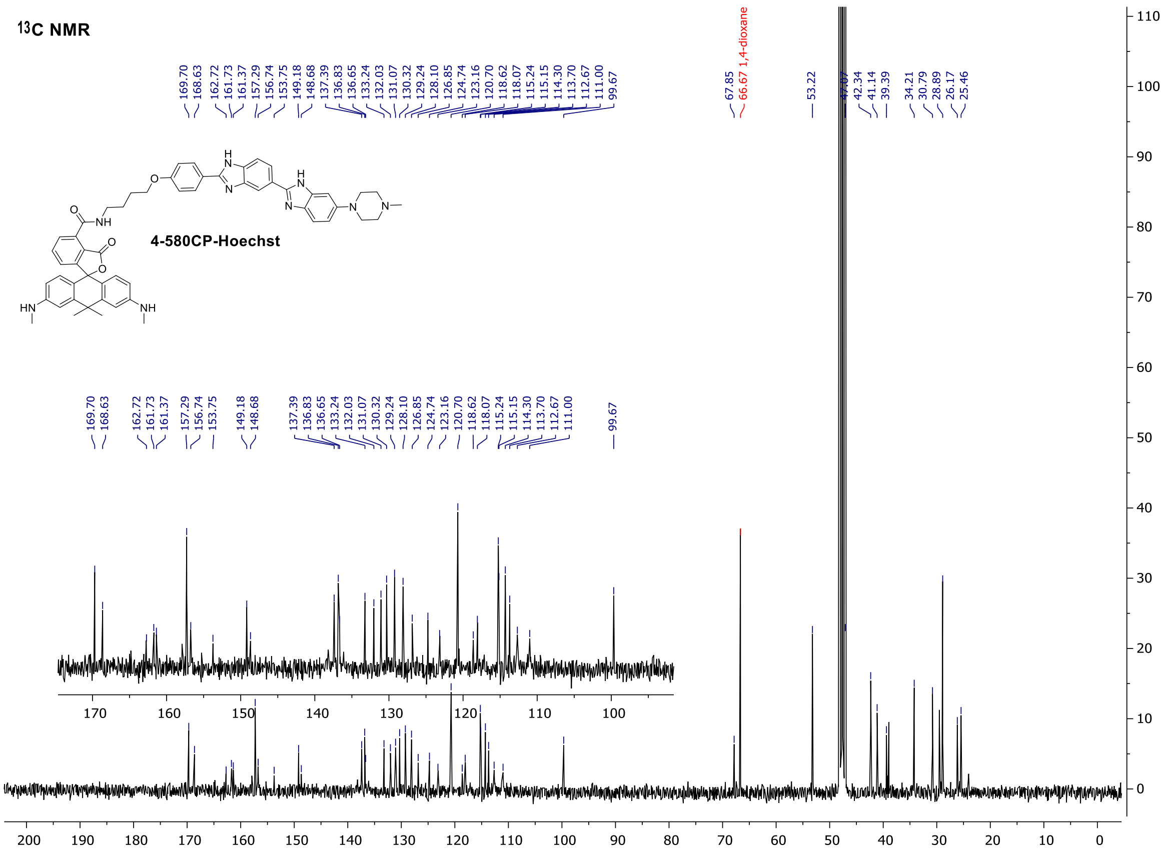
13C NMR
4-580CP-Hoechst
66.67 1,4-dioxane

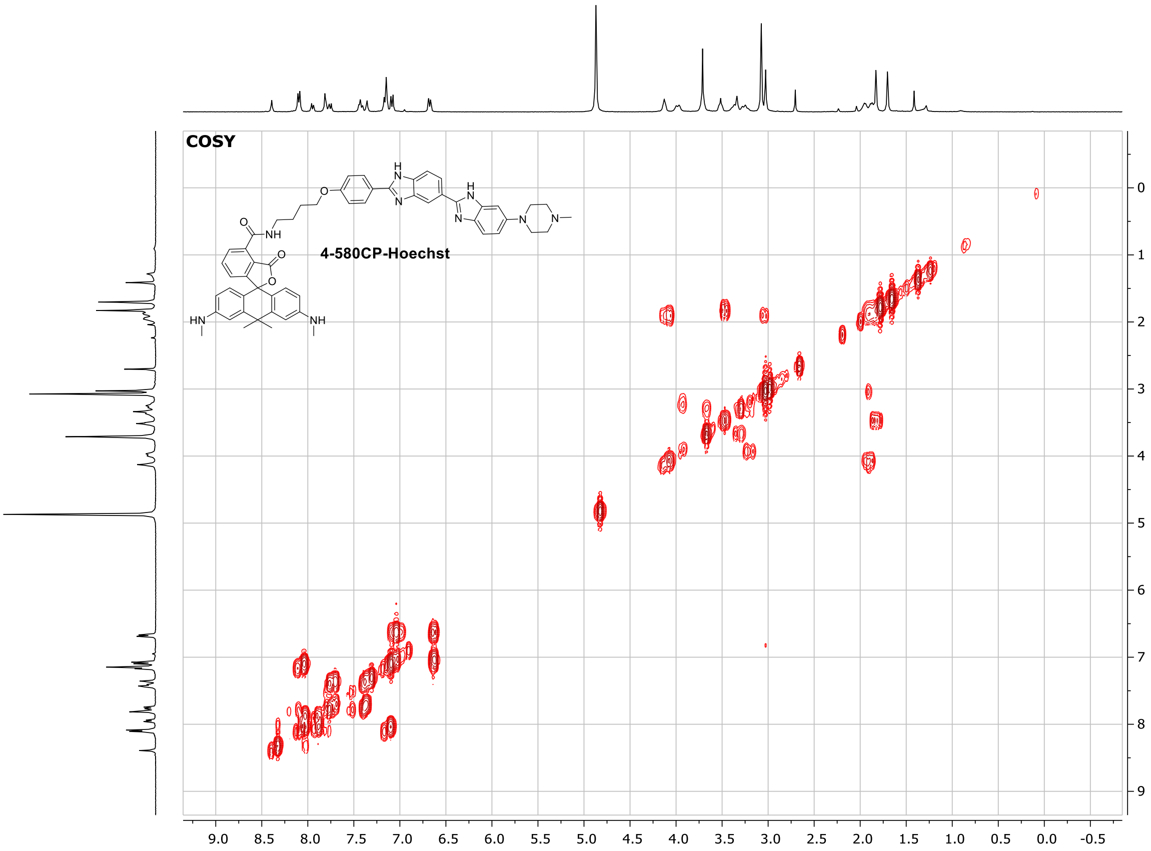
COSY
4-580CP-Hoechst

### 4-610CP-JASP (40)

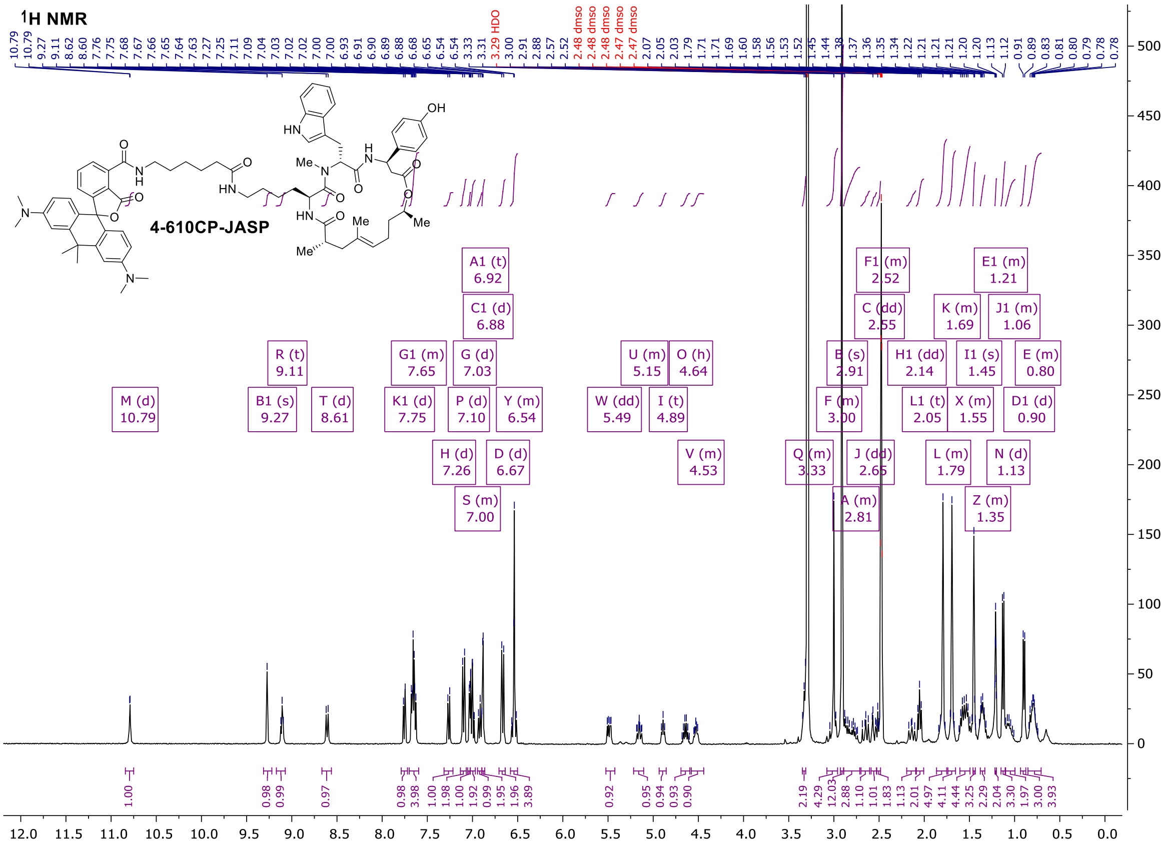

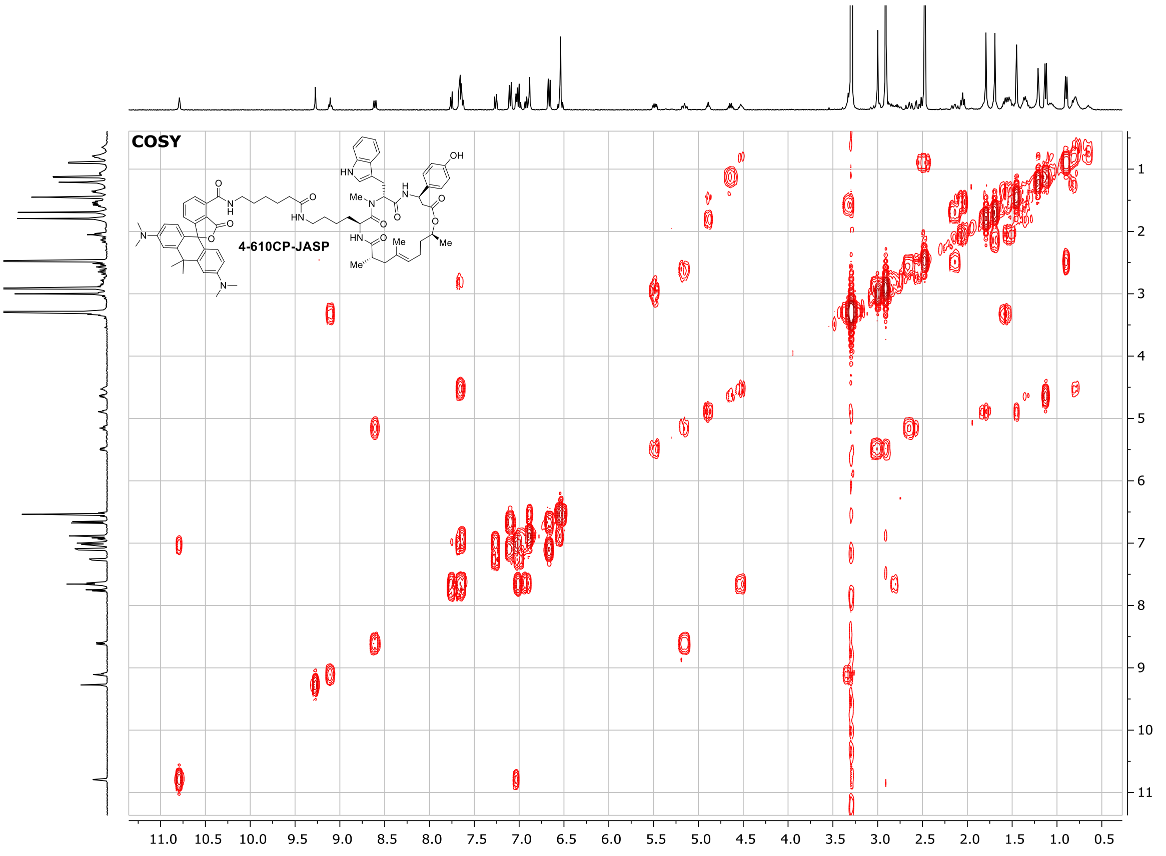
